# Temporal Notch signaling and Hes-mediated competitive de-repression regulate mucociliary cell fates in *Xenopus*

**DOI:** 10.1101/2023.02.15.528675

**Authors:** Magdalena Maria Brislinger-Engelhardt, Mona Hansen, Tim Litwin, Maximilian Haas, Aisha Andricek, Fabian Lorenz, Africa Temporal-Plo, Sarah Bowden, Sandra Hägele, Damian Weber, Alexia Tasca, Stefan Haug, Clemens Kreutz, Peter Walentek

**Affiliations:** Internal Medicine IV, Medical Center - University of Freiburg, Hugstetter Strasse 55, 79106 Freiburg, Germany; CIBSS Centre for Integrative Biological Signalling Studies, University of Freiburg, Schänzlestrasse 18, 79104 Freiburg, Germany; SGBM Spemann Graduate School for Biology and Medicine, University of Freiburg, Albertstrasse 19A, 79104 Freiburg, Germany; IMPRS-EBM International Max Planck Research School of Epigenetics, Biophysics, and Metabolism, Max Planck Institute of Immunobiology and Epigenetics, Stübeweg 51, 79108 Freiburg, Germany; Institute of Genetic Epidemiology, Medical Center - University of Freiburg, Hugstetter Strasse 55, 79106 Freiburg, Germany; IMBI Institute of Medical Biometry and Statistics, Institute of Medicine and Medical Center Freiburg, Stefan-Meier Strasse 26, 79104 Freiburg, Germany

**Keywords:** Xenopus, airway epithelium, development, multi-ciliated cells, mucus, basal cells, mathematical modeling

## Abstract

Mucociliary epithelia are found across different organs in animals, where they release bioactive substances and generate extracellular fluid flows. One key function of mucociliary epithelia is the clearance of pathogens, e.g. in the vertebrate lung and the epidermis of amphibian tadpoles. Mucociliary clearance relies on the correct balance between secretory cells that release mucus, ciliated cells that generate fluid flow as well as specialized cell types, including pH-regulating ionocytes and basal stem cells. Notch signaling, Hes repressors and cell type-inducing transcription factors (e.g. Foxi1, Mcidas, Spdef and Tp63) regulate cell fates and cell type proportions across mucociliary systems. Lateral inhibition was proposed to control mucociliary cell fates, but current models cannot explain how more than two cell types are generated and how Hes genes are employed as mediators during patterning. Using the *Xenopus* tadpole epidermis, we addressed these open questions in mucociliary biology through a combination of *in vivo* and organoid experiments, time-resolved transcriptomic studies and mathematical modeling. This revealed that ionocytes, ciliated cells, secretory cells and basal cells are preferentially specified at different time points and Notch signaling levels via sequentially expressed Hes factors. We termed this mode of patterning “competitive de-repression”, because cell fates are selected by suppression of alternative fate choices, and demonstrate that this relies on differential active repression of cell fate transcription factors. Mathematical modeling further indicated the need for a positively Notch-regulated patterning factor, and we provide evidence that Spdef mediates Notch input for secretory and basal cell specification. Collectively, this work presents a coherent model for Notch- and Hes-mediated mucociliary cell fate specification in a vertebrate tissue, which allows for the specification of more than two cell fates within the Notch lateral-inhibition paradigm.

## Introduction

Mucociliary epithelia line various tissues, including the mammalian respiratory tract, where they act as barrier against infections^1,2^. This is achieved through mucociliary clearance, a process in which secretory cells release mucus to trap inhaled pathogens and ciliated cells generate a directional fluid flow to remove mucus from the lung. This clearance function relies on the proper balance between the numbers of ciliated, secretory and additional cells in the tissue^3–5^.

The *Xenopus* embryonic mucociliary epidermis is an *in vivo* model for airway-like mucociliary epithelia^6^. Epidermis development is initiated by maternally deposited *sox3* and *foxi2*, which induce ectoderm formation at zygotic genome activation (ZGA)^7^. Subsequently, ectodermal progenitors with differential potential emerge in the outer epithelial cell layer as compared to deep cell layers, which is guided by maternally deposited signaling factors that are asymmetrically sorted along the apical-basal axis of dividing blastula cells^8,9^. Outer cells develop into goblet-like cells (goblet), while *foxi1*(+) multipotent mucociliary progenitors (MPPs) emerge among deep cell layers^10^ (**Fig. 1A**). During specification, MPPs commit to four different cell types: Ionocytes (ISCs), multi-ciliated cells (MCCs), small secretory cells (SSCs), and basal cells (BCs)^11,12^. After specification, MCCs actively migrate to generate an even MCC spacing pattern, which is not observed for ISCs and SSCs^13,14^. MCCs, ISCs and SSCs intercalate into the epithelium, while BCs position at the base and serve as tissue-specific stem cells^13,15–18^ (**Fig. S1A,B**). The proportions of mucociliary cell types are stable in the trunk region of the embryo, with 2/3 of goblet cells from the outer layer and 1/3 of intercalating cells produced at approximately equal numbers^11^. The patterning process is accompanied by morphogenetic changes from a multi-layered into a bi-layered epithelium driven by epiboly during gastrulation and radial intercalation during epidermis development^13,19^ (**Fig. S1A**). Due to its simplicity and stable cell type proportions, the *Xenopus* epidermis model allows to address how stereotypic cell type compositions are generated in mucociliary epithelia.

**Figure 1:**
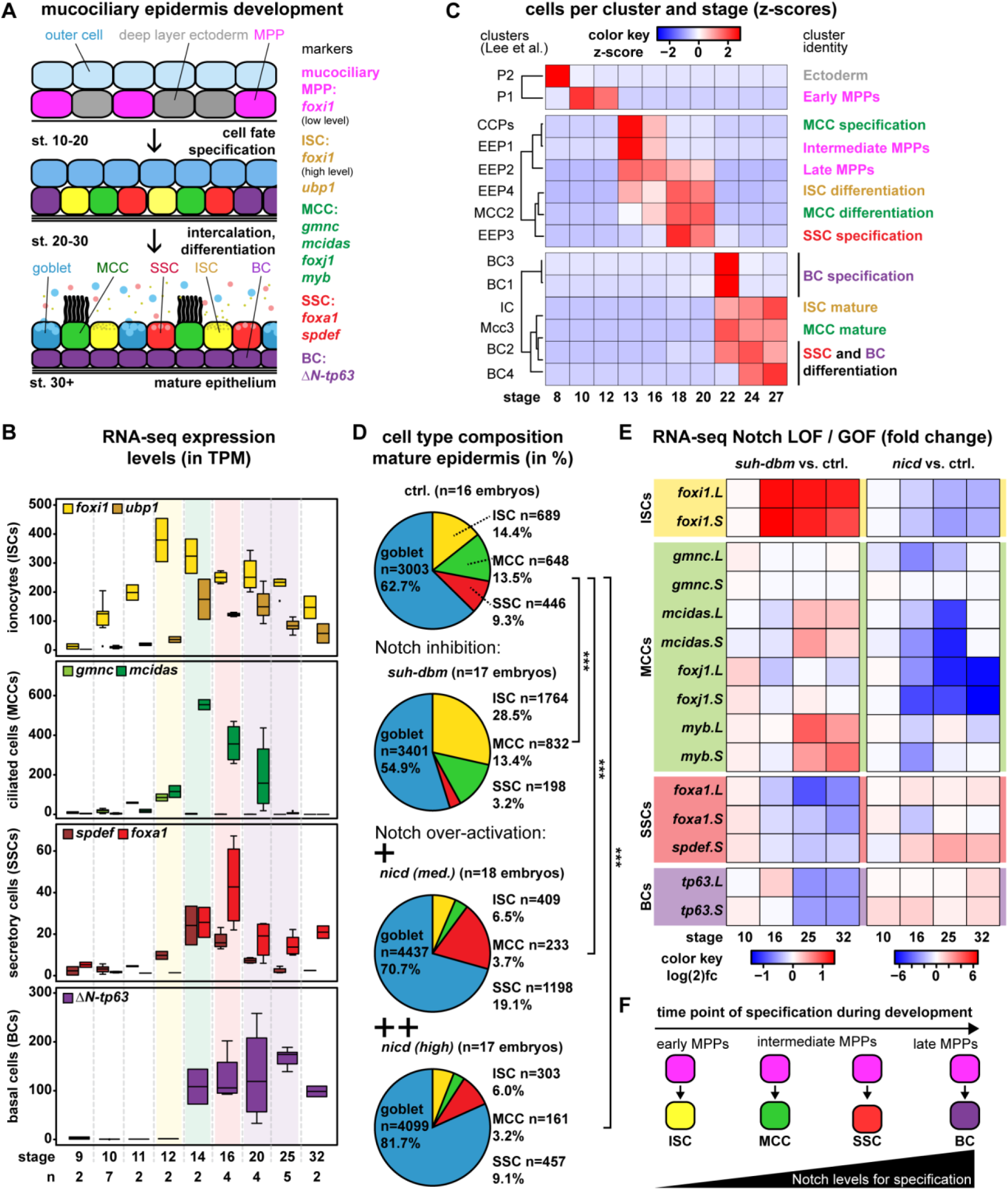
Cell fate specification and Notch signaling are temporally linked. **A:** Schematic representation of mucociliary epidermal development and cell types. **B:** Expression levels of indicated genes in unmanipulated mucociliary organoids across different stages of development (bulk RNA-seq). **C:** Cell numbers per cluster across different stages of development (single-cell RNA-seq from Lee et al.). **D:** Quantification of cell types after Notch manipulations and IF at st. 32. Representative examples are shown in Fig. S4B. X-test, *** = p < 0.001. **E:** Changes in cell type marker expression relative to stage-matched controls after manipulation of Notch signaling in mucociliary organoids (bulk RNA-seq). **F:** Schematic representation of correlations between MPPs, Notch levels and cell fates during developmental patterning.

Cell fates are conferred by evolutionarily conserved transcription factors when these reach critical levels within the cell (**Fig. 1A**): ISCs are specified by high levels of Foxi1 that activate Ubp1 and Dmrt2 during subsequent ISC subtype differentiation, while MPPs express *foxi1* at lower levels^10,15^ (**Fig. S1C**). Due to positive auto-regulation of *foxi1* expression, ISCs represent a “default” cell fate in the epidermis, e.g. when MCC specification is blocked^10,20^. Mcidas and Gmnc specify MCCs by competing with the cell cycle regulator Gmnn for E2f protein binding in a concentration-dependent manner^21–23^ and induced downstream cilia motility genes including *foxj1*^24,25^. Foxa1 induces SSC fate in *Xenopus*^26^, and Spdef is a pro-secretory cell factor conserved across epithelia^27^. High levels of ΔN-tp63 induce BC specification and dominantly block ISC, MCC and SSC fates^17^.

Notch signaling controls cell type specification across mucociliary tissues, and lateral inhibition was suggested to regulate MCCs in *Xenopus* as well as to decide between MCC and secretory cell fates in the mammalian airways^28,29^. However, proposed models of lateral inhibition deployment fail to explain how more than two cell types can be generated in mucociliary systems. Furthermore, in *Xenopus* ISCs and MCCs are both generated upon Notch inhibition, while SSCs are induced by Notch activation^15,30^. Single-cell RNA-sequencing (scRNA-seq) studies also suggested non-binary cell fate decisions from progenitors in *Xenopus*^12^. Furthermore, multiple Hairy enhancer of split (Hes/Hey) proteins have been implicated as direct Notch targets as well as in mucociliary patterning, but how they regulate cell fate decisions remains unresolved^29,31–35^. Collectively, mounting evidence argues for an alternative mode of Notch-regulated cell fate specification in mucociliary epithelia.

Here, we used the *Xenopus* mucociliary epidermis and mathematical modeling to investigate how Notch signaling and Hes transcription factors are employed to generate a stereotypic cell type composition with correct proportions of specialized cells *in vivo*.

## Results

### Mucociliary cell types are preferentially specified during different time points of development

We have previously reported that SSCs are specified later than ISCs and MCCs^16^. This suggested that secretory cells requiring higher Notch levels are specified after low Notch-requiring cell types, similar to the mammalian airways^29^. We used “animal cap” explants that develop autonomously into mucociliary organoids to investigate gene expression dynamics over the course of development^11^. These organoids recapitulate stages of *in vivo* epidermis development^36,37^ and maintain similar cell type proportions, with the limitation that some morphological differences are observed (e.g. the internal localization of SSCs and delayed planar cell polarization)^37^ (**Fig. S2A-D**). We harvested organoids over time (st. 9 - 32) for bulk mRNA-sequencing (RNA-seq) and plotted expression levels (normalized to TPM) of cell fate regulators over developmental time to determine when different cell types are specified.

RNA-seq data revealed that cell fate regulators were induced and peaked sequentially, suggesting that the probabilities for MPPs to acquire different cell fates changed over time (**Fig. 1B**). Based on our understanding of mucociliary transcription factor functions, we concluded from these data, that the ISC specification rate peaks between st. 12 - 14, while the main phase of MCC specification is between st. 14 - 16, followed by SSC specification around st. 16, and BC induction at st. 20 - 25 (**Fig. 1B**). The sequential appearance of cell type markers was also observed in embryos using chromogenic whole-mount *in situ* hybridization (WMISH) and multi-color hybridization chain-reaction (HCR), with the limitation that due to the dual role of Foxi1 in MPPs and ISCs, it is hard to determine the exact time point of ISC fate acquisition at the individual cell level (**Fig. S3A,B**).

Next, we used published scRNA-seq data from *Xenopus* mucociliary organoids collected during different time points of development (st. 8 - 27)^12^. In this dataset, MCCs were the first cell type to clearly separate based on differentially expressed transcripts^12^, which could be in part driven by the expression of *foxi1* in MPPs and ISCs^10^. Across all stages, 15 cell clusters were designated in the study. We analyzed the number of cells assigned to different clusters over time to visualize when cells with different properties are enriched during development. In this analysis, we excluded the goblet cell cluster to focus on cell fate decisions in deep layer cells. This revealed enrichment of cell clusters at specific time points (**Fig. 1 C**): P2 cells were enriched at st. 8, while P1 cells were detected at st. 10 - 12. At st. 13 - 16, CCP and EEP1 cells were specifically enriched, in contrast to EEP2, MCC2 and EEP4 clusters, which were enriched between st. 13 - 20. EEP3 cells were enriched at st. 18 - 20, followed by BC1 and BC3 cells at st. 22. Lastly, IC, MCC3, BC2 and BC4 cells were enriched at st. 22 – 27. These data supported the notion that the likelihood for different cell states varied over time.

To interpret which cluster reflected which cell state, we analyzed marker expressing cell numbers per cluster (**Fig. 1 C, S4A**): The P2 cluster did not show enrichment for any mucociliary cell type marker used, likely representing a general ectoderm progenitor state, while P1 cells represent early MPPs/ISCs based on the core-ISC gene *gadd45g*^20^, which is expressed in MPPs and early ISCs, similar to *foxi1*^10^. EEP4 and IC clusters are enriched for cells expressing *foxi1* as well as ISC differentiation genes (*ca12*, *slc26a4* and *dmrt2*), indicating early differentiating and mature ISCs, respectively. CCPs, MCC2 and MCC3 cells represent different states in MCCs, with CCPs only expressing early specification genes (*mcidas* and *cdc20b*), MCC2 additionally expressing the differentiation marker *tekt2*, and MCC3 expressing only differentiation markers (*tekt2* and *dnah5*). EEP1 and EEP2 cluster cells, were moderately enriched in *spdef* expressing cells, but no other markers, suggesting that these are later stage MPPs, while EEP3 cells were more strongly enriched in *spdef* as well as in *foxa1* and *tp63*, indicating SSC specification. BC1 and BC3 cluster cells were most strongly enriched in *tp63*, in line with a state of BC specification. In contrast, BC2 and BC4 clusters were enriched for cells expressing *spdef*, *foxa1* and *tp63* most likely representing a mixture of differentiating SSCs and BCs. Together, the scRNA-seq data supported our interpretation of bulk RNA-seq data, including the notion that MCCs might not be per se specified before ISCs.

These data identify sequential phases of preferred cell fate acquisition from MPPs and suggest that early, intermediate and late sub-populations of MPPs exist.

### All mucociliary cell types are specified at distinct Notch levels

To elucidate if and how Notch signaling regulates each mucociliary cell type, we inhibited Notch using *dominant-negative RBPJ* (*suh-dbm*) or overactivated it using *Notch intracellular domain* (*nicd*) by mRNA overexpression^20^. In the mature epidermis (st. 32), Notch inhibition increased ISC and decreased SSC numbers, but did not increase MCC numbers evaluated by immuno-fluorescence (IF) (**Fig. 1D, S4B**). Notch activation suppressed ISCs and MCCs, expanded SSC numbers at intermediate (med.) levels, while suppressing them at high levels (**Fig. 1D, S4B**). Assessment of cell numbers and TP63(+) BCs by IF revealed an increase in BCs in response to high Notch activation, while overall cell numbers were not changed (**Fig. S4C**). This indicated that the loss of intercalating cell types was due to specification of BCs from MPPs instead.

To investigate Notch effects during different phases of epidermal development, we analyzed mucociliary cell type marker sets^12,17,20^ by RNA-seq in organoids. Notch inhibition strongly induced ISC markers and suppressed SSC as well as BC markers (**Fig. 1E, S5A,B**). MCC marker expression was reduced at st. 16, but recovered during later stages, indicating that MCC specification was shifted to later time points. This explains why final MCC numbers were not changed at st. 32 (**Fig. 1D**). In contrast, Notch overactivation suppressed ISC and MCC markers, and increased SSC and BC markers, confirming that *ΔN-tp63* expression and BCs are also Notch regulated (**Fig. 1E, S5A,B**).

Contrary to the current assumption^28,38^, these data argue that MCCs are specified at higher Notch levels than ISCs, but at lower levels than SSCs, and that BCs are generated when Notch levels are further increased.

### Emergence of ligand-expressing MPPs leads to an increase of Notch signaling during cell fate specification

The correlation between required Notch level and time point of peak specification rates suggested a temporal Notch-dependent mechanism controlling early, intermediate and late MPP subpopulations (**Fig. 1F**). Such temporal patterning mode would require the gradual increase in systemic Notch signaling levels during cell fate specification stages. To test this, we used quantitative RT-PCR (qPCR) on mucociliary organoids injected with Notch reporters (*4xcsl::H2B-mvenus*^39^ and *mouse-hes1::dsRed*^40^) that were harvested at different time points during development. In line with our hypothesis, we detected a systemic increase in Notch signaling between st. 9 - 16, followed by a decrease in signaling activity after cell fate specification stages (**Fig. 2A**). Such an increase could be driven by either homogenous increase of Notch across all cells or through an increase in cell numbers expressing Notch signaling components.

**Figure 2:**
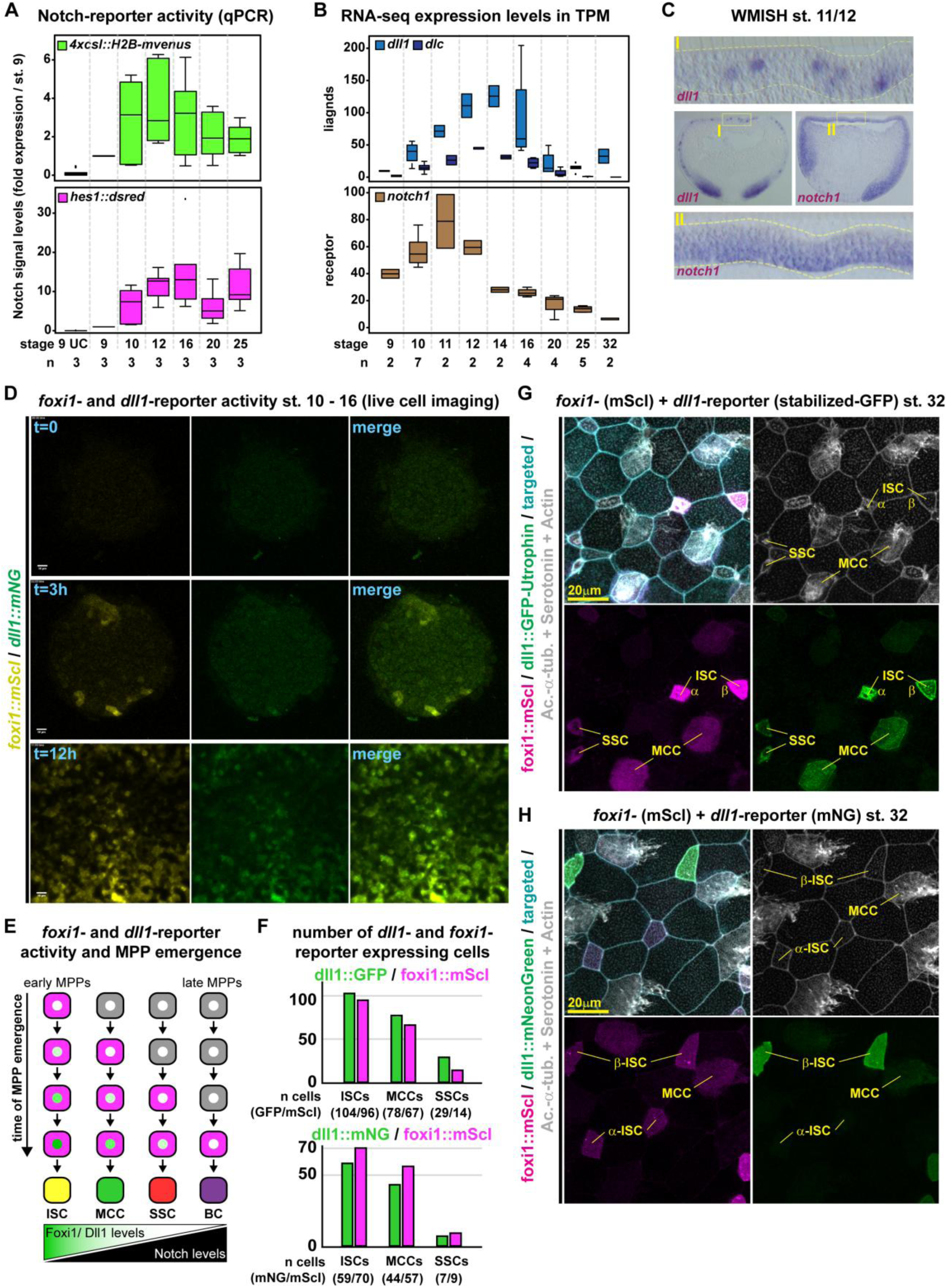
An increase in Dll1-expressing MPPs leads to increased Notch signaling during developmental patterning. **A:** Notch reporter activity in mucociliary organoids relative to reporter-injected st. 9 samples over time (qPCR). **B:** Expression levels of *dll1* and *notch1* in unmanipulated mucociliary organoids across different stages of development (bulk RNA-seq). **C:** Embryo sections and magnifications, WMISH for *dll1* and *notch1* at st. 11/12. Epidermis outlined in yellow. **D:** Stills from Movie1 depicting the increase in the number of *foxi1::mScarletI* and *dll1::mNeonGreen* cells during patterning stages in flat-mount mucociliary organoids imaged from the basal side. **E:** Schematic representation of potential reporter expression and cell fate acquisition in MPPs emerging at different time points during patterning. **F:** Number of *dll1::gfp-utrophin* or *dll1::mNeonGreen* expressing and *foxi1::mScarletI* expressing cells at st. 32. n = 7 embryos, for *dll1::gfp-utrophin*, and n = 5 embryos for *dll1::mNeonGreen*. **G,H:** Immunofluorescent images of epidermal cell types at st. 32. Acetylated-α-tubulin (Ac.-α-tub., grey) marks MCC cilia; Serotonin (grey) marks SSCs; F-actin (Actin, grey) marks cell borders and apical morphology; foxi1::mScarletI (magenta); dll1::gfp-utrophin or dll1::mNeonGreen (green). Representative samples used for reporter analysis in F. Cell types are annotated. Targeting was confirmed by co-injection of fluorescent membrane markers (cyan).

During epidermal development, MPPs express Notch ligands affecting cell fate decisions^10^. Cell fate specification terminates MPP ligand expression in differentiating ISCs, MCCs, SSCs and BCs^10,41^. Our RNA-seq data confirmed that *dll1* is the dominantly expressed ligand and that *notch1* is the dominantly expressed receptor during cell fate specification (**Fig. 2B, S6A**). WMISH revealed that *notch1* was homogenously expressed below the epithelial cell layer, suggesting that all precursors can respond to Notch signaling (**Fig. 2C**). In contrast, *dll1* expression was confined to interspersed MPPs, increased until st. 14 - 16 and was terminated at the end of cell fate specification by st. 20 (**Fig. 2B,C**). We next analyzed cell type composition by IF at st. 32 in control embryos and after morpholino-oligonucleotide (MO) knockdown of *dll1* or *notch1*. This uncovered a strong increase in ISCs, less pronounced effects on MCCs, and strong decrease in SSCs in MO-injected embryos, like Notch inhibition by mRNA overexpression (**Fig. 1D, S6B**). Co-injection of downstream-acting *nicd* reversed *dll1* MO and *notch1* MO effects (**Fig. S6B**). Hence, these knockdown and epistasis/rescue experiments demonstrate specificity of MOs and mRNAs as well as that Dll1 regulates cell fates via Notch1 in MPPs.

We have previously detected an increase in *foxi1*(+) MPPs during early patterning stages^10^. This suggested that an increasing number of ligand-expressing MPPs could drive the systemic increase in Notch signaling during cell fate specification stages. To investigate this possibility, we used a validated *foxi1*-reporter^10^ and a newly generated *dll1*-reporter that resembles endogenous *dll1* expression during patterning (**Fig. S6C,D**). We injected embryos with reporter-plasmids and imaged flat-mount epidermal explants from the basal side between st. 10 - 16 (**Movie 1**). This revealed that the number of cells with *foxi1*-reporter activity increased over time, not only through proliferation of reporter(+) cells, but also by activation of the reporter in additional deep layer cells (**Fig. 2D, Movie 1, Movie 2**). Furthermore, *dll1*-reporter expression shortly followed *foxi1*-reporter activation (**Fig. 2D, Movie 1, Movie 2**). Reporter(+) cells showed local mobility and frequently underwent a round of cell division, with both daughter cells maintaining similar levels of reporter activity (**Movie 2**). These data supported a model, in which an increasing number of ligand-expressing MPPs significantly contributes to the systemic increase in Notch signaling during cell fate specification stages.

MPPs emerging from epidermal precursors at early stages should be exposed to lower Notch levels than MPPs emerging at later time points, when “older” Dll1-expressing MPPs are already present. Since *foxi1* and *dll1* expression are regulated by negative Notch feedback^10,28^, early MPPs should express *foxi1* and *dll1* for a longer time, resulting in stronger reporter activation, and adopt earlier cell fates than late emerging MPPs (**Fig. 2E**). To test this, we injected *foxi1-* (*foxi1::mScarletI*) and *dll1-*reporter plasmids and investigated reporter levels in mature mucociliary cell types at st. 32 by IF. To trace *dll1*-expressing cells, we used reporter constructs containing a highly stable GFP-Utrophin fusion (*dll1::gfp-utrophin*) to identify cells even with low reporter activity, and compared the results to a construct containing a less stable mNeon-Green without Utrophin fusion (*dll1::mNeonGreen*) (**Fig. S6C**). As previously reported^10^, *foxi1-reporter* activity was detected most frequently in ISCs, frequently in MCCs and less frequently in SSCs (number of reporter expressing cells in **Fig. 2F**). The same was true for *dll1*-reporter expression using both constructs (**Fig. 2F-H, S7A,B**), in line with previous reports^42^. However, subtle differences between stable and less stable *dll1*-construct expression were observed. GFP-Utrophin was found in all mScarletI expressing ISCs, MCCs and SSCs as well as additional mScarletI(-) cells, indicating that the GFP reporter was more stable than foxi1::mScarletI. In contrast, mNeonGreen was more frequently found in mScarletI expressing ISCs than MCCs and SSCs (**Fig. 2F**). Additionally, we detected high *dll1*-reporter activities most frequently in ISCs, particularly β-ISCs (**Fig. 2G,H**), which were reported to be specified before α-ISCs that also require higher Notch activation during subtype selection^15^.

These experiments indicate that an increase in MPP numbers during cell fate specification stages leads to an increase in Notch ligand levels in the tissue and a progressive increase in Notch signaling levels over developmental time. Early emerging MPPs adopt ISC and MCC fates requiring low Notch levels, while later emerging MPPs are exposed to higher Notch levels leading to SSC and BC specification.

### Hes genes regulate cell fates by suppression of alternative fate choices

Next, we wondered how Notch-regulated *hes* genes could be employed for cell fate specification. We analyzed how all known *Xenopus hes*/*hey* genes react to Notch manipulations, and investigated their temporal expression dynamics in mucociliary organoids by RNA-seq (**Fig. S7C,D**). We found that *hes7* paralogs were expressed first, followed by *hes4* paralogs, and then *hes1/hes5* paralogs (**Fig. 3A, S7D**). Within the four paralog groups, *hes7.1.L (*the *hes7.1* gene expressed from long chromosome 3 of the allotetraploid frog^43^), *hes4.L/.S (*both alloalleles), and *hes5.10.L/.S (*both alloalleles) were the dominantly expressed Hes factors. *hes7.1* and *hes1* were inhibited by NICD (likely due to auto- or Notch-repression^44,45^), while *hes4* and *hes5.10* were induced by NICD, with *hes5.10* showing a stronger response (**Fig. S7C,E**). Next, we confirmed the expression of these *hes* genes by WMISH and found that *hes7.1* was first expressed and only in the deep layers, next *hes4* was increasingly expressed across cell layers followed by *hes5.10* (**Fig. 3B, S8A**). In contrast, *hes1* was expressed only weakly and in few cells across stages (**Fig. S7D, S8A,B**) arguing against a role in cell fate decisions in the epidermis, similar to findings from the mouse airways^33^. Hence, we identified a set of differentially Notch-regulated *hes* genes that are expressed sequentially during patterning.

**Figure 3:**
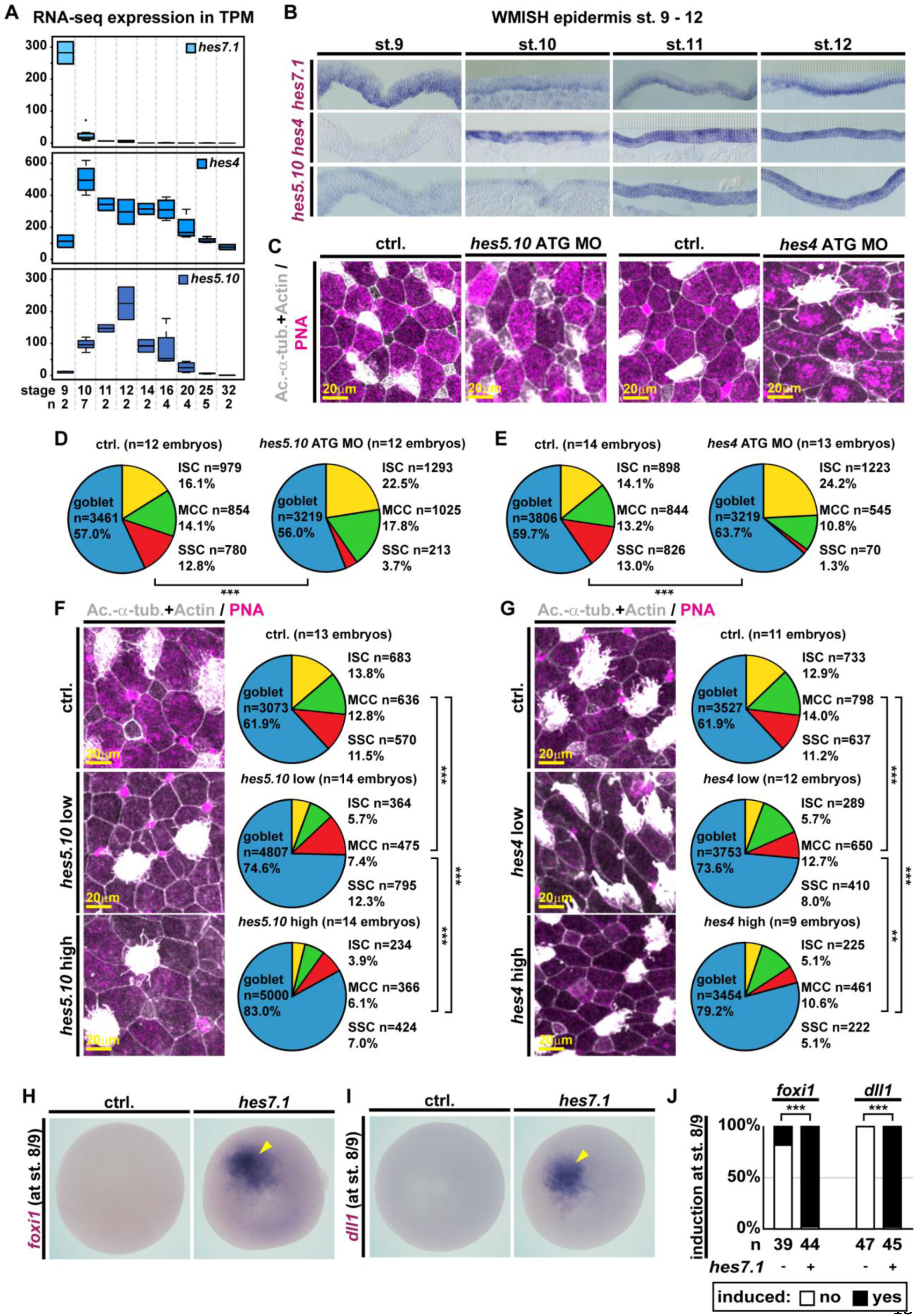
Sequentially expressed Hes genes regulate mucociliary development. **A:** Expression levels of indicated *hes* transcripts in unmanipulated mucociliary organoids across different stages of development (bulk RNA-seq). **B:** Epidermis sections, WMISH for indicated *hes* transcripts from st. 8 - 12. **C-G:** Immunofluorescent images of epidermal cell types at st. 32 (left) and quantification of results. Acetylated-α-tubulin (Ac.-α-tub., grey) marks MCC cilia; F-actin (Actin, grey) marks cell borders and apical morphology; peanut agglutinin (PNA, magenta) marks mucus in goblet cells and SSCs. MO-mediated knockdown of *hes5.10* (**C,D**) or *hes4* (**C,E**) using translation-blocking ATG MOs and quantification of cell types (**D,E**). Overexpression of low (15ng/ul) and high (50ng/ul) doses of hes5.10 (**F**) or of hes4 (**G**) and quantification of cell types (right). X-test, ** = p < 0.01; *** = p < 0.001. **H-J:** *Hes7.1* overexpression induces (yellow arrowhead) *foxi1* (**H,J**) and *dll1* **(I,J**) at st. 8/9. X-test, *** = p < 0.001.

We hypothesized that early expressed and negatively Notch-regulated Hes7.1 could be used to specify early cell types (ISCs and MCCs), while positively Notch-regulated Hes4 and Hes5.10 could be employed to specify late cell types (SSCs and BCs). In line with this, *hes5.10* (former name *esr6*) was previously shown to inhibit MCCs upon overexpression in *Xenopus*, but effects on other cell types or of other *hes* genes were not tested^28^. To test if endogenous Hes5.10 and Hes4 were required for the specification of SSCs and BCs, we knocked down *hes5.10* or *hes4* using MOs and analyzed cell type composition by IF at st. 32 (**Fig. 3C-E, S8C-E**). Knockdown of *hes5.10* using either a translation- (ATG MO) or a splice-blocking MO (SPL MO) increased the number of ISCs and MCCs, while reducing the number of SSCs (**Fig. 3C,D, S8C,E**). Knockdown of *hes4* using ATG or SPL MOs increased only the number of ISCs, MCCs were slightly reduced and SSCs were strongly reduced (**Fig. 3C,E, S8D,E**). The total number of epithelial cells or of basal Tp63(+) cells was not significantly changed in these morphants (**Fig. S9A**).

To test how different levels of Hes4 and Hes5.10 affect cell fate specification in the epidermis, we next used low (15ng/ul) and high (50ng/ul) concentrations of mRNAs for overexpression and investigated cell type composition by IF (st. 32). This showed that ISCs and MCCs were suppressed by low and high levels of *hes5.10*, while only high levels of *hes5.10* also efficiently inhibited SSCs (**Fig. 3F, S9B**). Additionally, the overall number of cells and particularly the number of Tp63(+) BCs were increased (**Fig. S9A**). These results were similar to the concentration-dependent effects of *nicd* (**Fig. S4B,C**) and suggested that Hes5.10 mediates medium and high Notch levels required for SSC and BC specification. Overexpression of *hes4* at low or high concentrations decreased ISC numbers, while SSC numbers were more reduced at high *hes4* concentrations (**Fig. 3G, S9C**). In contrast, MCC numbers were only mildly affected, even at high concentrations of *hes4* overexpression. Interestingly, the number of Tp63(+) BCs was strongly decreased, while the reduction in overall cell numbers was less pronounced (**Fig. S9A**). This indicated that Hes4 promotes MCC specification by inhibiting ISC, SSC and BC fates more efficiently.

To validate that these effects were driven by changes in cell fate specification rather than secondary loss of cells, we used ATG MO injections to knock down *hes5.10* or *hes4* on one side of the embryo and investigated cell type marker expression by WMISH at the end of cell fate specification (st. 17/18). Injection of *hes5.10* MO caused an increase of *foxi1* and in a subset of embryos also of *mcidas*, but *foxa1* was lost on the injected side of the embryo (**Fig. S10A,B**). Knockdown of *hes4* reduced *mcidas* and *foxa1* expression, while *foxi1* expression was increased (**Fig. S10A,B**). In some *hes4* and *hes5.10* morphants *ΔN-tp63* expression was also affected, but the effects were ambiguous and weak. These data confirmed that Hes knockdown affected decision making during cell fate specification.

Overexpression of *hes5.10* at both concentrations decreased *foxi1*, *foxj1* and *foxa1* expression on the injected side of the embryo, while *ΔN-tp63* expression was increased (**Fig. S10C,D**). This was in line with the st. 32 IF data and indicated that a moderate increase in Hes5.10 has delayed the specification of SSCs, similar to the delay in MCC specification after strong Notch inhibition. Overexpression of *hes4* at low or high concentrations reduced *foxi1*, *foxa1* and *ΔN-tp63* expression on the injected side, while *foxj1* was overall increased (**Fig. S10C,D**). In contrast to our initial hypothesis, these results indicated that sequential Hes4 and Hes5.10 expression could already regulate specification of the four mucociliary cell types: First, when Hes4 and Hes5.10 are low, ISCs are specified, because they are a “default” cell type in the epidermis^10^. Next, MCCs are specified when Hes4 is high and Hes5.10 is low. Finally, Hes4 levels decrease and SSCs are specified at lower Hes5.10 levels, while BCs are specified when Hes5.10 levels are high.

We therefore wondered which other role Hes7.1 might play, when it is not required for Notch-mediated cell fate decisions? Since it is only transiently expressed at ZGA and downregulated by st. 10 when patterning begins, we speculated that it might be involved in the regulation of the starting point of mucociliary development. To test this, we overexpressed *hes7.1* and analyzed *foxi1* and *dll1* expression at ZGA (st. 8/9) by WMISH. Indeed, this led to activation of both genes, indicating premature MPP generation (**Fig. 3H-J**). In contrast, overexpression of *hes7.1* and analysis of cell type markers by WMISH at the end of patterning as well as by IF in the mature mucociliary epidermis revealed a downregulation of cell type markers (**Fig. S10D,E, S11A**). Furthermore, the numbers of total cells and Tp63(+) BCs were reduced (**Fig. S11B**). Knockdown of *hes7.1* by MO injections increased *foxi1* expression at the end of patterning, while *mcidas*, *foxa1* and *ΔN-tp63* expression were reduced, suggesting that patterning was delayed (**Fig. S10B,E**). Analysis of cell types by IF in the mature epidermis further revealed an expansion of ISC and MCC numbers, while SSCs were decreased (**Fig. S11C**). This resembled a phenotype observed after knockdown of *ΔN-tp63*, where retention of MPPs for SSC specification is defective^17^. However, *hes7.1* MO did not lead to a similar loss of BCs (**Fig. S11B**). These results indicated a role for Hes7.1 in the regulation of transitions from early deep layer cells to MPPs, rather than Notch-mediated cell fate choices.

Together, this reveals that mucociliary cell fates are temporally regulated by the sequential expression of *hes4* and *hes5.10* genes and their ability to inhibit competing fate choices. Hence, cell fates are regulated by a “competitive de-repression” mechanism.

### Molecular basis of Hes-mediated competitive de-repression

To further elucidate the mechanism behind the observed Hes-mediated de-repression of cell fates, we first tested if Hes factors predominantly act through passive or active repression in the system^45^. Only active suppression by Hes factors requires the C-terminal WRPW-domain to bind TLE/Groucho, the actual repressor. We generated WRPW-deletion constructs, and overexpressed them at the same concentrations as full-length Hes-encoding mRNAs followed by IF at st. 32 to investigate cell type composition. Full-length *hes4* overexpression strongly inhibited ISCs while *hes4ΔWRPW* lost this ability (**Fig. 4A, S12A**). Similarly, overexpression of full-length *hes5.10* suppressed ISC and MCC formation while *hes5.10ΔWRPW* did not (**Fig. 4A, S12A**). This indicated that cell fate choices of ISCs, MCCs and SSCs required direct repression by positively Notch-regulated Hes factors during patterning.

**Figure 4:**
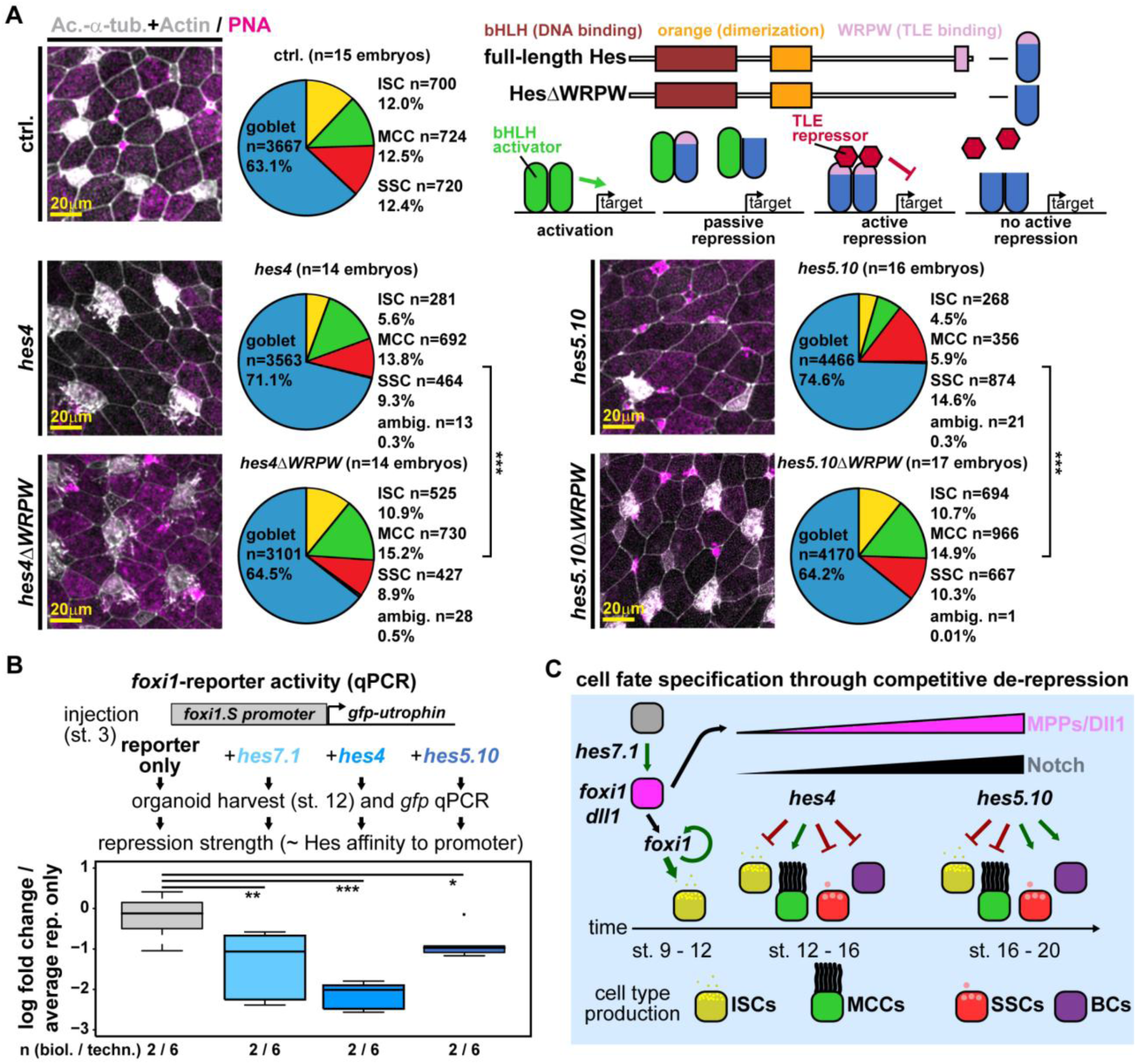
The molecular basis of Hes-mediated competitive de-repression. **A:** Immunofluorescent images of epidermal cell types at st. 32 (left) and quantification of results (right). Acetylated-α-tubulin (Ac.-α-tub., grey) marks MCC cilia; F-actin (Actin, grey) marks cell borders and apical morphology; peanut agglutinin (PNA, magenta) marks mucus in goblet cells and SSCs. Comparison of full-length *hes4* and *hes5.10* effects on cell types with effects after TLE/Groucho-binding domain deletion (*hes4ΔWRPW* and *hes5.10ΔWRPW*). The effects of active and passive repression by Hes factors are depicted schematically. X-test, *** = p < 0.001. **B:** Analysis of *foxi1::gfp-utrophin* reporter activity in reporter only or reporter and hes-mRNA co-injected organoids at st. 12 by qPCR. Mann-Whitney-test, * = p < 0.05; ** = p < 0.01; *** = p < 0.001. **C:** Schematic representation of cell fate specification through competitive de-repression of cell fates.

DNA binding in proximity to the regulated gene is a prerequisite for direct repression by Hes/TLE complexes^45^. The various Hes factors repressed cell types differentially when expressed at non-saturating levels, suggesting that they could operate through different binding affinities to promoters of cell fate regulators. To test this, we injected embryos with *foxi1*-reporter (*foxi1::GFP-utrophin*)^10^ alone or in combination with low (15ng/ul) concentrations of mRNAs encoding the Hes7.1, Hes4 and Hes5.10. Then, we generated organoids and assessed reporter activity by qPCR against *gfp* at st. 12. This revealed that Hes4 most strongly repressed reporter activity, while Hes7.1 and Hes5.10 had weaker effects (**Fig. 4B**). This was in line with the identified function of Hes4 as main repressor of ISCs during early patterning, required to facilitate MCC specification.

These results support the conclusion that differential binding affinities of Hes repressors to target gene promoters allow the selective, dose-dependent direct repression of fate choices during patterning. Since the number of ligand expressing cells increases over time, the probability for MPPs to be exposed to Notch ligands and to express Hes genes during cell fate decision making also increases with time. At the tissue level, these changes in probabilities are reflected in the changes of cell type production rates at different phased of mucociliary pattering (**Fig. 4C**).

### Mathematical modeling reveals the need for a positively Notch-regulated patterning factor

To test if Hes-mediated repression would be sufficient to generate stable proportions of mucociliary cell types, we built a dynamic mathematical model based on ordinary differential equations (ODE) using the Data2Dynamics environment^46^. Mathematical modeling allows us to extract information from systemic features (quantitative bulk RNA-seq data from organoids) and link them to the quantifications of cell type proportions from IF datasets to elucidate how dynamic gene expression leads to cell fate decisions. For this purpose, the temporal resolution of RNA-seq data has been further improved by fine-staging organoids (**Fig. S13A, Table S1**) and used as training data for the model.

Based on our observations (Movie1 and 2), this model assumes a well-mixed cell population, where deep layer cells, MPPs and specified cells constantly interact locally with their neighbors. The model defines cells, RNA levels and protein levels as functionally different model states: The combination of cells and their states determines the ligand production and translates into the system’s Notch level. Different Notch levels activate or inhibit the expression of genes, and the genes translate into their respective protein states that impact cell specification rates. During patterning stages, MPP, ISC, MCC, SSC and BC number estimates were based on the mRNA expression of markers. With our model structure defined, we were able to calibrate the a priori unknown model parameters (**Fig. 5A**) using time-resolved RNA-seq and st. 32 IF data via maximum likelihood estimation.

**Figure 5:**
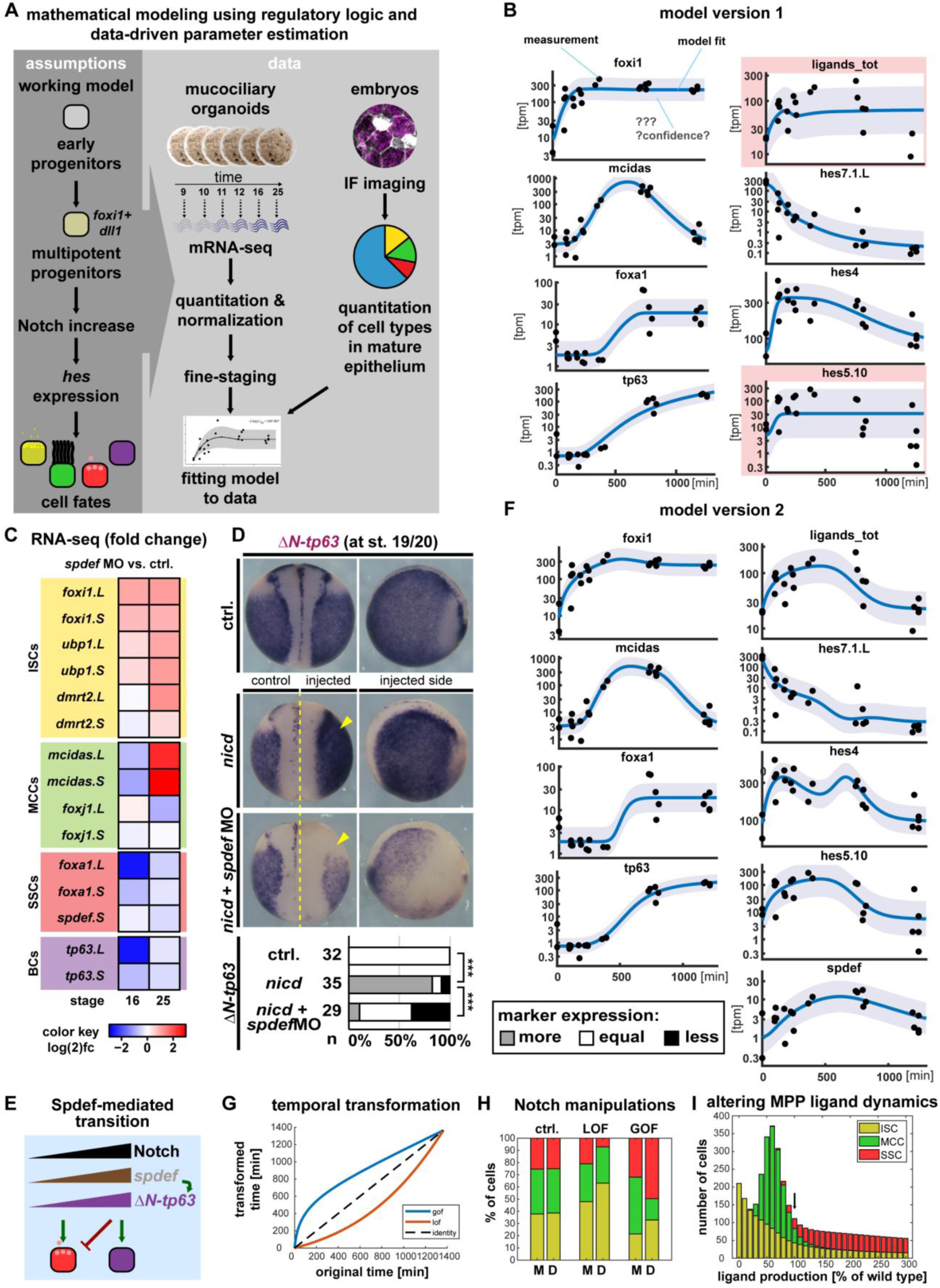
Mathematical modeling of patterning and Spdef functions. **A:** Schematic representation of modeling approach. **B:** Model 1 Visualization of fit (blue line) to data from RNA-seq (black dots). Light blue area represents the error model that describes the measurement noise. Please note that model 1 failed to correctly fit ligands_tot and hes5.10 levels (highlighted by red boxes). **C:** Changes in cell type marker expression relative to stage-matched controls after knockdown of *spdef* in mucociliary organoids (bulk RNA-seq). **D:** WMISH for *ΔN-tp63* transcripts in control and unilaterally *nicd* (100ng/ul) or *nicd* = *spdef* MO injected st. 19/20 embryos. Left panels, dorsal views, manipulated side is indicated. Right panels, lateral view on manipulated side of the embryo. Graph depicts quantification of results. Marker expression was assessed relative to the uninjected control side and is shown as more, equal or less than the control side. X-test, *** = p < 0.001. **E:** Schematic representation of Spdef functions during patterning. **F:** Model 2 Visualization of fit (blue line) to data from RNA-seq (black dots). Light blue area represents the error model that describes the measurement noise. **G:** The *in-silico* model could explain Notch loss- (LOF) and gain-of-function (GOF) experiments by deceleration (LOF) or acceleration (GOF) of the whole cell specification process via the depicted nonlinear transformation of the time axis. **H:** Comparison of modeled [M] and experimentally determined [D] cell type ratios of ISCs (yellow), MCCs (green) and SSCs (red). **I:** *In silico* manipulation of ligand expression levels from differentiating multipotent progenitors using model 2, 100% = normal control levels of ligands, manipulations in 10% steps, for negative (reduced, left) and positive (increased, right) manipulations of levels.

The model dynamics are initiated by a stable rate of MPP production that then undergo cell fate decisions in response to Notch signaling via Hes input (**Fig. 5A, S12B**):

1. MPPs (*foxi1* and *dll1*) undergo a transition into ISCs (*foxi1* only) when Notch signaling is low and only Hes7.1 is active.
2. When Notch levels (Notch ligand expression) increase, *hes7.1* is inhibited and *hes4* is activated. Hes4 represses ISCs and SSCs (*foxa1*) resulting in MCC (*mcidas*) specification.
3. When Notch levels further increase, *hes4* is less active and *hes5.10* gets increasingly activated. Hes5.10 represses ISCs and MCCs allowing first for SSCs to be specified and then for BC (*ΔN-tp63*) specification at high Hes5.10 levels.
4. The process is simulated until st. 25, when cell type proportions have been finally decided.

To increase model stability, competitive de-repression was phenomenologically simplified in the model as a combination of inhibition and activation of cell type specification genes by Hes factors (**Fig. S12B**).

The resulting model found a solution (model 1) that recapitulated the sequential specification of cell types (**Fig. S13B**), but failed to fit ligand and *hes5.10* levels (**Fig. 5B**). This indicated, that the model could not coherently describe how *ΔN-tp63* levels can remain high when Notch levels decrease, because the model required constant high Notch input for BC maintenance. Furthermore, model 1 failed to predict cell type production phases for Notch gain- and loss-of-function experiments or the effect of altering ligand levels expressed from MPPs (**Fig. S13C,D**). Together, this suggested that Hes-mediated repression alone was unlikely sufficient to regulate the patterning process.

The Ets-family transcription factor Spdef is Notch-regulated, required for secretory cells in the mammalian airways^27,47^ and is expressed in human airway basal cells (Human Protein Atlas)^48^. In *Xenopus*, *spdef* expression precedes *foxa1* as well as *ΔN-tp63*, and *spdef* is expressed in intermediate and late MPPs (**Fig. 1B, S4A**). Hence, Spdef could promote SSC-specification at lower levels, while promoting BC at higher levels. This would allow the system to terminate developmental patterning at peak Notch levels and transition towards homeostatic maintenance at lower Notch levels after st. 16 (**Fig. 2A,B**). We tested this hypothesis by *spdef* knockdown using MO injections followed by RNA-seq on control and manipulated mucociliary organoids at st. 16 and 25 (**Fig. 5C, S13E**). This revealed that loss of Spdef caused an upregulation of ISC markers (*foxi1*, *ubp1* and *dmrt2*), a loss of MCC markers at st. 16 followed by a recovery at st. 25 (*mcidas* and *foxj1*), and a strong decrease in SSC and BC markers (*foxa1*, *spdef*, *tp63*). These results resembled Notch inhibition in organoids (**Fig. 1E**) and supported a role for Spdef as positive mediator of Notch signaling input on cell fate decisions. Next, we validated that Spdef is required for Notch-mediated activation of *ΔN-tp63* expression in embryos. For that, we injected *nicd* mRNA to stimulate Notch on one side of the embryo with or without co-injection of *spdef* MO, and investigated the effects on epidermal *ΔN-tp63* by WMISH at the end of specification. This confirmed a strong upregulation of *ΔN-tp63* upon Notch activation, which was lost when *spdef* was simultaneously knocked down (**Fig. 5D**). Furthermore, overexpression of *spdef* mRNA increased *ΔN-tp63* expression and decreased ISC, MCC and SSC markers even when Notch activity was not stimulated (**Fig. S13F**). These experiments revealed that Spdef mediates positive input from Notch signaling to activate *ΔN-tp63* and to specify BCs.

Based on these experimental results, we added the regulatory connections to the model, re-fitted parameters, and modeled cell type production (model 2) (**Fig. 5E, S12B**). Model 2 improved fitting quality, including for ligand and *hes5.10* levels (**Fig. 5F, Table S2**). Model 2 could explain Notch gain- and loss-of-function effects by a temporal transformation representing de- (for loss) and acceleration (for gain) of cell type production rates (**Fig. 5G,H, S14A,B**). Furthermore, model 2 generated results in line with experimental observations on altering ligand expression from multipotent progenitors (**Fig. 5I, S14C**). Hence, the modelling approach can be used to predict critical gaps in our knowledge and to develop testable hypotheses.

To expand and challenge the model, we included data from additional stages (dataset 3, **Fig. S5A, S14D, Table S3**), Ubp1 as ISC regulator (**Fig. S12B**), and IF data from Hes manipulations on cell type composition. We then re-fitted parameters and modeled cell type production in model 2.1 (**Fig. S14E,F**). We tested if the model could predict the effects of *hes* manipulations on the expression of cell type markers at the end of specification (st. 16/17), which we have experimentally assessed by WMISH, but which were not included in the data used for parameter fitting (**Fig. S10, S15A,B**). The model predictions matched experimental results in 16 of 24 cases (66,66%) (**Fig. S15C**). Predictions for *hes7.1* manipulations were less correct (50%) than for Hes4 and Hes5.10 (75%), likely reflecting the over-simplified role for Hes7.1 as inducer of ISCs, which we have assigned to it for simplification of modeling. Furthermore, the effects on BCs were less correct (*ΔN-tp63*, 33.33%) than for ISCs (*foxi1*, 83.33%), MCCs (*mcidas/foxj1*, 83.33%) or SSCs (*foxa1*, 66.66%). This is probably due to the important co-regulation of *ΔN-tp63* by Wnt/β-catenin signaling^17^, which is not reflected in our model.

Taken together, the mathematical model represents key regulatory connections and recapitulates normal mucociliary patterning *in silico*. This can be used to reveal critical gaps in our understanding as well as to predict trends for biological effects caused by manipulations.

## Discussion

While Notch-regulation of mucociliary cell fates has been demonstrated across diverse systems, how Notch and Hes factors are applied in these systems to generate stable cell type compositions remains incompletely resolved^28,32,33,35,49^. Notch was proposed to act through lateral inhibition, but a “simple” version of lateral inhibition cannot explain how more than two cell fates are established. Our results are compatible with a modified application of the lateral inhibition paradigm between earlier and later emerging mucociliary progenitors (MPPs), where earlier formed MPPs express the majority of ligands and activate Notch in later emerging MPPs. This would not only lead to higher Notch activation levels in later emerging MPPs facilitating the specification of SSCs and BCs, but also protect early MPPs and differentiating ISCs/MCCs from excessive Notch activation via cis-inhibition^50^. Cis-inhibition can be achieved in early MPPs by high levels of Notch ligands expressed on their surface during cell fate specification stages. This is likely a relevant feature of this patterning mechanism, as overactivation of Notch in MCCs after cell fate specification leads to trans-differentiation or cell death in *Xenopus* and the mammalian airways^51,52^. While miRNA-mediated repression of *dll1* and *notch1* has been demonstrated during MCC differentiation^41^, cis-inhibition could support MCC differentiation early in the process. In contrast to initial development where an equipotent population of cells needs to respond to different Notch ligand levels, the cell type-specific Notch sensitivity can be established through differential Notch receptor expression in different mature cell types during homeostasis and regeneration^29,53^.

The increase in ligand expressing MPPs during patterning leads to an increase in Notch activity levels, regulating Hes gene expression. Hence, the proportions of cell types produced are in part controlled by the dynamics of MPP generation, their ligand expression and local movements. Changing these parameters should result in a different slope of signaling increase over time and lead to different proportions of cell types in the tissue as supported by our mathematical model. In the *Xenopus* epidermis, Hes7.1 seems to regulate the onset of MPP generation, but how this is achieved precisely will require more detailed investigation. In contrast, Hes4 and Hes5.10 drive differential cell fate choices via WRPW-domain dependent active repression. Hes4 and Hes5.10 seem to have different affinities to regulatory regions of cell fate determining transcription factors, and *hes4* is activated at lower Notch levels than *hes5.10*. Different combinations and levels of Hes factors can account for the selection of more than two cell types, e.g. MCCs (Hes4 high/Hes5.10 low), SSCs (Hes4 low/Hes5.10 medium) and BCs (Hes4 low/Hes5.10 high). Hes factors regulate cell fate specification by inhibiting alternative fate choices, a mode we termed “competitive de-repression” to highlight the negative-selection feature in this mechanism. While we cannot rule out a contribution of oscillations between ligand and *hes* expression during the patterning process, the two well-established oscillators in *Xenopus*, Hes7.1 and Hes1^45,54^, do not seem to regulate cell fate selection, but the onset of patterning and potentially proliferation, respectively. Oscillating behaviors for Notch components or Hes factors have not been reported in airway epithelial patterning either, and it was suggested that oscillatory behaviors rather regulate state transitions than cell fate acquisition during neurogenesis^55^. It is attractive to speculate that oscillatory behavior by Hes7.1 could regulate a similar transition during initiation of *Xenopus* epidermis development and thereby control the desynchronized emergence of MPPs during patterning. How Hes7.1 is activated in this context remains unresolved, but BMP signaling might play a role, as stimulation of deep layer cells with BMP in early stages of patterning was reported to induce uniform *dll1* expression and loss of mucociliary cell types^42^, similar to Hes7.1 overexpression.

While the identified mechanism of Hes-mediated competitive de-repression would be sufficient to regulate the fates of four cell types theoretically, it is likely not the sole mediator of Notch input towards cell fate decision output. Our mathematical modeling supported this conclusion: Based only on data derived from normal developing organoids and embryos, the model could fit most parameters to generate a correct cell type composition. However, not ligand and *hes5.10* expression, which decline after the phase of cell fate specification. This suggested that a state-transition is needed at the end of patterning in the system as well. Based on known functions and expression patterns of Spdef in mammalian mucociliary epithelia and cancers as well as our RNA-seq data, we hypothesized that Spdef could provide an activating component to the patterning process in response to Notch signaling. Indeed, scRNA-seq data suggested that early and intermediate MPPs have different *spdef* expression profiles, and *spdef* expressing cells were predominantly found among differentiating SSCs and BCs. We validated experimentally that Spdef is required for Notch activation of *ΔN-tp63*. This regulatory connection was implemented in our model version 2 in the form of: (1) Spdef levels increase with Notch levels, and (2) high Spdef levels activate *ΔN-tp63* expression. This effectively acts as switch to terminate cell fate specification, as ΔN-tp63 induces BC formation and suppresses further specification of ISCs, MCCs, and SSCs^17^. It will be interesting to test if Spdef also regulates *ΔN-tp63* in airway BCs, particularly because of Spdef’s important role as marker for progression of chronic airway diseases.

Taken together, we revealed that different cell types are predominantly produced during different phases of development and that this is driven by a temporal increase in Notch signaling levels acting on Hes genes that differentially regulate early and late emerging MPPs. This is a surprising deployment of Notch signaling during cell fate specification, but could be also applicable to other mucociliary systems in modified fashion. In the developing lung, such a temporal patterning mode would have to be deployed in a spatio-temporal manner, as lung buds grow at their distal tips^34,56^. Compatible with the mechanism described here being applied spatio-temporally, cells proliferate at the distal tips, then enter specification in more proximal regions, where low Notch-requiring MCCs are specified first and express Notch ligands on their surface, while high Notch-requiring secretory cells are formed in “older” more proximal parts of the epithelial lining^57^. This will be interesting to investigate further, and to test if our mathematical model could be applied in such manner as well. Our data-driven mathematical modeling approach allows to test validated and hypothetical regulatory connections during mucociliary patterning, and can be useful to highlight potential gaps in our understanding of regulatory processes. This can promote investigations by helping to prioritize which mechanistic aspects to investigate in order to reveal principles and adaptations of mucociliary patterning in development and disease.

Collectively, this work presents a coherent model for Notch- and Hes-mediated mucociliary cell fate specification in a vertebrate tissue, which allows for the specification of more than two cell fates within the Notch lateral-inhibition paradigm. We propose that tissue specific predispositions drive a default cell fate (ISCs), while additional cell fates (MCCs, SSCs and BCs) are induced by Hes-dependent competitive de-repression in combination with Spdef in response to Notch signaling levels.

## Supporting information

Figures S1-15, Tabels S1-9, Modeling Methods

Mathematical Model 1 and 2

Mathematical Model 2.1

## Acknowledgments

We thank: S. Schefold and C. Softley for expert technical help; W. Driever, G. Pyrowolakis, C. Kintner, L. Kodjabachian, J. Sedzinski and all Walentek lab members for discussions; Xenbase, EXRC for Xenopus resources; Light Imaging Center Freiburg, BiMiC and Aqua Core for microscope/animal resources; B. Grüning and the Freiburg Galaxy Team for bioinformatics platform and support. This study was supported by the Deutsche Forschungsgemeinschaft (DFG) under the Emmy Noether and Heisenberg Programmes (grant WA3365/2-1 and WA3365/5-1), by DFG/ANR (grant WA3365/4-1) and the NHLBI through a Pathway to Independence Award (K99HL127275) as well as by funding from the European Research Council (ERC) under the 2020 research and innovation programme (MAGIX; No. 101170596) to PW; And by funding under Germany’s Excellence Strategy (CIBSS – EXC-2189 – Project ID 390939984) to PW and CK; SHAU was funded by DFG CRC1453 grant 431984000.

## Author contribution

**Experiments:** MMBE: Notch and hes functions; MOH: imaging and reporter studies; MH: spdef data; AA: Hes-repression mode; ATP, SB, SH, DW, AT, PW: experimental support. **Bioinformatics and modeling:** PW, SHAU: bioinformatics analysis; TL, FL, CK: mathematical model implementation; MMBE, PW, TL, FL, CK: development of mathematical modeling approach. **Other:** PW: study design and supervision, coordinating collaborative work, manuscript preparation with input from all authors. MMBE, MOH contributed equally and can list themselves as first co-first authors.

## Materials and Methods

### Animal models and data availability

#### Xenopus laevis

Wild-type *Xenopus laevis* were obtained from the European Xenopus Resource Centre (EXRC) at University of Portsmouth, School of Biological Sciences, UK, or Xenopus 1, USA. Frog maintenance and care was conducted according to standard procedures in the AquaCore facility, University Freiburg, Medical Center (RI_00544) and based on recommendations provided by the international Xenopus community resource centers NXR (RRID:SCR_013731) and EXRC as well as by Xenbase (http://www.xenbase.org/, RRID:SCR_003280)^58^.

#### Ethics statements on animal experiments

This work was done in compliance with German animal protection laws and was approved under Registrier-Nr. G-18/76 and G-23/53 by the state of Baden-Württemberg.

#### Material, data and code availability

NGS data generated for this study was deposited at NCBI GEO under the accession numbers GSE215419 (RNA-seq dataset 1), GSE215373 (RNA-seq dataset 2) and GSE262944 (RNA-seq dataset 3). Additionally, published data from Haas *et al.* (GSE130448) were used^17^. Imaging data are available to the scientific community upon request to. Mathematical modeling details and code are available from Github (https://github.com/kreutz-lab/NotchPaper).

### Xenopus experiments

#### Manipulation of *Xenopus* Embryos

*X. laevis* eggs were collected and *in vitro*-fertilized, then cultured and microinjected by standard procedures^59,60^. Embryos were injected with Morpholino oligonucleotides (MOs, Gene Tools), mRNAs or plasmid DNA at two-cell to eight-cell stage using a PicoSpritzer setup in 1/3x Modified Frog Ringer’s solution (MR) with 2.5% Ficoll PM 400 (GE Healthcare, #17-0300-50), and were transferred after injection into 1/3x MR containing Gentamycin. Drop size was calibrated to about 7–8nL per injection.

Morpholino oligonucleotides (MOs) were obtained from Gene Tools targeting *dll1, hes4, hes5.10, hes7.1, notch1* and used at doses as indicated below.

##### Morpholino Sequences

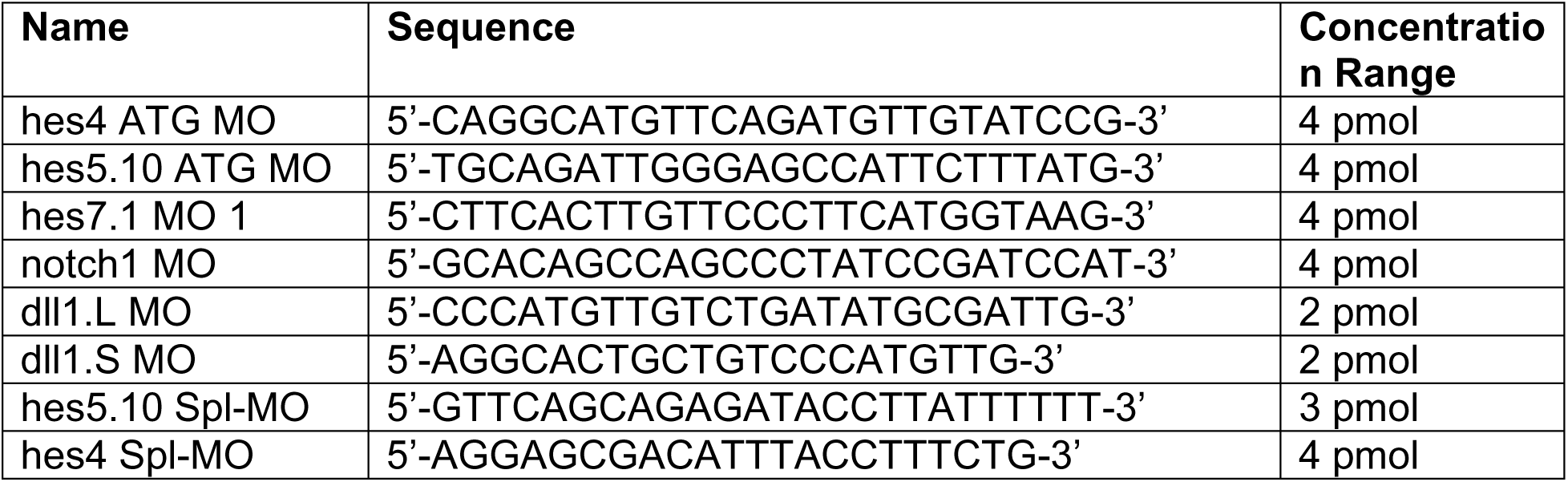
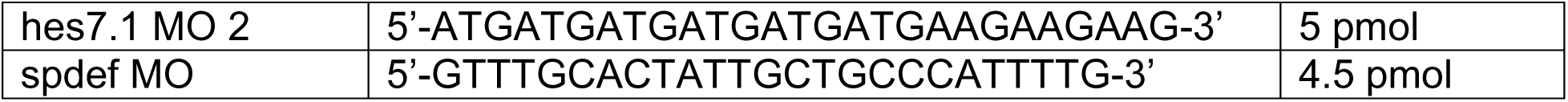

Full-length *hes4, hes5.10, hes7.1* constructs were cloned from total reverse-transcribed cDNA into pCS107 for overexpression. The *spdef* construct was donated by E. Dubaissi and encodes *Xenopus* tropicalis *spdef* in pCS2+ (matching reference sequence NM_001078944.1; mRNA was used at 100 ng/µL). mRNAs encoding membrane-BFP, membrane-GFP and Centrin-CFP (at 50ng/µL) or H2B-RFP (at 30ng/µL) were used in some experiments as lineage tracers. All mRNAs were prepared using the Ambion mMessage Machine kit using Sp6 (#AM1340) supplemented with RNAse Inhibitor (Promega #N251B).

##### ISH probe, full length and deletion construct cloning (3’-5’)

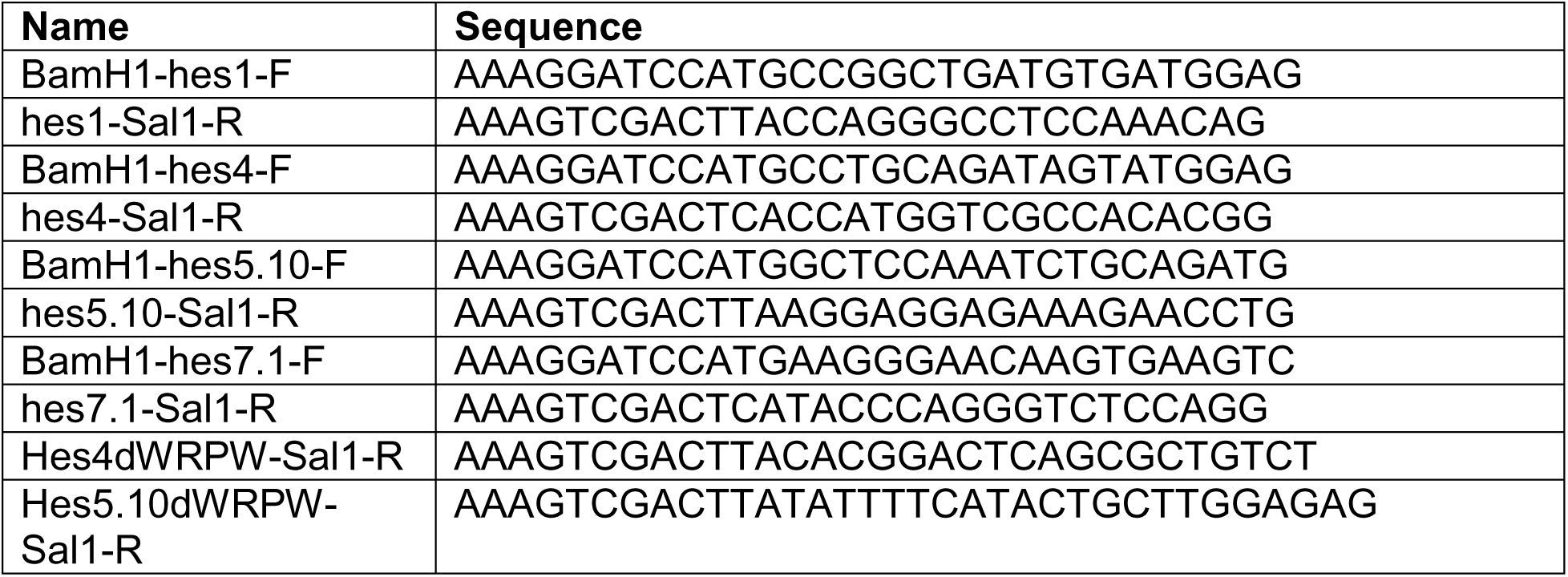

Plasmids were purified using the Pure Yield midiprep kit (Promega #2492). Notch reporter 4xcsl::H2B-mVenus^39^ and hes1::dsred^40^ plasmids were injected at 30ng/µl. Dll1 reporter plasmids *dll1*::mNeongreen and *dll1*::gfp-utrophin were injected at 10 ng/µl and *foxi1*::mScarletI at 5 ng/µl.

### *dll1*.S and *foxi1.S* reporter constructs

To generate the *dll1*.S::gfp-utrophin and *dll1*.S::mNeongreen reporter constructs, genomic DNA was prepared from *X. laevis* using the phenol/chloroform DNA purification (ThermoFisher #15593031 and associated protocol). A 1785 bp fragment (cf. Fig. S6C) of the dll1.S promoter was cloned from the genomic DNA using Easy-A Hi-Fi Cloning Enzyme (Agilent #600404) and primers listed in the table below. The *dll1*.S promoter fragment was subcloned into a-tub::gfp-utrophin(used in ^51^) after removal of the a-tub promoter sequence using HiFi DNA Assembly (NEB #E2621S) and Q5 High-Fidelity DNA Polymerase (NEB #M0491S). The *dll1*::mNeongreen reporter was generated by replacing the gfp-utrophin sequence in *dll1*::gfp-utrophin by the mNeongreen sequence using HiFi DNA Assembly and Q5 High-Fidelity DNA Polymerase, primers listed below. The *foxi1*::mScarletI construct was generated by replacing the gfp-utrophin sequence in a-*foxi1*::gfp-utrophin (used in ^10^) by the mScarletI sequence using Q5 High-Fidelity DNA Polymerase (NEB #M0491S) and HiFi DNA Assembly (NEB #E2621S), primers listed below. Final construct sequences were analyzed by whole-plasmid nanopore sequencing.

#### Cloning primers reporters (3’-5’)

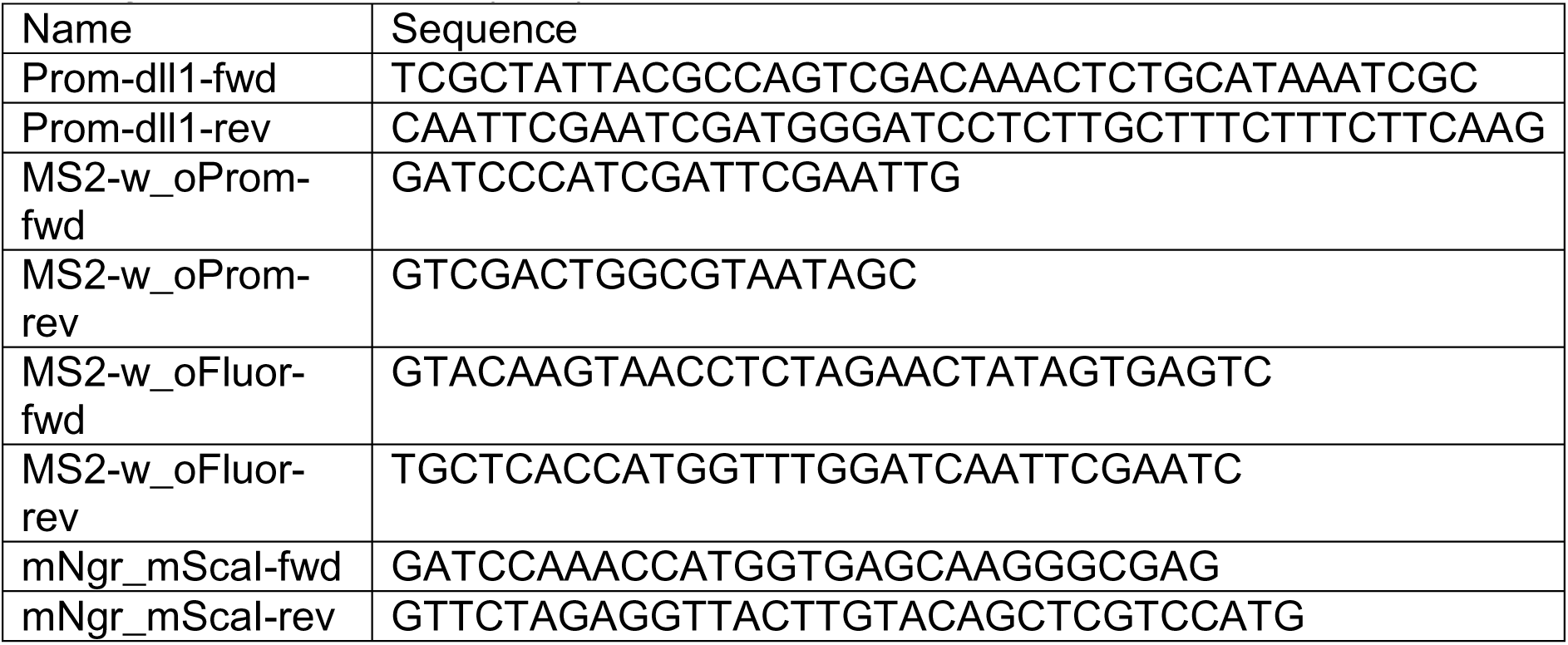

##### Preparation of mucociliary organoids (animal caps)

Injected and control embryos were cultured until st. 8. Animal caps were dissected in 1x Modified Barth’s solution (MBS)^11,61^ and transferred to 0.5x MBS + Gentamycin. 10-15 organoids were collected in Trizol (Thermo Fisher #15596026) per stage and used for qPCR or RNA-seq.

For live-cell imaging, human plasma fibronectin (Sigma #F0895) was diluted in 0.5x MBS (1:10), pipetted to a glass bottom (Ibidi #81218-200) and incubated overnight at 4°C. Animal caps were cut as described above and transferred immediately to the coated glass bottom dish filled with 0.5x MBS + Gentamycin. The deep cell layer of the animal cap was placed towards the fibronectin, allowing the tissue to attach and pressed gently down with a coverglass. After attachment (∼1 hr) the coverglass was removed and live-cell imaging was set up.

### Staining of Xenopus embryos and organoids

#### Immunofluorescence Staining

A detailed protocol was previously published^11^. In brief, whole *Xenopus* embryos, were fixed at indicated stages in 4% paraformaldehyde at 4°C overnight or 2h at RT, then washed 3x 15min with PBS, 2x 30min in PBSTx (0.1% Triton X-100 in PBS), and were blocked in PBSTx-CAS (90% PBS containing 0.1% Triton X-100, 10% CAS Blocking; ThermoFischer #00-8120) for 30min-1h at RT.

Mouse anti-Acetylated-α-tubulin (Sigma/Merck #T6793, 1:1000) or Mouse 594-conjugated Acetyl-Tubulin (proteinTech #CL594-66200, 1:500) primary antibody was used to mark cilia / MCCs, Rabbit Anti-serotonin (Sigma/Merck #S5545, 1:1000) primary antibody was used to mark SSCs, Rabbit anti-p63 (Abcam #ab124762, 1:500) primary antibody was used to mark BCs and Rabbit Anti-GFP (Invitrogen #A11122, 1:400) was used after HCR to boost *dll1*::gfp-utrophin signal. All primary antibodies were applied at 4°C overnight.

Secondary antibodies AlexaFlour-405-labeled goat anti-mouse (Invitrogen # A30104), AlexaFluor-488-labeled goat anti-mouse (Invitrogen #A11001), AlexaFlour-405-labeled goat anti-rabbit (Invitrogen #A31556), AlexaFlour-488-labeled goat anti-rabbit (Invitrogen #A11008) and AlexaFlour-647-labeled goat anti-rabbit (Invitrogen #A21245) were used for 2 h at RT (1:250). Antibodies were applied in 100% CAS Blocking (ThermoFischer #00-8120). Actin was stained by incubation (30-120 min at RT) with AlexaFluor 405-labeled Phalloidin (1:40 in PBSTx; Invitrogen #A30104), mucus-like compounds were stained by incubation (overnight at 4°C) with AlexaFluor-594- or -647-labeled PNA (1:500-1000 in PBSTx; Molecular Probes #L32459 and #L32460). Nuclei were stained with DAPI (Invitrogen #D1306, 1:40) 10 min at RT.

#### Whole mount *in situ* hybridization, sections and HCR

For WMISH antisense *in situ* hybridization probes, *notch1* and *dlc* fragments were cloned into pGEM-T Easy (Promega #A137A) from whole-embryo cDNAs derived from stages between 3 and 30 using primers listed below (ISH-primers). Anti-sense probe templates were either linearized with Sac2 (New England BioLabs #R0157S) and synthesized with SP6 RNA polymerase (Promega #P108G) or Sac1 (New England BioLabs #r3156S) and synthesized with T7 (Promega #P207E). All sequences were verified by Sanger sequencing. In addition, previously published probes were used: *foxi1*, *foxj1*, *mcidas*, *foxa1*, *tp63*, and *dll1*^10,11^.

Embryos were fixed in MEMFA (100mM MOPS pH7.4, 2mM EGTA, 1mM MgSO4, 3.7% (v/v) Formaldehyde) overnight at 4°C and stored in 100% Ethanol at -20°C until used. DNAs were purified using the PureYield Midiprep kit and were linearized before in vitro synthesis of anti-sense RNA probes using T7 or Sp6 polymerase (Promega, #P2077 and #P108G), RNAse inhibitor and dig-labeled rNTPs (Roche, #3359247910 and 11277057001). Embryos were in situ hybridized according to^62^, bleached after staining with BM Purple (Roche #11442074001) and imaged. Sections were made after embedding in gelatin-albumin with Glutaraldehyde at 50-70μm as described in ^63^.

##### Probe cloning primers (5’-3’)

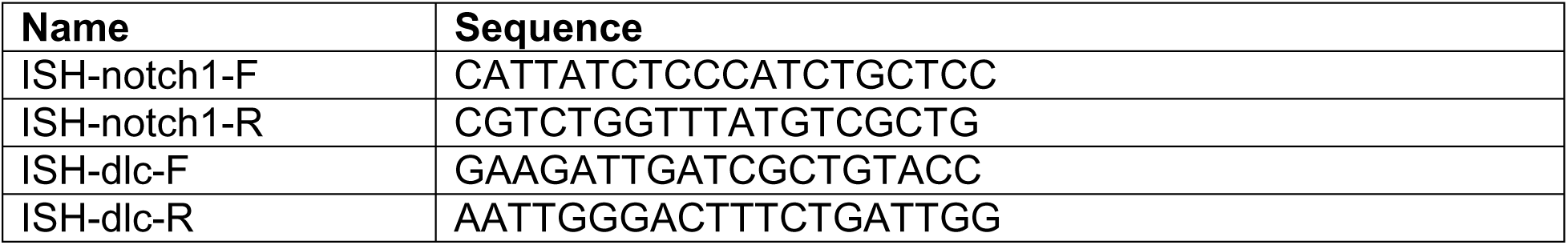

For HCR (hybridization Chain Reaction) whole *Xenopus* embryos were fixed at indicated stages in 10% MEMFA (100 mM MOPS pH 7.4, 2 mM EGTA, 1 mM MgSO_4_, 3.7% (v/v) Formaldehyde) for 1 h at RT, washed with PBSTw and stored in 100% Methanol at -20°C until use. Hybridization chain reaction was performed as previously described ^64^. *foxi1*, f*oxj1*, *foxa1*, *tp63* and *dll1* probes and amplifiers were designed and obtained from Molecular Instruments. IF staining was performed on samples after HCR following the steps described above. Hemisections for HCR stained samples were performed manually with a scalpel and samples were mounted in Fluoromount-G (ThermoFisher #00-4958-02).

#### Live-cell imaging

Images of Reporter injected planar animal caps were acquired with a confocal Zeiss Celldiscoverer 7 microscope using a 5x Objective and excitation lasers 488 nm for *dll1*::mNeongreen and 569 nm for *foxi1*::mScarletI. Time interval was set to 5 min and deep layer cells were imaged up to 12 hrs with temperature control (20 °C).

### Quantitative RT-PCR

Total RNA was extracted using a standard Trizol (Invitrogen #15596026) protocol and used for cDNA synthesis with iScript gDNA Clear cDNA Synthesis Kit (Bio-Rad #1725035). qPCR-reactions were conducted using Sso Advanced Universal SYBR Green Supermix (Bio-Rad #172-5275) on a CFX Connect Real-Time System (Bio-Rad) in 96-well PCR plates (Brand #781366).

Experiments for Notch reporter analysis was conducted in biological triplicates and technical duplicates and normalized by *ef1a* and *odc* expression levels. For dll1- and foxi1-reporter activity test analysis was conducted in biological duplicates and technical triplicates, expression levels were also normalized for injection levels (memRFP mRNA). Results are presented as log-transformed fold expression over average foxi1-reporter construct expression (Fig. 4B). Graphs were generated using R.

#### qPCR Primer Sequences

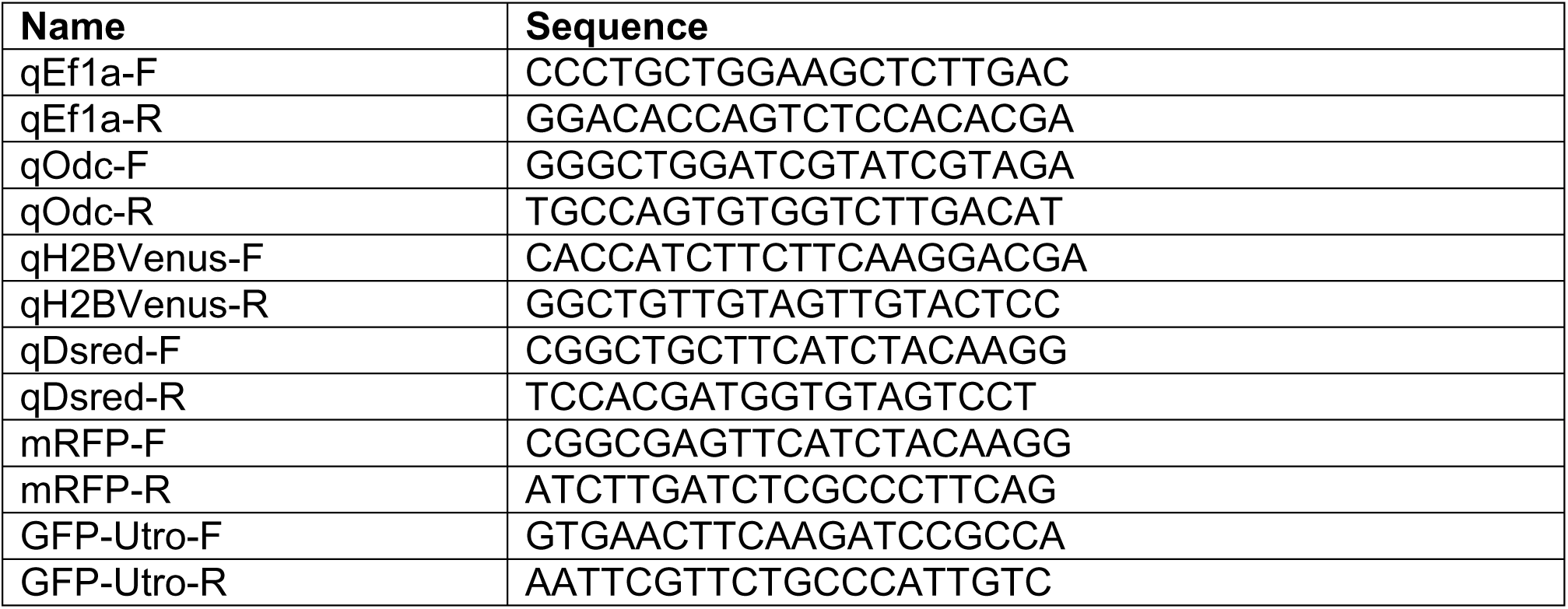

### Imaging and analysis of experiments

#### Confocal and bright field imaging and image processing

Confocal imaging was conducted using either a Zeiss LSM880, Zeiss LSM980 or Zeiss Celldiscoverer 7 microscope and Zeiss Zen software. Confocal images were adjusted for channel brightness/contrast, Z-stack projections were generated. A detailed protocol for quantification of *Xenopus* epidermal cell types was published^11^.

Images of embryos after *in situ* hybridization and corresponding sections were imaged using a Zeiss AxioZoom setup, Zeiss AxioImager.Z1 or Zeiss Stemi508 with Axiocam208-color, and images were adjusted for color balance, brightness and contrast using Adobe Photoshop.

#### Analysis

Embryos were staged according to Nieuwkoop and Faber (1994) Normal Table of *Xenopus laevis* (Daudin). Garland Publishing Inc, New York ISBN 0-8153-1896-0.

In Fig. 1D, 3D-G, 4A, S4B, S6B, S8C-E, S9B,C, S11A,C, S12A cell types were quantified based on their morphology using ImageJ^65^.

In Fig. 2F-H, S7B, images were analyzed for reporter activity and cell types were assigned based on morphological markers.

In Fig. 3H-J, expression of WMISH markers in the animal hemisphere was scored in whole mount embryos.

In Fig. 5D, S13F, expression of WMISH markers in the epidermis was scored based on differences observed between the uninjected control sides and manipulated sides of whole mount embryos.

In Fig. S2B-D, the number of marker(+) cells was scored in organoids and embryos from the same batches. Organoids were hemi-sected before staining and imaged from the surface as well as inner sides.

In Fig. S3B,C onset of marker gene expression was quantified after *in situ* hybridization according to the number of stained dots visible in (prospective) epidermal cells. Marker expression was categorized into three groups (no expression; <20 dots; >20 dots).

In Fig. S4C, S9A, S11B, the number of all nuclei (DAPI) and number of Tp63(+) cells was quantified.

In Fig. S10, expression level differences observed between the uninjected control sides and manipulated sides of embryos were scored in whole mount embryos, while depicted sections are shown for clarity. Please note that some changes appeared more clearly in sections (e.g. increase in *mcidas/foxj1* expression as well as increase/decrease in deep-layer *ΔN-tp63* expression) than in while mount quantifications.

### RNA-sequencing on *Xenopus* mucociliary organoids and bioinformatics analysis

*X. laevis* embryos were either injected 2-4x into the animal hemisphere at four-cell stage with mRNAs or MOs or remained uninjected, animal caps were prepared and used for RNA-sequencing at indicated stages. For RNA-seq dataset 1, uninjected organoids were collected at st. 9, st. 10.5 (labeled st. 10), st. 11.5 (labeled st. 11) and st. 12.5 (labeled st. 12) and organoids were derived from two independent experiments. For RNA-seq dataset 2, uninjected and manipulated (*suh-dbm*, *nicd*, *suh-dbm+dnmci*, *foxa1*MO) organoids were collected at st. 10.5 (labeled st. 10), st. 16 (st. 16) st. 25 (st. 25) and st. 32 (st. 32). Organoids were derived from 2 independent experiments. For RNA-seq dataset 3, uninjected organoids were collected at st. 14, st. 16 (please note that one replicate did not pass quality controls after sequencing and was excluded from further analysis) and st. 20. Organoids were derived from two independent experiments. A previously published dataset (*ΔN-tp63*MO RNA-seq, GSE130448) was used additionally. Datasets 1 and 3 were generated in collaboration with the NIG, University Medical Center Göttingen; dataset 2 was generated in collaboration with Genomics Facility at University of California, Berkeley.

500-1000 ng total RNA per sample was used, poly-A selection and RNA-sequencing library preparation was done using non strand massively-parallel cDNA sequencing (mRNA-Seq) protocol from Illumina, the TruSeq RNA Library Preparation Kit v2, Set A (Illumina #RS-122-2301) according to manufacturer’s recommendation. Quality and integrity of RNA was assessed with the Fragment Analyzer from Advanced Analytical by using the standard sensitivity RNA Analysis Kit (Advanced Analytical #DNF-471). For accurate quantitation of cDNA libraries, the QuantiFluor™dsDNA System from Promega was used. The size of final cDNA libraries was determined using the dsDNA 905 Reagent Kit (Advanced Bioanalytical #DNF-905) exhibiting a sizing of 300 bp on average. Libraries were pooled and paired-end 100bp sequencing on a HiSeq4000 was conducted at the Transcriptome and Genome Analysis Laboratory, University of Göttingen (RNA-seq dataset 1) or at the Functional Genomics Laboratory, University of California at Berkeley (RNA-seq dataset 2). Sequence images were transformed with Illumina software BaseCaller to BCL files, which was demultiplexed to fastq files with bcl2fastq v2.17.1.14. Quality control was done using FastQC v0.11.5 (Andrews, Simon (2010). “FastQC a quality-control tool for high-throughput sequence data” available at http://www.bioinformatics.babraham.ac.uk/projects/fastqc).

After adapter-trimming, paired-end reads were mapped to *Xenopus laevis* genome assembly v9.2 using RNA STAR v2.6.0b-1^66^. featureCounts v1.6.3^67^ was used to count uniquely mapped reads per gene and statistical analysis of differential gene expression was conducted in DEseq2 v1.22.1^68^. Heatmaps were generated in R v3.5.1 using ggplot2/heatmap2 v2.2.1. All bioinformatic analysis was performed on the Galaxy / Europe platform (usegalaxy.eu)^69^.

### Analysis of scRNA-sequencing data on *Xenopus* mucociliary organoids

Data from Lee et al.^12^ available at https://github.com/Natarajanlab/Xenopus-MCE-Atlas was imported into the NephGen scExplorer (https://nephgen.imbi.uni-freiburg.de/) using the Cerebro tool^70^. The number of cells assigned different clusters and time points as well as marker expressing cells per cluster and time point were exported into Excel and visualized as Heatmaps in R v3.5.1 using ggplot2/heatmap2 v2.2.1 performed on the Galaxy / Europe platform (usegalaxy.eu).

### Statistical evaluation

Stacked bar graphs were generated in Microsoft Excel, box plots (the line represents the median; 50% of values are represented by the box; 95% of values are represented within whiskers; values beyond 95% are depicted as outliers) were generated in R. Heatmaps and Venn diagrams were generated using the Galaxy Europe platform (usegalaxy.eu). Sample sizes for all experiments were chosen based on previous experience and used embryos derived from at least two different females. No randomization or blinding was applied. For some of the *in situ* and IF experiments shared controls were used in multiple graphs. X-test was conducted in Excel, Mann-Whitney-test was conducted at https://astatsa.com/WilcoxonTest/.

## References

1 Spassky, N. & Meunier, A. The development and functions of multiciliated epithelia. Nat Rev Mol Cell Biol 18, 423–436 (2017). 10.1038/nrm.2017.21

2 Walentek, P. Xenopus epidermal and endodermal epithelia as models for mucociliary epithelial evolution, disease, and metaplasia. Genesis 59, e23406 (2021). 10.1002/dvg.23406

3 Whitsett, J. A. Airway Epithelial Differentiation and Mucociliary Clearance. Ann Am Thorac Soc 15, S143–s148 (2018). 10.1513/AnnalsATS.201802-128AW

4 Walentek, P. Signaling Control of Mucociliary Epithelia: Stem Cells, Cell Fates, and the Plasticity of Cell Identity in Development and Disease. Cells Tissues Organs 211, 736–753 (2022). 10.1159/000514579

5 Hogan, B. L. et al. Repair and regeneration of the respiratory system: complexity, plasticity, and mechanisms of lung stem cell function. Cell Stem Cell 15, 123–138 (2014). 10.1016/j.stem.2014.07.012

6 Walentek, P. & Quigley, I. K. What we can learn from a tadpole about ciliopathies and airway diseases: Using systems biology in Xenopus to study cilia and mucociliary epithelia. Genesis 55 (2017). 10.1002/dvg.23001

7 Hendrickson, C. L. et al. Foxi2 and Sox3 are master transcription regulators that control ectoderm germ layer specification in Xenopus. PLoS Biol 23, e3003476 (2025). 10.1371/journal.pbio.3003476

8 Chalmers, A. D., Strauss, B. & Papalopulu, N. Oriented cell divisions asymmetrically segregate aPKC and generate cell fate diversity in the early Xenopus embryo. Development 130, 2657–2668 (2003). 10.1242/dev.00490

9 Huang, Y.-L. & Niehrs, C. Polarized Wnt Signaling Regulates Ectodermal Cell Fate in Xenopus. Developmental Cell 29, 250–257 (2014). 10.1016/j.devcel.2014.03.015

10 Bowden, S. et al. Foxi1 regulates multipotent mucociliary progenitors and ionocyte specification through transcriptional and epigenetic mechanisms. PLoS Biol 24, e3003583 (2026). 10.1371/journal.pbio.3003583

11 Walentek, P. Manipulating and Analyzing Cell Type Composition of the Xenopus Mucociliary Epidermis. Methods Mol Biol 1865, 251–263 (2018). 10.1007/978-1-4939-8784-9_18

12 Lee, J. et al. A single-cell, time-resolved profiling of Xenopus mucociliary epithelium reveals nonhierarchical model of development. Sci Adv 9, eadd5745 (2023). 10.1126/sciadv.add5745

13 Stubbs, J. L., Davidson, L., Keller, R. & Kintner, C. Radial intercalation of ciliated cells during Xenopus skin development. Development 133, 2507–2515 (2006). 10.1242/dev.02417

14 Chuyen, A. et al. The Scf/Kit pathway implements self-organized epithelial patterning. Dev Cell 56, 795–810.e797 (2021). 10.1016/j.devcel.2021.02.026

15 Quigley, I. K., Stubbs, J. L. & Kintner, C. Specification of ion transport cells in the Xenopus larval skin. Development 138, 705–714 (2011). 10.1242/dev.055699

16 Walentek, P. et al. A novel serotonin-secreting cell type regulates ciliary motility in the mucociliary epidermis of Xenopus tadpoles. Development 141, 1526–1533 (2014). 10.1242/dev.102343

17 Haas, M. et al. DeltaN-Tp63 Mediates Wnt/beta-Catenin-Induced Inhibition of Differentiation in Basal Stem Cells of Mucociliary Epithelia. Cell Rep 28, 3338–3352 e3336 (2019). 10.1016/j.celrep.2019.08.063

18 Collins, C. et al. Tubulin acetylation promotes penetrative capacity of cells undergoing radial intercalation. Cell Rep 36, 109556 (2021). 10.1016/j.celrep.2021.109556

19 Szabó, A. et al. The Molecular Basis of Radial Intercalation during Tissue Spreading in Early Development. Dev Cell 37, 213–225 (2016). 10.1016/j.devcel.2016.04.008

20 Quigley, I. K. & Kintner, C. Rfx2 Stabilizes Foxj1 Binding at Chromatin Loops to Enable Multiciliated Cell Gene Expression. PLoS Genet 13, e1006538 (2017). 10.1371/journal.pgen.1006538

21 Stubbs, J. L., Vladar, E. K., Axelrod, J. D. & Kintner, C. Multicilin promotes centriole assembly and ciliogenesis during multiciliate cell differentiation. Nat Cell Biol 14, 140–147 (2012). 10.1038/ncb2406

22 Lu, H. et al. Mcidas mutant mice reveal a two-step process for the specification and differentiation of multiciliated cells in mammals. Development 146 (2019). 10.1242/dev.172643

23 Arbi, M. et al. GemC1 controls multiciliogenesis in the airway epithelium. The EMBO Reports 17, 400–413 (2016). 10.15252/embr.201540882

24 Stubbs, J. L., Oishi, I., Izpisúa Belmonte, J. C. & Kintner, C. The forkhead protein Foxj1 specifies node-like cilia in Xenopus and zebrafish embryos. Nat Genet 40, 1454–1460 (2008). 10.1038/ng.267

25 Gomperts, B. N., Gong-Cooper, X. & Hackett, B. P. Foxj1 regulates basal body anchoring to the cytoskeleton of ciliated pulmonary epithelial cells. J Cell Sci 117, 1329–1337 (2004). 10.1242/jcs.00978

26 Dubaissi, E. et al. A secretory cell type develops alongside multiciliated cells, ionocytes and goblet cells, and provides a protective, anti-infective function in the frog embryonic mucociliary epidermis. Development 141, 1514–1525 (2014). 10.1242/dev.102426

27 Chen, G. et al. SPDEF is required for mouse pulmonary goblet cell differentiation and regulates a network of genes associated with mucus production. J Clin Invest 119, 2914–2924 (2009). 10.1172/jci39731

28 Deblandre, G. A., Wettstein, D. A., Koyano-Nakagawa, N. & Kintner, C. A two-step mechanism generates the spacing pattern of the ciliated cells in the skin of Xenopus embryos. Development 126, 4715–4728 (1999). 10.1242/dev.126.21.4715

29 Morimoto, M. et al. Canonical Notch signaling in the developing lung is required for determination of arterial smooth muscle cells and selection of Clara versus ciliated cell fate. J Cell Sci 123, 213–224 (2010). 10.1242/jcs.058669

30 Kurrle, Y., Kunesch, K., Bogusch, S. & Schweickert, A. Serotonin and MucXS release by small secretory cells depend on Xpod, a SSC specific marker gene. Genesis 58, e23344 (2020). 10.1002/dvg.23344

31 Ou-Yang, H. F., Wu, C. G., Qu, S. Y. & Li, Z. K. Notch signaling downregulates MUC5AC expression in airway epithelial cells through Hes1-dependent mechanisms. Respiration 86, 341–346 (2013). 10.1159/000350647

32 Gomi, K., Arbelaez, V., Crystal, R. G. & Walters, M. S. Activation of NOTCH1 or NOTCH3 signaling skews human airway basal cell differentiation toward a secretory pathway. PLoS One 10, e0116507 (2015). 10.1371/journal.pone.0116507

33 Tsao, P. N. et al. Notch signaling controls the balance of ciliated and secretory cell fates in developing airways. Development 136, 2297–2307 (2009). 10.1242/dev.034884

34 Tsao, P. N. et al. Gamma-secretase activation of notch signaling regulates the balance of proximal and distal fates in progenitor cells of the developing lung. J Biol Chem 283, 29532–29544 (2008). 10.1074/jbc.M801565200

35 Byrnes, L. E., Deleon, R., Reiter, J. F. & Choksi, S. P. Opposing transcription factors MYCL and HEY1 mediate the Notch-dependent airway stem cell fate decision. bioRxiv, 2022.2010.2005.511009 (2022). 10.1101/2022.10.05.511009

36 Song, R. et al. miR-34/449 miRNAs are required for motile ciliogenesis by repressing cp110. Nature 510, 115–120 (2014). 10.1038/nature13413

37 Chien, Y. H., Keller, R., Kintner, C. & Shook, D. R. Mechanical strain determines the axis of planar polarity in ciliated epithelia. Curr Biol 25, 2774–2784 (2015). 10.1016/j.cub.2015.09.015

38 Plasschaert, L. W. et al. A single-cell atlas of the airway epithelium reveals the CFTR-rich pulmonary ionocyte. Nature 560, 377–381 (2018). 10.1038/s41586-018-0394-6

39 Nowotschin, S., Xenopoulos, P., Schrode, N. & Hadjantonakis, A. K. A bright single-cell resolution live imaging reporter of Notch signaling in the mouse. BMC Dev Biol 13, 15 (2013). 10.1186/1471-213x-13-15

40 Matsuda, T. & Cepko, C. L. Controlled expression of transgenes introduced by in vivo electroporation. Proc Natl Acad Sci U S A 104, 1027–1032 (2007). 10.1073/pnas.0610155104

41 Marcet, B. et al. Control of vertebrate multiciliogenesis by miR-449 through direct repression of the Delta/Notch pathway. Nat Cell Biol 13, 693–699 (2011). 10.1038/ncb2241

42 Cibois, M. et al. BMP signalling controls the construction of vertebrate mucociliary epithelia. Development 142, 2352–2363 (2015). 10.1242/dev.118679

43 Session, A. M. et al. Genome evolution in the allotetraploid frog Xenopus laevis. Nature 538, 336–343 (2016). 10.1038/nature19840

44 Takada, H., Hattori, D., Kitayama, A., Ueno, N. & Taira, M. Identification of target genes for the Xenopus Hes-related protein XHR1, a prepattern factor specifying the midbrain-hindbrain boundary. Dev Biol 283, 253–267 (2005). 10.1016/j.ydbio.2005.04.020

45 Kageyama, R., Ohtsuka, T. & Kobayashi, T. The Hes gene family: repressors and oscillators that orchestrate embryogenesis. Development 134, 1243–1251 (2007). 10.1242/dev.000786

46 Raue, A. et al. Data2Dynamics: a modeling environment tailored to parameter estimation in dynamical systems. Bioinformatics 31, 3558–3560 (2015). 10.1093/bioinformatics/btv405

47 Turner, J. & Jones, C. E. Regulation of mucin expression in respiratory diseases. Biochem Soc Trans 37, 877–881 (2009). 10.1042/bst0370877

48 Uhlén, M. et al. Proteomics. Tissue-based map of the human proteome. Science 347, 1260419 (2015). 10.1126/science.1260419

49 Rock, J. R. et al. Notch-dependent differentiation of adult airway basal stem cells. Cell Stem Cell 8, 639–648 (2011). 10.1016/j.stem.2011.04.003

50. del Álamo, D., Rouault, H. & Schweisguth, F. Mechanism and Significance of *cis*-Inhibition in Notch Signalling. Current Biology 21, R40-R47 (2011). 10.1016/j.cub.2010.10.034

51 Tasca, A. et al. Notch signaling induces either apoptosis or cell fate change in multiciliated cells during mucociliary tissue remodeling. Dev Cell 56, 525–539 e526 (2021). 10.1016/j.devcel.2020.12.005

52 Tyner, J. W. et al. Blocking airway mucous cell metaplasia by inhibiting EGFR antiapoptosis and IL-13 transdifferentiation signals. J Clin Invest 116, 309–321 (2006). 10.1172/jci25167

53 Ruiz García, S., et al. Novel dynamics of human mucociliary differentiation revealed by single-cell RNA sequencing of nasal epithelial cultures. Development 146 (2019). 10.1242/dev.177428

54 Jen, W. C., Gawantka, V., Pollet, N., Niehrs, C. & Kintner, C. Periodic repression of Notch pathway genes governs the segmentation of Xenopus embryos. Genes Dev 13, 1486–1499 (1999). 10.1101/gad.13.11.1486

55 Biga, V. et al. Oscillatory Co-expression of HES1 and HES5 Enables a Hybrid State in a Bistable Transcription Factor Regulatory Motif. bioRxiv, 2025.2005.2022.655498 (2025). 10.1101/2025.05.22.655498

56 Nikolić, M. Z., Sun, D. & Rawlins, E. L. Human lung development: recent progress and new challenges. Development 145 (2018). 10.1242/dev.163485

57 Rawlins, E. L., Ostrowski, L. E., Randell, S. H. & Hogan, B. L. Lung development and repair: contribution of the ciliated lineage. Proc Natl Acad Sci U S A 104, 410–417 (2007). 10.1073/pnas.0610770104

58 Fisher, M. et al. Xenbase: key features and resources of the Xenopus model organism knowledgebase. Genetics 224 (2023). 10.1093/genetics/iyad018

59 Sive, H. L., Grainger, R. M. & Harland, R. M. Xenopus laevis In Vitro Fertilization and Natural Mating Methods. CSH Protoc 2007, pdb.prot4737 (2007). 10.1101/pdb.prot4737

60 Sive, H. L., Grainger, R. M. & Harland, R. M. Microinjection of Xenopus embryos. Cold Spring Harb Protoc 2010, pdb.ip81 (2010). 10.1101/pdb.ip81

61 Sive, H. L., Grainger, R. M. & Harland, R. M. Animal Cap Isolation from Xenopus laevis. CSH Protoc 2007, pdb.prot4744 (2007). 10.1101/pdb.prot4744

62 Harland, R. M. In situ hybridization: an improved whole-mount method for Xenopus embryos. Methods Cell Biol 36, 685–695 (1991). 10.1016/s0091-679x(08)60307-6

63 Walentek, P., Beyer, T., Thumberger, T., Schweickert, A. & Blum, M. ATP4a is required for Wnt-dependent Foxj1 expression and leftward flow in Xenopus left-right development. Cell Rep 1, 516–527 (2012). 10.1016/j.celrep.2012.03.005

64 Huber, P. B. & LaBonne, C. Small molecule-mediated reprogramming of Xenopus blastula stem cells to a neural crest state. Developmental Biology 505, 34–41 (2024). 10.1016/j.ydbio.2023.10.004

65. Schindelin, J., et al. Fiji: an open-source platform for biological-image analysis. Nat Methods 9, 676-682 (2012). 10.1038/nmeth.2019

66 Dobin, A. et al. STAR: ultrafast universal RNA-seq aligner. Bioinformatics 29, 15–21 (2013). 10.1093/bioinformatics/bts635

67 Danecek, P. et al. Twelve years of SAMtools and BCFtools. GigaScience 10 (2021). 10.1093/gigascience/giab008

68 Love, M. I., Huber, W. & Anders, S. Moderated estimation of fold change and dispersion for RNA-seq data with DESeq2. Genome Biol 15, 550 (2014). 10.1186/s13059-014-0550-8

69 Community, T. G. The Galaxy platform for accessible, reproducible, and collaborative data analyses: 2024 update. Nucleic Acids Research 52, W83–W94 (2024). 10.1093/nar/gkae410

70 Hillje, R., Pelicci, P. G. & Luzi, L. Cerebro: interactive visualization of scRNA-seq data. Bioinformatics 36, 2311–2313 (2019). 10.1093/bioinformatics/btz877

