## Supplementary material for "Temporal Notch signaling and Hes-mediated competitive de-repression regulate mucociliary cell fates in *Xenopus*": Figures S1-15, Tabels S1-9, Modeling Methods

### Brislinger, Hansen et al. Suppl Figure 1

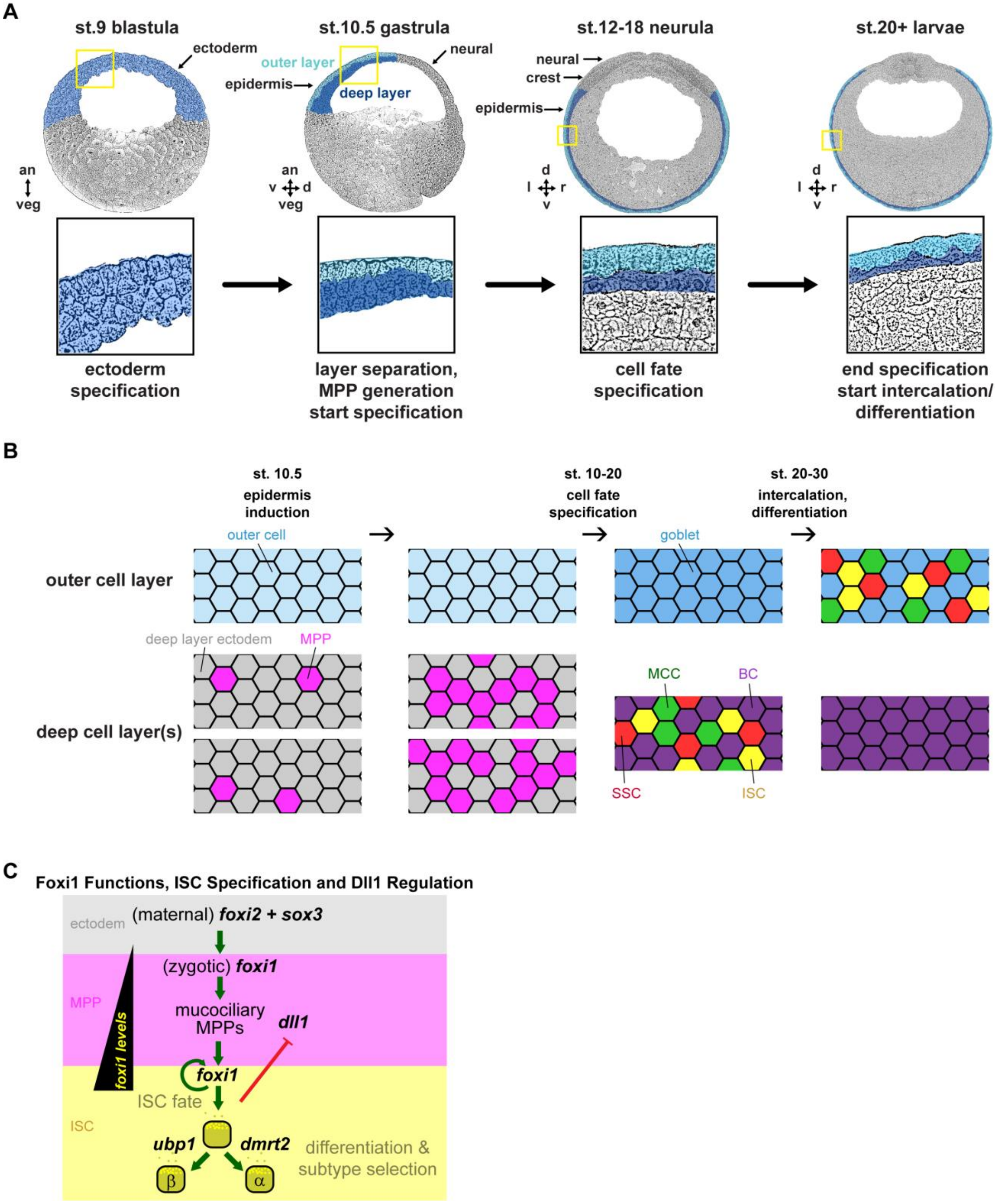

**Supplementary Figure S1: Background information on *Xenopus* mucociliary epidermis development**

**A:** Schematic representations of sections from a st. 9 blastula embryo (with prospective ectoderm expressing *foxi2* and *sox3* indicated in blue), of a st. 10.5 gastrula, a st. 12 neurula and a st. 20 embryo (with outer layer epidermal cells indicated in light blue and deep layer cells indicated in dark blue). Yellow boxes indicate areas shown magnified below. Please note the rearrangement of the deep cell layers into a single basal layer between st. 10.5 and st. 12+. Axes and embryo orientations are indicated: an = animal; veg = vegetal; d = dorsal; v = ventral; l = left; r = right. **B:** Schematic representation of outer and deep cell layers (en face view) at st. 10.5, during patterning (st. 10 – 20), after cell fate specification and at mature stages (st. 32). Outer layer cells = light blue; deep layer ectodermal precursors = grey; mucociliary multipotent progenitors (MPPs) = magenta; goblet cells = darker blue; ionocytes (ISCs) = yellow; multiciliated cells (MCCs) = green; small secretory cells (SSCs) = red; basal cells (BCs) = purple. **C:** Schematic representation of Foxi1 level-dependent generation of MPPs and ISCs in the *Xenopus* epidermis. Grey area depicts cell state as deep layer ectodermal progenitor, which changes into a low-level Foxi1- and Delta-like1 (Dll1) expressing MPP (magenta area), and then into one of the two ISC subtypes, driven by high Foxi1 expression achieved through positive auto-regulation of *foxi1* in cooperation with Ubp1 and Dmrt2, that drive β- and α-ISC fates, respectively. Please note that ISC differentiation and Ubp1 expression terminates *dll1* expression in differentiating MPPs/early ISCs during epidermal development.

**Brislinger, Hansen et al. Suppl Figure 2**

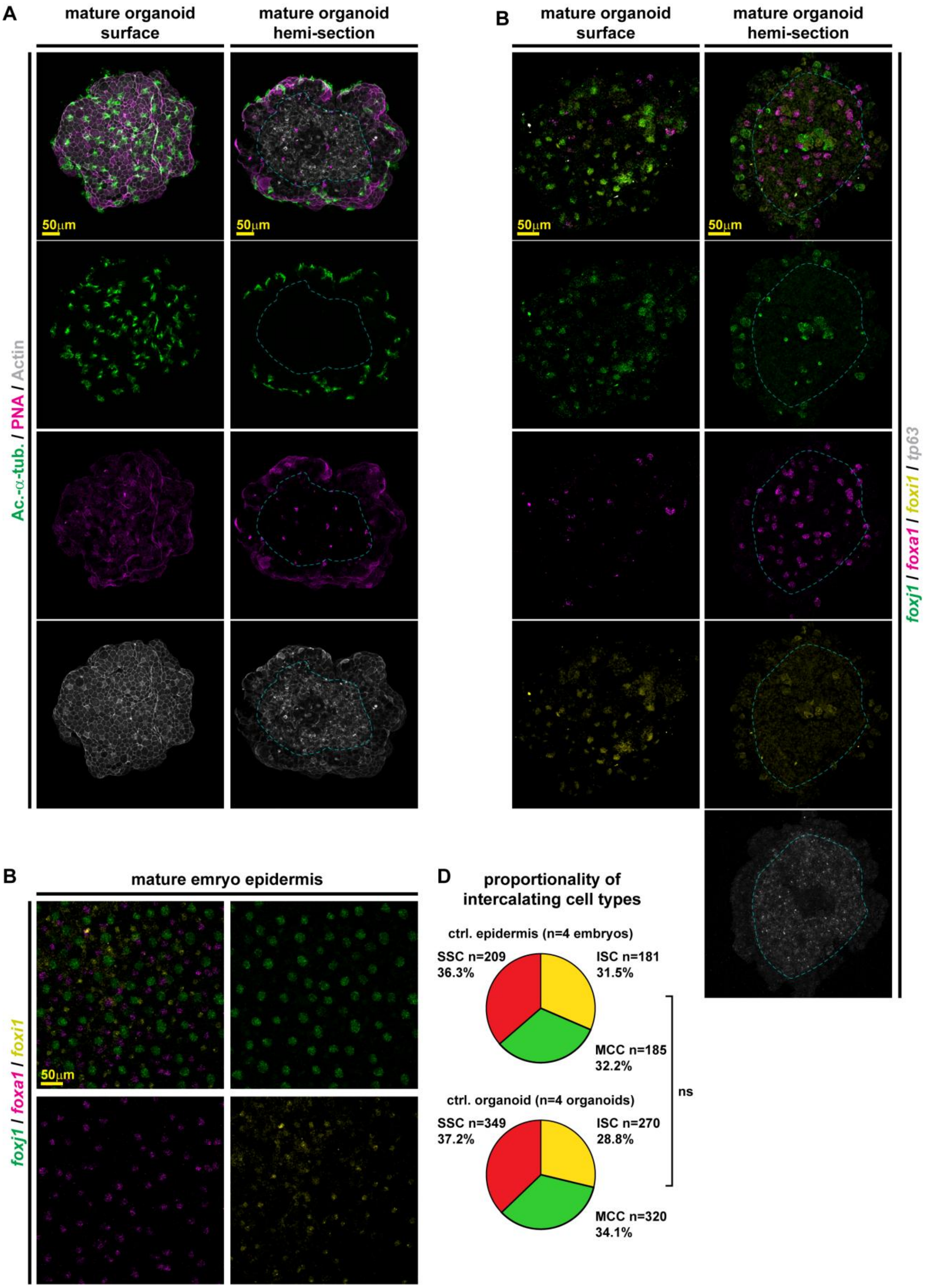

**Supplementary Figure S2: Comparison of cell type proportions between mucociliary organoids and the embryonic epidermis *in vivo***

**A:** Immunofluorescent images of mature (st. 32 - 35) mucociliary organoids stained for Acetylated-$\alpha$-tubulin (Ac.-$\alpha$-tub., green) that marks MCC cilia; for F-actin (Actin, grey) that marks cell borders and apical morphology; and with peanut agglutinin (PNA, magenta) that marks small mucus granules in goblet cells and large secretory vesicles in SSCs. Left panels = outside surface; right panels = internal view on the same hemi-sected organoid. Please note that SSCs with large PNA(+) granules frequently reside in the core of the organoid. **B,C:** Fluorescent micrographs of mature (st. 32 - 35) mucociliary organoids and the embryonic epidermis (derived from same batch of embryos) stained by fluorescent *in situ* hybridization chain reaction (HCR) using markers for ISCs (*foxi1*, yellow), MCCs (*foxj1*, green), SSCs (*foxa1*, magenta) and BCs (*tp63*, grey). **B:** Organoid. Left panels = outside surface; right panels = internal view on the same hemi-sected organoid. Please note that SSCs marked by *foxa1* frequently reside in the mostly *tp63*+ (BCs, grey) core of the organoid. **C:** En face view on the epidermis. **D:** Quantification of marker(+) cells in organoids and embryos. $X^2$-test, ns = $p > 0.05$.

**Brislinger, Hansen et al. Suppl Figure 3**

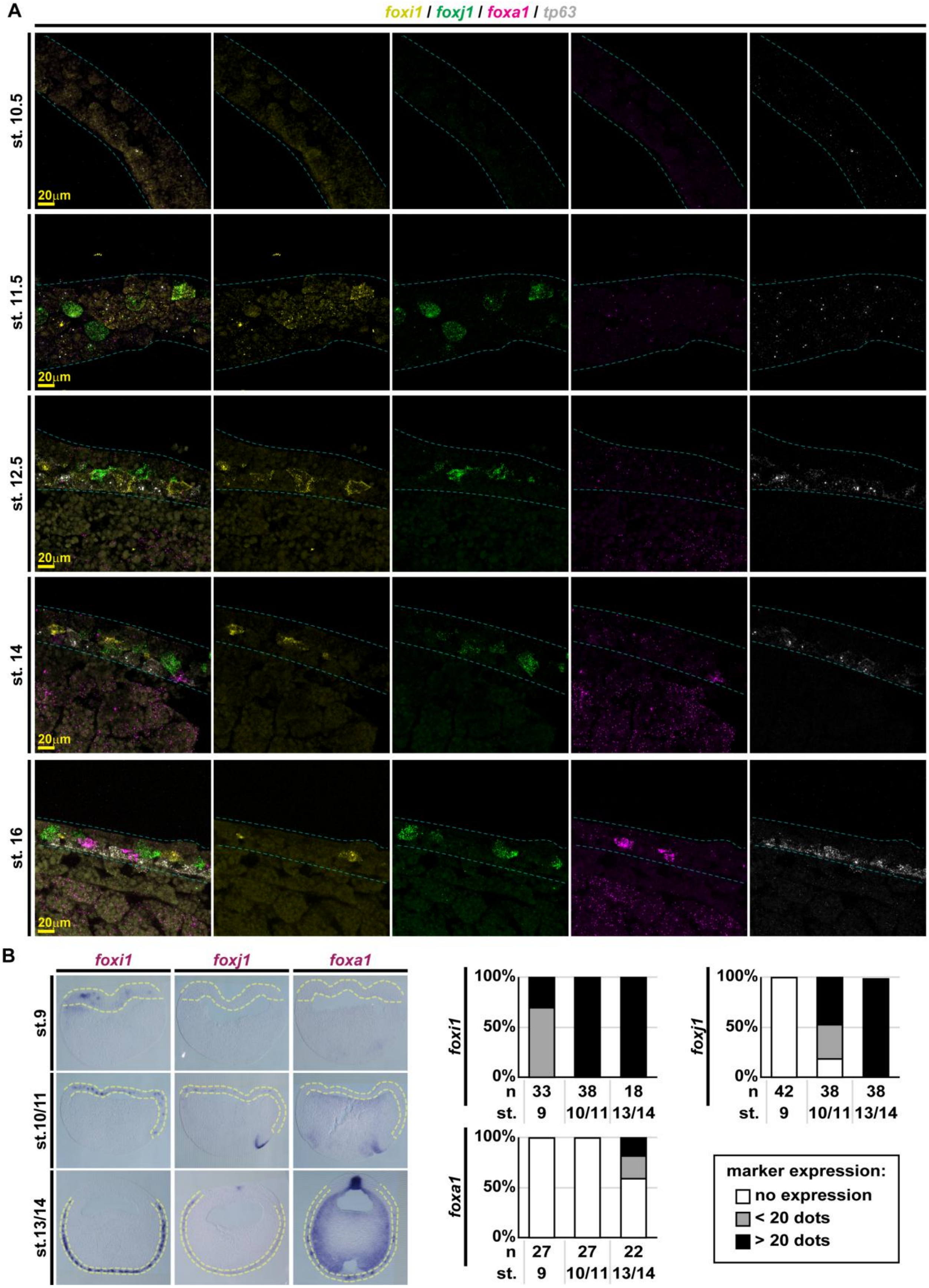

**Supplementary Figure S3: Sequential appearance of cell type markers during mucociliary epidermis development**

**A:** Fluorescent micrographs of sectioned embryos at different stages (st. 10.5 - 16) of epidermis development stained by fluorescent *in situ* hybridization chain reaction (HCR) using markers for MPPs/ISCs (*foxi1*, yellow), MCCs (*foxj1*, green), SSCs (*foxa1*, magenta) and BCs (*tp63*, grey). **B:** Embryo sections, chromogenic whole-mount *in situ* hybridization (WMISH) for indicated transcripts from st. 9 to st. 13/14. Epidermis outlined in yellow. Note the continuous low-level expression patches of *foxi1* vs. high-level expressing single cells at st.9. Graphs: Quantification of results from non-sectioned embryos. No expression = white; less than 20 high-level expressing cells = grey; more than 20 high-level expressing cells = black.

### Brislinger, Hansen et al. Suppl Figure 4

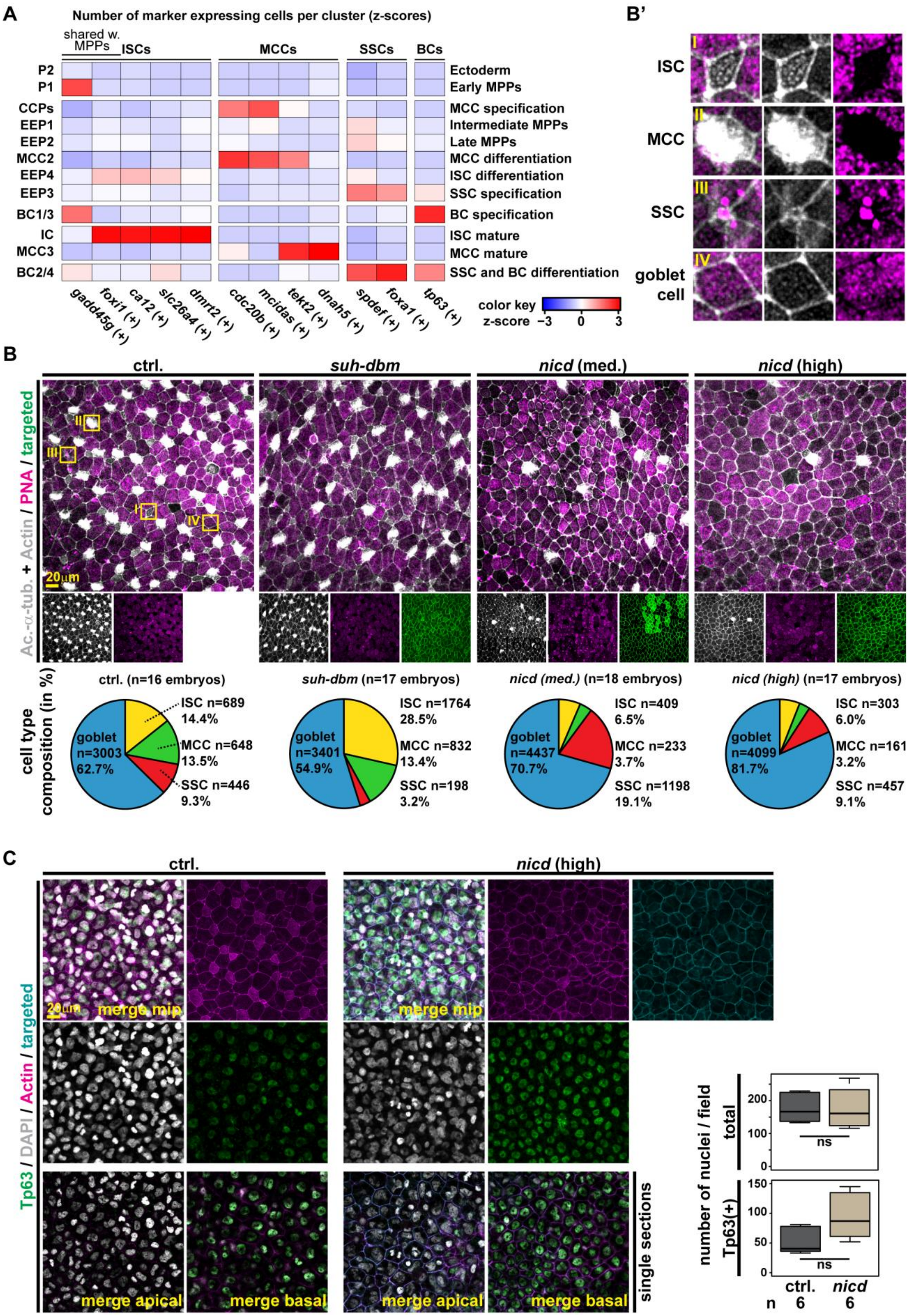

### Supplementary Figure S4: Cell state markers, cell type analysis and response to Notch manipulations

**A:** Heatmap of z-scores depicting enrichment of cells expressing markers within the different clusters (left side = cluster annotation by Lee et al.; right side = cluster identity annotation based on the markers indicated below each column). *Gadd45g* and *foxi1* are designated core-ISC genes also expressed from MPPs. **B:** Immunofluorescent images (IF) of epidermal cell types after Notch manipulations at st. 32. Suh-dbm = Notch loss-of-function; nicd = Notch gain-of-function; med. = medium level and high = high level nicd overexpression. Acetylated-α-tubulin (Ac.-α-tub., grey) marks MCC cilia; F-actin (Actin, grey) marks cell borders and apical morphology; peanut agglutinin (PNA, magenta) marks small mucus granules in goblet cells and large secretory vesicles in SSCs; membrane-GFP (targeted, green) marks targeted cells. Graphs below images: Quantification of results. **B':** Magnified views of individual cell types after marker staining. Location of depicted cells is indicated in control (ctrl.) image by yellow boxes. I = ISC; II = MCC; III = SSC; IV = goblet cell **C:** IF of epidermal cells in controls (ctrl.) and after high level nicd overexpression at st. 32. Tp63 (green) marks basal cells (BCs) in the deep layer; F-actin (Actin, magenta) marks cell borders and apical morphology; DAPI (grey) marks nuclei; membrane-GFP (targeted, cyan) marks targeted cells. Graphs on the right: Quantification of total cells and of Tp63(+) BCs per embryo/field. Mann-Whitney-test, ns = $p > 0.05$.

**Brislinger, Hansen et al. Suppl Figure 5**

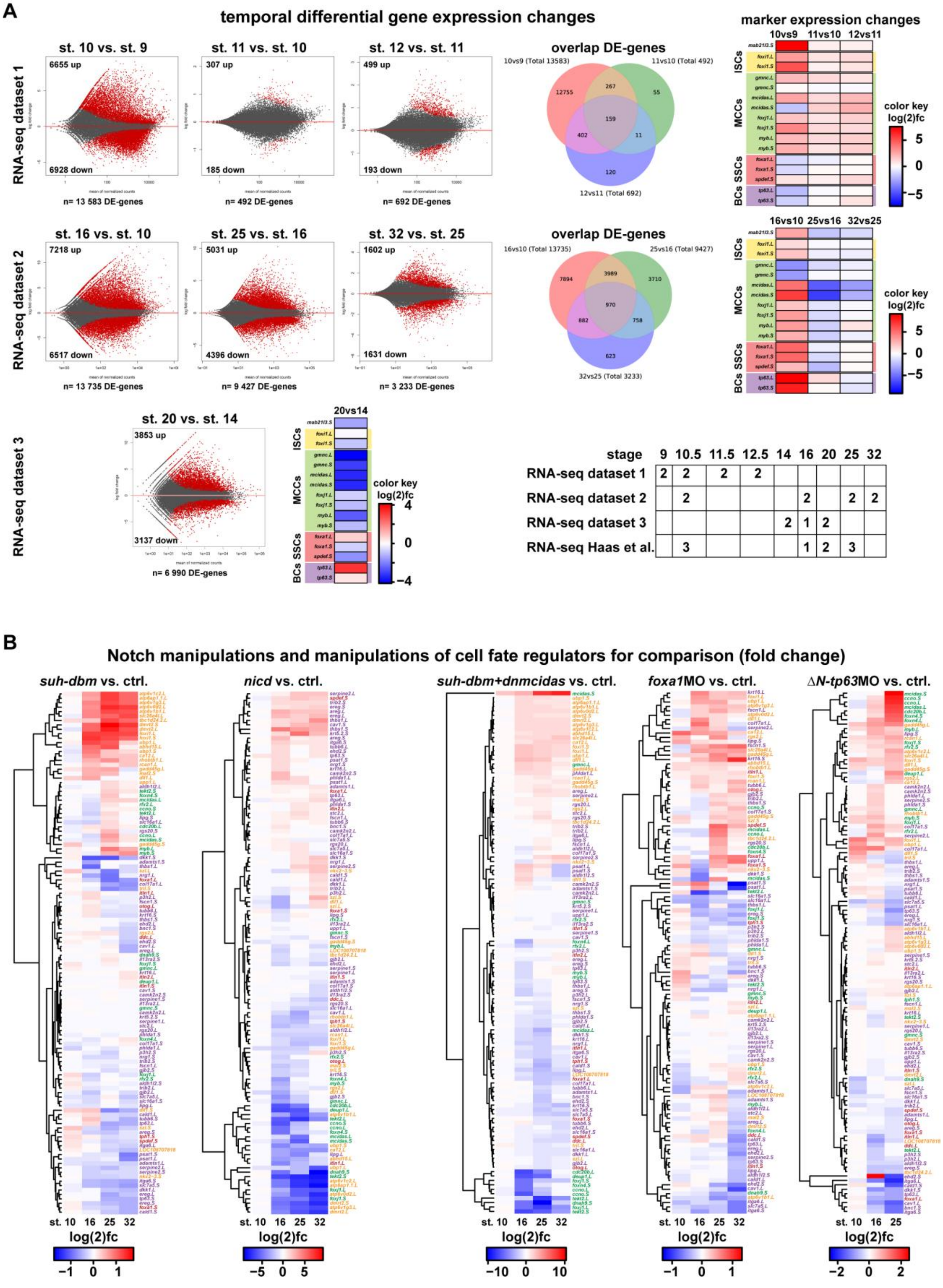

| stage | 9 | 10.5 | 11.5 | 12.5 | 14 | 16 | 20 | 25 | 32 |
|---|---|---|---|---|---|---|---|---|---|
| RNA-seq dataset 1 | 2 | 2 | 2 | 2 | | | | | |
| RNA-seq dataset 2 | | 2 | | | | 2 | | 2 | 2 |
| RNA-seq dataset 3 | | | | | 2 | 1 | 2 | | |
| RNA-seq Haas et al. | | 3 | | | | 1 | 2 | 3 | |

**Supplementary Figure S5: RNA-seq on control and Notch manipulated organoids**
**A:** RNA-seq datasets (n = 2 per stage and dataset; please note: in dataset 3, one st. 16 sample failed quality controls and was discarded) were used to investigate temporal expression dynamics in mucociliary organoids. Comparison of differentially expressed genes (red dots) across stages indicate an increase in early specification markers (heatmaps on the right side) between st. 9 and st. 16. Venn-diagrams depict unique and overlapping differentially expressed genes between stages. Distribution of RNA-seq samples and replicate numbers used in this study is shown as table. **B:** Heatmaps (log(2) fold-change) of RNA-seq datasets comparing sets of marker genes for ISCs (yellow), MCCs (green), SSCs/secretory cells (red) and BCs (purple) after various manipulations across multiple stages. *suh-dbm* vs. ctrl.: inhibition of Notch signaling (n = 2 per stage, depletion of SSCs and basal cells); *suh-dbm+dnmcidas* vs. ctrl.: inhibition of Notch signaling and inhibition of MCC formation by dominant-negative acting Mcidas (n = 2 per stage, depletion of SSCs and basal cells and MCCs); *nicd* vs. ctrl.: gain of Notch signaling (n = 2 per stage, depletion of ISCs and MCCs); *foxa1*MO vs. ctrl.: knockdown of *foxa1* (n = 2 per stage, depletion of SSCs); *ΔN-tp63*MO vs. ctrl.: knockdown of *ΔN-tp63* (n = 3 per stage, depletion of BCs).

### Brislinger, Hansen et al. Suppl Figure 6

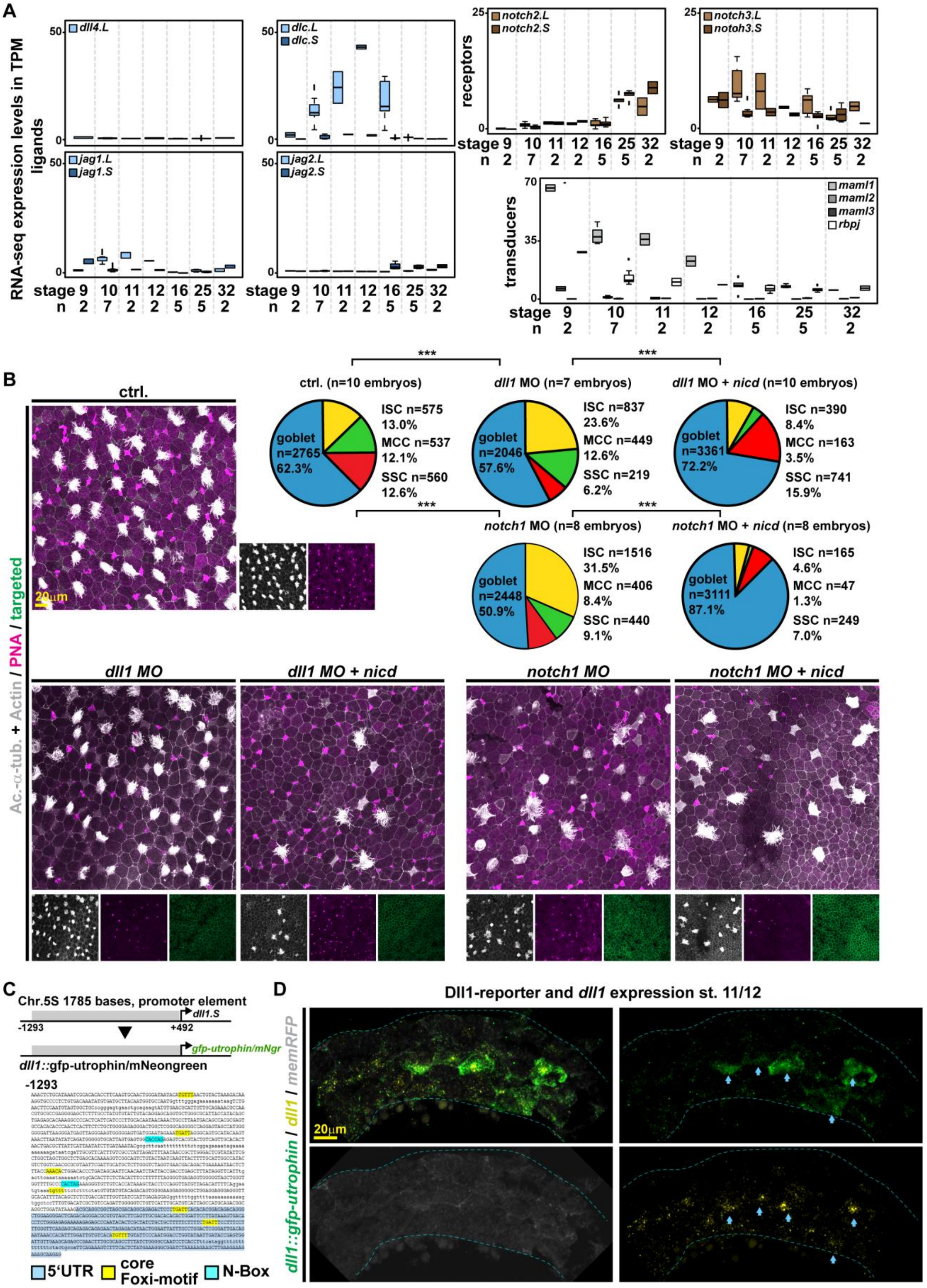

**Supplementary Figure S6: Expression and manipulation of Notch signaling components in mucociliary patterning**

**A:** Expression of additional Notch signaling components in unmanipulated mucociliary organoids (RNA-seq) over time. **B:** Immunofluorescent images of epidermal cell types at st. 32 in controls (ctrl.) and after knockdown of *dll1* (*dll1* MO) or *notch1* (*notch1* MO) alone and after co-injected with *nicd* mRNA. Acetylated-$\alpha$-tubulin (Ac.-$\alpha$-tub., grey) marks MCC cilia; F-actin (Actin, grey) marks cell borders and apical morphology; peanut agglutinin (PNA, magenta) marks small mucus granules in goblet cells and large secretory vesicles in SSCs; membrane-GFP (targeted, green) marks targeted cells. Quantification of results shown on the top right. $X^2$-test, *** = $p < 0.001$. **C,D:** *Dll1*-reporter generation. **C:** Schematic representation of cloned genomic *dll1.S* promoter locus (grey box), position of 5'UTR (darker blue), core-Foxi (yellow) and N-box (light blue) motifs. Two versions were generated expressing different fluorescent proteins under the *dll1*-promoter, GFP-utrophin fusion or mNeonGreen. **D:** Immunofluorescent image of sectioned epidermis at st. 11/12 from a *dll1*-reporter (green) injected embryo co-stained for endogenous *dll1* expression (HCR, yellow). Targeted cells labeled by co-injected membrane-RFP (mRFP, grey). Blue arrowheads indicate cells with *dll1*-reporter activity.

**Brislinger, Hansen et al. Suppl Figure 7**

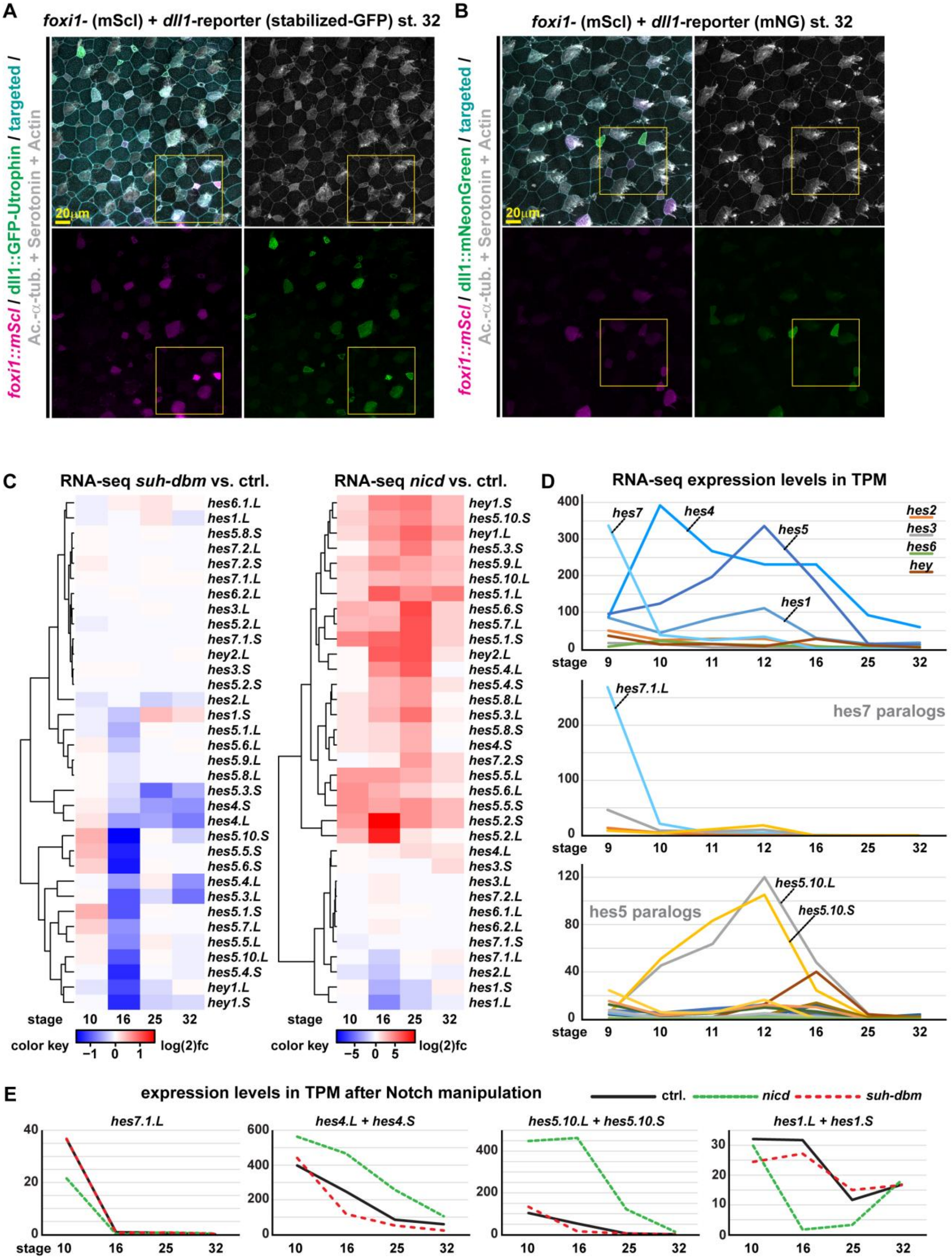

**Supplementary Figure S7: Expression of reporters and sequential expression of Hes transcription factors in the *Xenopus* epidermis**

**A,B:** Representative samples used for reporter analysis in Fig. 2F. Immunofluorescent images of epidermal cell types at st. 32 in reporter-injected embryos. Acetylated-α-tubulin (Ac.-α-tub., grey) marks MCC cilia; F-actin (Actin, grey) marks cell borders and apical morphology; Serotonin (grey) marks large SSC granules. Targeting was confirmed by co-injection of fluorescent membrane markers (cyan). *Dll1*-reporters (A: *dll1::gfp-utrophin* or B: *dll1::mNeonGreen*) are shown in green, *foxi1-mScarletI* (mScl; magenta) was co-injected to mark MPP derivative cells. Areas shown in Fig. 2F are indicated by yellow boxes in A and B. **C:** RNA-seq datasets comparing the expression of all known *Xenopus laevis hes/hey* genes after manipulation of Notch signaling across multiple stages. *suh-dbm* vs. ctrl.: inhibition of Notch signaling (n = 2 per stage); *nicd* vs. ctrl.: gain of Notch signaling (n = 2 per stage). **D:** Summed up expression levels of *hes* paralog groups in unmanipulated mucociliary organoids across different stages of development (RNA-seq), and expression levels of individual *hes* transcripts from *hes7* and *hes5* paralog groups identifies dominantly and sequentially expressed *hes* genes during mucociliary patterning. **E:** Expression levels of indicated *hes* transcripts in unmanipulated mucociliary organoids (black line) and after Notch manipulations (loss = red line; gain = green line) across different stages of development (RNA-seq).

**Brislinger, Hansen et al. Suppl Figure 8**

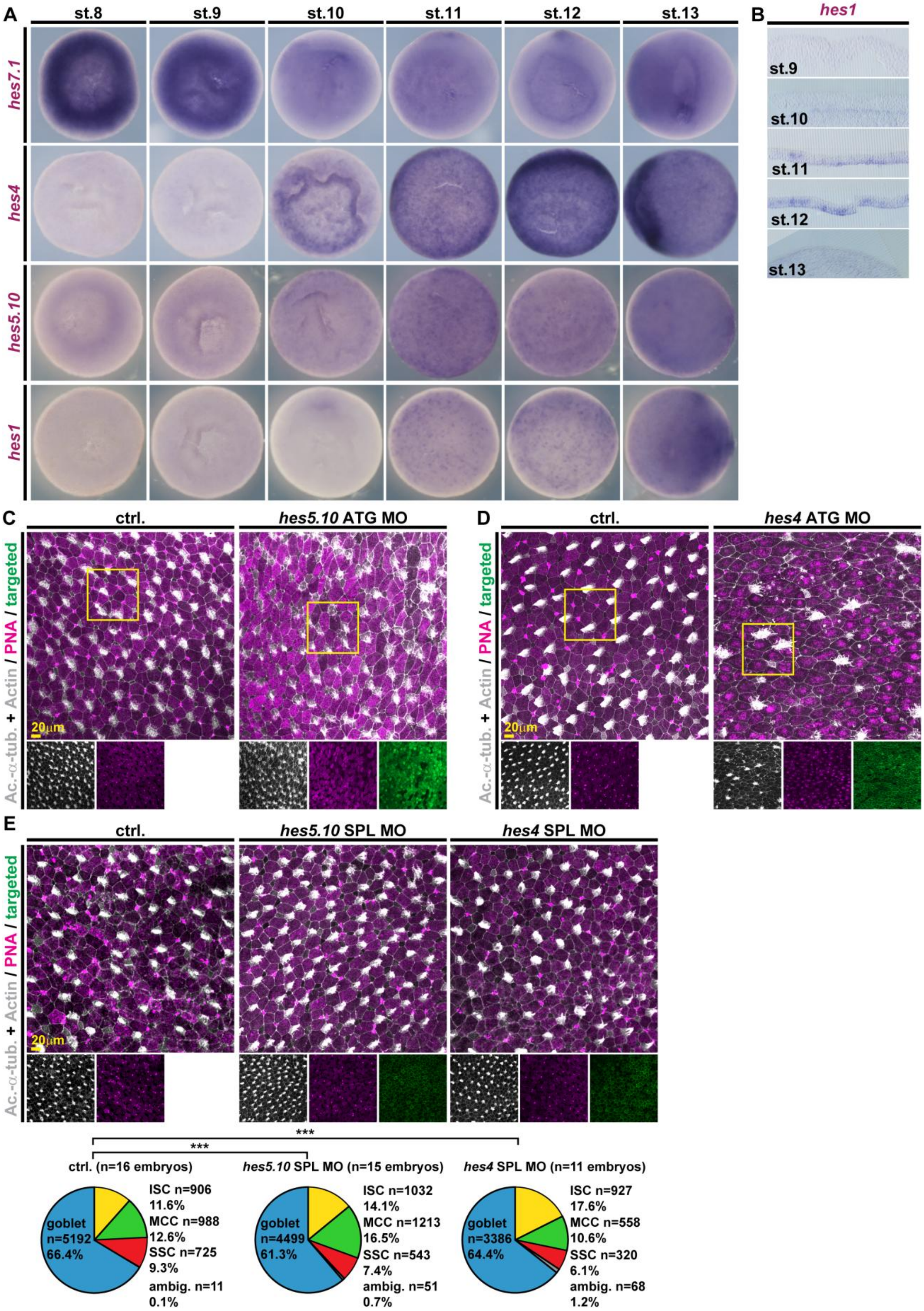

**Supplementary Figure S8: Expression and function of Hes transcription factors in the *Xenopus* epidermis**

**A,B:** Animal (st. 9 - 12) and ventral (st. 13) views, WMISH for *hes7.1*, *hes4*, *hes5.10* and *hes1* transcripts across stages of mucociliary cell fate specification. **B:** Epidermis sections, WMISH for *hes1* transcripts from st. 8 to st. 13. Related to Fig. 3B. **C-E:** Immunofluorescent images of epidermal cell types after indicated manipulations at st. 32. Acetylated-α-tubulin (Ac.-α-tub., grey) marks MCC cilia; F-actin (Actin, grey) marks cell borders and apical morphology; peanut agglutinin (PNA, magenta) marks small mucus granules in goblet cells and large secretory vesicles in SSCs; membrane-GFP (targeted, green) marks targeted cells. **C:** ATG MO-mediated knockdown of *hes5.10* (*hes5.10* ATG MO). Areas depicted in Fig. 3C are indicated by yellow boxes. **D:** ATG MO-mediated knockdown of *hes4* (*hes4* ATG MO). Areas depicted in Fig. 3C are indicated by yellow boxes. **E:** Splice-inhibiting MO-mediated knockdown of *hes5.10* (*hes5.10* SPL MO) or *hes4* (*hes4* SPL MO) and quantification of cell types (bottom). $\chi^2$-test, *** = $p < 0.001$.

Brislinger, Hansen et al. Suppl Figure 9

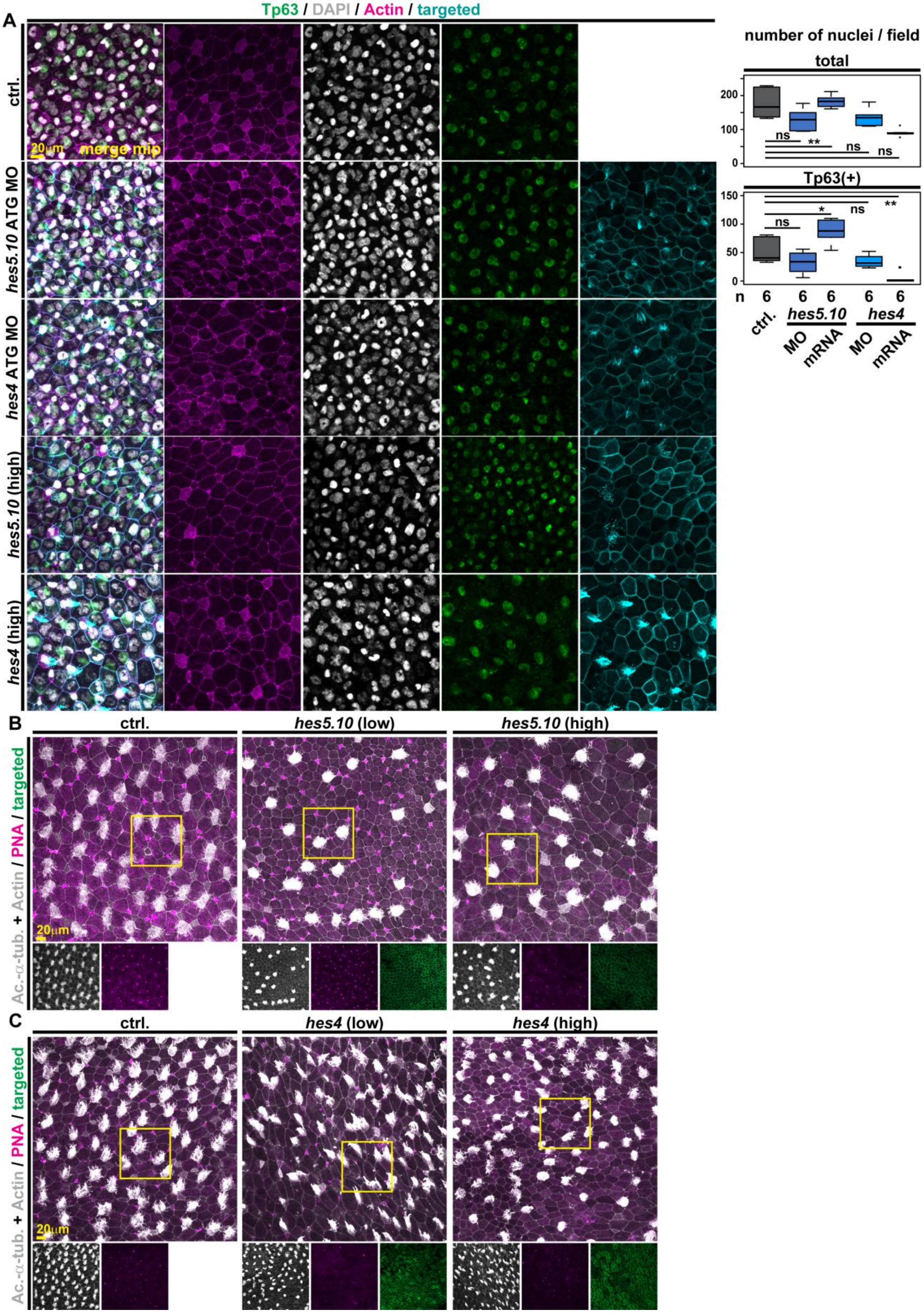

**Supplementary Figure S9: Hes4 and Hes5.10 functions in the *Xenopus* epidermis**
**A:** IF of epidermal cells in controls (ctrl.) and after ATG MO-mediated knockdown of *hes5.10* (*hes5.10* ATG MO) or *hes4* (*hes4* ATG MO), as well as after overexpression of high levels (50 ng/ul) of *hes5.10* or *hes4* mRNA at st. 32. Tp63 (green) marks basal cells (BCs) in the deep layer; F-actin (Actin, magenta) marks cell borders and apical morphology; DAPI (grey) marks nuclei; membrane-GFP (targeted, cyan) marks targeted cells. Graphs on the right: Quantification of total cells and of Tp63(+) BCs per embryo/field. Mann-Whitney-test, ns = $p > 0.05$; * = $p < 0.05$; ** = $p < 0.01$. **B,C:** Immunofluorescent images of epidermal cell types after indicated manipulations at st. 32. Acetylated-$\alpha$-tubulin (Ac.-$\alpha$-tub., grey) marks MCC cilia; F-actin (Actin, grey) marks cell borders and apical morphology; peanut agglutinin (PNA, magenta) marks small mucus granules in goblet cells and large secretory vesicles in SSCs; membrane-GFP (targeted, green) marks targeted cells. **B:** mRNA overexpression of *hes5.10* at low (15 ng/ul) and high (50 ng/ul) concentrations. Areas depicted in Fig. 3F are indicated by yellow boxes. **C:** mRNA overexpression of *hes4* at low (15 ng/ul) and high (50 ng/ul) concentrations. Areas depicted in Fig. 3G are indicated by yellow boxes.

**Brislinger, Hansen et al. Figure S10**

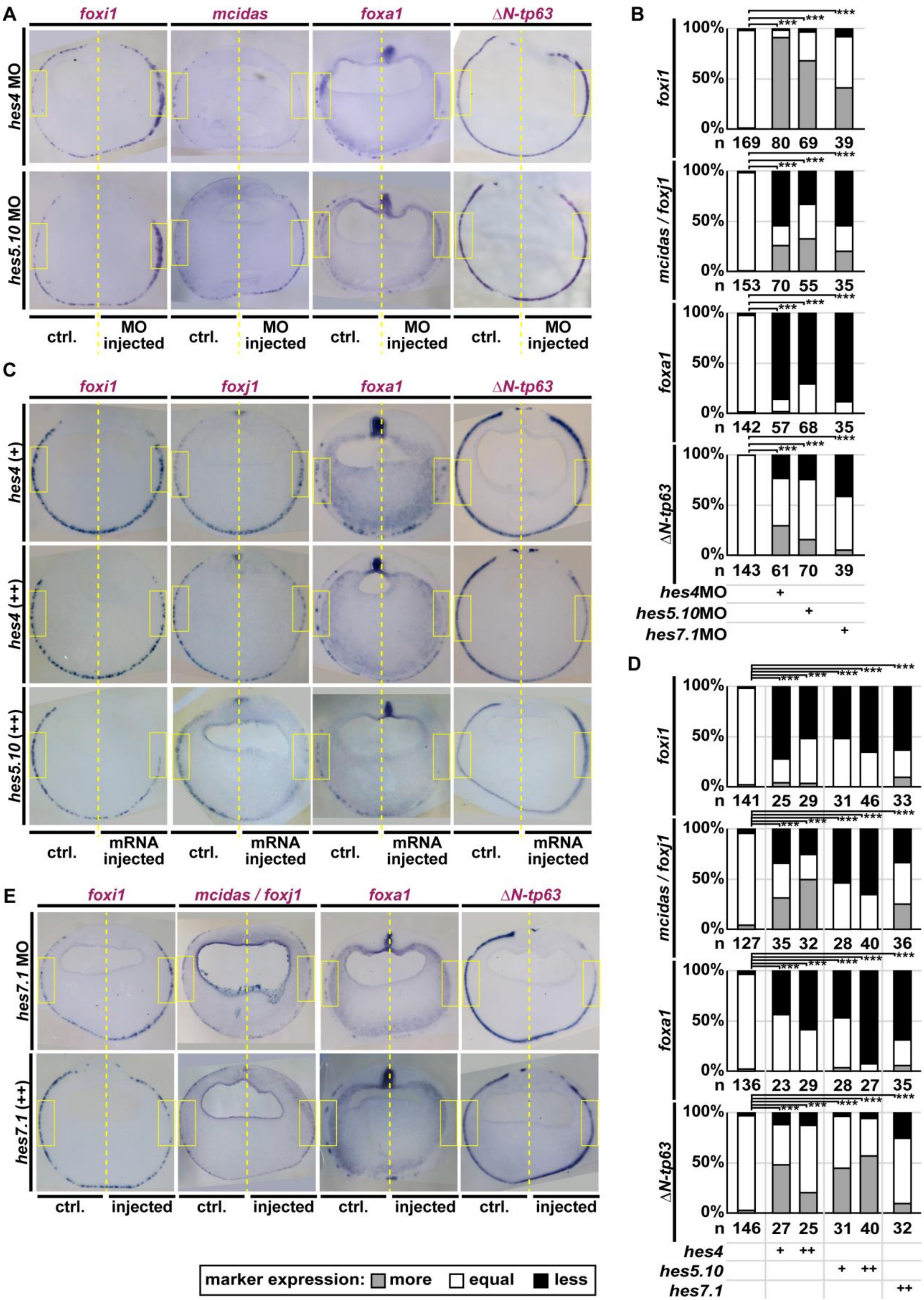

**Supplementary Figure S10: Hes transcription factors regulate cell fates assessed at the end of cell fate specification**

**A,C,E:** Sections, WMISH for indicated transcripts at st. 16/17 in controls and after *hes* manipulations. +=low dose, ++= high dose of mRNA overexpression of *hes* genes. Sections of embryos allow detailed comparison between injected and uninjected control (ctrl.) sides of the embryo. Yellow boxes indicate targeted and control areas. **B,D:** Quantification of results depicted in A, C and E. Expression levels were scored more, equal, less expression on the injected vs. uninjected side. $X^2$-test, *** = $p < 0.001$.

**Brislinger, Hansen et al. Suppl Figure 11**

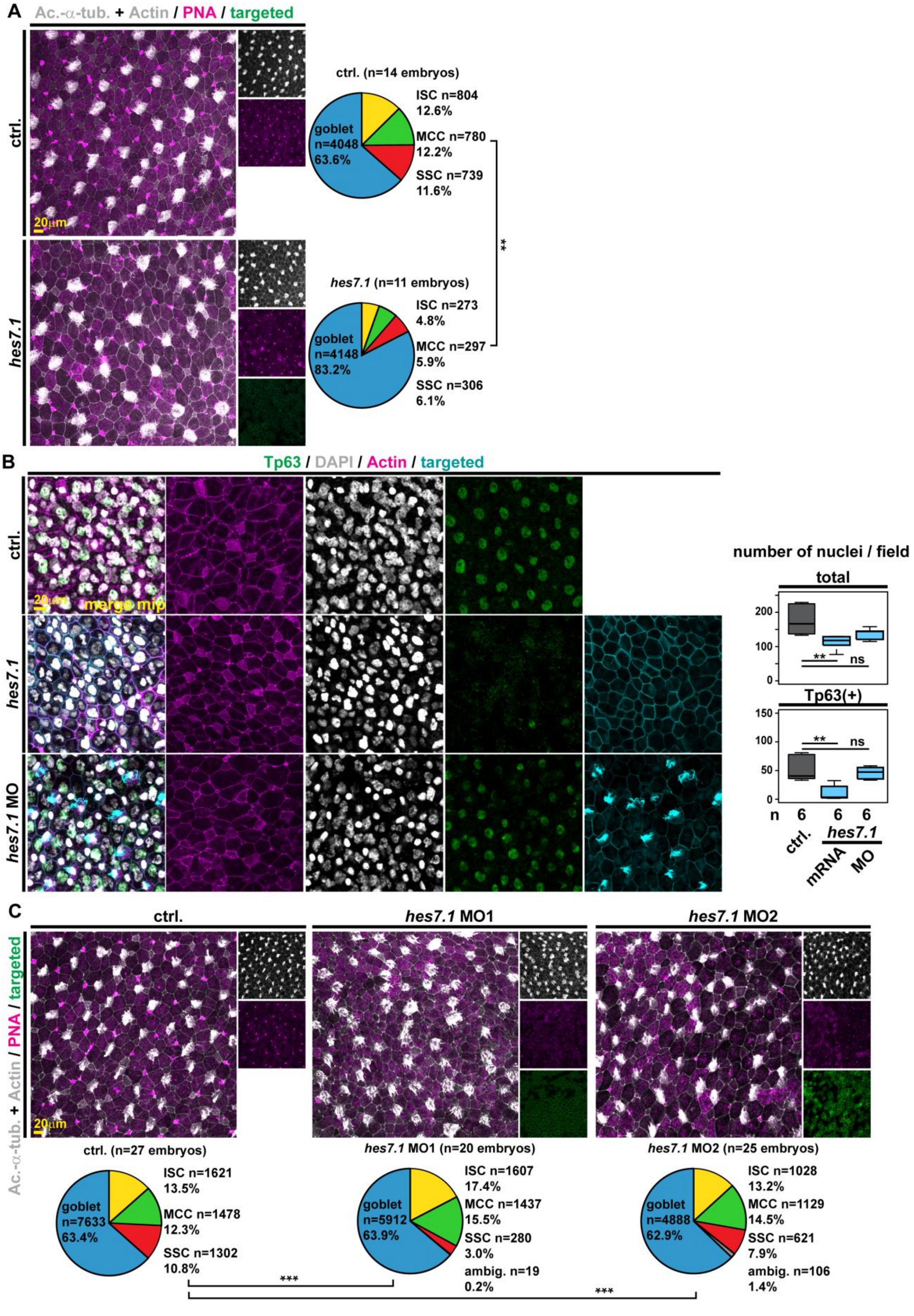

**Supplementary Figure S11: *hes7.1* functions in the *Xenopus* epidermis**

**A,C:** Immunofluorescent images of epidermal cell types after indicated manipulations at st. 32 and quantification of results. Acetylated-$\alpha$-tubulin (Ac.-$\alpha$-tub., grey) marks MCC cilia; F-actin (Actin, grey) marks cell borders and apical morphology; peanut agglutinin (PNA, magenta) marks small mucus granules in goblet cells and large secretory vesicles in SSCs; membrane-GFP (targeted, green) marks targeted cells. $X^2$-test, ** = $p < 0.01$, *** = $p < 0.001$. **A:** mRNA overexpression of *hes7.1* at high (50 ng/ul) concentrations. **C:** ATG MO-mediated knockdown of *hes7.1* using two non-overlapping ATG MOs (*hes7.1* MO1 and MO2). **B:** IF of epidermal cells in controls (ctrl.) and after ATG MO1-mediated knockdown of *hes7.1* (*hes7.1* MO) or after overexpression of high levels (50 ng/ul) of *hes7.1* mRNA at st. 32. Tp63 (green) marks basal cells (BCs) in the deep layer; F-actin (Actin, magenta) marks cell borders and apical morphology; DAPI (grey) marks nuclei; membrane-GFP (targeted, cyan) marks targeted cells. Graphs on the right: Quantification of total cells and of Tp63(+) BCs per embryo/field. Mann-Whitney-test, ns = $p > 0.05$; ** = $p < 0.01$.

### Brislinger, Hansen et al. Suppl Figure 12

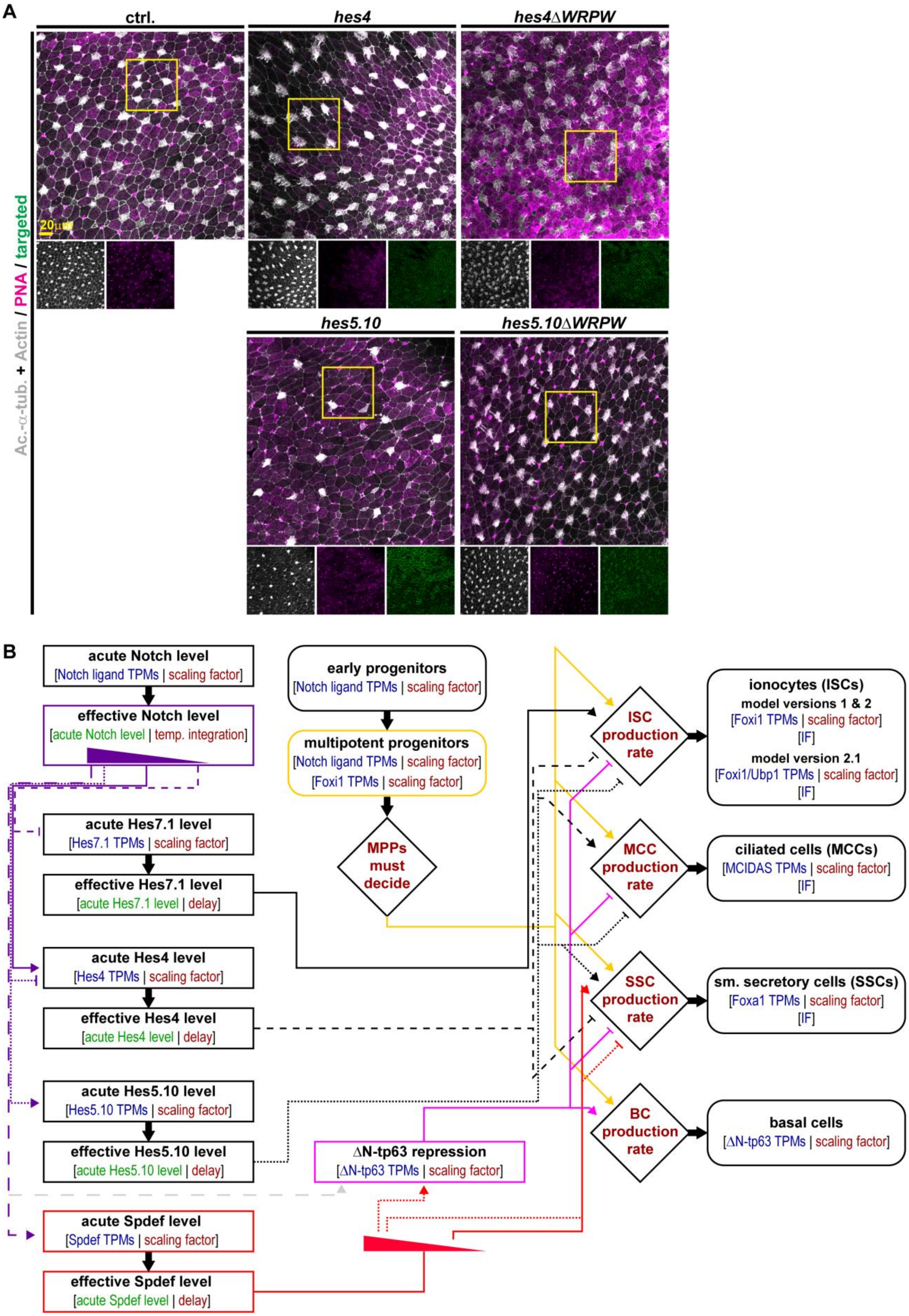

**Supplementary Figure S12: Effects of WRPW-deletion constructs and structure of mathematical model**

**A:** Immunofluorescent images of epidermal cell types after mRNA overexpression of full-length *hes4* or *hes5.10* or WRPW-deletions (*hes4ΔWPRW* or *hes5.10 ΔWPRW*) at 50 ng/ul at st. 32. Acetylated-α-tubulin (Ac.-α-tub., grey) marks MCC cilia; F-actin (Actin, grey) marks cell borders and apical morphology; peanut agglutinin (PNA, magenta) marks small mucus granules in goblet cells and large secretory vesicles in SSCs; membrane-GFP (targeted, green) marks targeted cells. Areas depicted in Fig. 4A are indicated by yellow boxes. **B:** Schematic representation of data and connections of parameters/functions throughout model 2.1. Details on variables and factors in the model are provided in Tables S3,4,5 and Supplementary Document S1. Arrowheads = activation; T-ends = inhibition; parameters in blue = measured data from RNA-seq (in TPM) or by IF (in N); parameters in green = derived from within the model and used for subsequent calculations; parameters in red = derived from fitting the model to data. Model 1 does not include Spdef regulation (bright red boxes and lines). Instead, high Notch levels act directly on ΔN-tp63 (grey dashed-line). In model 2.1, a combination of Foxi1 and Ubp1 levels was used to model ISC production.

**Brislinger, Hansen et al. Suppl Figure 13**

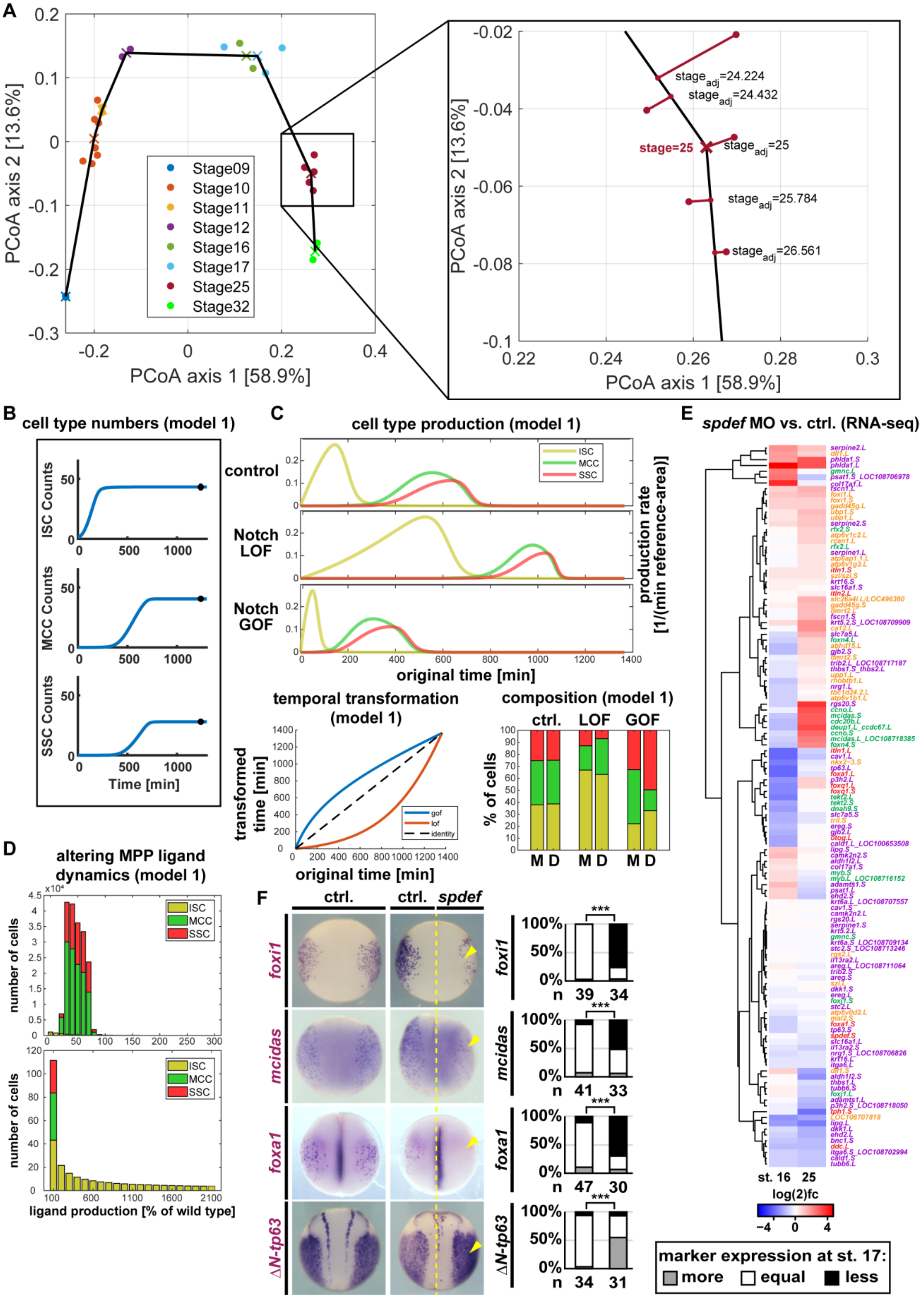

**Supplementary Figure S13: Mathematical modeling of mucociliary cell fate specification - Model1 and Spdef**

**A:** Fine-staging of temporal-resolved RNA-seq datasets. To reduce temporal bias, RNA-seq data were reduced to two dimensions using Principal Coordinate Analysis (PCoA). Here, Bray-Curtis dissimilarity was used as a measure of distance. Because we assumed that not all transcripts are regulated over time, we performed PCoA on 10,000 transcripts that exhibited the greatest variability across 25 sequencing samples from datasets 1 and 2 as well as from Haas et al.. Linear interpolation between the mean positions of the samples at the same stage was used to derive the trajectory of transcriptional regulation in this low-dimensional representation. **B-D:** Results from Model 1. **B:** Cell type numbers. **C:** Upper panels show effects of Notch manipulations on cell type production rates. LOF = loss-of-function (= *suh-dbm*); GOF = gain-of-function (= *nicd*). Lower panels show the temporal transformation (left) and results from the model (M) compared to experimental data (D). **D:** Modeling of how changes of ligand levels expression from MPPs would affect cell type composition. Upper panel shows 100 (% = normal) and modulations in steps of + (right) and - (left) 10 % from normal in the range from 0 % - 300 %. Lower panel 100 (% = normal) left and modulations in steps of 100 % from normal in the range up to 2200 % right. **E:** Heatmaps (log(2) fold-change) of RNA-seq comparing sets of marker genes for ISCs (yellow), MCCs (green), SSCs/secretory cells (red) and BCs (purple) after knockdown of *spdef* (*spdef* MO) vs. control (ctrl.). (n = 2 per stage). **F:** WMISH for *foxi1*, *mcidas*, *foxa1* and *ΔN-tp63* transcripts in control and unilaterally *spdef* mRNA (100ng/ul) injected st. 17/18 embryos. Left panels, dorsal views, manipulated side is indicated. Right panels show quantification of results. Marker expression was assessed relative to the uninjected control side and is shown as more, equal or less than the control side. $X^2$-test, *** = $p < 0.001$.

### Brislinger, Hansen et al. Suppl Figure 14

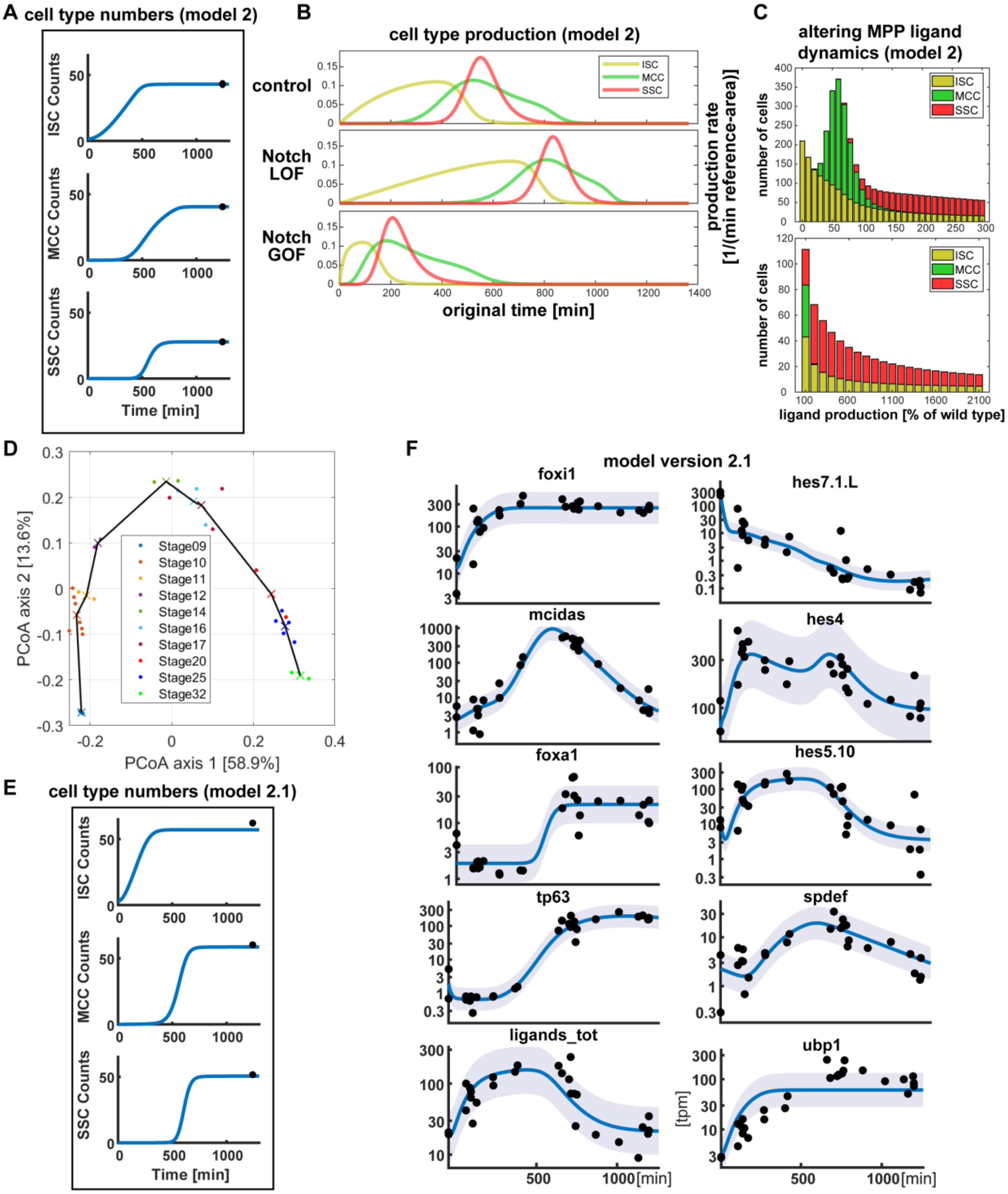

**Supplementary Figure S14: Mathematical modeling of mucociliary cell fate specification - Model2 and 2.1**

**A-C:** Results from Model 2. **A:** Cell type numbers. **B:** Effects of Notch manipulations on cell type production rates. LOF = loss-of-function (= *suh-dbm*); GOF = gain-of-function (= *nicd*). **C:** Modeling of how changes of ligand levels expression from MPPs would affect cell type composition. Upper panel shows 100 (% = normal) and modulations in steps of + (right) and - (left) 10 % from normal in the range from 0 % - 300 %. Lower panel 100 (% = normal) left and modulations in steps of 100 % from normal in the range up to 2200 % right. **D-F:** Additional data and results from Model 2.1. **D:** Fine-staging of temporal-resolved RNA-seq datasets. To reduce temporal bias, RNA-seq data were reduced to two dimensions using Principal Coordinate Analysis (PCoA). Here, Bray-Curtis dissimilarity was used as a measure of distance. Because we assumed that not all transcripts are regulated over time, we performed PCoA on 10,000 transcripts that exhibited the greatest variability across 30 sequencing samples from datasets 1, 2 and 3 as well as from Haas et al.. **E:** Cell type numbers. **F:** Visualization of fit (blue line) to data from RNA-seq (black dots). Light blue area represents the error model that describes the measurement noise.

**Brislinger, Hansen et al. Suppl Figure 15**

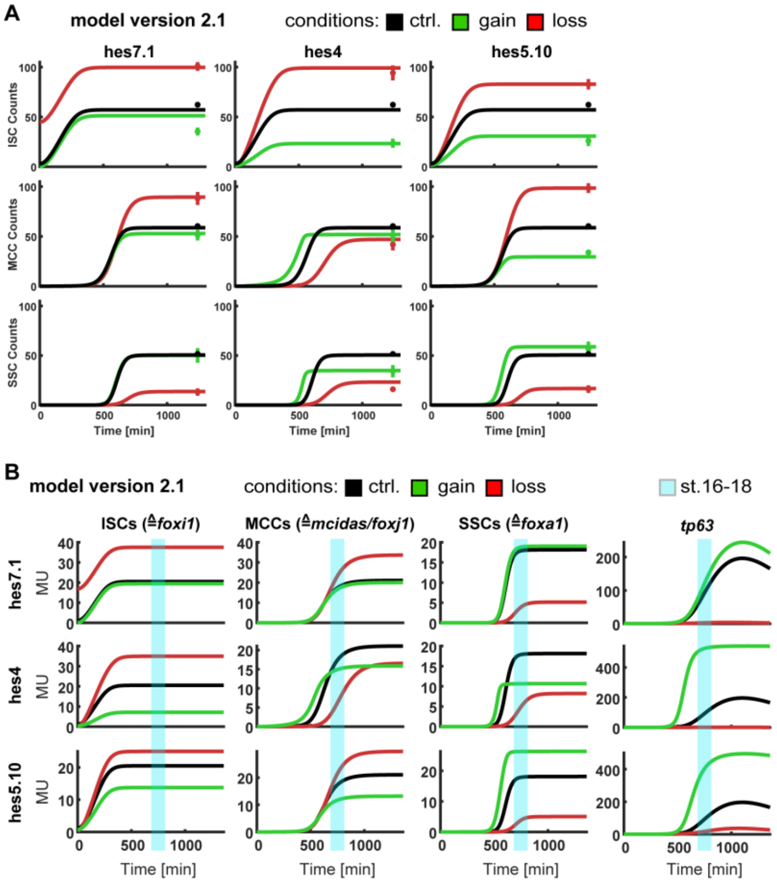

**C**

**comparison of model version 2.1 preditions and experimental results (cf. Fig. S10)**

| Condition | Marker | Model | WMISH |
|---|---|---|---|
| hes7.1 LOF | foxi1 | More | More |
| hes7.1 LOF | mcidas/foxj1 | More | Less |
| hes7.1 LOF | foxa1 | Less | Less |
| hes7.1 LOF | tp63 | Less | Less |
| hes4 LOF | foxi1 | More | More |
| hes4 LOF | mcidas/foxj1 | Less | Less |
| hes4 LOF | foxa1 | Less | Less |
| hes4 LOF | tp63 | Less | Neurtral |
| hes5.10 LOF | foxi1 | More | More |
| hes5.10 LOF | mcidas/foxj1 | More | More |
| hes5.10 LOF | foxa1 | Less | Less |
| hes5.10 LOF | tp63 | Less | Neutral |

| Condition | Marker | Model | WMISH (low GOF) |
|---|---|---|---|
| hes7.1 GOF | foxi1 | Neutral | Less |
| hes7.1 GOF | mcidas/foxj1 | Neutral | Neutral/Less |
| hes7.1 GOF | foxa1 | Neutral | Less |
| hes7.1 GOF | tp63 | Neutral/More | Less |
| hes4 GOF | foxi1 | Less | Less |
| hes4 GOF | mcidas/foxj1 | Neutral/Less | Neutral/More |
| hes4 GOF | foxa1 | Less | Less |
| hes4 GOF | tp63 | More | Less |
| hes5.10 GOF | foxi1 | Less | Less |
| hes5.10 GOF | mcidas/foxj1 | Less | Less |
| hes5.10 GOF | foxa1 | More | Less |
| hes5.10 GOF | tp63 | More | More |

model predition correct?
yes no

**Supplementary Figure S15: Mathematical modeling and model 2.1 predictions**

**A:** Modeling of cell numbers in controls (black line) and hes gain (green line) as well as loss (red line) of function manipulations using model version 2.1. **B:** Modeling of cell type production (marker equivalents) over time in controls (black line) and hes gain (green line) as well as loss (red line) of function manipulations using model version 2.1. The time range equivalent to st. 16-18 (end of specification) is highlighted in light blue. **C:** Trends in gene expression derived from plots in B were used for comparison with experimental results shown in Fig. S10 (WMISH).

| Sample | Stage | Stage adjusted |
|---|---|---|
| Exp3_St9-B | 9 | 8.997 |
| Exp3_St9-A | 9 | 9.003 |
| Exp1_E1S10Ctrl | 10 | 9.844 |
| Exp3_St10_5-B | 10 | 9.846 |
| Exp1_E2S10Ctrl | 10 | 9.908 |
| Exp3_St10_5-A | 10 | 9.947 |
| Exp2_st10_A1 | 10 | 10.542 |
| Exp2_st10_G1 | 10 | 10.589 |
| Exp3_St11_5-B | 11 | 10.921 |
| Exp3_St11_5-A | 11 | 11.034 |
| Exp2_st10_D1 | 10 | 11.078 |
| Exp3_St12_5-B | 12 | 11.909 |
| Exp3_St12_5-A | 12 | 12.132 |
| Exp2_st16-19_D2 | 17 | 15.256 |
| Exp1_E1S16Ctrl | 16 | 15.772 |
| Exp1_E2S16Ctrl | 16 | 17.432 |
| Exp2_st16-19_A2 | 17 | 17.64 |
| Exp2_st16-19_G2 | 17 | 18.184 |
| Exp2_st25_A3 | 25 | 24.224 |
| Exp1_E1S25Ctrl | 25 | 24.432 |
| Exp2_st25_G3 | 25 | 25 |
| Exp2_st25_D3 | 25 | 25.784 |
| Exp1_E2S25Ctrl | 25 | 26.561 |
| Exp1_E1S32Ctrl | 32 | 31.279 |
| Exp1_E2S32Ctrl | 32 | 32.721 |

**Table S1:** Fine-staging of RNA-seq data model 1 and 2 (datasets 1,2 and from Haas et al.).

| Observable | Model 1<br>-2 log Likelihood | Model 2<br>-2 log Likelihood | Improvement<br>Difference |
|---|---|---|---|
| foxi1 | 13.15 | 10.53 | 2.62 |
| mcidas | 15.80 | 16.65 | -0.85 |
| foxa1 | 14.75 | 15.03 | -0.28 |
| ligands_tot | 28.15 | 10.85 | 17.30 |
| hes4 | -10.17 | -13.16 | 2.99 |
| hes5 | 62.51 | 46.16 | 16.35 |
| hes7 | 48.92 | 45.31 | 3.62 |
| tp63 | 15.89 | 10.24 | 5.65 |
| spdef | - | 25.29 | - |
| isc_abs | 3.98 | 3.98 | 0.00 |
| mcc_abs | 3.39 | 3.39 | 0.00 |
| ssc_abs | 3.01 | 3.01 | 0.00 |
| **Sum** | **199.37** | **177.26** | **22.11** |

**Table S2:** Comparison of likelihood between models 1 and 2.

| Sample | Stage | Stage adjusted |
|---|---|---|
| st9A | 9 | 9.0065 |
| st9B | 9 | 8.9935 |
| st10_5A | 10 | 10.211 |
| st10_5B | 10 | 10.1495 |
| E1_10_5 | 10 | 9.9215 |
| E2_10_5 | 10 | 9.9285 |
| st10_5A1 | 10 | 10.2065 |
| st10_5D1 | 10 | 10.5335 |
| st10_5G1 | 10 | 10.3145 |
| st11_5A | 11 | 11.2635 |
| st11_5B | 11 | 11.25 |
| st12_5A | 12 | 12.301 |
| st12_5B | 12 | 12.2075 |
| S3E1_st14 | 14 | 14.297 |
| S3E2_st14 | 14 | 13.892 |
| E1_16 | 16 | 15.44 |
| E2_16 | 16 | 16.728 |
| st16D2 | 17 | 15.829 |
| S3E1_st16 | 16 | 15.892 |
| st19A2 | 17 | 17.0375 |
| st19G2 | 17 | 17.3345 |
| S3E1_st20 | 20 | 21.975 |
| S3E2_st20 | 20 | 19.637 |
| E1_25 | 25 | 24.3825 |
| E2_25 | 25 | 26.253 |
| st25A3 | 25 | 23.9375 |
| st25D3 | 25 | 25.434 |
| st25G3 | 25 | 24.9175 |

**Table S3:** Fine-staging of RNA-seq data model 2.1 (datasets 1,2, 3 and from Haas et al.).

| Nr. | States for molecules (cf. report) | Biological meaning (cf. model flow chart) |
|---|---|---|
| 1 | lig | acute Notch level (ligand RNA) |
| 2 | Lig | effective Notch level (NICD protein) |
| 3 | hes4 | acute Hes4 level (RNA) |
| 4 | hes5 | acute Hes5 level (RNA) |
| 5 | hes7 | acute Hes7.1 level (RNA) |
| 6 | hes4_d1 | auxiliary delay chain state without direct biological interpretation |
| 7 | hes5_d1 | auxiliary delay chain state without direct biological interpretation |
| 8 | hes7_d1 | auxiliary delay chain state without direct biological interpretation |
| 9 | hes4_d2 | auxiliary delay chain state without direct biological interpretation |
| 10 | hes5_d2 | auxiliary delay chain state without direct biological interpretation |
| 11 | hes7_d2 | auxiliary delay chain state without direct biological interpretation |
| 12 | hes4_d3 | auxiliary delay chain state without direct biological interpretation |
| 13 | hes5_d3 | auxiliary delay chain state without direct biological interpretation |
| 14 | hes7_d3 | auxiliary delay chain state without direct biological interpretation |
| 15 | hes4_d | effective Hes4 level (protein) |
| 16 | hes5_d | effective Hes5 level (protein) |
| 17 | hes7_d | effective Hes7.1 level (protein) |
| 18 | spdef | acute level (RNA) |
| 19 | Spdef | effective Spdef level (protein) |
| 20 | tp63 | ΔN-tp63 repression (RNA) |

**Table S4:** States representing molecule abundances for all models.

| Nr. | States for cell types (cf. report) | Biological meaning (cf. model flow chart) |
|---|---|---|
| 1 | ep | early progenitors (EPs) |
| 2 | mpp | multipotent progenitors (MPPs) |
| 3 | bc | basal cells (BCs) |
| 4 | isc | ionocytes (ISCs) |
| 5 | mcc | ciliated cells (MCCs) that express mcidas |
| 6 | mcc_late | ciliated cells (MCCs) that do not express mcidas |
| 7 | ssc | small secretory cells (SSCs) |

**Table S5:** States representing cell abundances for all models.

| Parameter pattern (cf. report) | Biological meaning |
|---|---|
| d_{x} | degradation rate of {x} |
| hes_delay | average time needed for translation from acute Hes* level to effective Hes* level |
| h_{x}_a_{y} | Hill coefficient for activation of {x} by {y} |
| h_{x}_i_{y} | Hill coefficient for inhibition of {x} by {y} |
| h_{x}_a | Hill coefficient for activation of {x} by effective Notch level |
| h_{x}_i | Hill coefficient for inhibition of {x} by effective Notch level |
| init_{x} | Initial value of dynamical state or observable {x}. Depending on the variable, it has to be multiplied with a scale parameter to get initial data value on the biological scale |
| k_cell_i | Hill threshold for inhibition of ISCs, MCCs and SSC by ΔN-tp63 repression |
| k_isc_a | Hill threshold for activation of ISCs by hes7 |
| k_mcc_a | Hill threshold for activation of MCCs by hes4 |
| k_ssc_a | Hill threshold for activation of SSCs by hes5 |
| k_{x}_a_{y}_delta | Hill threshold for activation of {x} by {y} measured from the steady state of {y} |
| k_{x}_i_{y}_delta | Hill threshold for inhibition of {x} by {y} measured from the steady state of {y} |
| p_mcc_late | transition rate from early to late MCC state, reciprocal of mean dwell time in early MCC state |
| p_lig_ep | cell specific ligand production rate of EPs (corresponds to initial Notch signaling levels) |
| p_lig_mpp_factor | MPPs produce p_lig_mpp_factor times as much ligands as EPs (>1) |
| p_lig_bc_factor | BCs produce p_lig_bc_factor times as much ligands as MPPs (<1) |
| p_lig_isc_factor | ISCs produce p_lig_isc_factor times as much ligands as MPPs (<1) |
| p_lig_mcc_factor | MCCs produce p_lig_mcc_factor times as much ligands as MPPs (<1) |
| p_lig_ssc_factor | SSCs produce p_lig_ssc_factor times as much ligands as MPPs (<1) |
| p_mpp | transition rate from EPs to MPPs |
| p_Spdef | production rate of effective Spdef level |
| p_Lig | production rate of effective Notch level |
| v_max | overall factor in<br>ISC, MCC and SSC production rates |
| v_max_bc | overall factor in BC production rates |
| s_{x} | scaling parameter / proportionality factor from dynamical state to observable for species {x} |
| sd_tpm_{x} | error parameter for constant error model for tpm values of {x} |

**Table S6:** Parameters in model 1 and 2.

| Parameter pattern (cf. report) | Biological meaning |
|---|---|
| d_{x} | degradation rate of {x} |
| hes_delay | average time needed for translation from acute Hes* level to effective Hes* level |
| h_{x}_a_{y} | Hill coefficient for activation of {x} by {y} |
| h_{x}_i_{y} | Hill coefficient for inhibition of {x} by {y} |
| h_{x}_a | Hill coefficient for activation of {x} by effective Notch level |
| h_{x}_i | Hill coefficient for inhibition of {x} by effective Notch level |
| init_{x} | Initial value of dynamical state or observable {x}. Depending on the variable, it has to be multiplied with a scale parameter to get initial data value on the biological scale |
| k_cell_i | Hill threshold for inhibition of ISCs, MCCs and SSC by ΔN-tp63 repression |
| k_isc_a | Hill threshold for activation of ISCs by hes7 |
| k_mcc_a | Hill threshold for activation of MCCs by hes4 |
| k_ssc_a | Hill threshold for activation of SSCs by hes5 |
| k_{x}_a_{y}_fc | Hill threshold for activation of {x} by {y} measured as a ratio to the initial state of {y}. |
| k_{x}_i_{y}_fc | Hill threshold for inhibition of {x} by {y} measured as ratio to the initial state of {y}. |
| p_mcc_late | transition rate from early to late MCC state, reciprocal of mean dwell time in early MCC state |
| p_lig_ep_fc | EPs produce p_lig_ep_fc times as much ligands as MPPs (<1) |
| p_lig_bc_fc | BCs produce p_lig_bc_fc times as much ligands as MPPs (<1) |
| p_lig_isc_fc | ISCs produce p_lig_isc_fc times as much ligands as MPPs (<1) |
| p_lig_mcc_fc | MCCs produce p_lig_mcc_fc times as much ligands as MPPs (<1) |
| p_lig_ssc_fc | SSCs produce p_lig_ssc_fc times as much ligands as MPPs (<1) |
| p_mpp | transition rate from EPs to MPPs |
| p_Spdef | Production rate of effective Spdef level |
| p_Lig | Production rate of effective Notch level |
| prior_p_{x} | Production rate prior for RNA component {x} to restrict model to biologically reasonable behavior. These are not fitted. |
| reg_p_{x} | Penalized fold-change of estimated model production rate to production rate prior |
| reg_{x} | Penalized fold-change of scale parameters for RNA component {x}. The difference in scale should be small, thus strong deviations are penalized. |
| v_max_cell | Mean Maximal ISC, MCC and SSC production rates |
| vmax_{cell}_fc | Fold change of maximal production rate for the different cell types {cell}. This parameter is strongly penalized |
| v_max_bc | Overall factor in BC production rates |
| s_{x} | Scaling parameter / proportionality factor from dynamical state to observable for species {x} |
| s_{x}_fc | Scale if defined as fold-change to different scale |
| sd_RNA | Error parameter for constant error model for tpm values |
| sd_cell | Error Parameter for constant error model for all cell count data |

| | |
|---|---|
| transl_{hes}_lof_fc | Loss of function fold-change for the translation rate of {hes} in {hes} knockdown experiments compared to the wildtype condition |
| init_{hes}_gof_lc_fc | Gain of function fold-change for the initial amount of {hes} RNA compared to the wildtype condition |
| init_{cell}_{P}_{hes}_fc | Change of {cell} initial in the {hes} gain/loss of function ({P}) experiment compared to the wildtype condition. This parameter is penalized to force biologically reasonable solutions. |

**Table S7:** Parameters in model 2.1. Compared to previous model versions, there were some reparametrizations and parameters for the loss of function/gain of function experiments were added. The parameters in red are the critical additions from model 2.1. compared to model 2 to allow the model to describe the gain/loss of function experiments.

| stage | physical time [min] |
|---|---|
| 9 | 0 |
| 10 | 117 |
| 10.5 | 169 |
| 11 | 238 |
| 11.5 | 313 |
| 12 | 377 |
| 12.5 | 461 |
| 13 | 624 |
| 14 | 688 |
| 15 | 735 |
| 18 | 811 |
| 25 | 1240.3 |

**Table S8:** Calculating physical time from stages

| Nr. | Observables (cf. report) | Biological meaning (cf. model flow chart) |
|---|---|---|
| 1 | foxi1 | Foxi1 TPMs |
| 2 | mcidas | MCIDAS TPMs |
| 3 | foxa1 | Foxa1 TPMs |
| 4 | lig | Foxi1 TPMs |
| 5 | hes4 | Hes4 TPMs |
| 6 | hes5 | Hes5.10 TPMs |
| 7 | hes7 | Hes7.1 TPMs |
| 8 | tp63 | ΔN-tp63 TPMs |
| 9 | spdef | Spdef TPMs |
| 10 | isc_abs | ISCs (number of cells, IF) |
| 11 | mcc_abs | MCCs (number of cells, IF) |
| 12 | ssc_abs | SSCs (number of cells, IF) |
| 13 | ubp1 | ubp1 TPMs |

**Table S9:** Observables in model 1 and 2. Ubp1 was added in model 2.1 as a marker for ISCs.

### Mathematical modeling

A Notch signaling ODE model for cell fate specification in mucociliary tissue

Our final ordinary differential equation (ODE) model (model 2) consists of 27 dynamical states, 38 reactions, 12 observables, 209 data points and 98 fitted parameters. Model 2.1 consists of 27 dynamical states, 37 reactions, 13 observables, 757 data points and 127 fitted parameters. In the following, it is explained what these quantities represent and how they are linked to each other in the model.

Dynamical states: Molecules and cells

The ODE model elucidates the influence of different molecular compounds on the cell type composition. Thus, the model acts on two different scales: the molecular scale on the one hand, the cellular scale on the other. Therefore, unlike most ODE models in systems biology, the 27 dynamical states only partly represent the abundance of molecular compounds (**Table S4**). Instead, 7 of them represent cell type abundances (**Table S5**).

Time evolution of the states

The rate equations, which govern the time evolution of system, can be divided into 3 categories:

1. Rate equations that represent cell state transitions (for simplicity, we still call them "reactions"): Initially, (nearly) all cells start as early progenitors (EPs). In a first step, these early progenitors are assumed to become multipotent progenitors (MPPs) at a constant rate, i.e. independent from any molecular abundances. This is different in the second step: MPPs themselves turn into ionocytes (ISCs), ciliated cells (MCCs), small secretory cells (SSCs) or basal cells (BCs) based on ΔN-tp63 repression and the effective levels of Hes4, Hes5, Hes7.1 and Spdef. The production rates of ISCs, MCCs, SSCs and BCs are governed by hill equations, where each hill equation represents an activating or inhibiting effect from one of the mentioned transcription factors. The precise activations/inhibitions assumed in the model are depicted in the model flow chart (**Fig.S12C**). Most transcription factors act on a cell type either only activating or only inhibiting. The only exception is the effective Spdef level: It activates SSCs at lower levels but inhibits SSCs at higher levels. It was presumed that MPPs have to decide on a cell fate. This is ensured by a prior, that penalizes high MPP abundances compared to BC abundances at a late time point. For MCCs, two different dynamical states are introduced, because the related cell marker *mcidas* is only transiently expressed. The transition from the MCC marker expressing state to the later MCC state is assumed at a constant rate, i.e. independently of transcription factors represented in the model.

2. Rate equations that represent production and degradation of molecules: All the dynamical states representing molecular abundances have a production rate and a degradation rate. The degradation rates are assumed to be proportional to the abundances of all molecular states with individual rate constants. In contrast, the production rate depends on either the molecular abundances or on the cell type composition:

   a.) For all the dynamical states representing RNA except the acute Notch level, the production rates depend on other molecular abundances. The precise activation/inhibition connections between the transcription factors are depicted in the model flow chart (**Fig. S12B**). Hes4 is activated for low and inhibited for high effective Notch levels. This is reflected by a product of two hill equations at the according production rate. All others are given by a single hill equation. For all the dynamical states representing protein abundances, the production rates depend linearly or via a delay chain of length n=4 on the related RNA state.

   b.) The production rate for the acute Notch level depends on the cell composition. It is defined as the weighted mean over the production rates of the individual cell types. The weights are given by the cell type proportions, i.e. the cell type composition. The starting point of modeling the whole specification process is the transition from EPs to MPPs and the resulting change in cell composition and acute Notch level production. All other transcription factors are direct or intermediate targets of the acute Notch level.

<u>Model Parametrization</u>

The model uses a set of model parameters to generate the dynamics based on an ordinary differential equation model implied by the model scheme in Figure S9C. The model parameters (for models 1 and 2: **Table S6**; for model 2.1: **Table S7**) are chosen though numerical optimization such that the model predictions follow the observed data as closely as possible. In order to restrict the choice of possible parameter combinations to be biologically reasonable and feasible for numerical optimization, some parameters were regularized by quadratic priors and steady states were used within the delay chain.

<u>Embryonic stage representation in the model</u>

The mapping between development stages and physical time in minutes was implemented according to Xenbase.org (http://www.xenbase.org/anatomy/static/xenopustimetemp.jsp). In particular, the values for *Xenopus laevis*, 22°C, average were used. Time point zero (physical time) was chosen to be at stage 9. Stages, where only one sample is present in the weblink table (i.e., where the columns "Average", "Minimum" and "Maximum" coincide), were omitted. The value for the last considered stage, i.e. stage 25, is not present/taken in/from the table, but represents an interpolated value for stage 16 + 8 hours. Values for the physical time are obtained by linear interpolating the values from the columns A and B (**Table S8**).

Link to data

The parameters of the ODE model were fitted by two different types of data: time-resolved RNA-seq TPM data and cell counts from immunofluorescence (IF) images at a single later time of development (st. 32). The RNA-seq data observables (cf. **Table S9**) are linked to the dynamical ODE states via scaling parameters / proportionality factors (cf. **Table S6 and S7**), and in some cases offsets (*mcidas, foxa1*). The magnitude of the measurement errors for each observable were fitted by assuming constant error models on the logarithmic scale. The cell count data was fitted with one mean scaling parameter for all experiments which was varied by experiment-specific fold-changes. These fold-changes were penalized with priors such that they do not deviate too much from the mean scaling parameter. The cell count error model comprises only one standard deviation on the absolute cell scale, which prevents the model from using a large standard deviation for specific experiments that cannot be easily explained by the model and instead forces the model to find an explanation if possible.

Hes Gain/Loss of Function (in model 2.1 only)

The developed model was challenged by including cell count data for hes4, hes5 and hes7 loss/gain of function experiments. Please note: For hes gain of function, only low level hes overexpression data was used, as high level expression leads to general concentration-dependent inhibition of cell types. In order to validate if all the important model mechanisms are included, the previously developed model was extended with only the canonical changes to allow for description of loss/gain of function experiments. These changes include a fold change parameter for the initial hes RNA counts in the gain of function conditions and a fold change in the translation rate of hes RNA in the loss of function conditions. Additionally, the model allowed for different initial ISC/MCC counts in the gain/loss of function conditions, since the mathematical model effectively only starts at developmental stage 9, although deviating from the wildtype initial cell counts was penalized by use of a prior to prevent the model to abuse this freedom in parametrization.

All model parameters have been re-optimized to fit wildtype RNA and cell data as well as hes loss/gain of function cell counts simultaneously. Overall, the model structure was flexible enough to reasonably fit most data sets. This suggests that the assumed model appropriately captures the intermediate stage mechanisms of the process. The fit quality of the hes7 perturbations is lacking compared to hes4/hes5, which suggests that the model is not capturing the early stage mechanisms adequately enough to explain such perturbations.

Software used

All modelling analyses were conducted using the Data2Dynamics modelling toolbox (https://github.com/Data2Dynamics/d2d)[62] which runs under Matlab, The MathWorks, Natick, MA.
