## Supplementary material for "Temporal Notch signaling and Hes-mediated competitive de-repression regulate mucociliary cell fates in *Xenopus*": Mathematical Model 1 and 2

### Data2Dynamics Software – Modeling Report

Fabian Lorenz

22-Dec-2022 19:00:38

**Website:** `https://github.com/Data2Dynamics/d2d`

**Key reference:**

- Data2Dynamics: a modeling environment tailored to parameter estimation in dynamical systems. A. Raue, B. Steiert, M. Schelker, C. Kreutz, T. Maiwald, H. Hass, J. Vanlier, C. Tönsing, L. Adlung, R. Engesser, W. Mader, T. Heinemann, J. Hasenauer, M. Schilling, T. Höfer, E. Klipp, F. Theis, U. Klingmüller, B. Schoeberl and J. Timmer. *Bioinformatics*, **31**(21), 3558-3560, 2015.
- Lessons learned from quantitative dynamical modeling in systems biology. A. Raue, M. Schilling, J. Bachmann, A. Matteson, M. Schelker, D. Kaschek, S. Hug, C. Kreutz, BD. Harms, F. Theis, U. Klingmüller, and J. Timmer. *PLOS ONE*, **8**(9), e74335, 2013.

**2 Estimated model parameters** **21**

**3 Uncertainty analysis of model parameters by the profile likelihood** **24**

**4 Confidence intervals for the model parameters** **26**

### 1 Model including spdef

#### 1.1 Comments

Model for the cell composition in mucociliary tissue.

#### 1.2 Dynamic variables

The model contains 27 dynamic variables. The dynamics of those variables evolve according to a system of ordinary differential equations (ODE) as will be defined in the following. The following list indicates the unique variable names and their initial conditions, i.e. their values at time point zero. In order to account for the delay between the transcription of the $\mathrm{hes}$ transcription factors and their transcriptional effects, the so-called linear chain trick has been applied, i.e. a sequence of five states $\mathrm{hes} \rightarrow \mathrm{hes\_d1} \rightarrow \mathrm{hes\_d2} \rightarrow \mathrm{hes\_d3} \rightarrow \mathrm{hes\_d}$ was introduced. Within this chain, production of $\mathrm{hes}$ denotes transcription and the transcriptional effect is mediated by the delayed increase of state $\mathrm{hes\_d}$.

- **Dynamic variable 1:** mpp

$$[\mathrm{mpp}](\mathrm{t}=0) = \mathrm{init\_mpp} \tag{1}$$

- **Dynamic variable 2:** ep

$$[\mathrm{ep}](\mathrm{t}=0) = \mathrm{init\_ep} \tag{2}$$

- **Dynamic variable 3:** bc

$$[\mathrm{bc}](\mathrm{t}=0) = \mathrm{init\_bc} \tag{3}$$

- **Dynamic variable 4:** isc

$$[\mathrm{isc}](\mathrm{t}=0) = \mathrm{init\_isc} \tag{4}$$

- **Dynamic variable 5:** mcc

$$[\mathrm{mcc}](\mathrm{t}=0) = \mathrm{init\_mcc} \tag{5}$$

- **Dynamic variable 6:** mcc_late

$$[\mathrm{mcc\_late}](\mathrm{t}=0) = \mathrm{init\_mcc\_late} \tag{6}$$

- **Dynamic variable 7:** ssc

$$[\mathrm{ssc}](\mathrm{t}=0) = \mathrm{init\_ssc} \tag{7}$$

- **Dynamic variable 8:** lig

$$[\mathrm{lig}](\mathrm{t}=0) = \mathrm{init\_lig} \tag{8}$$

- **Dynamic variable 9:** Lig

$$[\mathrm{Lig}](\mathrm{t}=0) = \mathrm{init\_Lig} \tag{9}$$

- **Dynamic variable 10:** hes4

$$[\mathrm{hes4}](\mathrm{t}=0) = \mathrm{init\_hes4} \tag{10}$$

- **Dynamic variable 11:** hes5

$$[\text{hes5}](t=0) = \text{init\_hes5} \tag{11}$$

- **Dynamic variable 12:** hes7

$$[\text{hes7}](t=0) = \text{init\_hes7} \tag{12}$$

- **Dynamic variable 13:** hes4_d1

$$[\text{hes4\_d1}](t=0) = \text{init\_hes4\_d1} \tag{13}$$

- **Dynamic variable 14:** hes5_d1

$$[\text{hes5\_d1}](t=0) = \text{init\_hes5\_d1} \tag{14}$$

- **Dynamic variable 15:** hes7_d1

$$[\text{hes7\_d1}](t=0) = \text{init\_hes7\_d1} \tag{15}$$

- **Dynamic variable 16:** hes4_d2

$$[\text{hes4\_d2}](t=0) = \text{init\_hes4\_d2} \tag{16}$$

- **Dynamic variable 17:** hes5_d2

$$[\text{hes5\_d2}](t=0) = \text{init\_hes5\_d2} \tag{17}$$

- **Dynamic variable 18:** hes7_d2

$$[\text{hes7\_d2}](t=0) = \text{init\_hes7\_d2} \tag{18}$$

- **Dynamic variable 19:** hes4_d3

$$[\text{hes4\_d3}](t=0) = \text{init\_hes4\_d3} \tag{19}$$

- **Dynamic variable 20:** hes5_d3

$$[\text{hes5\_d3}](t=0) = \text{init\_hes5\_d3} \tag{20}$$

- **Dynamic variable 21:** hes7_d3

$$[\text{hes7\_d3}](t=0) = \text{init\_hes7\_d3} \tag{21}$$

- **Dynamic variable 22:** hes4_d

$$[\text{hes4\_d}](t=0) = \text{init\_hes4\_d} \tag{22}$$

- **Dynamic variable 23:** hes5_d

$$[\text{hes5\_d}](t=0) = \text{init\_hes5\_d} \tag{23}$$

- **Dynamic variable 24:** hes7_d

$$[\text{hes7\_d}](t=0) = \text{init\_hes7\_d} \tag{24}$$

- **Dynamic variable 25:** tp63

$$[\text{tp63}](t=0) = \text{init\_tp63} \tag{25}$$

- **Dynamic variable 26:** spdef

$$[\text{spdef}](t=0) = \text{init\_spdef} \tag{26}$$

- **Dynamic variable 27:** Spdef

$$[\text{Spdef}](t=0) = \text{init\_Spdef} \tag{27}$$

#### 1.3 Reactions

Altogether, the model contains 38 reactions. Reactions define interactions between dynamics variables and were translated into the ODE systems by the rate-equation approach. The following list indicates the reactions and their corresponding reaction rate equations. In the reaction rate equations, dynamic and input variables are indicated by square brackets. The remaining variables are model parameters that were estimated from the data and remain constant over time.

- **Reaction 1:**

$$\mathrm{ep} \longrightarrow \mathrm{mpp}$$

$$v_1 = [\mathrm{ep}] \cdot \mathrm{p\_mpp} \tag{28}$$

- **Reaction 2:**

$$\varnothing \longrightarrow \mathrm{mpp}$$

$$v_2 = \frac{\mathrm{k\_cell\_i}^{\mathrm{h\_cell\_i}}}{\mathrm{k\_cell\_i}^{\mathrm{h\_cell\_i}} + [\mathrm{tp63}]^{\mathrm{h\_cell\_i}}} \tag{29}$$

- **Reaction 3:**

$$\varnothing \longrightarrow \mathrm{isc}$$

$$v_3 = \frac{[\mathrm{hes7\_d}]^{\mathrm{h\_isc\_a}} \cdot \mathrm{k\_cell\_i}^{\mathrm{h\_cell\_i}} \cdot \mathrm{k\_isc\_i\_hes4}^{\mathrm{h\_isc\_i\_hes4}} \cdot \mathrm{k\_isc\_i\_hes5}^{\mathrm{h\_isc\_i\_hes5}} \cdot [\mathrm{mpp}] \cdot \mathrm{v\_max\_isc}}{\left([\mathrm{hes7\_d}]^{\mathrm{h\_isc\_a}} + \mathrm{k\_isc\_a}^{\mathrm{h\_isc\_a}}\right) \cdot \left([\mathrm{hes4\_d}]^{\mathrm{h\_isc\_i\_hes4}} + \mathrm{k\_isc\_i\_hes4}^{\mathrm{h\_isc\_i\_hes4}}\right) \cdot \left(\mathrm{k\_cell\_i}^{\mathrm{h\_cell\_i}} + [\mathrm{tp63}]^{\mathrm{h\_cell\_i}}\right) \cdot \left([\mathrm{hes5\_d}]^{\mathrm{h\_isc\_i\_hes5}} + \mathrm{k\_isc\_i\_hes5}^{\mathrm{h\_isc\_i\_hes5}}\right)} \tag{30}$$

- **Reaction 4:**

$$\varnothing \longrightarrow \mathrm{mcc}$$

$$v_4 = \frac{[\mathrm{hes4\_d}]^{\mathrm{h\_mcc\_a}} \cdot \mathrm{k\_cell\_i}^{\mathrm{h\_cell\_i}} \cdot \mathrm{k\_mcc\_i\_hes5}^{\mathrm{h\_mcc\_i\_hes5}} \cdot [\mathrm{mpp}] \cdot \mathrm{v\_max\_mcc}}{\left([\mathrm{hes4\_d}]^{\mathrm{h\_mcc\_a}} + \mathrm{k\_mcc\_a}^{\mathrm{h\_mcc\_a}}\right) \cdot \left(\mathrm{k\_cell\_i}^{\mathrm{h\_cell\_i}} + [\mathrm{tp63}]^{\mathrm{h\_cell\_i}}\right) \cdot \left([\mathrm{hes5\_d}]^{\mathrm{h\_mcc\_i\_hes5}} + \mathrm{k\_mcc\_i\_hes5}^{\mathrm{h\_mcc\_i\_hes5}}\right)} \tag{31}$$

- **Reaction 5:**

$$\varnothing \longrightarrow \mathrm{ssc}$$

$$v_5 = \frac{[\mathrm{Spdef}]^{\mathrm{h\_ssc\_a\_spdef}} \cdot [\mathrm{hes5\_d}]^{\mathrm{h\_ssc\_a}} \cdot \mathrm{k\_cell\_i}^{\mathrm{h\_cell\_i}} \cdot \mathrm{k\_ssc\_i\_hes4}^{\mathrm{h\_ssc\_i\_hes4}} \cdot \mathrm{k\_ssc\_i\_spdef}^{\mathrm{h\_ssc\_i\_spdef}} \cdot [\mathrm{mpp}] \cdot \mathrm{v\_max\_ssc}}{\left([\mathrm{Spdef}]^{\mathrm{h\_ssc\_a\_spdef}} + \mathrm{k\_ssc\_a\_spdef}^{\mathrm{h\_ssc\_a\_spdef}}\right) \cdot \left([\mathrm{hes5\_d}]^{\mathrm{h\_ssc\_a}} + \mathrm{k\_ssc\_a}^{\mathrm{h\_ssc\_a}}\right) \cdot \left([\mathrm{Spdef}]^{\mathrm{h\_ssc\_i\_spdef}} + \mathrm{k\_ssc\_i\_spdef}^{\mathrm{h\_ssc\_i\_spdef}}\right) \cdot \left(\mathrm{k\_cell\_i}^{\mathrm{h\_cell\_i}} + [\mathrm{tp63}]^{\mathrm{h\_cell\_i}}\right) \cdot \left([\mathrm{hes4\_d}]^{\mathrm{h\_ssc\_i\_hes4}} + \mathrm{k\_ssc\_i\_hes4}^{\mathrm{h\_ssc\_i\_hes4}}\right)} \tag{32}$$

- **Reaction 6:**

$$\mathrm{mpp} \longrightarrow \mathrm{bc}$$

$$v_6 = [\mathrm{mpp}] \cdot \mathrm{v\_max\_bc} \cdot \left( \frac{[\mathrm{Lig}]^{\mathrm{h\_tp63\_a}}}{[\mathrm{Lig}]^{\mathrm{h\_tp63\_a}} + \mathrm{k\_tp63\_a}^{\mathrm{h\_tp63\_a}}} + \frac{[\mathrm{Spdef}]^{\mathrm{h\_tp63\_a\_spdef}}}{[\mathrm{Spdef}]^{\mathrm{h\_tp63\_a\_spdef}} + \mathrm{k\_tp63\_a\_spdef}^{\mathrm{h\_tp63\_a\_spdef}}} \right) \tag{33}$$

- **Reaction 7:**

$$\mathrm{mcc} \longrightarrow \mathrm{mcc\_late}$$

$$v_7 = [\mathrm{mcc}] \cdot \mathrm{p\_mcc\_late} \tag{34}$$

- **Reaction 8:**

$$\mathrm{lig} \longrightarrow \varnothing$$

$$v_8 = \mathrm{d\_lig} \cdot [\mathrm{lig}] \tag{35}$$

- **Reaction 9:**

$$\varnothing \longrightarrow \mathrm{lig}$$

$$v_9 = \frac{[\mathrm{bc}] \cdot \mathrm{p\_lig\_bc} + [\mathrm{ep}] \cdot \mathrm{p\_lig\_bsc} + [\mathrm{isc}] \cdot \mathrm{p\_lig\_isc} + [\mathrm{mcc}] \cdot \mathrm{p\_lig\_mcc} + [\mathrm{mcc\_late}] \cdot \mathrm{p\_lig\_mcc} + [\mathrm{mpp}] \cdot \mathrm{p\_lig\_mpp} + \mathrm{p\_lig\_ssc} \cdot [\mathrm{ssc}]}{[\mathrm{bc}] + [\mathrm{ep}] + [\mathrm{isc}] + [\mathrm{mcc}] + [\mathrm{mcc\_late}] + [\mathrm{mpp}] + [\mathrm{ssc}]} \tag{36}$$

- **Reaction 10:**

$$\varnothing \longrightarrow \mathrm{Lig}$$

$$v_{10} = [\mathrm{lig}] \cdot \mathrm{p\_Lig} \tag{37}$$

- **Reaction 11:**

$$\text{Lig} \longrightarrow \varnothing$$

$$v_{11} = [\text{Lig}] \cdot \text{d\_Lig} \tag{38}$$

- **Reaction 12:**

$$\text{hes4} \longrightarrow \varnothing$$

$$v_{12} = \text{d\_hes4} \cdot [\text{hes4}] \tag{39}$$

- **Reaction 13:**

$$\text{hes5} \longrightarrow \varnothing$$

$$v_{13} = \text{d\_hes5} \cdot [\text{hes5}] \tag{40}$$

- **Reaction 14:**

$$\text{hes7} \longrightarrow \varnothing$$

$$v_{14} = \text{d\_hes7} \cdot [\text{hes7}] \tag{41}$$

- **Reaction 15:**

$$\varnothing \longrightarrow \text{hes4}$$

$$v_{15} = \frac{[\text{Lig}]^{\text{h\_hes4\_a}} \cdot \text{k\_hes4\_i}^{\text{h\_hes4\_i}} \cdot \text{v\_max\_hes4}}{\left([\text{Lig}]^{\text{h\_hes4\_a}} + \text{k\_hes4\_a}^{\text{h\_hes4\_a}}\right) \cdot \left([\text{Lig}]^{\text{h\_hes4\_i}} + \text{k\_hes4\_i}^{\text{h\_hes4\_i}}\right)} \tag{42}$$

- **Reaction 16:**

$$\emptyset \longrightarrow \text{hes5}$$

$$v_{16} = \frac{[\text{Lig}]^{\text{h\_hes5\_a}} \cdot \text{v\_max\_hes5}}{[\text{Lig}]^{\text{h\_hes5\_a}} + \text{k\_hes5\_a}^{\text{h\_hes5\_a}}} \quad (43)$$

- **Reaction 17:**

$$\emptyset \longrightarrow \text{hes7}$$

$$v_{17} = \frac{\text{k\_hes7\_i}^{\text{h\_hes7\_i}} \cdot \text{k\_hes7\_i\_spdef}^{\text{h\_hes7\_i\_spdef}} \cdot \text{v\_max\_hes7}}{\left([\text{Lig}]^{\text{h\_hes7\_i}} + \text{k\_hes7\_i}^{\text{h\_hes7\_i}}\right) \cdot \left([\text{Spdef}]^{\text{h\_hes7\_i\_spdef}} + \text{k\_hes7\_i\_spdef}^{\text{h\_hes7\_i\_spdef}}\right)} \quad (44)$$

- **Reaction 18:**

$$\emptyset \longrightarrow \text{hes4\_d1}$$

$$v_{18} = \frac{4 \cdot [\text{hes4}]}{\text{hes\_delay}} \quad (45)$$

- **Reaction 19:**

$$\emptyset \longrightarrow \text{hes5\_d1}$$

$$v_{19} = \frac{4 \cdot [\text{hes5}]}{\text{hes\_delay}} \quad (46)$$

- **Reaction 20:**

$$\emptyset \longrightarrow \text{hes7\_d1}$$

$$v_{20} = \frac{4 \cdot [\text{hes7}]}{\text{hes\_delay}} \quad (47)$$

- **Reaction 21:**

$$\text{hes4\_d1} \longrightarrow \text{hes4\_d2}$$

$$v_{21} = \frac{4 \cdot [\text{hes4\_d1}]}{\text{hes\_delay}} \tag{48}$$

- **Reaction 22:**

$$\text{hes5\_d1} \longrightarrow \text{hes5\_d2}$$

$$v_{22} = \frac{4 \cdot [\text{hes5\_d1}]}{\text{hes\_delay}} \tag{49}$$

- **Reaction 23:**

$$\text{hes7\_d1} \longrightarrow \text{hes7\_d2}$$

$$v_{23} = \frac{4 \cdot [\text{hes7\_d1}]}{\text{hes\_delay}} \tag{50}$$

- **Reaction 24:**

$$\text{hes4\_d2} \longrightarrow \text{hes4\_d3}$$

$$v_{24} = \frac{4 \cdot [\text{hes4\_d2}]}{\text{hes\_delay}} \tag{51}$$

- **Reaction 25:**

$$\text{hes5\_d2} \longrightarrow \text{hes5\_d3}$$

$$v_{25} = \frac{4 \cdot [\text{hes5\_d2}]}{\text{hes\_delay}} \tag{52}$$

- **Reaction 26:**

$$\text{hes7\_d2} \longrightarrow \text{hes7\_d3}$$

$$v_{26} = \frac{4 \cdot [\text{hes7\_d2}]}{\text{hes\_delay}} \tag{53}$$

- **Reaction 27:**

$$\text{hes4\_d3} \longrightarrow \text{hes4\_d}$$

$$v_{27} = \frac{4 \cdot [\text{hes4\_d3}]}{\text{hes\_delay}} \tag{54}$$

- **Reaction 28:**

$$\text{hes5\_d3} \longrightarrow \text{hes5\_d}$$

$$v_{28} = \frac{4 \cdot [\text{hes5\_d3}]}{\text{hes\_delay}} \tag{55}$$

- **Reaction 29:**

$$\text{hes7\_d3} \longrightarrow \text{hes7\_d}$$

$$v_{29} = \frac{4 \cdot [\text{hes7\_d3}]}{\text{hes\_delay}} \tag{56}$$

- **Reaction 30:**

$$\text{hes4\_d} \longrightarrow \varnothing$$

$$v_{30} = \text{d\_hes4\_d} \cdot [\text{hes4\_d}] \tag{57}$$

- **Reaction 31:**

$$\text{hes5\_d} \longrightarrow \varnothing$$

$$v_{31} = \text{d\_hes5\_d} \cdot [\text{hes5\_d}] \quad (58)$$

- **Reaction 32:**

$$\text{hes7\_d} \longrightarrow \varnothing$$

$$v_{32} = \text{d\_hes7\_d} \cdot [\text{hes7\_d}] \quad (59)$$

- **Reaction 33:**

$$\text{tp63} \longrightarrow \varnothing$$

$$v_{33} = \text{d\_tp63} \cdot [\text{tp63}] \quad (60)$$

- **Reaction 34:**

$$\varnothing \longrightarrow \text{tp63}$$

$$v_{34} = \text{v\_max\_tp63} \cdot \left( \frac{[\text{Lig}]^{\text{h\_tp63\_a}}}{[\text{Lig}]^{\text{h\_tp63\_a}} + \text{k\_tp63\_a}^{\text{h\_tp63\_a}}} + \frac{[\text{Spdef}]^{\text{h\_tp63\_a\_spdef}}}{[\text{Spdef}]^{\text{h\_tp63\_a\_spdef}} + \text{k\_tp63\_a\_spdef}^{\text{h\_tp63\_a\_spdef}}} \right) \quad (61)$$

- **Reaction 35:**

$$\text{spdef} \longrightarrow \varnothing$$

$$v_{35} = \text{d\_spdef} \cdot [\text{spdef}] \quad (62)$$

- **Reaction 36:**

$$\varnothing \longrightarrow \text{spdef}$$

$$v_{36} = \frac{[\text{Lig}]^{\text{h\_spdef\_a}} \cdot \text{v\_max\_spdef}}{[\text{Lig}]^{\text{h\_spdef\_a}} + \text{k\_spdef\_a}^{\text{h\_spdef\_a}}} \tag{63}$$

- **Reaction 37:**

$$\text{Spdef} \longrightarrow \varnothing$$

$$v_{37} = [\text{Spdef}] \cdot \text{d\_Spdef} \tag{64}$$

- **Reaction 38:**

$$\varnothing \longrightarrow \text{Spdef}$$

$$v_{38} = \text{p\_Spdef} \cdot [\text{spdef}] \tag{65}$$

#### 1.4 ODE system

The specified reation laws and rate equations $v$ determine an ODE system. The time evolution of the dynamical variables is calculated by solving this equation system.

$$\begin{aligned}
\text{d[mpp]/dt} &= +v_1 + v_2 - v_6 \\
\text{d[ep]/dt} &= -v_1 \\
\text{d[bc]/dt} &= +v_6 \\
\text{d[isc]/dt} &= +v_3 \\
\text{d[mcc]/dt} &= +v_4 - v_7 \\
\text{d[mcc_late]/dt} &= +v_7 \\
\text{d[ssc]/dt} &= +v_5 \\
\text{d[lig]/dt} &= -v_8 + v_9 \\
\text{d[Lig]/dt} &= +v_{10} - v_{11} \\
\text{d[hes4]/dt} &= -v_{12} + v_{15} \\
\text{d[hes5]/dt} &= -v_{13} + v_{16} \\
\text{d[hes7]/dt} &= -v_{14} + v_{17} \\
\text{d[hes4_d1]/dt} &= +v_{18} - v_{21} \\
\text{d[hes5_d1]/dt} &= +v_{19} - v_{22}
\end{aligned}$$

$$
\begin{aligned}
\mathrm{d[hes7\_d1]/dt} &= +v_{20} - v_{23} \\
\mathrm{d[hes4\_d2]/dt} &= +v_{21} - v_{24} \\
\mathrm{d[hes5\_d2]/dt} &= +v_{22} - v_{25} \\
\mathrm{d[hes7\_d2]/dt} &= +v_{23} - v_{26} \\
\mathrm{d[hes4\_d3]/dt} &= +v_{24} - v_{27} \\
\mathrm{d[hes5\_d3]/dt} &= +v_{25} - v_{28} \\
\mathrm{d[hes7\_d3]/dt} &= +v_{26} - v_{29} \\
\mathrm{d[hes4\_d]/dt} &= +v_{27} - v_{30} \\
\mathrm{d[hes5\_d]/dt} &= +v_{28} - v_{31} \\
\mathrm{d[hes7\_d]/dt} &= +v_{29} - v_{32} \\
\mathrm{d[tp63]/dt} &= -v_{33} + v_{34} \\
\mathrm{d[spdef]/dt} &= -v_{35} + v_{36} \\
\mathrm{d[Spdef]/dt} &= -v_{37} + v_{38}
\end{aligned}
$$

The ODE system was solved by a parallelized implementation of the CVODES algorithm [1]. It also supplies the parameter sensitivities utilized for parameter estimation.

#### 1.5 Derived variables

The model contains 15 derived variables. Derived variables are calculated after the ODE system was solved. Dynamic and input variables are indicated by square brackets. The remaining variables are model parameters that remain constant over time.

- **Derived variable 1:** bsc

$$\mathrm{[bsc](t)} = \mathrm{[bc]} + \mathrm{[ep]} \tag{66}$$

- **Derived variable 2:** mcc_total

$$\mathrm{[mcc\_total](t)} = \mathrm{[mcc]} + \mathrm{[mcc\_late]} \tag{67}$$

- **Derived variable 3:** n_cells

$$\mathrm{[n\_cells](t)} = \mathrm{[bc]} + \mathrm{[ep]} + \mathrm{[isc]} + \mathrm{[mcc]} + \mathrm{[mcc\_late]} + \mathrm{[mpp]} + \mathrm{[ssc]} \tag{68}$$

- **Derived variable 4:** p_isc

$$\mathrm{[p\_isc](t)} = \frac{\mathrm{[hes7\_d]}^{\mathrm{h\_isc\_a}} \cdot \mathrm{k\_cell\_i}^{\mathrm{h\_cell\_i}} \cdot \mathrm{k\_isc\_i\_hes4}^{\mathrm{h\_isc\_i\_hes4}} \cdot \mathrm{k\_isc\_i\_hes5}^{\mathrm{h\_isc\_i\_hes5}} \cdot \mathrm{v\_max\_isc}}{\left(\mathrm{[hes7\_d]}^{\mathrm{h\_isc\_a}} + \mathrm{k\_isc\_a}^{\mathrm{h\_isc\_a}}\right) \cdot \left(\mathrm{[hes4\_d]}^{\mathrm{h\_isc\_i\_hes4}} + \mathrm{k\_isc\_i\_hes4}^{\mathrm{h\_isc\_i\_hes4}}\right) \cdot \left(\mathrm{k\_cell\_i}^{\mathrm{h\_cell\_i}} + \mathrm{[tp63]}^{\mathrm{h\_cell\_i}}\right) \cdot \left(\mathrm{[hes5\_d]}^{\mathrm{h\_isc\_i\_hes5}} + \mathrm{k\_isc\_i\_hes5}^{\mathrm{h\_isc\_i\_hes5}}\right)} \tag{69}$$

- **Derived variable 5:** p_mcc

$$\mathrm{[p\_mcc](t)} = \frac{\mathrm{[hes4\_d]}^{\mathrm{h\_mcc\_a}} \cdot \mathrm{k\_cell\_i}^{\mathrm{h\_cell\_i}} \cdot \mathrm{k\_mcc\_i\_hes5}^{\mathrm{h\_mcc\_i\_hes5}} \cdot \mathrm{v\_max\_mcc}}{\left(\mathrm{[hes4\_d]}^{\mathrm{h\_mcc\_a}} + \mathrm{k\_mcc\_a}^{\mathrm{h\_mcc\_a}}\right) \cdot \left(\mathrm{k\_cell\_i}^{\mathrm{h\_cell\_i}} + \mathrm{[tp63]}^{\mathrm{h\_cell\_i}}\right) \cdot \left(\mathrm{[hes5\_d]}^{\mathrm{h\_mcc\_i\_hes5}} + \mathrm{k\_mcc\_i\_hes5}^{\mathrm{h\_mcc\_i\_hes5}}\right)} \tag{70}$$

- **Derived variable 6:** p_ssc

$$\mathrm{[p\_ssc](t)} = \frac{\mathrm{[Spdef]}^{\mathrm{h\_ssc\_a\_spdef}} \cdot \mathrm{[hes5\_d]}^{\mathrm{h\_ssc\_a}} \cdot \mathrm{k\_cell\_i}^{\mathrm{h\_cell\_i}} \cdot \mathrm{k\_ssc\_i\_hes4}^{\mathrm{h\_ssc\_i\_hes4}} \cdot \mathrm{k\_ssc\_i\_spdef}^{\mathrm{h\_ssc\_i\_spdef}} \cdot \mathrm{v\_max\_ssc}}{\left(\mathrm{[Spdef]}^{\mathrm{h\_ssc\_a\_spdef}} + \mathrm{k\_ssc\_a\_spdef}^{\mathrm{h\_ssc\_a\_spdef}}\right) \cdot \left(\mathrm{[hes5\_d]}^{\mathrm{h\_ssc\_a}} + \mathrm{k\_ssc\_a}^{\mathrm{h\_ssc\_a}}\right) \cdot \left(\mathrm{[Spdef]}^{\mathrm{h\_ssc\_i\_spdef}} + \mathrm{k\_ssc\_i\_spdef}^{\mathrm{h\_ssc\_i\_spdef}}\right) \cdot \left(\mathrm{k\_cell\_i}^{\mathrm{h\_cell\_i}} + \mathrm{[tp63]}^{\mathrm{h\_cell\_i}}\right) \cdot \left(\mathrm{[hes4\_d]}^{\mathrm{h\_ssc\_i\_hes4}} [illegible]\right.} \tag{71}$$

- **Derived variable 7:** p_bc

$$[\mathrm{p\_bc}](t) = \mathrm{v\_max\_bc} \cdot \left( \frac{[\mathrm{Lig}]^{\mathrm{h\_tp63\_a}}}{[\mathrm{Lig}]^{\mathrm{h\_tp63\_a}} + \mathrm{k\_tp63\_a}^{\mathrm{h\_tp63\_a}}} + \frac{[\mathrm{Spdef}]^{\mathrm{h\_tp63\_a\_spdef}}}{[\mathrm{Spdef}]^{\mathrm{h\_tp63\_a\_spdef}} + \mathrm{k\_tp63\_a\_spdef}^{\mathrm{h\_tp63\_a\_spdef}}} \right) \tag{72}$$

- **Derived variable 8:** p_lig

$$[\mathrm{p\_lig}](t) = \frac{[\mathrm{bc}] \cdot \mathrm{p\_lig\_bc} + [\mathrm{ep}] \cdot \mathrm{p\_lig\_bsc} + [\mathrm{isc}] \cdot \mathrm{p\_lig\_isc} + [\mathrm{mcc}] \cdot \mathrm{p\_lig\_mcc} + [\mathrm{mcc\_late}] \cdot \mathrm{p\_lig\_mcc} + [\mathrm{mpp}] \cdot \mathrm{p\_lig\_mpp} + \mathrm{p\_lig\_ssc} \cdot [\mathrm{ssc}]}{[\mathrm{bc}] + [\mathrm{ep}] + [\mathrm{isc}] + [\mathrm{mcc}] + [\mathrm{mcc\_late}] + [\mathrm{mpp}] + [\mathrm{ssc}]} \tag{73}$$

- **Derived variable 9:** p_hes4

$$[\mathrm{p\_hes4}](t) = \frac{[\mathrm{Lig}]^{\mathrm{h\_hes4\_a}} \cdot \mathrm{k\_hes4\_i}^{\mathrm{h\_hes4\_i}} \cdot \mathrm{v\_max\_hes4}}{\left([\mathrm{Lig}]^{\mathrm{h\_hes4\_a}} + \mathrm{k\_hes4\_a}^{\mathrm{h\_hes4\_a}}\right) \cdot \left([\mathrm{Lig}]^{\mathrm{h\_hes4\_i}} + \mathrm{k\_hes4\_i}^{\mathrm{h\_hes4\_i}}\right)} \tag{74}$$

- **Derived variable 10:** p_hes5

$$[\mathrm{p\_hes5}](t) = \frac{[\mathrm{Lig}]^{\mathrm{h\_hes5\_a}} \cdot \mathrm{v\_max\_hes5}}{[\mathrm{Lig}]^{\mathrm{h\_hes5\_a}} + \mathrm{k\_hes5\_a}^{\mathrm{h\_hes5\_a}}} \tag{75}$$

- **Derived variable 11:** p_hes7

$$[\mathrm{p\_hes7}](t) = \frac{\mathrm{k\_hes7\_i}^{\mathrm{h\_hes7\_i}} \cdot \mathrm{k\_hes7\_i\_spdef}^{\mathrm{h\_hes7\_i\_spdef}} \cdot \mathrm{v\_max\_hes7}}{\left([\mathrm{Lig}]^{\mathrm{h\_hes7\_i}} + \mathrm{k\_hes7\_i}^{\mathrm{h\_hes7\_i}}\right) \cdot \left([\mathrm{Spdef}]^{\mathrm{h\_hes7\_i\_spdef}} + \mathrm{k\_hes7\_i\_spdef}^{\mathrm{h\_hes7\_i\_spdef}}\right)} \tag{76}$$

- **Derived variable 12:** p_spdef

$$[\mathrm{p\_spdef}](t) = \frac{[\mathrm{Lig}]^{\mathrm{h\_spdef\_a}} \cdot \mathrm{v\_max\_spdef}}{[\mathrm{Lig}]^{\mathrm{h\_spdef\_a}} + \mathrm{k\_spdef\_a}^{\mathrm{h\_spdef\_a}}} \tag{77}$$

- **Derived variable 13:** p_tp63

$$[\mathrm{p\_tp63}](t) = \mathrm{v\_max\_tp63} \cdot \left( \frac{[\mathrm{Lig}]^{\mathrm{h\_tp63\_a}}}{[\mathrm{Lig}]^{\mathrm{h\_tp63\_a}} + \mathrm{k\_tp63\_a}^{\mathrm{h\_tp63\_a}}} + \frac{[\mathrm{Spdef}]^{\mathrm{h\_tp63\_a\_spdef}}}{[\mathrm{Spdef}]^{\mathrm{h\_tp63\_a\_spdef}} + \mathrm{k\_tp63\_a\_spdef}^{\mathrm{h\_tp63\_a\_spdef}}} \right) \tag{78}$$

- **Derived variable 14:** tp63_hill

$$[\mathrm{tp63\_hill}](t) = \frac{[\mathrm{Lig}]^{\mathrm{h\_tp63\_a}}}{[\mathrm{Lig}]^{\mathrm{h\_tp63\_a}} + \mathrm{k\_tp63\_a}^{\mathrm{h\_tp63\_a}}} + \frac{[\mathrm{Spdef}]^{\mathrm{h\_tp63\_a\_spdef}}}{[\mathrm{Spdef}]^{\mathrm{h\_tp63\_a\_spdef}} + \mathrm{k\_tp63\_a\_spdef}^{\mathrm{h\_tp63\_a\_spdef}}} \tag{79}$$

#### 1.6 Conditions

Conditions modify the model according to replacement rules. New model parameters can be introduced or relations between existing model parameters can be implemented. The following list are default conditions that can be replace my experiment specific conditions defined seperately for each data set.

In our project, we used replacements primarily to set dynamic variable to steady states. The steady states were derived by setting the right-hand side of the ODEs to zero and solve for the dynamic variables. In addition, some initial conditions which are not assumed in the steady state were replaced by the initial value of the corresponding observation to better control parameter bounds and initialization of parameter optimization.

$$\mathrm{init\_bc} \rightarrow 0$$

$$\mathrm{init\_hes4} \rightarrow \frac{\mathrm{init\_hes4}}{\mathrm{s\_hes4}}$$

$$\text{init\_hes4\_d} \rightarrow \frac{4 \cdot \text{init\_hes4}}{\text{d\_hes4\_d} \cdot \text{hes\_delay} \cdot \text{s\_hes4}}$$

$$\text{init\_hes4\_d1} \rightarrow \frac{\text{init\_hes4}}{\text{s\_hes4}}$$

$$\text{init\_hes4\_d2} \rightarrow \frac{\text{init\_hes4}}{\text{s\_hes4}}$$

$$\text{init\_hes4\_d3} \rightarrow \frac{\text{init\_hes4}}{\text{s\_hes4}}$$

$$\text{init\_hes5} \rightarrow \frac{\text{init\_hes5}}{\text{s\_hes5}}$$

$$\text{init\_hes5\_d} \rightarrow \frac{4 \cdot \text{init\_hes5}}{\text{d\_hes5\_d} \cdot \text{hes\_delay} \cdot \text{s\_hes5}}$$

$$\text{init\_hes5\_d1} \rightarrow \frac{\text{init\_hes5}}{\text{s\_hes5}}$$

$$\text{init\_hes5\_d2} \rightarrow \frac{\text{init\_hes5}}{\text{s\_hes5}}$$

$$\text{init\_hes5\_d3} \rightarrow \frac{\text{init\_hes5}}{\text{s\_hes5}}$$

$$\text{init\_hes7} \rightarrow \frac{\text{init\_hes7}}{\text{s\_hes7}}$$

$$\text{init\_hes7\_d} \rightarrow \frac{4 \cdot \text{init\_hes7}}{\text{d\_hes7\_d} \cdot \text{hes\_delay} \cdot \text{s\_hes7}}$$

$$\text{init\_hes7\_d1} \rightarrow \frac{\text{init\_hes7}}{\text{s\_hes7}}$$

$$\text{init\_hes7\_d2} \rightarrow \frac{\text{init\_hes7}}{\text{s\_hes7}}$$

$$\text{init\_hes7\_d3} \rightarrow \frac{\text{init\_hes7}}{\text{s\_hes7}}$$

$$\text{init\_lig} \rightarrow \frac{\text{p\_lig\_bsc} \cdot (\text{init\_ep} + \text{init\_mpp} \cdot \text{p\_lig\_mpp\_factor} + \text{init\_isc} \cdot \text{p\_lig\_isc\_factor} \cdot \text{p\_lig\_mpp\_factor})}{\text{d\_lig} \cdot (\text{init\_ep} + \text{init\_isc} + \text{init\_mpp})}$$

$$\text{init\_mcc} \rightarrow 0$$

$$\text{init\_mcc\_late} \rightarrow 0$$

$$\text{init\_Spdef} \rightarrow \frac{\text{init\_spdef} \cdot \text{p\_Spdef}}{\text{d\_Spdef} \cdot \text{s\_spdef}}$$

$$\text{init\_spdef} \rightarrow \frac{\text{init\_spdef}}{\text{s\_spdef}}$$

$$\text{init\_ssc} \rightarrow 0$$

$$\text{init\_Lig} \rightarrow \frac{\text{p\_Lig} \cdot \text{p\_lig\_bsc} \cdot (\text{init\_ep} + \text{init\_mpp} \cdot \text{p\_lig\_mpp\_factor} + \text{init\_isc} \cdot \text{p\_lig\_isc\_factor} \cdot \text{p\_lig\_mpp\_factor})}{\text{d\_Lig} \cdot \text{d\_lig} \cdot (\text{init\_ep} + \text{init\_isc} + \text{init\_mpp})}$$

$$\text{init\_tp63} \rightarrow \frac{\text{init\_tp63}}{\text{s\_tp63}}$$

$$\text{k\_hes4\_i} \rightarrow \text{k\_hes4\_a\_delta} + \text{k\_hes4\_i\_delta} + \frac{\text{p\_Lig} \cdot \left( \frac{\text{init\_ep} \cdot \text{p\_lig\_bsc}}{\text{init\_ep} + \text{init\_isc} + \text{init\_mpp}} + \frac{\text{init\_mpp} \cdot \text{p\_lig\_bsc} \cdot \text{p\_lig\_mpp\_factor}}{\text{init\_ep} + \text{init\_isc} + \text{init\_mpp}} + \frac{\text{init\_isc} \cdot \text{p\_lig\_bsc} \cdot \text{p\_lig\_isc\_factor} \cdot \text{p\_lig\_mpp\_factor}}{\text{init\_ep} + \text{init\_isc} + \text{init\_mpp}} \right)}{\text{d\_Lig} \cdot \text{d\_lig}}$$

$$\text{k\_hes4\_a} \rightarrow \frac{\text{init\_ep} \cdot \text{p\_Lig} \cdot \text{p\_lig\_bsc} + \text{d\_Lig} \cdot \text{d\_lig} \cdot \text{init\_ep} \cdot \text{k\_hes4\_a\_delta} + \text{d\_Lig} \cdot \text{d\_lig} \cdot \text{init\_isc} \cdot \text{k\_hes4\_a\_delta} + \text{d\_Lig} \cdot \text{d\_lig} \cdot \text{init\_mpp} \cdot \text{k\_hes4\_a\_delta} + \text{init\_mpp} \cdot \text{p\_Lig} \cdot \text{p\_lig\_bsc} \cdot \text{p\_lig\_mpp\_factor} + \text{init\_is}}{\text{d\_Lig} \cdot \text{d\_lig} \cdot (\text{init\_ep} + \text{init\_isc} + \text{init\_mpp})}$$ [illegible]

$$\text{k\_hes5\_a} \rightarrow \frac{\text{init\_ep} \cdot \text{p\_Lig} \cdot \text{p\_lig\_bsc} + \text{d\_Lig} \cdot \text{d\_lig} \cdot \text{init\_ep} \cdot \text{k\_hes5\_a\_delta} + \text{d\_Lig} \cdot \text{d\_lig} \cdot \text{init\_isc} \cdot \text{k\_hes5\_a\_delta} + \text{d\_Lig} \cdot \text{d\_lig} \cdot \text{init\_mpp} \cdot \text{k\_hes5\_a\_delta} + \text{init\_mpp} \cdot \text{p\_Lig} \cdot \text{p\_lig\_bsc} \cdot \text{p\_lig\_mpp\_factor} + \text{init\_is}}{\text{d\_Lig} \cdot \text{d\_lig} \cdot (\text{init\_ep} + \text{init\_isc} + \text{init\_mpp})}$$ [illegible]

$$\text{k\_hes7\_i} \rightarrow \frac{\text{init\_ep} \cdot \text{p\_Lig} \cdot \text{p\_lig\_bsc} + \text{d\_Lig} \cdot \text{d\_lig} \cdot \text{init\_ep} \cdot \text{k\_hes7\_i\_delta} + \text{d\_Lig} \cdot \text{d\_lig} \cdot \text{init\_isc} \cdot \text{k\_hes7\_i\_delta} + \text{d\_Lig} \cdot \text{d\_lig} \cdot \text{init\_mpp} \cdot \text{k\_hes7\_i\_delta} + \text{init\_mpp} \cdot \text{p\_Lig} \cdot \text{p\_lig\_bsc} \cdot \text{p\_lig\_mpp\_factor} + \text{init\_isc}}{\text{d\_Lig} \cdot \text{d\_lig} \cdot (\text{init\_ep} + \text{init\_isc} + \text{init\_mpp})}$$ [illegible]

$$\mathrm{k\_hes7\_i\_spdef} \rightarrow \mathrm{k\_hes7\_i\_spdef\_delta} + \frac{\mathrm{init\_spdef} \cdot \mathrm{p\_Spdef}}{\mathrm{d\_Spdef} \cdot \mathrm{s\_spdef}}$$

$$\mathrm{k\_isc\_i\_hes4} \rightarrow \mathrm{k\_isc\_i\_hes4\_delta} + \frac{4 \cdot \mathrm{init\_hes4}}{\mathrm{d\_hes4\_d} \cdot \mathrm{hes\_delay} \cdot \mathrm{s\_hes4}}$$

$$\mathrm{k\_isc\_i\_hes5} \rightarrow \mathrm{k\_isc\_i\_hes5\_delta} + \frac{4 \cdot \mathrm{init\_hes5}}{\mathrm{d\_hes5\_d} \cdot \mathrm{hes\_delay} \cdot \mathrm{s\_hes5}}$$

$$\mathrm{k\_mcc\_i\_hes5} \rightarrow \mathrm{k\_mcc\_i\_hes5\_delta} + \frac{4 \cdot \mathrm{init\_hes5}}{\mathrm{d\_hes5\_d} \cdot \mathrm{hes\_delay} \cdot \mathrm{s\_hes5}}$$

$$\mathrm{k\_spdef\_a} \rightarrow \frac{\mathrm{init\_ep} \cdot \mathrm{p\_Lig} \cdot \mathrm{p\_lig\_bsc} + \mathrm{d\_Lig} \cdot \mathrm{d\_lig} \cdot \mathrm{init\_ep} \cdot \mathrm{k\_spdef\_a\_delta} + \mathrm{d\_Lig} \cdot \mathrm{d\_lig} \cdot \mathrm{init\_isc} \cdot \mathrm{k\_spdef\_a\_delta} + \mathrm{d\_Lig} \cdot \mathrm{d\_lig} \cdot \mathrm{init\_mpp} \cdot \mathrm{k\_spdef\_a\_delta} + \mathrm{init\_mpp} \cdot \mathrm{p\_Lig} \cdot \mathrm{p\_lig\_bsc} \cdot \mathrm{p\_lig\_mpp\_factor} + \mathrm{in}\ldots}{\mathrm{d\_Lig} \cdot \mathrm{d\_lig} \cdot (\mathrm{init\_ep} + \mathrm{init\_isc} + \mathrm{init\_mpp})}$$ [illegible]

$$\mathrm{k\_ssc\_i\_hes4} \rightarrow \mathrm{k\_ssc\_i\_hes4\_delta} + \frac{4 \cdot \mathrm{init\_hes4}}{\mathrm{d\_hes4\_d} \cdot \mathrm{hes\_delay} \cdot \mathrm{s\_hes4}}$$

$$\mathrm{k\_ssc\_i\_spdef} \rightarrow \mathrm{k\_ssc\_i\_spdef\_delta} + \frac{\mathrm{init\_spdef} \cdot \mathrm{p\_Spdef}}{\mathrm{d\_Spdef} \cdot \mathrm{s\_spdef}}$$

$$\mathrm{k\_ssc\_a\_spdef} \rightarrow \mathrm{k\_ssc\_a\_spdef\_delta} + \frac{\mathrm{init\_spdef} \cdot \mathrm{p\_Spdef}}{\mathrm{d\_Spdef} \cdot \mathrm{s\_spdef}}$$

$$\mathrm{k\_tp63\_a} \rightarrow \frac{\mathrm{init\_ep} \cdot \mathrm{p\_Lig} \cdot \mathrm{p\_lig\_bsc} + \mathrm{d\_Lig} \cdot \mathrm{d\_lig} \cdot \mathrm{init\_ep} \cdot \mathrm{k\_tp63\_a\_delta} + \mathrm{d\_Lig} \cdot \mathrm{d\_lig} \cdot \mathrm{init\_isc} \cdot \mathrm{k\_tp63\_a\_delta} + \mathrm{d\_Lig} \cdot \mathrm{d\_lig} \cdot \mathrm{init\_mpp} \cdot \mathrm{k\_tp63\_a\_delta} + \mathrm{init\_mpp} \cdot \mathrm{p\_Lig} \cdot \mathrm{p\_lig\_bsc} \cdot \mathrm{p\_lig\_mpp\_factor} + \mathrm{init\_i}\ldots}{\mathrm{d\_Lig} \cdot \mathrm{d\_lig} \cdot (\mathrm{init\_ep} + \mathrm{init\_isc} + \mathrm{init\_mpp})}$$ [illegible]

$$\mathrm{k\_tp63\_a\_spdef} \rightarrow \mathrm{k\_tp63\_a\_spdef\_delta} + \frac{\mathrm{init\_spdef} \cdot \mathrm{p\_Spdef}}{\mathrm{d\_Spdef} \cdot \mathrm{s\_spdef}}$$

$$\mathrm{p\_lig\_bc} \rightarrow \mathrm{p\_lig\_bc\_factor} \cdot \mathrm{p\_lig\_bsc} \cdot \mathrm{p\_lig\_mpp\_factor}$$

$$\mathrm{p\_lig\_isc} \rightarrow \mathrm{p\_lig\_bsc} \cdot \mathrm{p\_lig\_isc\_factor} \cdot \mathrm{p\_lig\_mpp\_factor}$$

$$\mathrm{p\_lig\_mcc} \rightarrow \mathrm{p\_lig\_bsc} \cdot \mathrm{p\_lig\_mcc\_factor} \cdot \mathrm{p\_lig\_mpp\_factor}$$

$$\mathrm{p\_lig\_mpp} \rightarrow \mathrm{p\_lig\_bsc} \cdot \mathrm{p\_lig\_mpp\_factor}$$

$$\mathrm{p\_lig\_ssc} \rightarrow \mathrm{p\_lig\_bsc} \cdot \mathrm{p\_lig\_mpp\_factor} \cdot \mathrm{p\_lig\_ssc\_factor}$$

$$\mathrm{v\_max\_isc} \rightarrow \mathrm{v\_max}$$

$$\mathrm{v\_max\_mcc} \rightarrow \mathrm{v\_max}$$

$$\mathrm{v\_max\_ssc} \rightarrow \mathrm{v\_max}$$

$$\mathrm{v\_max\_hes4} \rightarrow \frac{\mathrm{d\_hes4} \cdot \mathrm{init\_hes4} \cdot \left(\left(\frac{\mathrm{init\_ep} \cdot \mathrm{p\_Lig} \cdot \mathrm{p\_lig\_bsc}+\mathrm{d\_Lig} \cdot \mathrm{d\_lig} \cdot \mathrm{init\_ep} \cdot \mathrm{k\_hes4\_a\_delta}+\mathrm{d\_Lig} \cdot \mathrm{d\_lig} \cdot \mathrm{init\_ep} \cdot \mathrm{k\_hes4\_i\_delta}+\mathrm{d\_Lig} \cdot \mathrm{d\_lig} \cdot \mathrm{init\_isc} \cdot \mathrm{k\_hes4\_a\_delta}+\mathrm{d\_Lig} \cdot \mathrm{d\_lig} \cdot \mathrm{init\_isc} \cdot \mathrm{k\_hes4\_i\_delta}+\mathrm{d\_Lig} \cdot \mathrm{d\_lig} \cdot \mathrm{init\_mpp} \cdot \mathrm{k\_hes4\_a\_delta}+\mathrm{d\_Lig} \cdot \mathrm{d\_lig} \cdot \mathrm{init\_mpp} \cdot \mathrm{k\_hes4\_i\_delta}+\mathrm{init\_mpp} \cdot \mathrm{p\_Lig} \cdot \mathrm{p\_lig\_bsc} \cdot \mathrm{p\_lig\_mpp\_factor}+\mathrm{init\_isc} \cdot \mathrm{p\_Lig} \cdot \mathrm{p\_lig\_bsc} \cdot \mathrm{p\_lig\_isc\_factor} \cdot \mathrm{p\_lig\_mpp\_factor}}{\mathrm{d\_Lig} \cdot \mathrm{d\_lig} \cdot (\mathrm{init\_ep}+\mathrm{init\_isc}+\mathrm{init\_mpp})}\right)^{\mathrm{h\_hes4\_i}}+\left(\frac{\mathrm{p\_Lig} \cdot \mathrm{p\_lig\_bsc} \cdot (\mathrm{init\_ep}+\mathrm{init\_mpp} \cdot \mathrm{p\_lig\_mpp\_factor}+\mathrm{init\_isc} \cdot \mathrm{p\_lig\_isc\_factor} \cdot \mathrm{p\_lig\_mpp\_factor})}{\mathrm{d\_Lig} \cdot \mathrm{d\_lig} \cdot (\mathrm{init\_ep}+\mathrm{init\_isc}+\mathrm{init\_mpp})}\right)^{\mathrm{h\_hes4\_i}}\right) \cdot \left(\left(\frac{\mathrm{init\_ep} \cdot \mathrm{p\_Lig} \cdot \mathrm{p\_lig\_bsc}+\mathrm{d\_Lig} \cdot \mathrm{d\_lig} \cdot \mathrm{init\_ep} \cdot \mathrm{k\_hes4\_a\_delta}+\mathrm{d\_Lig} \cdot \mathrm{d\_lig} \cdot \mathrm{init\_isc} \cdot \mathrm{k\_hes4\_a\_delta}+\mathrm{d\_Lig} \cdot \mathrm{d\_lig} \cdot \mathrm{init\_mpp} \cdot \mathrm{k\_hes4\_a\_delta}+\mathrm{init\_mpp} \cdot \mathrm{p\_Lig} \cdot \mathrm{p\_lig\_bsc} \cdot \mathrm{p\_lig\_mpp\_factor}+\mathrm{init\_isc} \cdot \mathrm{p\_Lig} \cdot \mathrm{p\_lig\_bsc} \cdot \mathrm{p\_lig\_isc\_factor} \cdot \mathrm{p\_lig\_mpp\_factor}}{\mathrm{d\_Lig} \cdot \mathrm{d\_lig} \cdot (\mathrm{init\_ep}+\mathrm{init\_isc}+\mathrm{init\_mpp})}\right)^{\mathrm{h\_hes4\_a}}+\left(\frac{\mathrm{p\_Lig} \cdot \mathrm{p\_lig\_bsc} \cdot (\mathrm{init\_ep}+\mathrm{init\_mpp} \cdot \mathrm{p\_lig\_mpp\_factor}+\mathrm{init\_isc} \cdot \mathrm{p\_lig\_isc\_factor} \cdot \mathrm{p\_lig\_mpp\_factor})}{\mathrm{d\_Lig} \cdot \mathrm{d\_lig} \cdot (\mathrm{init\_ep}+\mathrm{init\_isc}+\mathrm{init\_mpp})}\right)^{\mathrm{h\_hes4\_a}}\right)}{\mathrm{s\_hes4} \cdot \left(\frac{\mathrm{init\_ep} \cdot \mathrm{p\_Lig} \cdot \mathrm{p\_lig\_bsc}+\mathrm{d\_Lig} \cdot \mathrm{d\_lig} \cdot \mathrm{init\_ep} \cdot \mathrm{k\_hes4\_a\_delta}+\mathrm{d\_Lig} \cdot \mathrm{d\_lig} \cdot \mathrm{init\_ep} \cdot \mathrm{k\_hes4\_i\_delta}+\mathrm{d\_Lig} \cdot \mathrm{d\_lig} \cdot \mathrm{init\_isc} \cdot \mathrm{k\_hes4\_a\_delta}+\mathrm{d\_Lig} \cdot \mathrm{d\_lig} \cdot \mathrm{init\_isc} \cdot \mathrm{k\_hes4\_i\_delta}+\mathrm{d\_Lig} \cdot \mathrm{d\_lig} \cdot \mathrm{init\_mpp} \cdot \mathrm{k\_hes4\_a\_delta}+\mathrm{d\_Lig} \cdot \mathrm{d\_lig} \cdot \mathrm{init\_mpp} \cdot \mathrm{k\_hes4\_i\_delta}+\mathrm{init\_mpp} \cdot \mathrm{p\_Lig} \cdot \mathrm{p\_lig\_bsc} \cdot \mathrm{p\_lig\_mpp\_factor}+\mathrm{init\_isc} \cdot \mathrm{p\_Lig} \cdot \mathrm{p\_lig\_bsc} \cdot \mathrm{p\_lig\_isc\_factor} \cdot \mathrm{p\_lig\_mpp\_factor}}{\mathrm{d\_Lig} \cdot \mathrm{d\_lig} \cdot (\mathrm{init\_ep}+\mathrm{init\_isc}+\mathrm{init\_mpp})}\right)^{\mathrm{h\_hes4\_i}} \cdot \left(\frac{\mathrm{p\_Lig} \cdot \mathrm{p\_lig\_bsc} \cdot (\mathrm{init\_ep}+\mathrm{init\_mpp} \cdot \mathrm{p\_lig\_mpp\_factor}+\mathrm{init\_isc} \cdot \mathrm{p\_lig\_isc\_factor} \cdot \mathrm{p\_lig\_mpp\_factor})}{\mathrm{d\_Lig} \cdot \mathrm{d\_lig} \cdot (\mathrm{init\_ep}+\mathrm{init\_isc}+\mathrm{init\_mpp})}\right)^{\mathrm{h\_hes4\_a}}}$$

$$\mathrm{v\_max\_hes5} \rightarrow \frac{\mathrm{d\_hes5} \cdot \mathrm{init\_hes5} \cdot \left(\left(\frac{\mathrm{init\_ep} \cdot \mathrm{p\_Lig} \cdot \mathrm{p\_lig\_bsc}+\mathrm{d\_Lig} \cdot \mathrm{d\_lig} \cdot \mathrm{init\_ep} \cdot \mathrm{k\_hes5\_a\_delta}+\mathrm{d\_Lig} \cdot \mathrm{d\_lig} \cdot \mathrm{init\_isc} \cdot \mathrm{k\_hes5\_a\_delta}+\mathrm{d\_Lig} \cdot \mathrm{d\_lig} \cdot \mathrm{init\_mpp} \cdot \mathrm{k\_hes5\_a\_delta}+\mathrm{init\_mpp} \cdot \mathrm{p\_Lig} \cdot \mathrm{p\_lig\_bsc} \cdot \mathrm{p\_lig\_mpp\_factor}+\mathrm{init\_isc} \cdot \mathrm{p\_Lig} \cdot \mathrm{p\_lig\_bsc} \cdot \mathrm{p\_lig\_isc\_factor} \cdot \mathrm{p\_lig\_mpp\_factor}}{\mathrm{d\_Lig} \cdot \mathrm{d\_lig} \cdot (\mathrm{init\_ep}+\mathrm{init\_isc}+\mathrm{init\_mpp})}\right)^{\mathrm{h\_hes5\_a}}+\left(\frac{\mathrm{p\_Lig} \cdot \mathrm{p\_lig\_bsc} \cdot (\mathrm{init\_ep}+\mathrm{init\_mpp} \cdot \mathrm{p\_lig\_mpp\_factor}+\mathrm{init\_isc} \cdot \mathrm{p\_lig\_isc\_factor} \cdot \mathrm{p\_lig\_mpp\_factor})}{\mathrm{d\_Lig} \cdot \mathrm{d\_lig} \cdot (\mathrm{init\_ep}+\mathrm{init\_isc}+\mathrm{init\_mpp})}\right)^{\mathrm{h\_hes5\_a}}\right)}{\mathrm{s\_hes5} \cdot \left(\frac{\mathrm{p\_Lig} \cdot \mathrm{p\_lig\_bsc} \cdot (\mathrm{init\_ep}+\mathrm{init\_mpp} \cdot \mathrm{p\_lig\_mpp\_factor}+\mathrm{init\_isc} \cdot \mathrm{p\_lig\_isc\_factor} \cdot \mathrm{p\_lig\_mpp\_factor})}{\mathrm{d\_Lig} \cdot \mathrm{d\_lig} \cdot (\mathrm{init\_ep}+\mathrm{init\_isc}+\mathrm{init\_mpp})}\right)^{\mathrm{h\_hes5\_a}}}$$

$$\mathrm{v\_max\_hes7} \rightarrow \frac{\mathrm{d\_hes7} \cdot \mathrm{init\_hes7} \cdot \left(\left(\frac{\mathrm{init\_spdef} \cdot \mathrm{p\_Spdef}+\mathrm{d\_Spdef} \cdot \mathrm{k\_hes7\_i\_spdef\_delta} \cdot \mathrm{s\_spdef}}{\mathrm{d\_Spdef} \cdot \mathrm{s\_spdef}}\right)^{\mathrm{h\_hes7\_i\_spdef}}+\left(\frac{\mathrm{init\_spdef} \cdot \mathrm{p\_Spdef}}{\mathrm{d\_Spdef} \cdot \mathrm{s\_spdef}}\right)^{\mathrm{h\_hes7\_i\_spdef}}\right) \cdot \left(\left(\frac{\mathrm{init\_ep} \cdot \mathrm{p\_Lig} \cdot \mathrm{p\_lig\_bsc}+\mathrm{d\_Lig} \cdot \mathrm{d\_lig} \cdot \mathrm{init\_ep} \cdot \mathrm{k\_hes7\_i\_delta}+\mathrm{d\_Lig} \cdot \mathrm{d\_lig} \cdot \mathrm{init\_isc} \cdot \mathrm{k\_hes7\_i\_delta}+\mathrm{d\_Lig} \cdot \mathrm{d\_lig} \cdot \mathrm{init\_mpp} \cdot \mathrm{k\_hes7\_i\_delta}+\mathrm{init\_mpp} \cdot \mathrm{p\_Lig} \cdot \mathrm{p\_lig\_bsc} \cdot \mathrm{p\_lig\_mpp\_factor}+\mathrm{init\_isc} \cdot \mathrm{p\_Lig} \cdot \mathrm{p\_lig\_bsc} \cdot \mathrm{p\_lig\_isc\_factor} \cdot \mathrm{p\_lig\_mpp\_factor}}{\mathrm{d\_Lig} \cdot \mathrm{d\_lig} \cdot (\mathrm{init\_ep}+\mathrm{init\_isc}+\mathrm{init\_mpp})}\right)^{\mathrm{h\_hes7\_i}}+\left(\frac{\mathrm{p\_Lig} \cdot \mathrm{p\_lig\_bsc} \cdot (\mathrm{init\_ep}+\mathrm{init\_mpp} \cdot \mathrm{p\_lig\_mpp\_factor}+\mathrm{init\_isc} \cdot \mathrm{p\_lig\_isc\_factor} \cdot \mathrm{p\_lig\_mpp\_factor})}{\mathrm{d\_Lig} \cdot \mathrm{d\_lig} \cdot (\mathrm{init\_ep}+\mathrm{init\_isc}+\mathrm{init\_mpp})}\right)^{\mathrm{h\_hes7\_i}}\right)}{\mathrm{s\_hes7} \cdot \left(\frac{\mathrm{init\_spdef} \cdot \mathrm{p\_Spdef}+\mathrm{d\_Spdef} \cdot \mathrm{k\_hes7\_i\_spdef\_delta} \cdot \mathrm{s\_spdef}}{\mathrm{d\_Spdef} \cdot \mathrm{s\_spdef}}\right)^{\mathrm{h\_hes7\_i\_spdef}} \cdot \left(\frac{\mathrm{init\_ep} \cdot \mathrm{p\_Lig} \cdot \mathrm{p\_lig\_bsc}+\mathrm{d\_Lig} \cdot \mathrm{d\_lig} \cdot \mathrm{init\_ep} \cdot \mathrm{k\_hes7\_i\_delta}+\mathrm{d\_Lig} \cdot \mathrm{d\_lig} \cdot \mathrm{init\_isc} \cdot \mathrm{k\_hes7\_i\_delta}+\mathrm{d\_Lig} \cdot \mathrm{d\_lig} \cdot \mathrm{init\_mpp} \cdot \mathrm{k\_hes7\_i\_delta}+\mathrm{init\_mpp} \cdot \mathrm{p\_Lig} \cdot \mathrm{p\_lig\_bsc} \cdot \mathrm{p\_lig\_mpp\_factor}+\mathrm{init\_isc} \cdot \mathrm{p\_Lig} \cdot \mathrm{p\_lig\_bsc} \cdot \mathrm{p\_lig\_isc\_factor} \cdot \mathrm{p\_lig\_mpp\_factor}}{\mathrm{d\_Lig} \cdot \mathrm{d\_lig} \cdot (\mathrm{init\_ep}+\mathrm{init\_isc}+\mathrm{init\_mpp})}\right)^{\mathrm{h\_hes7\_i}}}$$

$$\mathrm{v\_max\_spdef} \rightarrow \frac{\mathrm{d\_spdef} \cdot \mathrm{init\_spdef} \cdot \left(\left(\frac{\mathrm{init\_ep} \cdot \mathrm{p\_Lig} \cdot \mathrm{p\_lig\_bsc}+\mathrm{d\_Lig} \cdot \mathrm{d\_lig} \cdot \mathrm{init\_ep} \cdot \mathrm{k\_spdef\_a\_delta}+\mathrm{d\_Lig} \cdot \mathrm{d\_lig} \cdot \mathrm{init\_isc} \cdot \mathrm{k\_spdef\_a\_delta}+\mathrm{d\_Lig} \cdot \mathrm{d\_lig} \cdot \mathrm{init\_mpp} \cdot \mathrm{k\_spdef\_a\_delta}+\mathrm{init\_mpp} \cdot \mathrm{p\_Lig} \cdot \mathrm{p\_lig\_bsc} \cdot \mathrm{p\_lig\_mpp\_factor}+\mathrm{init\_isc} \cdot \mathrm{p\_Lig} \cdot \mathrm{p\_lig\_bsc} \cdot \mathrm{p\_lig\_isc\_factor} \cdot \mathrm{p\_lig\_mpp\_factor}}{\mathrm{d\_Lig} \cdot \mathrm{d\_lig} \cdot (\mathrm{init\_ep}+\mathrm{init\_isc}+\mathrm{init\_mpp})}\right)^{\mathrm{h\_spdef\_a}}+\left(\frac{\mathrm{p\_Lig} \cdot \mathrm{p\_lig\_bsc} \cdot (\mathrm{init\_ep}+\mathrm{init\_mpp} \cdot \mathrm{p\_lig\_mpp\_factor}+\mathrm{init\_isc} \cdot \mathrm{p\_lig\_isc\_factor} \cdot \mathrm{p\_lig\_mpp\_factor})}{\mathrm{d\_Lig} \cdot \mathrm{d\_lig} \cdot (\mathrm{init\_ep}+\mathrm{init\_isc}+\mathrm{init\_mpp})}\right)^{\mathrm{h\_spdef\_a}}\right)}{\mathrm{s\_spdef} \cdot \left(\frac{\mathrm{p\_Lig} \cdot \mathrm{p\_lig\_bsc} \cdot (\mathrm{init\_ep}+\mathrm{init\_mpp} \cdot \mathrm{p\_lig\_mpp\_factor}+\mathrm{init\_isc} \cdot \mathrm{p\_lig\_isc\_factor} \cdot \mathrm{p\_lig\_mpp\_factor})}{\mathrm{d\_Lig} \cdot \mathrm{d\_lig} \cdot (\mathrm{init\_ep}+\mathrm{init\_isc}+\mathrm{init\_mpp})}\right)^{\mathrm{h\_spdef\_a}}}$$

$$\mathrm{v\_max\_tp63} \rightarrow \mathrm{d\_tp63} \cdot \mathrm{init\_tp63} / \left(\mathrm{s\_tp63} \cdot \left(\frac{\left(\frac{\mathrm{init\_spdef} \cdot \mathrm{p\_Spdef}}{\mathrm{d\_Spdef} \cdot \mathrm{s\_spdef}}\right)^{\mathrm{h\_tp63\_a\_spdef}}}{\left(\frac{\mathrm{init\_spdef} \cdot \mathrm{p\_Spdef}+\mathrm{d\_Spdef} \cdot \mathrm{k\_tp63\_a\_spdef\_delta} \cdot \mathrm{s\_spdef}}{\mathrm{d\_Spdef} \cdot \mathrm{s\_spdef}}\right)^{\mathrm{h\_tp63\_a\_spdef}}+\left(\frac{\mathrm{init\_spdef} \cdot \mathrm{p\_Spdef}}{\mathrm{d\_Spdef} \cdot \mathrm{s\_spdef}}\right)^{\mathrm{h\_tp63\_a\_spdef}}} + \frac{\left(\frac{\mathrm{p\_Lig} \cdot \mathrm{p\_lig\_bsc} \cdot (\mathrm{init\_ep}+\mathrm{init\_mpp} \cdot \mathrm{p\_lig\_mpp\_factor}+\mathrm{init\_isc} \cdot \mathrm{p\_lig\_isc\_factor} \cdot \mathrm{p\_lig\_mpp\_factor})}{\mathrm{d\_Lig} \cdot \mathrm{d\_lig} \cdot (\mathrm{init\_ep}+\mathrm{init\_isc}+\mathrm{init\_mpp})}\right)^{\mathrm{h\_tp63\_a}}}{\left(\frac{\mathrm{init\_ep} \cdot \mathrm{p\_Lig} \cdot \mathrm{p\_lig\_bsc}+\mathrm{d\_Lig} \cdot \mathrm{d\_lig} \cdot \mathrm{init\_ep} \cdot \mathrm{k\_tp63\_a\_delta}+\mathrm{d\_Lig} \cdot \mathrm{d\_lig} \cdot \mathrm{init\_isc} \cdot \mathrm{k\_tp63\_a\_delta}+\mathrm{d\_Lig} \cdot \mathrm{d\_lig} \cdot \mathrm{init\_mpp} \cdot \mathrm{k\_tp63\_a\_delta}+\mathrm{init\_mpp} \cdot \mathrm{p\_Lig} \cdot \mathrm{p\_lig\_bsc} \cdot \mathrm{p\_lig\_mpp\_factor}+\mathrm{init\_isc} \cdot \mathrm{p\_Lig} \cdot \mathrm{p\_lig\_bsc} \cdot \mathrm{p\_lig\_isc\_factor} \cdot \mathrm{p\_lig\_mpp\_factor}}{\mathrm{d\_Lig} \cdot \mathrm{d\_lig} \cdot (\mathrm{init\_ep}+\mathrm{init\_isc}+\mathrm{init\_mpp})}\right)^{\mathrm{h\_tp63\_a}}+\left(\frac{\mathrm{p\_Lig} \cdot \mathrm{p\_lig\_bsc} \cdot (\mathrm{init\_ep}+\mathrm{init\_mpp} \cdot \mathrm{p\_lig\_mpp\_factor}+\mathrm{init\_isc} \cdot \mathrm{p\_lig\_isc\_factor} \cdot \mathrm{p\_lig\_mpp\_factor})}{\mathrm{d\_Lig} \cdot \mathrm{d\_lig} \cdot (\mathrm{init\_ep}+\mathrm{init\_isc}+\mathrm{init\_mpp})}\right)^{\mathrm{h\_tp63\_a}}}\right)\right)$$

#### 1.7 Experiment: RNAseq_data_WT_controls_correctedTimes

##### 1.7.1 Comments

Time-resolved RNAseq data for the most important molecular players in the mdoelled mucociliary tissue.

##### 1.7.2 Observables

The following observables are modified in this data set.

- **Observable:** foxi1

$$\text{foxi1(t)} = \log_{10}(\text{s\_isc} \cdot \left(\frac{[\text{isc}]}{[\text{n\_cells}]} + \frac{[\text{mpp}] \cdot \text{s\_mpp\_foxi1}}{[\text{n\_cells}]}\right)) \tag{80}$$

$$\sigma\{\text{foxi1}\}(\text{t}) = \text{sd\_tpm\_foxi1} \tag{81}$$

- **Observable:** mcidas

$$\text{mcidas(t)} = \log_{10}(\text{background\_mcidas} + \frac{[\text{mcc}] \cdot \text{s\_mcc}}{[\text{n\_cells}]}) \tag{82}$$

$$\sigma\{\text{mcidas}\}(\text{t}) = \text{sd\_tpm\_mcidas} \tag{83}$$

- **Observable:** foxa1

$$\text{foxa1(t)} = \log_{10}(\text{background\_foxa1} + \frac{\text{s\_ssc} \cdot [\text{ssc}]}{[\text{n\_cells}]}) \tag{84}$$

$$\sigma\{\text{foxa1}\}(\text{t}) = \text{sd\_tpm\_foxa1} \tag{85}$$

- **Observable:** lig

$$[\text{lig}](\text{t}) = \log_{10}([\text{lig}] \cdot \text{s\_lig}) \tag{86}$$

$$\sigma\{[\text{lig}]\}(\text{t}) = \text{sd\_tpm\_ligands\_tot} \tag{87}$$

- **Observable:** hes4

$$[\text{hes4}](\text{t}) = \log_{10}([\text{hes4}] \cdot \text{s\_hes4}) \tag{88}$$

$$\sigma\{[\text{hes4}]\}(\text{t}) = \text{sd\_tpm\_hes4} \tag{89}$$

- **Observable:** hes5

$$[\text{hes5}](\text{t}) = \log_{10}([\text{hes5}] \cdot \text{s\_hes5}) \tag{90}$$

$$\sigma\{[\text{hes5}]\}(\text{t}) = \text{sd\_tpm\_hes5} \tag{91}$$

- **Observable:** hes7

$$[\text{hes7}](\text{t}) = \log_{10}([\text{hes7}] \cdot \text{s\_hes7}) \tag{92}$$

$$\sigma\{[\text{hes7}]\}(\text{t}) = \text{sd\_tpm\_hes7} \tag{93}$$

- **Observable:** tp63

$$[\text{tp63}](\text{t}) = \log_{10}(\text{s\_tp63} \cdot [\text{tp63}]) \tag{94}$$

$$\sigma\{[\text{tp63}]\}(\text{t}) = \text{sd\_tpm\_tp63} \tag{95}$$

- **Observable:** spdef

$$[\text{spdef}](\text{t}) = \log_{10}(\text{s\_spdef} \cdot [\text{spdef}]) \tag{96}$$

$$\sigma\{[\text{spdef}]\}(\text{t}) = \text{sd\_tpm\_spdef} \tag{97}$$

| time [min] | foxi1 [tpm] | mcidas [tpm] | foxa1 [tpm] | lig [tpm] | hes4 [tpm] | hes5 [tpm] | hes7 [tpm] | tp63 [tpm] | spdef [tpm] |
|---|---|---|---|---|---|---|---|---|---|
| 0 | 21.4122 | 5.63667 | 4.01337 | 19.3056 | 58.2492 | 8.03 | 228.636 | 5.21332 | 0.290805 |
| 0 | 3.73601 | 2.72262 | 6.49646 | 20.66 | 117.69 | 13.0757 | 308.907 | 0.688885 | 4.33647 |
| 110.799 | 138.971 | 3.067 | 1.54264 | 77.022 | 385.925 | 119.701 | 26.119 | 0.776347 | 3.20894 |
| 98.982 | 125.131 | 4.70885 | 2.00375 | 87.2581 | 354.278 | 95.7696 | 12.3979 | 0.59586 | 3.28351 |
| 98.748 | 15.7956 | 1.12522 | 1.57711 | 42.0192 | 587.846 | 136.407 | 75.7261 | 0.8056 | 2.69489 |
| 106.236 | 242.094 | 8.74825 | 1.5337 | 100.201 | 169.706 | 6.45534 | 0.55149 | 0.641397 | 6.06226 |
| 174.796 | 127.625 | 4.70015 | 2.05203 | 63.4676 | 431.103 | 49.7569 | 8.41104 | 0.804862 | 5.66696 |
| 249.7 | 94.1405 | 8.29909 | 2.08001 | 54.0713 | 455.313 | 32.8595 | 5.49451 | 0.760972 | 1.50088 |
| 181.282 | 77.2064 | 0.879445 | 1.60553 | 27.4377 | 321.448 | 39.3183 | 21.3358 | 0.259333 | 0.686017 |
| 243.1 | 172.065 | 9.72742 | 1.1865 | 93.9638 | 238.221 | 158.562 | 5.81861 | 0.820473 | 4.31496 |
| 227.098 | 224.839 | 25.6547 | 1.25839 | 123.09 | 297.602 | 135.976 | 3.79494 | 0.783333 | 4.7794 |
| 399.176 | 453.483 | 145.236 | 1.40541 | 180.745 | 291.355 | 174.592 | 7.44881 | 1.56658 | 11.6615 |
| 365.352 | 304.528 | 85.8069 | 1.41914 | 146.701 | 171.716 | 276.509 | 2.0362 | 1.38043 | 7.8565 |
| 754.557 | 271.455 | 494.72 | 64.7823 | 115.075 | 319.437 | 109.47 | 12.0144 | 115.268 | 15.3678 |
| 796.611 | 321.149 | 302.605 | 25.9136 | 72.1299 | 157.068 | 5.03945 | 1.05558 | 134.161 | 17.6462 |
| 741.485 | 239.25 | 295.523 | 67.2722 | 237.103 | 271.849 | 117.662 | 0.232522 | 92.876 | 16.6473 |
| 801.88 | 229.908 | 224.015 | 5.97821 | 25.2432 | 244.478 | 9.11634 | 0.22395 | 33.1506 | 6.4619 |
| 822.284 | 343.726 | 437.127 | 13.7004 | 70.0158 | 142.549 | 16.9482 | 0.261772 | 80.1973 | 8.65471 |
| 1205.47 | 195.257 | 4.33881 | 21.2258 | 24.3812 | 210.398 | 69.681 | 0.170823 | 205.952 | 1.82997 |
| 1240.3 | 278.599 | 4.45848 | 25.328 | 34.7456 | 79.1663 | 0.357583 | NaN | 158.667 | 3.76917 |
| 1192.71 | 220.195 | 8.22757 | 13.7736 | 9.01661 | 82.3656 | 1.90134 | 0.0895419 | 188.719 | 4.53755 |
| 1240.3 | 233.701 | 3.55422 | 9.9331 | 21.8985 | 110.998 | 7.04825 | 0.116594 | 174.176 | 1.55596 |
| 1240.3 | 244.843 | 17.3731 | 10.5134 | 19.673 | 99.3332 | 1.86039 | 0.187399 | 154.876 | 1.35181 |

**Table 1: Experimental data for the experiment RNAseq_data_WT_controls_correctedTimes**

##### 1.7.3 Experiment specific conditions

To evaluate the model for this experiment the following conditions are applied.

- **Local condition #1:**

$$\text{background\_foxa1} \rightarrow \text{init\_foxa1}$$
$$\text{background\_mcidas} \rightarrow \text{init\_mcidas}$$
$$\text{s\_lig} \rightarrow \frac{\text{d\_lig} \cdot \text{init\_lig} \cdot (\text{init\_ep} + \text{init\_isc} + \text{init\_mpp})}{\text{p\_lig\_bsc} \cdot (\text{init\_ep} + \text{init\_mpp} \cdot \text{p\_lig\_mpp\_factor} + \text{init\_isc} \cdot \text{p\_lig\_isc\_factor} \cdot \text{p\_lig\_mpp\_factor})}$$

##### 1.7.4 Experimental data and model fit

The model observables and the experimental data is shown in Figure 1. The agreement of the model observables and the experimental data, given in Table 1, yields a value of the objective function $\chi^2 = -211.726$ for 206 data points in this data set.

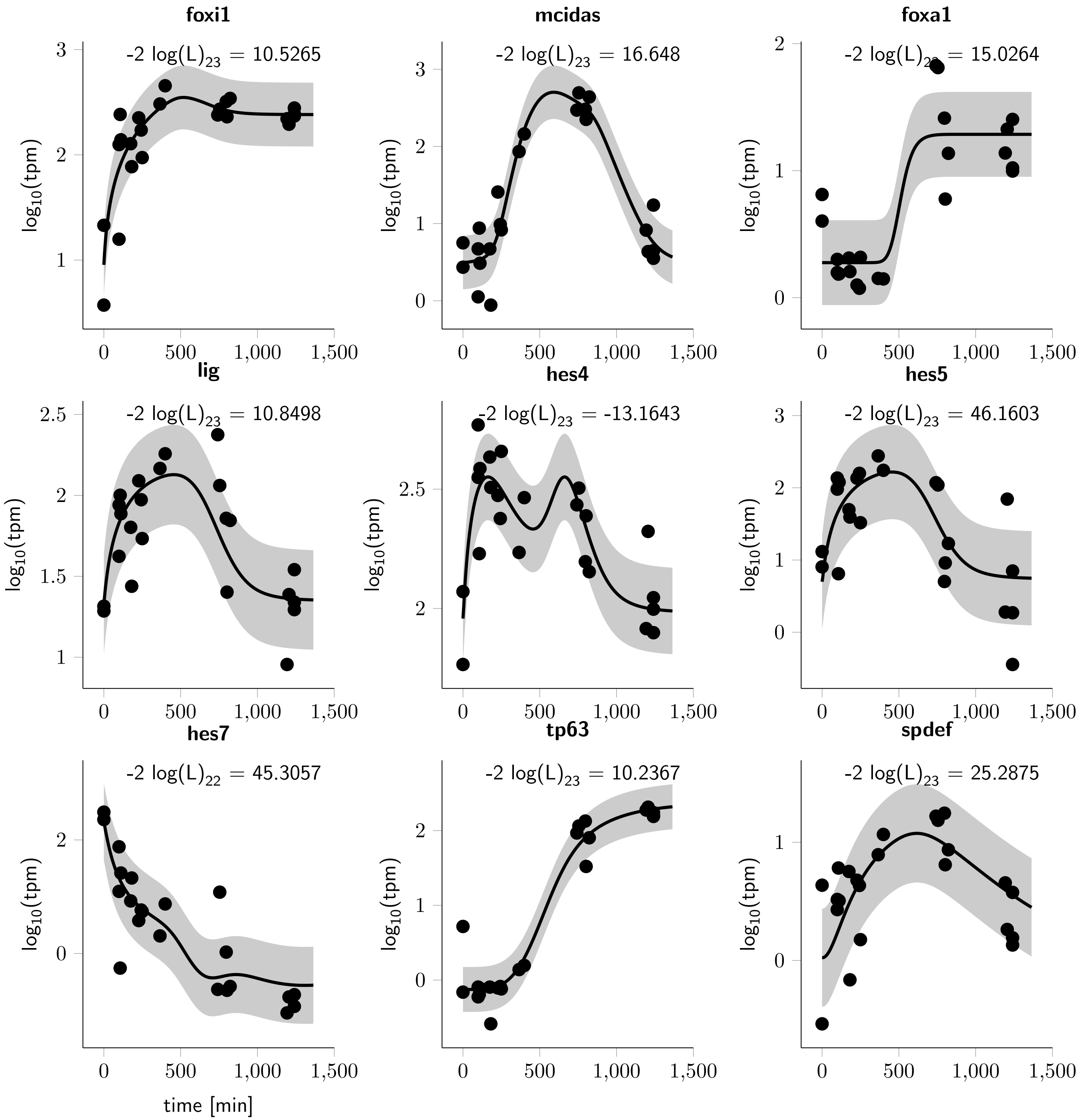

**Figure 1: RNAseq_data_WT_controls_correctedTimes observables and experimental data for the experiment.** The observables are displayed as solid lines. The error model that describes the measurement noise is indicated by shades.

| time [min] | isc_abs [number of cells] | mcc_abs [number of cells] | ssc_abs [number of cells] |
|---|---|---|---|
| 1240.3 | 43.0625 | 40.5 | 27.875 |

**Table 2: Experimental data for the experiment cellQuant_data_ctrl_absNums**

#### 1.8 Experiment: cellQuant_data_ctrl_absNums

##### 1.8.1 Comments

Cell counts for the specified cell types from immunoflourescence images.

##### 1.8.2 Observables

The following observables are modified in this data set.

- **Observable:** isc_abs

$$\mathrm{isc\_abs(t)} = [\mathrm{isc}] \quad (98)$$

$$\sigma\{\mathrm{isc\_abs}\}(\mathrm{t}) = 1 \quad (99)$$

- **Observable:** mcc_abs

$$\mathrm{mcc\_abs(t)} = [\mathrm{mcc\_total}] \quad (100)$$

$$\sigma\{\mathrm{mcc\_abs}\}(\mathrm{t}) = 1 \quad (101)$$

- **Observable:** ssc_abs

$$\mathrm{ssc\_abs(t)} = [\mathrm{ssc}] \quad (102)$$

$$\sigma\{\mathrm{ssc\_abs}\}(\mathrm{t}) = 1 \quad (103)$$

##### 1.8.3 Experimental data and model fit

The agreement of the model observables and the experimental data, given in Table 2, yields a value of the objective function $\chi^2 = 4.87169$ for 3 data points in this data set.

| time [min] | bc_frac au [au] | mcc_abs au [au] |
|---|---|---|
| 1240.3 | 0 | NaN |
| 180 | NaN | 0 |

**Table 3: Prior information**

#### 1.9 Constraints for cell composition

##### 1.9.1 Comments

A prior data point is introduced ensuring that finally the multipotent progenitors have to decide for one cell fate. This prior information was integrated as a penalty, i.e. like a data point with small uncertainty.

##### 1.9.2 Observables

The following observables are defined in this prior information.

- **Observable:** bsc2_frac

$$\mathrm{bsc2\_frac(t)} = \log_{10}\left([\mathrm{bc}] + 0.01\right) - \log_{10}\left([\mathrm{bc}] + [\mathrm{ep}] + [\mathrm{mpp}] + 0.01\right) \tag{104}$$

$$\sigma\{\mathrm{bsc2\_frac}\}(\mathrm{t}) = 0.0001 \tag{105}$$

- **Observable:** mcc_abs

$$\mathrm{mcc\_abs(t)} = [\mathrm{mcc\_total}] \tag{106}$$

$$\sigma\{\mathrm{mcc\_abs}\}(\mathrm{t}) = 0.0001 \tag{107}$$

##### 1.9.3 Experimental prior and model fit

The agreement of the model observables and the prior, given in Table 3, yields a value of the objective function $\chi^2 = -7.71213$ for 1 data points in this data set.

#### 2 Estimated model parameters

In total 89 parameters are estimated from the experimental data. The best fit yields a value of the objective function $-2\log(L) = 258.67$ for a total of 210 data points. The model parameters were estimated by maximum likelihood estimation. In Table 4 – 5 the estimated parameter values are given. Parameters highlighted in red color indicate parameter values close to their bounds. The parameter name prefix init_ indicates the initial value of a dynamic variable.

| | name | $\theta_{min}$ | $\hat{\theta}$ | $\theta_{max}$ | log | non-log $\hat{\theta}$ | estimated |
|---|---|---|---|---|---|---|---|
| 1 | d_hes4 | -5 | +2.9999 | +3 | 1 | $+1.00 \cdot 10^{+03}$ | 1 |
| 2 | d_hes4_d | -5 | +0.5361 | +3 | 1 | $+3.44 \cdot 10^{+00}$ | 1 |
| 3 | d_hes5 | -5 | +0.6605 | +3 | 1 | $+4.58 \cdot 10^{+00}$ | 1 |
| 4 | d_hes5_d | -5 | +0.3789 | +3 | 1 | $+2.39 \cdot 10^{+00}$ | 1 |
| 5 | d_hes7 | -5 | +2.9987 | +3 | 1 | $+9.97 \cdot 10^{+02}$ | 1 |
| 6 | d_hes7_d | -5 | -3.3104 | +3 | 1 | $+4.89 \cdot 10^{-04}$ | 1 |
| 7 | d_lig | -5 | +0.6448 | +3 | 1 | $+4.41 \cdot 10^{+00}$ | 1 |
| 8 | d_Spdef | -5 | -3.5018 | +0 | 1 | $+3.15 \cdot 10^{-04}$ | 1 |
| 9 | d_spdef | -5 | -2.5138 | +3 | 1 | $+3.06 \cdot 10^{-03}$ | 1 |
| 10 | d_tp63 | -5 | -2.5007 | +3 | 1 | $+3.16 \cdot 10^{-03}$ | 1 |
| 11 | d_Lig | -3 | -0.0000 | +0 | 1 | $+1.00 \cdot 10^{+00}$ | 1 |
| 12 | hes_delay | -5 | +2.6730 | +3 | 1 | $+4.71 \cdot 10^{+02}$ | 1 |
| 13 | h_cell_i | +0 | +1.0000 | +1 | 1 | $+1.00 \cdot 10^{+01}$ | 1 |
| 14 | h_hes4_i | +0 | +0.7311 | +1 | 1 | $+5.38 \cdot 10^{+00}$ | 1 |
| 15 | h_hes4_a | +0 | +0.1326 | +1 | 1 | $+1.36 \cdot 10^{+00}$ | 1 |
| 16 | h_hes5_a | +0 | +0.2786 | +1 | 1 | $+1.90 \cdot 10^{+00}$ | 1 |
| 17 | h_hes7_i | +0 | +0.3996 | +1 | 1 | $+2.51 \cdot 10^{+00}$ | 1 |
| 18 | h_hes7_i_spdef | +0 | +0.9999 | +1 | 1 | $+1.00 \cdot 10^{+01}$ | 1 |
| 19 | h_isc_i_hes4 | +0 | +0.4489 | +1 | 1 | $+2.81 \cdot 10^{+00}$ | 1 |
| 20 | h_isc_i_hes5 | +0 | +0.9765 | +1 | 1 | $+9.47 \cdot 10^{+00}$ | 1 |
| 21 | h_isc_a | +0 | +0.3744 | +1 | 1 | $+2.37 \cdot 10^{+00}$ | 1 |
| 22 | h_mcc_i_hes5 | +0 | +0.7002 | +1 | 1 | $+5.01 \cdot 10^{+00}$ | 1 |
| 23 | h_mcc_a | +0 | +0.8016 | +1 | 1 | $+6.33 \cdot 10^{+00}$ | 1 |
| 24 | h_spdef_a | +0 | +0.1792 | +1 | 1 | $+1.51 \cdot 10^{+00}$ | 1 |
| 25 | h_ssc_i_hes4 | +0 | +0.4238 | +1 | 1 | $+2.65 \cdot 10^{+00}$ | 1 |
| 26 | h_ssc_i_spdef | +0 | +1.0000 | +1 | 1 | $+1.00 \cdot 10^{+01}$ | 1 |
| 27 | h_ssc_a | +0 | +0.9981 | +1 | 1 | $+9.96 \cdot 10^{+00}$ | 1 |
| 28 | h_ssc_a_spdef | +0 | +0.9664 | +1 | 1 | $+9.26 \cdot 10^{+00}$ | 1 |
| 29 | h_tp63_a | +0 | +0.9393 | +1 | 1 | $+8.70 \cdot 10^{+00}$ | 1 |
| 30 | h_tp63_a_spdef | +0 | +0.9074 | +1 | 1 | $+8.08 \cdot 10^{+00}$ | 1 |
| 31 | init_ep | +0 | +3.0000 | +3 | 1 | $+1.00 \cdot 10^{+03}$ | 1 |
| 32 | init_foxa1 | +0 | +1.8881 | +3 | 0 | $+1.89 \cdot 10^{+00}$ | 1 |
| 33 | init_hes4 | +5e+01 | +90.7147 | +1e+02 | 0 | $+9.07 \cdot 10^{+01}$ | 1 |
| 34 | init_hes5 | +5 | +5.0001 | +2e+01 | 0 | $+5.00 \cdot 10^{+00}$ | 1 |
| 35 | init_hes7 | +2e+02 | +201.0276 | +4e+02 | 0 | $+2.01 \cdot 10^{+02}$ | 1 |
| 36 | init_isc | -5 | +0.1612 | +0.7 | 1 | $+1.45 \cdot 10^{+00}$ | 1 |
| 37 | init_lig | +2e+01 | +21.2875 | +2e+01 | 0 | $+2.13 \cdot 10^{+01}$ | 1 |
| 38 | init_mcidas | +1 | +3.1395 | +6 | 0 | $+3.14 \cdot 10^{+00}$ | 1 |
| 39 | init_mpp | -5 | -3.9264 | +3 | 1 | $+1.18 \cdot 10^{-04}$ | 1 |
| 40 | init_spdef | -5 | +0.0232 | +3 | 1 | $+1.05 \cdot 10^{+00}$ | 1 |
| 41 | init_tp63 | -0.5 | -0.1259 | +0.2 | 1 | $+7.48 \cdot 10^{-01}$ | 1 |
| 42 | k_cell_i | -5 | +0.2319 | +3 | 1 | $+1.71 \cdot 10^{+00}$ | 1 |
| 43 | k_hes4_i_delta | -5 | -3.8454 | +3 | 1 | $+1.43 \cdot 10^{-04}$ | 1 |
| 44 | k_hes4_a_delta | -7 | -0.7017 | +3 | 1 | $+1.99 \cdot 10^{-01}$ | 1 |
| 45 | k_hes5_a_delta | -5 | +2.9636 | +3 | 1 | $+9.20 \cdot 10^{+02}$ | 1 |
| 46 | k_hes7_i_delta | -7 | -6.8845 | +3 | 1 | $+1.30 \cdot 10^{-07}$ | 1 |

**Table 4: Estimated parameter values**
$\hat{\theta}$ indicates the estimated value of the parameters. $\theta_{min}$ and $\theta_{max}$ indicate the upper and lower bounds for the parameters. The log-column indicates whether a parameter was log-transformed for parameter estimation and uncertainty analysis. If log $\equiv$ 1 the non-log-column indicates the non-logarithmic value of the estimate. The estimated-column indicates if the parameter value was estimated (1).

| | name | $\theta_{min}$ | $\hat{\theta}$ | $\theta_{max}$ | log | non-log $\hat{\theta}$ | estimated |
|---|---|---|---|---|---|---|---|
| 47 | sd_tpm_mcidas | -5 | -0.4605 | +3 | 1 | $+3.46 \cdot 10^{-01}$ | 1 |
| 48 | sd_tpm_spdef | -5 | -0.3813 | +3 | 1 | $+4.16 \cdot 10^{-01}$ | 1 |
| 49 | sd_tpm_tp63 | -5 | -0.5216 | +3 | 1 | $+3.01 \cdot 10^{-01}$ | 1 |
| 50 | k_hes7_i_spdef_delta | -5 | +2.9045 | +3 | 1 | $+8.03 \cdot 10^{+02}$ | 1 |
| 51 | k_isc_i_hes4_delta | -5 | +2.4573 | +3 | 1 | $+2.87 \cdot 10^{+02}$ | 1 |
| 52 | k_isc_i_hes5_delta | -5 | -0.0801 | +3 | 1 | $+8.32 \cdot 10^{-01}$ | 1 |
| 53 | k_isc_a | -5 | +3.5723 | +4 | 1 | $+3.74 \cdot 10^{+03}$ | 1 |
| 54 | k_mcc_i_hes5_delta | -5 | +2.7790 | +3 | 1 | $+6.01 \cdot 10^{+02}$ | 1 |
| 55 | k_mcc_a | -5 | +0.9750 | +3 | 1 | $+9.44 \cdot 10^{+00}$ | 1 |
| 56 | k_spdef_a_delta | -5 | +2.0526 | +3 | 1 | $+1.13 \cdot 10^{+02}$ | 1 |
| 57 | k_ssc_i_hes4_delta | -5 | +2.9002 | +3 | 1 | $+7.95 \cdot 10^{+02}$ | 1 |
| 58 | k_ssc_i_spdef_delta | -5 | -0.1586 | +3 | 1 | $+6.94 \cdot 10^{-01}$ | 1 |
| 59 | k_ssc_a | -5 | +0.0454 | +3 | 1 | $+1.11 \cdot 10^{+00}$ | 1 |
| 60 | k_ssc_a_spdef_delta | -5 | +1.7781 | +3 | 1 | $+6.00 \cdot 10^{+01}$ | 1 |
| 61 | k_tp63_a_delta | -5 | +2.4913 | +3 | 1 | $+3.10 \cdot 10^{+02}$ | 1 |
| 62 | k_tp63_a_spdef_delta | -5 | +3.2543 | +6 | 1 | $+1.80 \cdot 10^{+03}$ | 1 |
| 63 | p_lig_bsc | -5 | -0.5317 | +4 | 1 | $+2.94 \cdot 10^{-01}$ | 1 |
| 64 | p_lig_bc_factor | -5 | -1.1530 | +0 | 1 | $+7.03 \cdot 10^{-02}$ | 1 |
| 65 | p_lig_isc_factor | -5 | -0.0003 | +0 | 1 | $+9.99 \cdot 10^{-01}$ | 1 |
| 66 | p_lig_mcc_factor | -5 | -0.0417 | +0 | 1 | $+9.08 \cdot 10^{-01}$ | 1 |
| 67 | p_lig_mpp_factor | +0 | +0.8918 | +3 | 1 | $+7.79 \cdot 10^{+00}$ | 1 |
| 68 | p_lig_ssc_factor | -5 | -2.2787 | +0 | 1 | $+5.26 \cdot 10^{-03}$ | 1 |
| 69 | p_mcc_late | -3 | -1.9092 | -1 | 1 | $+1.23 \cdot 10^{-02}$ | 1 |
| 70 | p_mpp | -7 | -2.4034 | -1 | 1 | $+3.95 \cdot 10^{-03}$ | 1 |
| 71 | p_Spdef | -5 | -3.0131 | +3 | 1 | $+9.70 \cdot 10^{-04}$ | 1 |
| 72 | p_Lig | -5 | -0.2068 | +3 | 1 | $+6.21 \cdot 10^{-01}$ | 1 |
| 73 | v_max | -5 | -0.8104 | +3 | 1 | $+1.55 \cdot 10^{-01}$ | 1 |
| 74 | v_max_bc | -5 | -1.8857 | +3 | 1 | $+1.30 \cdot 10^{-02}$ | 1 |
| 75 | s_hes4 | -5 | -0.7141 | +3 | 1 | $+1.93 \cdot 10^{-01}$ | 1 |
| 76 | s_hes5 | -5 | -0.4943 | +3 | 1 | $+3.20 \cdot 10^{-01}$ | 1 |
| 77 | s_hes7 | -5 | +1.2324 | +3 | 1 | $+1.71 \cdot 10^{+01}$ | 1 |
| 78 | s_isc | -5 | +3.7943 | +4 | 1 | $+6.23 \cdot 10^{+03}$ | 1 |
| 79 | s_mcc | -5 | +4.8292 | +5 | 1 | $+6.75 \cdot 10^{+04}$ | 1 |
| 80 | s_mpp_foxi1 | +0 | +0.0243 | +1 | 0 | $+2.43 \cdot 10^{-02}$ | 1 |
| 81 | s_spdef | -5 | -2.7174 | +3 | 1 | $+1.92 \cdot 10^{-03}$ | 1 |
| 82 | s_ssc | -5 | +2.8422 | +4 | 1 | $+6.95 \cdot 10^{+02}$ | 1 |
| 83 | s_tp63 | -5 | +1.7951 | +3 | 1 | $+6.24 \cdot 10^{+01}$ | 1 |
| 84 | sd_tpm_foxa1 | -5 | -0.4761 | +3 | 1 | $+3.34 \cdot 10^{-01}$ | 1 |
| 85 | sd_tpm_foxi1 | -5 | -0.5182 | +3 | 1 | $+3.03 \cdot 10^{-01}$ | 1 |
| 86 | sd_tpm_hes4 | -5 | -0.7415 | +3 | 1 | $+1.81 \cdot 10^{-01}$ | 1 |
| 87 | sd_tpm_hes5 | -5 | -0.1853 | +3 | 1 | $+6.53 \cdot 10^{-01}$ | 1 |
| 88 | sd_tpm_hes7 | -5 | -0.1688 | +3 | 1 | $+6.78 \cdot 10^{-01}$ | 1 |
| 89 | sd_tpm_lig | -5 | -0.5123 | +3 | 1 | $+3.07 \cdot 10^{-01}$ | 1 |

**Table 5: Estimated parameter values (CONT)**

$\hat{\theta}$ indicates the estimated value of the parameters. $\theta_{min}$ and $\theta_{max}$ indicate the upper and lower bounds for the parameters. The log-column indicates whether a parameter was log-transformed for parameter estimation and uncertainty analysis. If log $\equiv 1$ the non-log-column indicates the non-logarithmic value of the estimate. The estimated-column indicates if the parameter value was estimated (1).

### 3 Uncertainty analysis of model parameters by the profile likelihood

In order to evaluate uncertainties of the estimated model parameters, to derive confidence intervals and assess identifiability, the profile likelihood was calculated for all parameters. The profile likelihood of a parameter is calculated by fixing the value of this parameter to a specific number and then refitting all remaining parameters. This is repeated to find all values where the refitted likelihood is below the significance threshold [2].

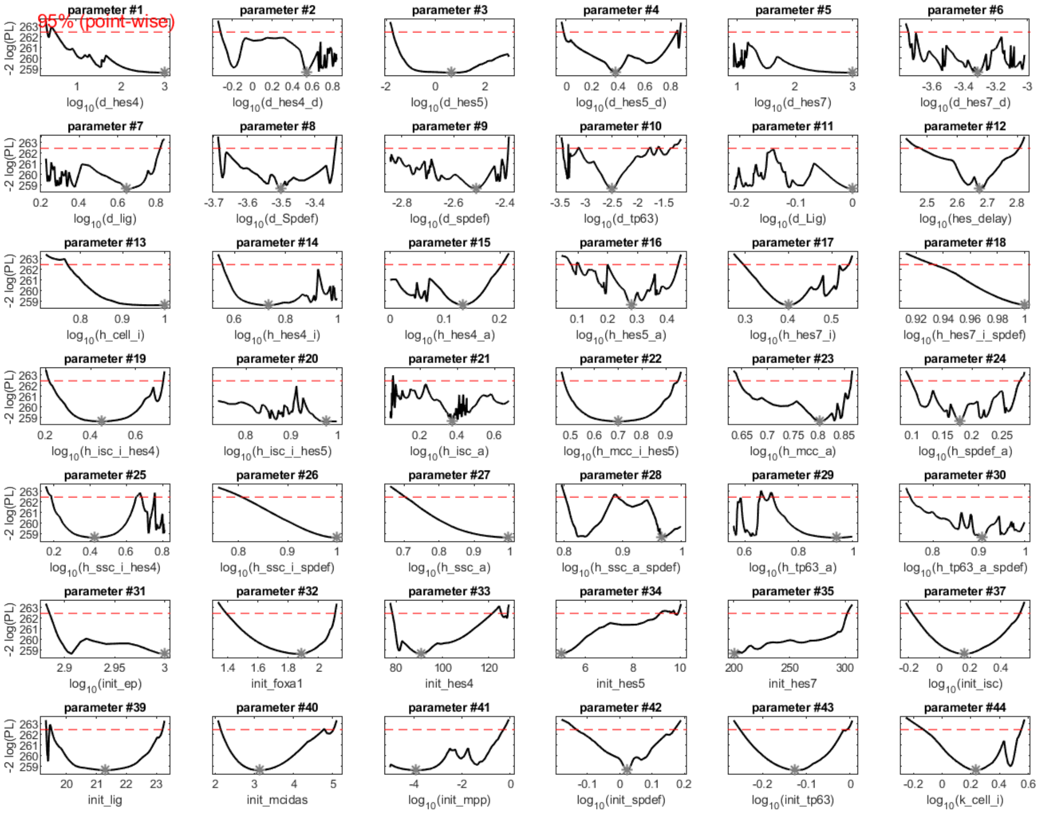

**Figure 2:** Profile likelihood for the estimated model parameters. The horizontal axis denote the parameter values, the vertical axis is $-2\log(\text{Likelihood})$. The minima of the curves indicate to the maximum likelihood estimates. 95% confidence intervals are given by the intersection of the profile likelihood with the thresholds depicted as red dashed lines.

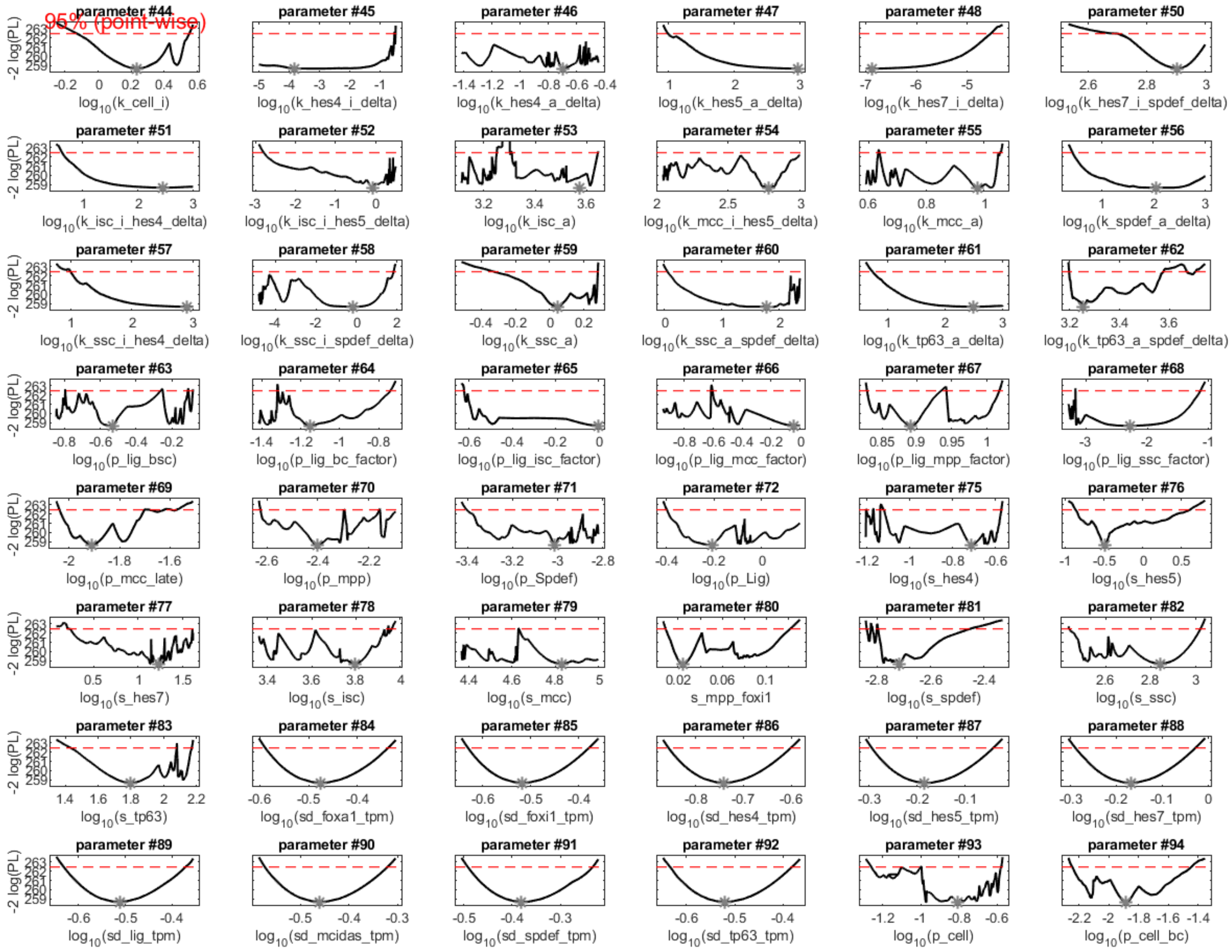

**Figure 3:** Profile likelihood for the estimated model parameters. The horizontal axis denote the parameter values, the vertical axis is $-2\log(\text{Likelihood})$. The minima of the curves indicate to the maximum likelihood estimates. 95% confidence intervals are given by the intersection of the profile likelihood with the thresholds depicted as red dashed lines.

| | name | $\hat{\theta}$ | $\sigma^-$ | $\sigma^+$ |
|---|---|---|---|---|
| 1 | d_hes4 | +3.000 | +0.342 | +Inf |
| 2 | d_hes4_d | +0.536 | -0.316 | +Inf |
| 3 | d_hes5 | +0.660 | -1.734 | +Inf |
| 4 | d_hes5_d | +0.379 | -0.019 | +0.867 |
| 5 | d_hes7 | +2.999 | -Inf | +Inf |
| 6 | d_hes7_d | -3.310 | -3.746 | +Inf |
| 7 | d_lig | +0.645 | -Inf | +0.820 |
| 8 | d_Spdef | -3.502 | -3.684 | -3.340 |
| 9 | d_spdef | -2.514 | -Inf | -2.389 |
| 10 | d_tp63 | -2.501 | -3.444 | -1.343 |
| 11 | d_Lig | -0.000 | -Inf | +Inf |
| 12 | hes_delay | +2.673 | +2.474 | +2.806 |
| 13 | h_cell_i | +1.000 | +0.768 | +Inf |
| 14 | h_hes4_i | +0.731 | +0.552 | +Inf |
| 15 | h_hes4_a | +0.133 | -Inf | +0.204 |
| 16 | h_hes5_a | +0.279 | +0.076 | +0.438 |
| 17 | h_hes7_i | +0.400 | +0.295 | +0.540 |
| 18 | h_hes7_i_spdef | +1.000 | +0.932 | +Inf |
| 19 | h_isc_i_hes4 | +0.449 | +0.215 | +0.724 |
| 20 | h_isc_i_hes5 | +0.976 | -Inf | +Inf |
| 21 | h_isc_a | +0.374 | -Inf | +Inf |
| 22 | h_mcc_i_hes5 | +0.700 | +0.471 | +0.952 |
| 23 | h_mcc_a | +0.802 | +0.644 | +0.863 |
| 24 | h_spdef_a | +0.179 | +0.098 | +0.280 |
| 25 | h_ssc_i_hes4 | +0.424 | +0.184 | +Inf |
| 26 | h_ssc_i_spdef | +1.000 | +0.806 | +Inf |
| 27 | h_ssc_a | +0.998 | +0.703 | +Inf |
| 28 | h_ssc_a_spdef | +0.966 | +0.802 | +Inf |
| 29 | h_tp63_a | +0.939 | -Inf | +Inf |

**Table 6: Confidence intervals for the estimated parameter values derived by the profile likelihood**
$\hat{\theta}$ indicates the estimated optimal parameter value. $\sigma^-$ and $\sigma^+$ indicate 95% point-wise confidence intervals.

### 4 Confidence intervals for the model parameters

In Table 6 – 8, 95% confidence intervals for the estimated parameter values derived by the profile likelihood [2] are given.

### References

[1] Alan C Hindmarsh, Peter N Brown, Keith E Grant, Steven L Lee, Radu Serban, Dan E Shumaker, and Carol S Woodward. SUNDIALS: Suite of nonlinear and differential/algebraic equation solvers. *ACM Transactions on Mathematical Software*, 31(3):363–396, sep 2005.

[2] A Raue, C Kreutz, T Maiwald, J Bachmann, M Schilling, U Klingmüller, and J Timmer. Structural and practical identifiability analysis of partially observed dynamical models by exploiting the profile likelihood. *Bioinformatics*, 25(15):1923–1929, 2009.

| | name | $\hat{\theta}$ | $\sigma^-$ | $\sigma^+$ |
|---|---|---|---|---|
| 30 | h_tp63_a_spdef | +0.907 | +0.751 | +Inf |
| 31 | init_ep | +3.000 | +2.885 | +Inf |
| 32 | init_foxa1 | +1.888 | +1.387 | +2.113 |
| 33 | init_hes4 | +90.715 | +78.331 | +128.117 |
| 34 | init_hes5 | +5.000 | -Inf | +9.920 |
| 35 | init_hes7 | +201.028 | -Inf | +301.798 |
| 37 | init_isc | +0.161 | -0.177 | +0.525 |
| 39 | init_lig | +21.287 | +19.330 | +23.213 |
| 40 | init_mcidas | +3.139 | +2.142 | +5.027 |
| 41 | init_mpp | -3.926 | -Inf | -0.263 |
| 42 | init_spdef | +0.023 | -0.135 | +0.170 |
| 43 | init_tp63 | -0.126 | -0.253 | -0.006 |
| 44 | k_cell_i | +0.232 | -0.133 | +0.552 |
| 45 | k_hes4_i_delta | -3.845 | -Inf | -0.495 |
| 46 | k_hes4_a_delta | -0.702 | -Inf | +Inf |
| 47 | k_hes5_a_delta | +2.964 | +0.988 | +Inf |
| 48 | k_hes7_i_delta | -6.885 | -Inf | -4.535 |
| 50 | k_hes7_i_spdef_delta | +2.905 | +2.670 | +Inf |
| 51 | k_isc_i_hes4_delta | +2.457 | +0.622 | +Inf |
| 52 | k_isc_i_hes5_delta | -0.080 | -2.804 | +Inf |
| 53 | k_isc_a | +3.572 | -Inf | +Inf |
| 54 | k_mcc_i_hes5_delta | +2.779 | -Inf | +Inf |
| 55 | k_mcc_a | +0.975 | -Inf | +1.057 |
| 56 | k_spdef_a_delta | +2.053 | +0.440 | +Inf |
| 57 | k_ssc_i_hes4_delta | +2.900 | +0.998 | +Inf |
| 58 | k_ssc_i_spdef_delta | -0.159 | -Inf | +1.863 |
| 59 | k_ssc_a | +0.045 | -0.336 | +0.281 |
| 60 | k_ssc_a_spdef_delta | +1.778 | +0.052 | +Inf |
| 61 | k_tp63_a_delta | +2.491 | +0.731 | +Inf |
| 62 | k_tp63_a_spdef_delta | +3.254 | +3.197 | +3.705 |

**Table 7: Confidence intervals for the estimated parameter values derived by the profile likelihood**
$\hat{\theta}$ indicates the estimated optimal parameter value. $\sigma^-$ and $\sigma^+$ indicate 95% point-wise confidence intervals.

| | name | $\hat{\theta}$ | $\sigma^-$ | $\sigma^+$ |
|---|---|---|---|---|
| 63 | p_lig_bsc | -0.532 | -Inf | -0.084 |
| 64 | p_lig_bc_factor | -1.153 | -Inf | -0.739 |
| 65 | p_lig_isc_factor | -0.000 | -0.624 | +Inf |
| 66 | p_lig_mcc_factor | -0.042 | -Inf | +Inf |
| 67 | p_lig_mpp_factor | +0.892 | +0.829 | +1.018 |
| 68 | p_lig_ssc_factor | -2.279 | -Inf | -1.131 |
| 69 | p_mcc_late | -1.909 | -2.039 | -1.570 |
| 70 | p_mpp | -2.403 | -2.625 | +Inf |
| 71 | p_Spdef | -3.013 | -3.405 | +Inf |
| 72 | p_Lig | -0.207 | -0.398 | +Inf |
| 75 | s_hes4 | -0.714 | -Inf | -0.572 |
| 76 | s_hes5 | -0.494 | -0.873 | +0.616 |
| 77 | s_hes7 | +1.232 | +0.213 | +Inf |
| 78 | s_isc | +3.794 | -Inf | +3.963 |
| 79 | s_mcc | +4.829 | -Inf | +Inf |
| 80 | s_mpp_foxi1 | +0.024 | +0.008 | +0.122 |
| 81 | s_spdef | -2.717 | -2.839 | -2.447 |
| 82 | s_ssc | +2.842 | +2.437 | +3.022 |
| 83 | s_tp63 | +1.795 | +1.425 | +2.173 |
| 84 | sd_tpm_foxa1 | -0.476 | -0.589 | -0.336 |
| 85 | sd_tpm_foxi1 | -0.518 | -0.632 | -0.378 |
| 86 | sd_tpm_hes4 | -0.742 | -0.857 | -0.600 |
| 87 | sd_tpm_hes5 | -0.185 | -0.295 | -0.041 |
| 88 | sd_tpm_hes7 | -0.169 | -0.290 | -0.026 |
| 89 | sd_tpm_lig | -0.512 | -0.632 | -0.371 |
| 90 | sd_tpm_mcidas | -0.461 | -0.574 | -0.320 |
| 91 | sd_tpm_spdef | -0.381 | -0.493 | -0.232 |
| 92 | sd_tpm_tp63 | -0.522 | -0.635 | -0.380 |
| 93 | v_max | -0.810 | -1.261 | -0.579 |
| 94 | v_max_bc | -1.886 | -2.249 | -1.443 |

**Table 8: Confidence intervals for the estimated parameter values derived by the profile likelihood**
$\hat{\theta}$ indicates the estimated optimal parameter value. $\sigma^-$ and $\sigma^+$ indicate 95% point-wise confidence intervals.
