## Supplementary material for "Temporal Notch signaling and Hes-mediated competitive de-repression regulate mucociliary cell fates in *Xenopus*": Mathematical Model 2.1

### Data2Dynamics Software – Modeling Report

29-Mar-2024

This is a curated version of the automatically generated modelling report in the *Data2Dynamics (D2D)* modelling framework in order to provide data and equations comprehensively. The code of D2D is available at `https://github.com/Data2Dynamics/d2d`

**Key references:**

- Data2Dynamics: a modeling environment tailored to parameter estimation in dynamical systems. A. Raue, B. Steiert, M. Schelker, C. Kreutz, T. Maiwald, H. Hass, J. Vanlier, C. Tönsing, L. Adlung, R. Engesser, W. Mader, T. Heinemann, J. Hasenauer, M. Schilling, T. Höfer, E. Klipp, F. Theis, U. Klingmüller, B. Schoeberl and J. Timmer. *Bioinformatics*, **31**(21), 3558-3560, 2015.
- Lessons learned from quantitative dynamical modeling in systems biology. A. Raue, M. Schilling, J. Bachmann, A. Matteson, M. Schelker, D. Kaschek, S. Hug, C. Kreutz, BD. Harms, F. Theis, U. Klingmüller, and J. Timmer. *PLOS ONE*, **8**(9), e74335, 2013.

**2 Estimated model parameters** **84**

### 1 Model version 2.1

#### 1.1 Comments

Model establised for describing Notch-induced differentiation and cell type composition in mucociliary tissue regulated via $hes$ genes.

#### 1.2 Dynamic variables

The model contains 27 dynamic variables. The dynamics of those variables evolve according to a system of ordinary differential equations (ODE) as will be defined in the following. The following list indicates the unique variable names and their initial conditions, i.e. their values at time point zero. In order to account for the delay between the transcription of the $\mathrm{hes}$ transcription factors and their transcriptional effects, the so-called linear chain trick has been applied, i.e. a sequence of five states $\mathrm{hes} \rightarrow \mathrm{hes\_d1} \rightarrow \mathrm{hes\_d2} \rightarrow \mathrm{hes\_d3} \rightarrow \mathrm{hes\_d}$ was introduced. Within this chain, production of $\mathrm{hes}$ denotes transcription and the transcriptional effect is mediated by the delayed increase of state $\mathrm{hes\_d}$.

- **Dynamic variable 1:** mpp
$$[\mathrm{mpp}](\mathrm{t}=0) = \mathrm{init\_mpp} \quad (1)$$
- **Dynamic variable 2:** ep
$$[\mathrm{ep}](\mathrm{t}=0) = \mathrm{init\_ep} \quad (2)$$
- **Dynamic variable 3:** bc
$$[\mathrm{bc}](\mathrm{t}=0) = \mathrm{init\_bc} \quad (3)$$
- **Dynamic variable 4:** isc
$$[\mathrm{isc}](\mathrm{t}=0) = \mathrm{init\_isc} \quad (4)$$
- **Dynamic variable 5:** mcc
$$[\mathrm{mcc}](\mathrm{t}=0) = \mathrm{init\_mcc} \quad (5)$$
- **Dynamic variable 6:** mcc_late
$$[\mathrm{mcc\_late}](\mathrm{t}=0) = \mathrm{init\_mcc\_late} \quad (6)$$
- **Dynamic variable 7:** ssc
$$[\mathrm{ssc}](\mathrm{t}=0) = \mathrm{init\_ssc} \quad (7)$$
- **Dynamic variable 8:** lig
$$[\mathrm{lig}](\mathrm{t}=0) = \mathrm{init\_lig} \quad (8)$$
- **Dynamic variable 9:** Lig
$$[\mathrm{Lig}](\mathrm{t}=0) = \mathrm{init\_Lig} \quad (9)$$
- **Dynamic variable 10:** hes4
$$[\mathrm{hes4}](\mathrm{t}=0) = \mathrm{init\_hes4} \quad (10)$$

- **Dynamic variable 11:** hes5

$$[\mathrm{hes5}](t=0) = \mathrm{init\_hes5} \tag{11}$$

- **Dynamic variable 12:** hes7

$$[\mathrm{hes7}](t=0) = \mathrm{init\_hes7} \tag{12}$$

- **Dynamic variable 13:** hes4_d1

$$[\mathrm{hes4\_d1}](t=0) = \mathrm{init\_hes4\_d1} \tag{13}$$

- **Dynamic variable 14:** hes5_d1

$$[\mathrm{hes5\_d1}](t=0) = \mathrm{init\_hes5\_d1} \tag{14}$$

- **Dynamic variable 15:** hes7_d1

$$[\mathrm{hes7\_d1}](t=0) = \mathrm{init\_hes7\_d1} \tag{15}$$

- **Dynamic variable 16:** hes4_d2

$$[\mathrm{hes4\_d2}](t=0) = \mathrm{init\_hes4\_d2} \tag{16}$$

- **Dynamic variable 17:** hes5_d2

$$[\mathrm{hes5\_d2}](t=0) = \mathrm{init\_hes5\_d2} \tag{17}$$

- **Dynamic variable 18:** hes7_d2

$$[\mathrm{hes7\_d2}](t=0) = \mathrm{init\_hes7\_d2} \tag{18}$$

- **Dynamic variable 19:** hes4_d3

$$[\mathrm{hes4\_d3}](t=0) = \mathrm{init\_hes4\_d3} \tag{19}$$

- **Dynamic variable 20:** hes5_d3

$$[\mathrm{hes5\_d3}](t=0) = \mathrm{init\_hes5\_d3} \tag{20}$$

- **Dynamic variable 21:** hes7_d3

$$[\mathrm{hes7\_d3}](t=0) = \mathrm{init\_hes7\_d3} \tag{21}$$

- **Dynamic variable 22:** hes4_d

$$[\mathrm{hes4\_d}](t=0) = \mathrm{init\_hes4\_d} \tag{22}$$

- **Dynamic variable 23:** hes5_d

$$[\mathrm{hes5\_d}](t=0) = \mathrm{init\_hes5\_d} \tag{23}$$

- **Dynamic variable 24:** hes7_d

$$[\mathrm{hes7\_d}](t=0) = \mathrm{init\_hes7\_d} \tag{24}$$

- **Dynamic variable 25:** tp63

$$[\mathrm{tp63}](t=0) = \mathrm{init\_tp63} \tag{25}$$

- **Dynamic variable 26:** spdef

$$[\mathrm{spdef}](t=0) = \mathrm{init\_spdef} \tag{26}$$

- **Dynamic variable 27:** Spdef

$$[\mathrm{Spdef}](t=0) = \mathrm{init\_Spdef} \tag{27}$$

#### 1.3 Reactions

Altogether, the model contains 37 reactions. Reactions define interactions between dynamics variables and were translated into the ODE systems by the rate-equation approach. The following list indicates the reactions and their corresponding reaction rate equations. In the reaction rate equations, dynamic and input variables are indicated by square brackets. The remaining variables are model parameters that were estimated from the data and remain constant over time.

- **Reaction 1:**

$$\mathrm{ep} \longrightarrow \mathrm{mpp}$$

$$v_1 = [\mathrm{ep}] \cdot \mathrm{p\_mpp} \tag{28}$$

- **Reaction 2:**

$$\mathrm{mpp} \longrightarrow \mathrm{isc}$$

$$v_2 = \frac{[\mathrm{hes7\_d}]^{\mathrm{h\_isc\_a}} \cdot \mathrm{k\_cell\_i}^{\mathrm{h\_cell\_i}} \cdot \mathrm{k\_isc\_i\_hes4}^{\mathrm{h\_isc\_i\_hes4}} \cdot \mathrm{k\_isc\_i\_hes5}^{\mathrm{h\_isc\_i\_hes5}} \cdot [\mathrm{mpp}] \cdot \mathrm{v\_max\_isc}}{\left([\mathrm{hes7\_d}]^{\mathrm{h\_isc\_a}} + \mathrm{k\_isc\_a}^{\mathrm{h\_isc\_a}}\right) \cdot \left([\mathrm{hes4\_d}]^{\mathrm{h\_isc\_i\_hes4}} + \mathrm{k\_isc\_i\_hes4}^{\mathrm{h\_isc\_i\_hes4}}\right) \cdot \left(\mathrm{k\_cell\_i}^{\mathrm{h\_cell\_i}} + [\mathrm{tp63}]^{\mathrm{h\_cell\_i}}\right) \cdot \left([\mathrm{hes5\_d}]^{\mathrm{h\_isc\_i\_hes5}} + \mathrm{k\_isc\_i\_hes5}^{\mathrm{h\_isc\_i\_hes5}}\right)} \tag{29}$$

- **Reaction 3:**

$$\mathrm{mpp} \longrightarrow \mathrm{mcc}$$

$$v_3 = \frac{[\mathrm{hes4\_d}]^{\mathrm{h\_mcc\_a}} \cdot \mathrm{k\_cell\_i}^{\mathrm{h\_cell\_i}} \cdot \mathrm{k\_mcc\_i\_hes5}^{\mathrm{h\_mcc\_i\_hes5}} \cdot [\mathrm{mpp}] \cdot \mathrm{v\_max\_mcc}}{\left([\mathrm{hes4\_d}]^{\mathrm{h\_mcc\_a}} + \mathrm{k\_mcc\_a}^{\mathrm{h\_mcc\_a}}\right) \cdot \left(\mathrm{k\_cell\_i}^{\mathrm{h\_cell\_i}} + [\mathrm{tp63}]^{\mathrm{h\_cell\_i}}\right) \cdot \left([\mathrm{hes5\_d}]^{\mathrm{h\_mcc\_i\_hes5}} + \mathrm{k\_mcc\_i\_hes5}^{\mathrm{h\_mcc\_i\_hes5}}\right)} \tag{30}$$

- **Reaction 4:**

$$\mathrm{mpp} \longrightarrow \mathrm{ssc}$$

$$v_4 = \frac{[\mathrm{Spdef}]^{\mathrm{h\_ssc\_a\_spdef}} \cdot [\mathrm{hes5\_d}]^{\mathrm{h\_ssc\_a}} \cdot \mathrm{k\_cell\_i}^{\mathrm{h\_cell\_i}} \cdot \mathrm{k\_ssc\_i\_hes4}^{\mathrm{h\_ssc\_i\_hes4}} \cdot \mathrm{k\_ssc\_i\_spdef}^{\mathrm{h\_ssc\_i\_spdef}} \cdot [\mathrm{mpp}] \cdot \mathrm{v\_max\_ssc}}{\left([\mathrm{Spdef}]^{\mathrm{h\_ssc\_a\_spdef}} + \mathrm{k\_ssc\_a\_spdef}^{\mathrm{h\_ssc\_a\_spdef}}\right) \cdot \left([\mathrm{hes5\_d}]^{\mathrm{h\_ssc\_a}} + \mathrm{k\_ssc\_a}^{\mathrm{h\_ssc\_a}}\right) \cdot \left([\mathrm{Spdef}]^{\mathrm{h\_ssc\_i\_spdef}} + \mathrm{k\_ssc\_i\_spdef}^{\mathrm{h\_ssc\_i\_spdef}}\right) \cdot \left(\mathrm{k\_cell\_i}^{\mathrm{h\_cell\_i}} + [\mathrm{tp63}]^{\mathrm{h\_cell\_i}}\right) \cdot A}$$

with $$A = \left([\mathrm{hes4\_d}]^{\mathrm{h\_ssc\_i\_hes4}} + \mathrm{k\_ssc\_i\_hes4}^{\mathrm{h\_ssc\_i\_hes4}}\right) \tag{31}$$

- **Reaction 5:**

$$\mathrm{mpp} \longrightarrow \mathrm{bc}$$

$$v_5 = [\mathrm{mpp}] \cdot \mathrm{v\_max\_bc} \cdot \left(\frac{[\mathrm{Lig}]^{\mathrm{h\_tp63\_a}}}{[\mathrm{Lig}]^{\mathrm{h\_tp63\_a}} + \mathrm{k\_tp63\_a}^{\mathrm{h\_tp63\_a}}} + \frac{[\mathrm{Spdef}]^{\mathrm{h\_tp63\_a\_spdef}}}{[\mathrm{Spdef}]^{\mathrm{h\_tp63\_a\_spdef}} + \mathrm{k\_tp63\_a\_spdef}^{\mathrm{h\_tp63\_a\_spdef}}}\right) \tag{32}$$

- **Reaction 6:**

$$\mathrm{mcc} \longrightarrow \mathrm{mcc\_late}$$

$$v_6 = [\mathrm{mcc}] \cdot \mathrm{p\_mcc\_late} \tag{33}$$

- **Reaction 7:**

$$\mathrm{lig} \longrightarrow \varnothing$$

$$v_7 = \mathrm{d\_lig} \cdot [\mathrm{lig}] \tag{34}$$

- **Reaction 8:**

$$\varnothing \longrightarrow \mathrm{lig}$$

$$v_8 = \frac{[\mathrm{mpp}] \cdot \mathrm{prior\_p\_lig\_mpp} \cdot \mathrm{reg\_p\_lig\_mpp\_fc} + [\mathrm{bc}] \cdot \mathrm{p\_lig\_bc} \cdot \mathrm{reg\_lig\_fc} \cdot \mathrm{s\_RNA} + [\mathrm{ep}] \cdot \mathrm{p\_lig\_ep} \cdot \mathrm{reg\_lig\_fc} \cdot \mathrm{s\_RNA} + [\mathrm{isc}] \cdot \mathrm{p\_lig\_isc} \cdot \mathrm{reg\_lig\_fc} \cdot \mathrm{s\_RNA} + A}{\mathrm{reg\_lig\_fc} \cdot \mathrm{s\_RNA} \cdot ([\mathrm{bc}] + [\mathrm{ep}] + [\mathrm{isc}] + [\mathrm{mcc}] + [\mathrm{mcc\_late}] + [\mathrm{mpp}] + [\mathrm{ssc}])}$$
$$\text{with } A = [\mathrm{mcc}] \cdot \mathrm{p\_lig\_mcc} \cdot \mathrm{reg\_lig\_fc} \cdot \mathrm{s\_RNA} + [\mathrm{mcc\_late}] \cdot \mathrm{p\_lig\_mcc} \cdot \mathrm{reg\_lig\_fc} \cdot \mathrm{s\_RNA} + \mathrm{p\_lig\_ssc} \cdot \mathrm{reg\_lig\_fc} \cdot \mathrm{s\_RNA} \cdot [\mathrm{ssc}] \tag{35}$$

- **Reaction 9:**

$$\varnothing \longrightarrow \mathrm{Lig}$$

$$v_9 = [\mathrm{lig}] \cdot \mathrm{p\_Lig} \tag{36}$$

- **Reaction 10:**

$$\mathrm{Lig} \longrightarrow \varnothing$$

$$v_{10} = [\mathrm{Lig}] \cdot \mathrm{d\_lig\_prot} \tag{37}$$

- **Reaction 11:**

$$\mathrm{hes4} \longrightarrow \varnothing$$

$$v_{11} = \mathrm{d\_hes4} \cdot [\mathrm{hes4}] \tag{38}$$

- **Reaction 12:**

$$\text{hes5} \longrightarrow \varnothing$$

$$v_{12} = \text{d\_hes5} \cdot [\text{hes5}] \tag{39}$$

- **Reaction 13:**

$$\text{hes7} \longrightarrow \varnothing$$

$$v_{13} = \text{d\_hes7} \cdot [\text{hes7}] \tag{40}$$

- **Reaction 14:**

$$\varnothing \longrightarrow \text{hes4}$$

$$v_{14} = \frac{[\text{Lig}]^{\text{h\_hes4\_a}} \cdot \text{k\_hes4\_i}^{\text{h\_hes4\_i}} \cdot \text{prior\_p\_hes4} \cdot \text{reg\_p\_hes4\_fc}}{\text{reg\_hes4\_fc} \cdot \text{s\_RNA} \cdot \left([\text{Lig}]^{\text{h\_hes4\_a}} + \text{k\_hes4\_a}^{\text{h\_hes4\_a}}\right) \cdot \left([\text{Lig}]^{\text{h\_hes4\_i}} + \text{k\_hes4\_i}^{\text{h\_hes4\_i}}\right)} \tag{41}$$

- **Reaction 15:**

$$\varnothing \longrightarrow \text{hes5}$$

$$v_{15} = \frac{[\text{Lig}]^{\text{h\_hes5\_a}} \cdot \text{prior\_p\_hes5} \cdot \text{reg\_p\_hes5\_fc}}{\text{reg\_hes5\_fc} \cdot \text{s\_RNA} \cdot \left([\text{Lig}]^{\text{h\_hes5\_a}} + \text{k\_hes5\_a}^{\text{h\_hes5\_a}}\right)} \tag{42}$$

- **Reaction 16:**

$$\varnothing \longrightarrow \text{hes7}$$

$$v_{16} = \frac{\text{k\_hes7\_i}^{\text{h\_hes7\_i}} \cdot \text{k\_hes7\_i\_spdef}^{\text{h\_hes7\_i\_spdef}} \cdot \text{prior\_p\_hes7} \cdot \text{reg\_p\_hes7\_fc}}{\text{reg\_hes7\_fc} \cdot \text{s\_RNA} \cdot \left([\text{Lig}]^{\text{h\_hes7\_i}} + \text{k\_hes7\_i}^{\text{h\_hes7\_i}}\right) \cdot \left([\text{Spdef}]^{\text{h\_hes7\_i\_spdef}} + \text{k\_hes7\_i\_spdef}^{\text{h\_hes7\_i\_spdef}}\right)} \tag{43}$$

- **Reaction 17:**

$$\varnothing \longrightarrow \text{hes4\_d1}$$

$$v_{17} = \frac{4 \cdot [\text{hes4}] \cdot \text{transl\_hes4\_lof\_fc}}{\text{hes\_delay}} \tag{44}$$

- **Reaction 18:**

$$\emptyset \longrightarrow \text{hes5\_d1}$$

$$v_{18} = \frac{4 \cdot [\text{hes5}] \cdot \text{transl\_hes5\_lof\_fc}}{\text{hes\_delay}} \tag{45}$$

- **Reaction 19:**

$$\emptyset \longrightarrow \text{hes7\_d1}$$

$$v_{19} = \frac{4 \cdot [\text{hes7}] \cdot \text{transl\_hes7\_lof\_fc}}{\text{hes\_delay}} \tag{46}$$

- **Reaction 20:**

$$\text{hes4\_d1} \longrightarrow \text{hes4\_d2}$$

$$v_{20} = \frac{4 \cdot [\text{hes4\_d1}]}{\text{hes\_delay}} \tag{47}$$

- **Reaction 21:**

$$\text{hes5\_d1} \longrightarrow \text{hes5\_d2}$$

$$v_{21} = \frac{4 \cdot [\text{hes5\_d1}]}{\text{hes\_delay}} \tag{48}$$

- **Reaction 22:**

$$\text{hes7\_d1} \longrightarrow \text{hes7\_d2}$$

$$v_{22} = \frac{4 \cdot [\text{hes7\_d1}]}{\text{hes\_delay}} \tag{49}$$

- **Reaction 23:**

$$\text{hes4\_d2} \longrightarrow \text{hes4\_d3}$$

$$v_{23} = \frac{4 \cdot [\text{hes4\_d2}]}{\text{hes\_delay}} \tag{50}$$

- **Reaction 24:**

$$\text{hes5\_d2} \longrightarrow \text{hes5\_d3}$$

$$v_{24} = \frac{4 \cdot [\text{hes5\_d2}]}{\text{hes\_delay}} \tag{51}$$

- **Reaction 25:**

$$\text{hes7\_d2} \longrightarrow \text{hes7\_d3}$$

$$v_{25} = \frac{4 \cdot [\text{hes7\_d2}]}{\text{hes\_delay}} \tag{52}$$

- **Reaction 26:**

$$\text{hes4\_d3} \longrightarrow \text{hes4\_d}$$

$$v_{26} = \frac{4 \cdot [\text{hes4\_d3}]}{\text{hes\_delay}} \tag{53}$$

- **Reaction 27:**

$$\text{hes5\_d3} \longrightarrow \text{hes5\_d}$$

$$v_{27} = \frac{4 \cdot [\text{hes5\_d3}]}{\text{hes\_delay}} \tag{54}$$

- **Reaction 28:**

$$\text{hes7\_d3} \longrightarrow \text{hes7\_d}$$

$$v_{28} = \frac{4 \cdot [\text{hes7\_d3}]}{\text{hes\_delay}} \tag{55}$$

- **Reaction 29:**

$$\text{hes4\_d} \longrightarrow \emptyset$$

$$v_{29} = \text{d\_hes4\_prot} \cdot [\text{hes4\_d}] \tag{56}$$

- **Reaction 30:**

$$\text{hes5\_d} \longrightarrow \varnothing$$

$$v_{30} = \text{d\_hes5\_prot} \cdot [\text{hes5\_d}] \tag{57}$$

- **Reaction 31:**

$$\text{hes7\_d} \longrightarrow \varnothing$$

$$v_{31} = \text{d\_hes7\_prot} \cdot [\text{hes7\_d}] \tag{58}$$

- **Reaction 32:**

$$\text{tp63} \longrightarrow \varnothing$$

$$v_{32} = \text{d\_tp63} \cdot [\text{tp63}] \tag{59}$$

- **Reaction 33:**

$$\varnothing \longrightarrow \text{tp63}$$

$$v_{33} = \frac{\text{prior\_p\_tp63} \cdot \text{reg\_p\_tp63\_fc} \cdot \left(\frac{[\text{Lig}]^{\text{h\_tp63\_a}}}{[\text{Lig}]^{\text{h\_tp63\_a}} + \text{k\_tp63\_a}^{\text{h\_tp63\_a}}} + \frac{[\text{Spdef}]^{\text{h\_tp63\_a\_spdef}}}{[\text{Spdef}]^{\text{h\_tp63\_a\_spdef}} + \text{k\_tp63\_a\_spdef}^{\text{h\_tp63\_a\_spdef}}}\right)}{\text{reg\_tp63\_fc} \cdot \text{s\_RNA}} \tag{60}$$

- **Reaction 34:**

$$\text{spdef} \longrightarrow \varnothing$$

$$v_{34} = \text{d\_spdef} \cdot [\text{spdef}] \tag{61}$$

- **Reaction 35:**

$$\varnothing \longrightarrow \text{spdef}$$

$$v_{35} = \frac{[\text{Lig}]^{\text{h\_spdef\_a}} \cdot \text{prior\_p\_spdef} \cdot \text{reg\_p\_spdef\_fc}}{\text{reg\_spdef\_fc} \cdot \text{s\_RNA} \cdot \left([\text{Lig}]^{\text{h\_spdef\_a}} + \text{k\_spdef\_a}^{\text{h\_spdef\_a}}\right)} \tag{62}$$

- **Reaction 36:**

$$\text{Spdef} \longrightarrow \varnothing$$

$$v_{36} = [\text{Spdef}] \cdot \text{d\_Spdef} \quad (63)$$

- **Reaction 37:**

$$\varnothing \longrightarrow \text{Spdef}$$

$$v_{37} = \text{p\_Spdef} \cdot [\text{spdef}] \quad (64)$$

#### 1.4 ODE system

The specified reation laws and rate equations $v$ determine an ODE system. The time evolution of the dynamical variables is calculated by solving this equation system.

$$\begin{aligned}
\text{d[mpp]/dt} &= +v_1 - v_2 - v_3 - v_4 - v_5 \\
\text{d[ep]/dt} &= -v_1 \\
\text{d[bc]/dt} &= +v_5 \\
\text{d[isc]/dt} &= +v_2 \\
\text{d[mcc]/dt} &= +v_3 - v_6 \\
\text{d[mcc_late]/dt} &= +v_6 \\
\text{d[ssc]/dt} &= +v_4 \\
\text{d[lig]/dt} &= -v_7 + v_8 \\
\text{d[Lig]/dt} &= +v_9 - v_{10} \\
\text{d[hes4]/dt} &= -v_{11} + v_{14} \\
\text{d[hes5]/dt} &= -v_{12} + v_{15} \\
\text{d[hes7]/dt} &= -v_{13} + v_{16} \\
\text{d[hes4_d1]/dt} &= +v_{17} - v_{20} \\
\text{d[hes5_d1]/dt} &= +v_{18} - v_{21} \\
\text{d[hes7_d1]/dt} &= +v_{19} - v_{22} \\
\text{d[hes4_d2]/dt} &= +v_{20} - v_{23} \\
\text{d[hes5_d2]/dt} &= +v_{21} - v_{24} \\
\text{d[hes7_d2]/dt} &= +v_{22} - v_{25} \\
\text{d[hes4_d3]/dt} &= +v_{23} - v_{26} \\
\text{d[hes5_d3]/dt} &= +v_{24} - v_{27} \\
\text{d[hes7_d3]/dt} &= +v_{25} - v_{28} \\
\text{d[hes4_d]/dt} &= +v_{26} - v_{29} \\
\text{d[hes5_d]/dt} &= +v_{27} - v_{30} \\
\text{d[hes7_d]/dt} &= +v_{28} - v_{31} \\
\text{d[tp63]/dt} &= -v_{32} + v_{33}
\end{aligned}$$

$$\text{d[spdef]/dt} = -v_{34} + v_{35}$$
$$\text{d[Spdef]/dt} = -v_{36} + v_{37}$$

The ODE system was solved by a parallelized implementation of the CVODES algorithm [1]. It also supplies the parameter sensitivities utilized for parameter estimation.

#### 1.5 Derived variables

The model contains 15 derived variables. Derived variables are calculated after the ODE system was solved. Dynamic and input variables are indicated by square brackets. The remaining variables are model parameters that remain constant over time.

- **Derived variable 1:** bsc

$$[\text{bsc}](\text{t}) = [\text{bc}] + [\text{ep}] \tag{65}$$

- **Derived variable 2:** mcc_total

$$[\text{mcc\_total}](\text{t}) = [\text{mcc}] + [\text{mcc\_late}] \tag{66}$$

- **Derived variable 3:** n_cells

$$[\text{n\_cells}](\text{t}) = [\text{bc}] + [\text{ep}] + [\text{isc}] + [\text{mcc}] + [\text{mcc\_late}] + [\text{mpp}] + [\text{ssc}] \tag{67}$$

- **Derived variable 4:** p_isc

$$[\text{p\_isc}](\text{t}) = \frac{[\text{hes7\_d}]^{\text{h\_isc\_a}} \cdot \text{k\_cell\_i}^{\text{h\_cell\_i}} \cdot \text{k\_isc\_i\_hes4}^{\text{h\_isc\_i\_hes4}} \cdot \text{k\_isc\_i\_hes5}^{\text{h\_isc\_i\_hes5}} \cdot \text{v\_max\_isc}}{\left([\text{hes7\_d}]^{\text{h\_isc\_a}} + \text{k\_isc\_a}^{\text{h\_isc\_a}}\right) \cdot \left([\text{hes4\_d}]^{\text{h\_isc\_i\_hes4}} + \text{k\_isc\_i\_hes4}^{\text{h\_isc\_i\_hes4}}\right) \cdot \left(\text{k\_cell\_i}^{\text{h\_cell\_i}} + [\text{tp63}]^{\text{h\_cell\_i}}\right) \cdot \left([\text{hes5\_d}]^{\text{h\_isc\_i\_hes5}} + \text{k\_isc\_i\_hes5}^{\text{h\_isc\_i\_hes5}}\right)} \tag{68}$$

- **Derived variable 5:** p_mcc

$$[\text{p\_mcc}](\text{t}) = \frac{[\text{hes4\_d}]^{\text{h\_mcc\_a}} \cdot \text{k\_cell\_i}^{\text{h\_cell\_i}} \cdot \text{k\_mcc\_i\_hes5}^{\text{h\_mcc\_i\_hes5}} \cdot \text{v\_max\_mcc}}{\left([\text{hes4\_d}]^{\text{h\_mcc\_a}} + \text{k\_mcc\_a}^{\text{h\_mcc\_a}}\right) \cdot \left(\text{k\_cell\_i}^{\text{h\_cell\_i}} + [\text{tp63}]^{\text{h\_cell\_i}}\right) \cdot \left([\text{hes5\_d}]^{\text{h\_mcc\_i\_hes5}} + \text{k\_mcc\_i\_hes5}^{\text{h\_mcc\_i\_hes5}}\right)} \tag{69}$$

- **Derived variable 6:** p_ssc

$$[\text{p\_ssc}](\text{t}) = \frac{[\text{Spdef}]^{\text{h\_ssc\_a\_spdef}} \cdot [\text{hes5\_d}]^{\text{h\_ssc\_a}} \cdot \text{k\_cell\_i}^{\text{h\_cell\_i}} \cdot \text{k\_ssc\_i\_hes4}^{\text{h\_ssc\_i\_hes4}} \cdot \text{k\_ssc\_i\_spdef}^{\text{h\_ssc\_i\_spdef}} \cdot \text{v\_max\_ssc}}{\left([\text{Spdef}]^{\text{h\_ssc\_a\_spdef}} + \text{k\_ssc\_a\_spdef}^{\text{h\_ssc\_a\_spdef}}\right) \cdot \left([\text{hes5\_d}]^{\text{h\_ssc\_a}} + \text{k\_ssc\_a}^{\text{h\_ssc\_a}}\right) \cdot \left([\text{Spdef}]^{\text{h\_ssc\_i\_spdef}} + \text{k\_ssc\_i\_spdef}^{\text{h\_ssc\_i\_spdef}}\right) \cdot \left(\text{k\_cell\_i}^{\text{h\_cell\_i}} + [\text{tp63}]^{\text{h\_cell\_i}}\right) \cdot A}$$
$$\text{with } A = \left([\text{hes4\_d}]^{\text{h\_ssc\_i\_hes4}} + \text{k\_ssc\_i\_hes4}^{\text{h\_ssc\_i\_hes4}}\right) \tag{70}$$

- **Derived variable 7:** p_bc

$$[\text{p\_bc}](\text{t}) = \text{v\_max\_bc} \cdot \left(\frac{[\text{Lig}]^{\text{h\_tp63\_a}}}{[\text{Lig}]^{\text{h\_tp63\_a}} + \text{k\_tp63\_a}^{\text{h\_tp63\_a}}} + \frac{[\text{Spdef}]^{\text{h\_tp63\_a\_spdef}}}{[\text{Spdef}]^{\text{h\_tp63\_a\_spdef}} + \text{k\_tp63\_a\_spdef}^{\text{h\_tp63\_a\_spdef}}}\right) \tag{71}$$

- **Derived variable 8:** p_lig

$$[\text{p\_lig}](\text{t}) =$$
$$\frac{[\text{mpp}] \cdot \text{prior\_p\_lig\_mpp} \cdot \text{reg\_p\_lig\_mpp\_fc} + [\text{bc}] \cdot \text{p\_lig\_bc} \cdot \text{reg\_lig\_fc} \cdot \text{s\_RNA} + [\text{ep}] \cdot \text{p\_lig\_ep} \cdot \text{reg\_lig\_fc} \cdot \text{s\_RNA} + [\text{isc}] \cdot \text{p\_lig\_isc} \cdot \text{reg\_lig\_fc} \cdot \text{s\_RNA} + [\text{mcc}] \cdot \text{p\_lig\_mcc} \cdot \text{reg\_lig\_fc} \cdot \text{s\_RNA} + A}{\text{reg\_lig\_fc} \cdot \text{s\_RNA} \cdot ([\text{bc}] + [\text{ep}] + [\text{isc}] + [\text{mcc}] + [\text{mcc\_late}] + [\text{mpp}] + [\text{ssc}])}$$
$$\text{with } A = [\text{mcc\_late}] \cdot \text{p\_lig\_mcc} \cdot \text{reg\_lig\_fc} \cdot \text{s\_RNA} + \text{p\_lig\_ssc} \cdot \text{reg\_lig\_fc} \cdot \text{s\_RNA} \cdot [\text{ssc}] \tag{72}$$

- **Derived variable 9:** p_hes4

$$[\text{p\_hes4}](\text{t}) = \frac{[\text{Lig}]^{\text{h\_hes4\_a}} \cdot \text{k\_hes4\_i}^{\text{h\_hes4\_i}} \cdot \text{prior\_p\_hes4} \cdot \text{reg\_p\_hes4\_fc}}{\text{reg\_hes4\_fc} \cdot \text{s\_RNA} \cdot \left([\text{Lig}]^{\text{h\_hes4\_a}} + \text{k\_hes4\_a}^{\text{h\_hes4\_a}}\right) \cdot \left([\text{Lig}]^{\text{h\_hes4\_i}} + \text{k\_hes4\_i}^{\text{h\_hes4\_i}}\right)} \tag{73}$$

- **Derived variable 10:** p_hes5

$$[\text{p\_hes5}](\text{t}) = \frac{[\text{Lig}]^{\text{h\_hes5\_a}} \cdot \text{prior\_p\_hes5} \cdot \text{reg\_p\_hes5\_fc}}{\text{reg\_hes5\_fc} \cdot \text{s\_RNA} \cdot \left([\text{Lig}]^{\text{h\_hes5\_a}} + \text{k\_hes5\_a}^{\text{h\_hes5\_a}}\right)} \tag{74}$$

- **Derived variable 11:** p_hes7

$$[\text{p\_hes7}](\text{t}) = \frac{\text{k\_hes7\_i}^{\text{h\_hes7\_i}} \cdot \text{k\_hes7\_i\_spdef}^{\text{h\_hes7\_i\_spdef}} \cdot \text{prior\_p\_hes7} \cdot \text{reg\_p\_hes7\_fc}}{\text{reg\_hes7\_fc} \cdot \text{s\_RNA} \cdot \left([\text{Lig}]^{\text{h\_hes7\_i}} + \text{k\_hes7\_i}^{\text{h\_hes7\_i}}\right) \cdot \left([\text{Spdef}]^{\text{h\_hes7\_i\_spdef}} + \text{k\_hes7\_i\_spdef}^{\text{h\_hes7\_i\_spdef}}\right)} \tag{75}$$

- **Derived variable 12:** p_spdef

$$[\text{p\_spdef}](\text{t}) = \frac{[\text{Lig}]^{\text{h\_spdef\_a}} \cdot \text{prior\_p\_spdef} \cdot \text{reg\_p\_spdef\_fc}}{\text{reg\_spdef\_fc} \cdot \text{s\_RNA} \cdot \left([\text{Lig}]^{\text{h\_spdef\_a}} + \text{k\_spdef\_a}^{\text{h\_spdef\_a}}\right)} \tag{76}$$

- **Derived variable 13:** p_tp63

$$[\text{p\_tp63}](\text{t}) = \frac{\text{prior\_p\_tp63} \cdot \text{reg\_p\_tp63\_fc} \cdot \left(\frac{[\text{Lig}]^{\text{h\_tp63\_a}}}{[\text{Lig}]^{\text{h\_tp63\_a}} + \text{k\_tp63\_a}^{\text{h\_tp63\_a}}} + \frac{[\text{Spdef}]^{\text{h\_tp63\_a\_spdef}}}{[\text{Spdef}]^{\text{h\_tp63\_a\_spdef}} + \text{k\_tp63\_a\_spdef}^{\text{h\_tp63\_a\_spdef}}}\right)}{\text{reg\_tp63\_fc} \cdot \text{s\_RNA}} \tag{77}$$

- **Derived variable 14:** tp63_hill

$$[\text{tp63\_hill}](\text{t}) = \frac{[\text{Lig}]^{\text{h\_tp63\_a}}}{[\text{Lig}]^{\text{h\_tp63\_a}} + \text{k\_tp63\_a}^{\text{h\_tp63\_a}}} + \frac{[\text{Spdef}]^{\text{h\_tp63\_a\_spdef}}}{[\text{Spdef}]^{\text{h\_tp63\_a\_spdef}} + \text{k\_tp63\_a\_spdef}^{\text{h\_tp63\_a\_spdef}}} \tag{78}$$

#### 1.6 Conditions

Conditions modify the model according to replacement rules. New model parameters can be introduced or relations between existing model parameters can be implemented. The following list are default conditions that can be replace my experiment specific conditions defined seperately for each data set.

In our project, we used replacements primarily to set dynamic variable to steady states. The steady states were derived by setting the right-hand side of the ODEs to zero and solve for the dynamic variables. In addition, some initial conditions which are not assumed in the steady state were replaced by the initial value of the corresponding observation to better control parameter bounds and initialization of parameter optimization.

$$\begin{aligned}
\text{d_hes4_prot} &\rightarrow \text{d_hes4_prot_fc} \cdot \text{d_hes_prot} \\
\text{d_hes4} &\rightarrow \text{d_hes4_fc} \cdot \text{d_hes_rna} \\
\text{d_hes5_prot} &\rightarrow \text{d_hes5_prot_fc} \cdot \text{d_hes_prot} \\
\text{d_hes5} &\rightarrow \text{d_hes5_fc} \cdot \text{d_hes_rna} \\
\text{d_hes7_prot} &\rightarrow \text{d_hes7_prot_fc} \cdot \text{d_hes_prot} \\
\text{d_hes7} &\rightarrow \text{d_hes7_fc} \cdot \text{d_hes_rna} \\
\text{init_bc} &\rightarrow 0 \\
\text{init_hes4} &\rightarrow \frac{\text{init_hes4} \cdot \text{init_hes4_gof_hc_fc} \cdot \text{init_hes4_gof_lc_fc}}{\text{reg_hes4_fc} \cdot \text{s_RNA}} \\
\text{init_hes4_d} &\rightarrow \frac{4 \cdot \text{init_hes4} \cdot \text{init_hes4_gof_hc_fc} \cdot \text{init_hes4_gof_lc_fc} \cdot \text{transl_hes4_lof_fc}}{\text{d_hes4_prot_fc} \cdot \text{d_hes_prot} \cdot \text{hes_delay} \cdot \text{reg_hes4_fc} \cdot \text{s_RNA}}
\end{aligned}$$

$$\text{init\_hes4\_d1} \rightarrow \frac{\text{init\_hes4} \cdot \text{init\_hes4\_gof\_hc\_fc} \cdot \text{init\_hes4\_gof\_lc\_fc} \cdot \text{transl\_hes4\_lof\_fc}}{\text{reg\_hes4\_fc} \cdot \text{s\_RNA}}$$

$$\text{init\_hes4\_d2} \rightarrow \frac{\text{init\_hes4} \cdot \text{init\_hes4\_gof\_hc\_fc} \cdot \text{init\_hes4\_gof\_lc\_fc} \cdot \text{transl\_hes4\_lof\_fc}}{\text{reg\_hes4\_fc} \cdot \text{s\_RNA}}$$

$$\text{init\_hes4\_d3} \rightarrow \frac{\text{init\_hes4} \cdot \text{init\_hes4\_gof\_hc\_fc} \cdot \text{init\_hes4\_gof\_lc\_fc} \cdot \text{transl\_hes4\_lof\_fc}}{\text{reg\_hes4\_fc} \cdot \text{s\_RNA}}$$

$$\text{init\_hes5} \rightarrow \frac{\text{init\_hes5} \cdot \text{init\_hes5\_gof\_hc\_fc} \cdot \text{init\_hes5\_gof\_lc\_fc}}{\text{reg\_hes5\_fc} \cdot \text{s\_RNA}}$$

$$\text{init\_hes5\_d} \rightarrow \frac{4 \cdot \text{init\_hes5} \cdot \text{init\_hes5\_gof\_hc\_fc} \cdot \text{init\_hes5\_gof\_lc\_fc} \cdot \text{transl\_hes5\_lof\_fc}}{\text{d\_hes5\_prot\_fc} \cdot \text{d\_hes\_prot} \cdot \text{hes\_delay} \cdot \text{reg\_hes5\_fc} \cdot \text{s\_RNA}}$$

$$\text{init\_hes5\_d1} \rightarrow \frac{\text{init\_hes5} \cdot \text{init\_hes5\_gof\_hc\_fc} \cdot \text{init\_hes5\_gof\_lc\_fc} \cdot \text{transl\_hes5\_lof\_fc}}{\text{reg\_hes5\_fc} \cdot \text{s\_RNA}}$$

$$\text{init\_hes5\_d2} \rightarrow \frac{\text{init\_hes5} \cdot \text{init\_hes5\_gof\_hc\_fc} \cdot \text{init\_hes5\_gof\_lc\_fc} \cdot \text{transl\_hes5\_lof\_fc}}{\text{reg\_hes5\_fc} \cdot \text{s\_RNA}}$$

$$\text{init\_hes5\_d3} \rightarrow \frac{\text{init\_hes5} \cdot \text{init\_hes5\_gof\_hc\_fc} \cdot \text{init\_hes5\_gof\_lc\_fc} \cdot \text{transl\_hes5\_lof\_fc}}{\text{reg\_hes5\_fc} \cdot \text{s\_RNA}}$$

$$\text{init\_hes7} \rightarrow \frac{\text{init\_hes7} \cdot \text{init\_hes7\_gof\_hc\_fc} \cdot \text{init\_hes7\_gof\_lc\_fc}}{\text{reg\_hes7\_fc} \cdot \text{s\_RNA}}$$

$$\text{init\_hes7\_d} \rightarrow \frac{4 \cdot \text{init\_hes7} \cdot \text{init\_hes7\_gof\_hc\_fc} \cdot \text{init\_hes7\_gof\_lc\_fc} \cdot \text{transl\_hes7\_lof\_fc}}{\text{d\_hes7\_prot\_fc} \cdot \text{d\_hes\_prot} \cdot \text{hes\_delay} \cdot \text{reg\_hes7\_fc} \cdot \text{s\_RNA}}$$

$$\text{init\_hes7\_d1} \rightarrow \frac{\text{init\_hes7} \cdot \text{init\_hes7\_gof\_hc\_fc} \cdot \text{init\_hes7\_gof\_lc\_fc} \cdot \text{transl\_hes7\_lof\_fc}}{\text{reg\_hes7\_fc} \cdot \text{s\_RNA}}$$

$$\text{init\_hes7\_d2} \rightarrow \frac{\text{init\_hes7} \cdot \text{init\_hes7\_gof\_hc\_fc} \cdot \text{init\_hes7\_gof\_lc\_fc} \cdot \text{transl\_hes7\_lof\_fc}}{\text{reg\_hes7\_fc} \cdot \text{s\_RNA}}$$

$$\text{init\_hes7\_d3} \rightarrow \frac{\text{init\_hes7} \cdot \text{init\_hes7\_gof\_hc\_fc} \cdot \text{init\_hes7\_gof\_lc\_fc} \cdot \text{transl\_hes7\_lof\_fc}}{\text{reg\_hes7\_fc} \cdot \text{s\_RNA}}$$

$$\text{init\_lig} \rightarrow \frac{\text{init\_ligand}}{\text{reg\_lig\_fc} \cdot \text{s\_RNA}}$$

$$\text{init\_mcc\_late} \rightarrow 0$$

$$\text{init\_Spdef} \rightarrow \frac{\text{init\_spdef} \cdot \text{p\_Spdef}}{\text{d\_Spdef} \cdot \text{reg\_spdef\_fc} \cdot \text{s\_RNA}}$$

$$\text{init\_spdef} \rightarrow \frac{\text{init\_spdef}}{\text{reg\_spdef\_fc} \cdot \text{s\_RNA}}$$

$$\text{init\_ssc} \rightarrow 0$$

$$\text{init\_Lig} \rightarrow \frac{\text{init\_ligand} \cdot \text{p\_Lig}}{\text{d\_lig\_prot} \cdot \text{reg\_lig\_fc} \cdot \text{s\_RNA}}$$

$$\text{init\_tp63} \rightarrow \frac{\text{init\_tp63}}{\text{reg\_tp63\_fc} \cdot \text{s\_RNA}}$$

$$\text{k\_cell\_i} \rightarrow \frac{\text{init\_tp63} \cdot \text{k\_cell\_i\_fc}}{\text{reg\_tp63\_fc} \cdot \text{s\_RNA}}$$

$$\text{k\_hes4\_i} \rightarrow \frac{\text{init\_ligand} \cdot \text{k\_hes4\_a\_fc} \cdot \text{k\_hes4\_i\_fc} \cdot \text{p\_Lig}}{\text{d\_lig\_prot} \cdot \text{reg\_lig\_fc} \cdot \text{s\_RNA}}$$

$$\text{k\_hes4\_a} \rightarrow \frac{\text{init\_ligand} \cdot \text{k\_hes4\_a\_fc} \cdot \text{p\_Lig}}{\text{d\_lig\_prot} \cdot \text{reg\_lig\_fc} \cdot \text{s\_RNA}}$$

$$\text{k\_hes5\_a} \rightarrow \frac{\text{init\_ligand} \cdot \text{k\_hes5\_a\_fc} \cdot \text{p\_Lig}}{\text{d\_lig\_prot} \cdot \text{reg\_lig\_fc} \cdot \text{s\_RNA}}$$

$$\text{k\_hes7\_i} \rightarrow \frac{\text{init\_ligand} \cdot \text{p\_Lig}}{\text{d\_lig\_prot} \cdot \text{reg\_lig\_fc} \cdot \text{s\_RNA}}$$

$$\text{k\_hes7\_i\_spdef} \rightarrow \frac{\text{init\_spdef} \cdot \text{k\_hes7\_i\_spdef\_fc} \cdot \text{p\_Spdef}}{\text{d\_Spdef} \cdot \text{reg\_spdef\_fc} \cdot \text{s\_RNA}}$$

$$\text{k\_isc\_i\_hes4} \rightarrow \frac{4 \cdot \text{init\_hes4} \cdot \text{k\_isc\_i\_hes4\_fc}}{\text{d\_hes4\_prot\_fc} \cdot \text{d\_hes\_prot} \cdot \text{hes\_delay} \cdot \text{reg\_hes4\_fc} \cdot \text{s\_RNA}}$$

$$\text{k\_isc\_i\_hes5} \rightarrow \frac{4 \cdot \text{init\_hes5} \cdot \text{k\_isc\_i\_hes5\_fc}}{\text{d\_hes5\_prot\_fc} \cdot \text{d\_hes\_prot} \cdot \text{hes\_delay} \cdot \text{reg\_hes5\_fc} \cdot \text{s\_RNA}}$$

$$\text{k\_isc\_a} \rightarrow \frac{4 \cdot \text{init\_hes7} \cdot \text{k\_isc\_a\_fc}}{\text{d\_hes7\_prot\_fc} \cdot \text{d\_hes\_prot} \cdot \text{hes\_delay} \cdot \text{reg\_hes7\_fc} \cdot \text{s\_RNA}}$$

$$\text{k\_mcc\_i\_hes5} \rightarrow \frac{4 \cdot \text{init\_hes5} \cdot \text{k\_mcc\_i\_hes5\_fc}}{\text{d\_hes5\_prot\_fc} \cdot \text{d\_hes\_prot} \cdot \text{hes\_delay} \cdot \text{reg\_hes5\_fc} \cdot \text{s\_RNA}}$$

$$\text{k\_mcc\_a} \rightarrow \frac{4 \cdot \text{init\_hes4} \cdot \text{k\_mcc\_a\_fc}}{\text{d\_hes4\_prot\_fc} \cdot \text{d\_hes\_prot} \cdot \text{hes\_delay} \cdot \text{reg\_hes4\_fc} \cdot \text{s\_RNA}}$$

$$\text{k\_spdef\_a} \rightarrow \frac{\text{init\_ligand} \cdot \text{k\_spdef\_a\_fc} \cdot \text{p\_Lig}}{\text{d\_lig\_prot} \cdot \text{reg\_lig\_fc} \cdot \text{s\_RNA}}$$

$$\text{k\_ssc\_i\_hes4} \rightarrow \frac{4 \cdot \text{init\_hes4} \cdot \text{k\_ssc\_i\_hes4\_fc}}{\text{d\_hes4\_prot\_fc} \cdot \text{d\_hes\_prot} \cdot \text{hes\_delay} \cdot \text{reg\_hes4\_fc} \cdot \text{s\_RNA}}$$

$$\text{k\_ssc\_i\_spdef} \rightarrow \frac{\text{init\_spdef} \cdot \text{k\_ssc\_a\_spdef\_fc} \cdot \text{k\_ssc\_i\_spdef\_fc} \cdot \text{p\_Spdef}}{\text{d\_Spdef} \cdot \text{reg\_spdef\_fc} \cdot \text{s\_RNA}}$$

$$\text{k\_ssc\_a} \rightarrow \frac{4 \cdot \text{init\_hes5} \cdot \text{k\_ssc\_a\_fc}}{\text{d\_hes5\_prot\_fc} \cdot \text{d\_hes\_prot} \cdot \text{hes\_delay} \cdot \text{reg\_hes5\_fc} \cdot \text{s\_RNA}}$$

$$\text{k\_ssc\_a\_spdef} \rightarrow \frac{\text{init\_spdef} \cdot \text{k\_ssc\_a\_spdef\_fc} \cdot \text{p\_Spdef}}{\text{d\_Spdef} \cdot \text{reg\_spdef\_fc} \cdot \text{s\_RNA}}$$

$$\text{k\_tp63\_a} \rightarrow \frac{\text{init\_ligand} \cdot \text{k\_tp63\_a\_fc} \cdot \text{p\_Lig}}{\text{d\_lig\_prot} \cdot \text{reg\_lig\_fc} \cdot \text{s\_RNA}}$$

$$\text{k\_tp63\_a\_spdef} \rightarrow \frac{\text{init\_spdef} \cdot \text{k\_tp63\_a\_spdef\_fc} \cdot \text{p\_Spdef}}{\text{d\_Spdef} \cdot \text{reg\_spdef\_fc} \cdot \text{s\_RNA}}$$

$$\text{p\_lig\_ep} \rightarrow \frac{\text{p\_lig\_ep\_fc} \cdot \text{prior\_p\_lig\_mpp} \cdot \text{reg\_p\_lig\_mpp\_fc}}{\text{reg\_lig\_fc} \cdot \text{s\_RNA}}$$

$$\text{p\_lig\_bc} \rightarrow \frac{\text{p\_lig\_bc\_fc} \cdot \text{prior\_p\_lig\_mpp} \cdot \text{reg\_p\_lig\_mpp\_fc}}{\text{reg\_lig\_fc} \cdot \text{s\_RNA}}$$

$$\text{p\_lig\_isc} \rightarrow \frac{\text{p\_lig\_isc\_fc} \cdot \text{prior\_p\_lig\_mpp} \cdot \text{reg\_p\_lig\_mpp\_fc}}{\text{reg\_lig\_fc} \cdot \text{s\_RNA}}$$

$$\text{p\_lig\_mcc} \rightarrow \frac{\text{p\_lig\_mcc\_fc} \cdot \text{prior\_p\_lig\_mpp} \cdot \text{reg\_p\_lig\_mpp\_fc}}{\text{reg\_lig\_fc} \cdot \text{s\_RNA}}$$

$$\text{p\_lig\_ssc} \rightarrow \frac{\text{p\_lig\_ssc\_fc} \cdot \text{prior\_p\_lig\_mpp} \cdot \text{reg\_p\_lig\_mpp\_fc}}{\text{reg\_lig\_fc} \cdot \text{s\_RNA}}$$

$$\text{v\_max\_isc} \rightarrow \text{v\_max\_cell} \cdot \text{vmax\_isc\_fc}$$

$$\text{v\_max\_mcc} \rightarrow \text{v\_max\_cell} \cdot \text{vmax\_mcc\_fc}$$

$$\text{v\_max\_ssc} \rightarrow \text{v\_max\_cell} \cdot \text{vmax\_ssc\_fc}$$

#### 1.7 Experiment: RNAseq_data_WT_controls

##### 1.7.1 Comments

Time-resolved RNAseq data for the most important molecular players in the mdoelled mucociliary tissue.

##### 1.7.2 Observables

The following observables are modified in this data set.

- **Observable:** ubp1

$$\mathrm{ubp1(t)} = \log_{10}(\mathrm{background\_ubp1} + \frac{[\mathrm{isc}] \cdot \mathrm{s\_isc\_pure}}{[\mathrm{n\_cells}]}) \tag{79}$$

$$\sigma\{\mathrm{ubp1}\}(\mathrm{t}) = \mathrm{sd\_RNA} \tag{80}$$

- **Observable:** foxi1

$$\mathrm{foxi1(t)} = \log_{10}(\mathrm{s\_isc} \cdot \left(\frac{[\mathrm{isc}]}{[\mathrm{n\_cells}]} + \frac{[\mathrm{mpp}] \cdot \mathrm{s\_mpp\_foxi1\_fc}}{[\mathrm{n\_cells}]}\right)) \tag{81}$$

$$\sigma\{\mathrm{foxi1}\}(\mathrm{t}) = \mathrm{sd\_RNA} \tag{82}$$

- **Observable:** mcidas

$$\mathrm{mcidas(t)} = \log_{10}(\mathrm{background\_mcidas} + \frac{[\mathrm{mcc}] \cdot \mathrm{s\_mcc}}{[\mathrm{n\_cells}]}) \tag{83}$$

$$\sigma\{\mathrm{mcidas}\}(\mathrm{t}) = \mathrm{sd\_RNA} \tag{84}$$

- **Observable:** foxa1

$$\mathrm{foxa1(t)} = \log_{10}(\mathrm{background\_foxa1} + \frac{\mathrm{s\_ssc} \cdot [\mathrm{ssc}]}{[\mathrm{n\_cells}]}) \tag{85}$$

$$\sigma\{\mathrm{foxa1}\}(\mathrm{t}) = \mathrm{sd\_RNA} \tag{86}$$

- **Observable:** lig

$$[\mathrm{lig}](\mathrm{t}) = \log_{10}([\mathrm{lig}] \cdot \mathrm{reg\_lig\_fc} \cdot \mathrm{s\_RNA}) \tag{87}$$

$$\sigma\{[\mathrm{lig}]\}(\mathrm{t}) = \mathrm{sd\_RNA} \tag{88}$$

- **Observable:** hes4

$$[\mathrm{hes4}](\mathrm{t}) = \log_{10}([\mathrm{hes4}] \cdot \mathrm{reg\_hes4\_fc} \cdot \mathrm{s\_RNA}) \tag{89}$$

$$\sigma\{[\mathrm{hes4}]\}(\mathrm{t}) = \mathrm{sd\_RNA} \tag{90}$$

- **Observable:** hes5

$$[\mathrm{hes5}](\mathrm{t}) = \log_{10}([\mathrm{hes5}] \cdot \mathrm{reg\_hes5\_fc} \cdot \mathrm{s\_RNA}) \tag{91}$$

$$\sigma\{[\mathrm{hes5}]\}(\mathrm{t}) = \mathrm{sd\_RNA} \tag{92}$$

- **Observable:** hes7

$$[\mathrm{hes7}](\mathrm{t}) = \log_{10}([\mathrm{hes7}] \cdot \mathrm{reg\_hes7\_fc} \cdot \mathrm{s\_RNA}) \tag{93}$$

$$\sigma\{[\mathrm{hes7}]\}(\mathrm{t}) = \mathrm{sd\_RNA} \tag{94}$$

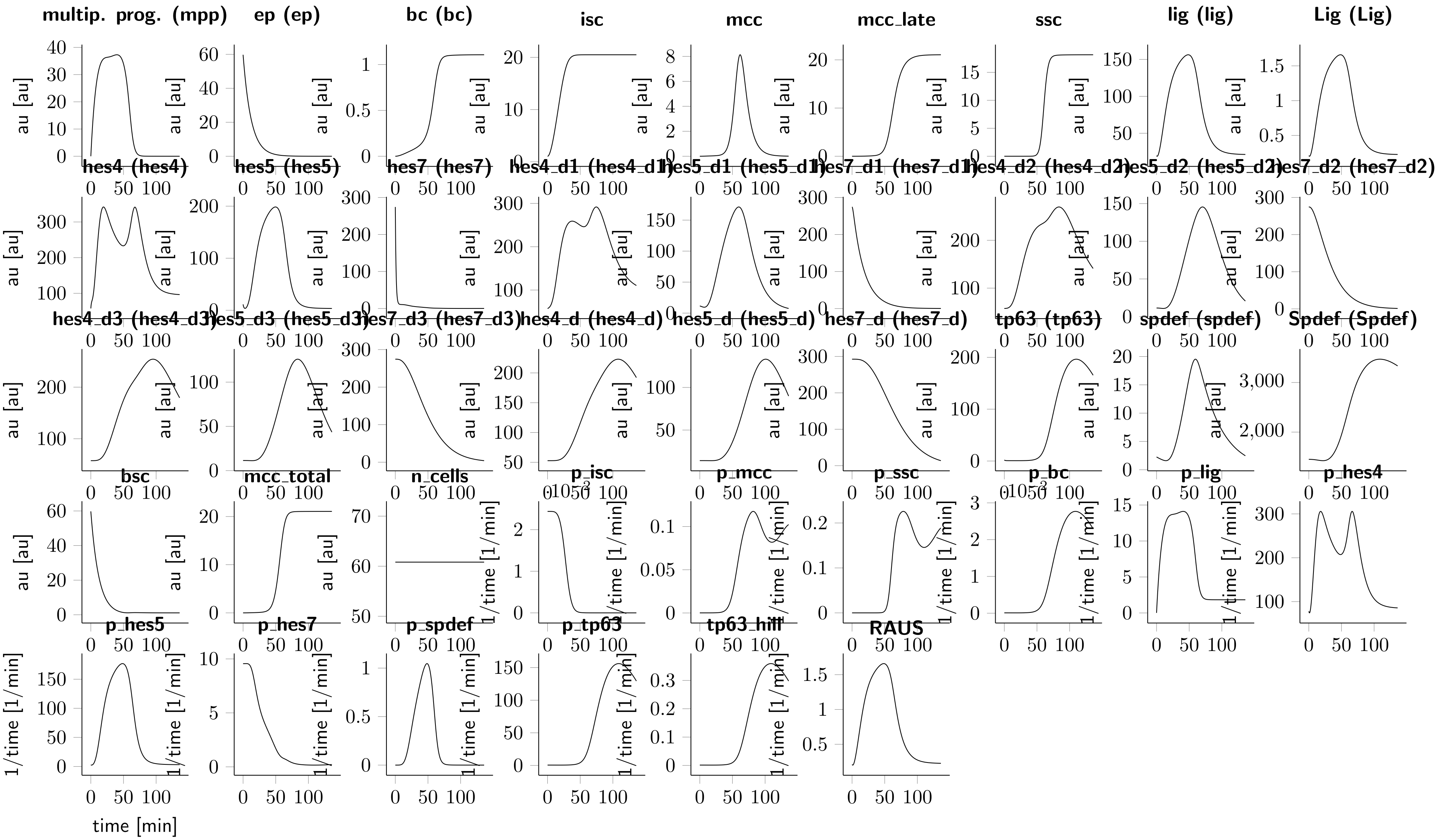

**Figure 1:** Fitted dynamics for the wildtype condition.

| time [min] | ubp1 [tpm] | foxi1 [tpm] | mcidas [tpm] | foxa1 [tpm] | lig [tpm] | hes4 [tpm] | hes5 [tpm] | hes7 [tpm] | tp63 [tpm] | spdef [tpm] |
|---|---|---|---|---|---|---|---|---|---|---|
| 0.07605 | 2.85433 | 21.4122 | 5.63667 | 4.01337 | 19.3056 | 58.2492 | 8.03 | 228.636 | 5.21332 | 0.290805 |
| 0 | 2.73866 | 3.73601 | 2.72262 | 6.49646 | 20.66 | 117.69 | 13.0757 | 308.907 | 0.688885 | 4.33647 |
| 13.8944 | 8.295 | 138.971 | 3.067 | 1.54264 | 77.022 | 385.925 | 119.701 | 26.119 | 0.776347 | 3.20894 |
| 13.2548 | 11.2875 | 125.131 | 4.70885 | 2.00375 | 87.2581 | 354.278 | 95.7696 | 12.3979 | 0.59586 | 3.28351 |
| 10.7815 | 4.63822 | 15.7956 | 1.12522 | 1.57711 | 42.0192 | 587.846 | 136.407 | 75.7261 | 0.8056 | 2.69489 |
| 10.8634 | 12.879 | 242.094 | 8.74825 | 1.5337 | 100.201 | 169.706 | 6.45534 | 0.55149 | 0.641397 | 6.06226 |
| 13.8476 | 16.0414 | 127.625 | 4.70015 | 2.05203 | 63.4676 | 431.103 | 49.7569 | 8.41104 | 0.804862 | 5.66696 |
| 17.3623 | 6.78132 | 94.1405 | 8.29909 | 2.08001 | 54.0713 | 455.313 | 32.8595 | 5.49451 | 0.760972 | 1.50088 |
| 14.9708 | 10.6195 | 77.2064 | 0.879445 | 1.60553 | 27.4377 | 321.448 | 39.3183 | 21.3358 | 0.259333 | 0.686017 |
| 27.7525 | 15.8445 | 172.065 | 9.72742 | 1.1865 | 93.9638 | 238.221 | 158.562 | 5.81861 | 0.820473 | 4.31496 |
| 27.55 | 24.2631 | 224.839 | 25.6547 | 1.25839 | 123.09 | 297.602 | 135.976 | 3.79494 | 0.783333 | 4.7794 |
| 42.7568 | 46.0573 | 453.483 | 145.236 | 1.40541 | 180.745 | 291.355 | 174.592 | 7.44881 | 1.56658 | 11.6615 |
| 41.186 | 26.4713 | 304.528 | 85.8069 | 1.41914 | 146.701 | 171.716 | 276.509 | 2.0362 | 1.38043 | 7.8565 |
| 74.6147 | 115.37 | 271.455 | 494.72 | 64.7823 | 115.075 | 319.437 | 109.47 | 12.0144 | 115.268 | 15.3678 |
| 77.8776 | 130.039 | 321.149 | 302.605 | 25.9136 | 72.1299 | 157.068 | 5.03945 | 1.05558 | 134.161 | 17.6462 |
| 75.6001 | 123.196 | 239.25 | 295.523 | 67.2722 | 237.103 | 271.849 | 117.662 | 0.232522 | 92.876 | 16.6473 |
| 78.6617 | 148.113 | 229.908 | 224.015 | 5.97821 | 25.2432 | 244.478 | 9.11634 | 0.22395 | 33.1506 | 6.4619 |
| 79.4141 | 237.712 | 343.726 | 437.127 | 13.7004 | 70.0158 | 142.549 | 16.9482 | 0.261772 | 80.1973 | 8.65471 |
| 120.243 | 51.5053 | 195.257 | 4.33881 | 21.2258 | 24.3812 | 210.398 | 69.681 | 0.170823 | 205.952 | 1.82997 |
| 124.03 | 83.5617 | 278.599 | 4.45848 | 25.328 | 34.7456 | 79.1663 | 0.357583 | 0.07 | 158.667 | 3.76917 |
| 117.514 | 99.3555 | 220.195 | 8.22757 | 13.7736 | 9.01661 | 82.3656 | 1.90134 | 0.0895419 | 188.719 | 4.53755 |
| 124.03 | 72.8822 | 233.701 | 3.55422 | 9.9331 | 21.8985 | 110.998 | 7.04825 | 0.116594 | 174.176 | 1.55596 |
| 123.524 | 114.484 | 244.843 | 17.3731 | 10.5134 | 19.673 | 99.3332 | 1.86039 | 0.187399 | 154.876 | 1.35181 |
| 70.1959 | 105.99 | 264.478 | 579.28 | 32.8523 | 139.89 | 219.607 | 72.7118 | 0.369308 | 143.998 | 33.4576 |
| 68.1088 | 244.008 | 382.802 | 528.308 | 18.4052 | 178.454 | 271.641 | 111.901 | 0.528732 | 72.6389 | 14.8366 |
| 75.7597 | 121.548 | 260.502 | 469.433 | 30.819 | 72.4332 | 211.752 | 44.8717 | 0.21379 | 202.117 | 23.181 |
| 105.478 | 91.1138 | 200.401 | 18.5748 | 25.0674 | 15.0344 | 120.313 | 9.19256 | 0.317352 | 258.23 | 8.01231 |
| 91.1395 | 149.945 | 271.79 | 92.0883 | 24.5292 | 19.0681 | 109.704 | 13.2948 | 0.497726 | 157.796 | 6.01985 |

**Table 1: Experimental data for the experiment RNAseq_data_WT_controls**

- **Observable:** tp63

$$[\mathrm{tp63}](\mathrm{t}) = \log_{10}(\mathrm{reg\_tp63\_fc} \cdot \mathrm{s\_RNA} \cdot [\mathrm{tp63}]) \tag{95}$$

$$\sigma\{[\mathrm{tp63}]\}(\mathrm{t}) = \mathrm{sd\_RNA} \tag{96}$$

- **Observable:** spdef

$$[\mathrm{spdef}](\mathrm{t}) = \log_{10}(\mathrm{reg\_spdef\_fc} \cdot \mathrm{s\_RNA} \cdot [\mathrm{spdef}]) \tag{97}$$

$$\sigma\{[\mathrm{spdef}]\}(\mathrm{t}) = \mathrm{sd\_RNA} \tag{98}$$

##### 1.7.3 Experimental data and model fit

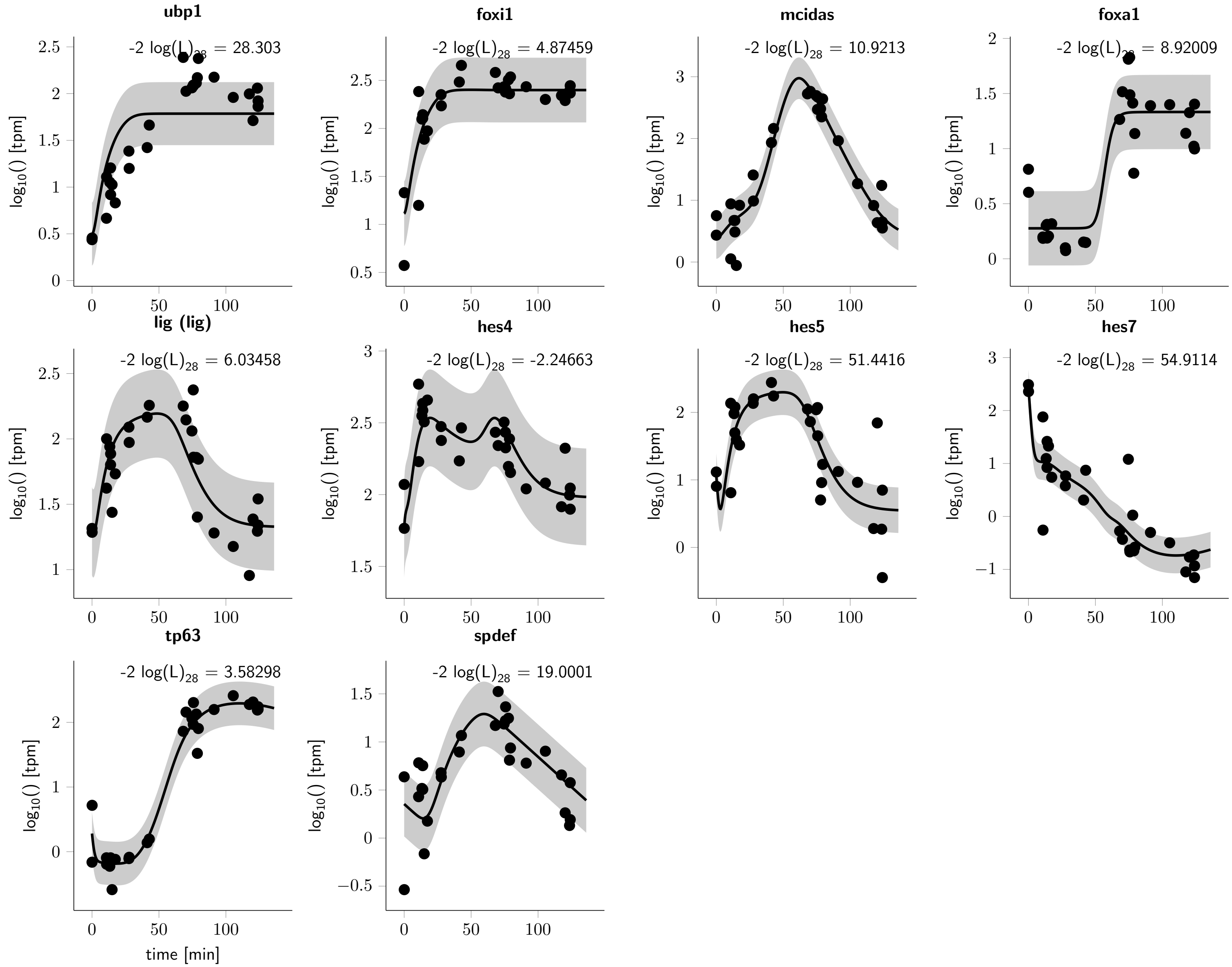

**Figure 2: RNAseq_data_WT_controls observables and experimental data for the experiment.** The observables are displayed as solid lines. The error model that describes the measurement noise is indicated by shades.

#### 1.8 Cell_prior3

##### 1.8.1 Comments

Cell counts for the specified cell types from immunoflourescence images.

##### 1.8.2 Observables

The following observables are modified in this data set.

- **Observable:** bsc2_frac

$$\text{bsc2\_frac}(t) = 0.434294 \cdot \ln\left([\text{bc}] + 0.01\right) - 0.43429448 \cdot \ln\left([\text{bc}] + [\text{ep}] + [\text{mpp}] + 0.01\right) \tag{99}$$

$$\sigma\{\text{bsc2\_frac}\}(t) = 0.0001 \tag{100}$$

##### 1.8.3 Experiment specific conditions

To evaluate the model for this experiment the following conditions are applied.

- **Local condition #2 (global condition #1):**

$$\text{init\_hes4} \rightarrow \frac{\text{init\_hes4}}{\text{reg\_hes4\_fc} \cdot \text{s\_RNA}}$$

$$\text{init\_hes4\_d} \rightarrow \frac{4 \cdot \text{init\_hes4}}{\text{d\_hes4\_prot\_fc} \cdot \text{d\_hes\_prot} \cdot \text{hes\_delay} \cdot \text{reg\_hes4\_fc} \cdot \text{s\_RNA}}$$

$$\text{init\_hes4\_d1} \rightarrow \frac{\text{init\_hes4}}{\text{reg\_hes4\_fc} \cdot \text{s\_RNA}}$$

$$\text{init\_hes4\_d2} \rightarrow \frac{\text{init\_hes4}}{\text{reg\_hes4\_fc} \cdot \text{s\_RNA}}$$

$$\text{init\_hes4\_d3} \rightarrow \frac{\text{init\_hes4}}{\text{reg\_hes4\_fc} \cdot \text{s\_RNA}}$$

$$\text{init\_hes5} \rightarrow \frac{\text{init\_hes5}}{\text{reg\_hes5\_fc} \cdot \text{s\_RNA}}$$

$$\text{init\_hes5\_d} \rightarrow \frac{4 \cdot \text{init\_hes5}}{\text{d\_hes5\_prot\_fc} \cdot \text{d\_hes\_prot} \cdot \text{hes\_delay} \cdot \text{reg\_hes5\_fc} \cdot \text{s\_RNA}}$$

$$\text{init\_hes5\_d1} \rightarrow \frac{\text{init\_hes5}}{\text{reg\_hes5\_fc} \cdot \text{s\_RNA}}$$

$$\text{init\_hes5\_d2} \rightarrow \frac{\text{init\_hes5}}{\text{reg\_hes5\_fc} \cdot \text{s\_RNA}}$$

$$\text{init\_hes5\_d3} \rightarrow \frac{\text{init\_hes5}}{\text{reg\_hes5\_fc} \cdot \text{s\_RNA}}$$

$$\text{init\_hes7} \rightarrow \frac{\text{init\_hes7}}{\text{reg\_hes7\_fc} \cdot \text{s\_RNA}}$$

$$\text{init\_hes7\_d} \rightarrow \frac{4 \cdot \text{init\_hes7}}{\text{d\_hes7\_prot\_fc} \cdot \text{d\_hes\_prot} \cdot \text{hes\_delay} \cdot \text{reg\_hes7\_fc} \cdot \text{s\_RNA}}$$

$$\text{init\_hes7\_d1} \rightarrow \frac{\text{init\_hes7}}{\text{reg\_hes7\_fc} \cdot \text{s\_RNA}}$$

$$\text{init\_hes7\_d2} \rightarrow \frac{\text{init\_hes7}}{\text{reg\_hes7\_fc} \cdot \text{s\_RNA}}$$
$$\text{init\_hes7\_d3} \rightarrow \frac{\text{init\_hes7}}{\text{reg\_hes7\_fc} \cdot \text{s\_RNA}}$$
$$\text{init\_isc} \rightarrow \text{init\_isc}$$
$$\text{init\_mcc} \rightarrow \text{init\_mcc}$$
$$\text{init\_mpp} \rightarrow \text{init\_mpp}$$
$$\text{init\_Spdef} \rightarrow \frac{\text{init\_spdef} \cdot \text{p\_Spdef}}{\text{d\_Spdef} \cdot \text{reg\_spdef\_fc} \cdot \text{s\_RNA}}$$
$$\text{init\_spdef} \rightarrow \frac{\text{init\_spdef}}{\text{reg\_spdef\_fc} \cdot \text{s\_RNA}}$$
$$\text{init\_tp63} \rightarrow \frac{\text{init\_tp63}}{\text{reg\_tp63\_fc} \cdot \text{s\_RNA}}$$
$$\text{k\_hes7\_i\_spdef} \rightarrow \frac{\text{init\_spdef} \cdot \text{k\_hes7\_i\_spdef\_fc} \cdot \text{p\_Spdef}}{\text{d\_Spdef} \cdot \text{reg\_spdef\_fc} \cdot \text{s\_RNA}}$$
$$\text{k\_ssc\_i\_spdef} \rightarrow \frac{\text{init\_spdef} \cdot \text{k\_ssc\_a\_spdef\_fc} \cdot \text{k\_ssc\_i\_spdef\_fc} \cdot \text{p\_Spdef}}{\text{d\_Spdef} \cdot \text{reg\_spdef\_fc} \cdot \text{s\_RNA}}$$
$$\text{k\_ssc\_a\_spdef} \rightarrow \frac{\text{init\_spdef} \cdot \text{k\_ssc\_a\_spdef\_fc} \cdot \text{p\_Spdef}}{\text{d\_Spdef} \cdot \text{reg\_spdef\_fc} \cdot \text{s\_RNA}}$$
$$\text{k\_tp63\_a\_spdef} \rightarrow \frac{\text{init\_spdef} \cdot \text{k\_tp63\_a\_spdef\_fc} \cdot \text{p\_Spdef}}{\text{d\_Spdef} \cdot \text{reg\_spdef\_fc} \cdot \text{s\_RNA}}$$
$$\text{transl\_hes4\_lof\_fc} \rightarrow 1$$
$$\text{transl\_hes5\_lof\_fc} \rightarrow 1$$
$$\text{transl\_hes7\_lof\_fc} \rightarrow 1$$

- **Local condition #3 (global condition #2):**

$$\text{init\_hes4} \rightarrow \frac{\text{init\_hes4}}{\text{reg\_hes4\_fc} \cdot \text{s\_RNA}}$$
$$\text{init\_hes4\_d} \rightarrow \frac{4 \cdot \text{init\_hes4}}{\text{d\_hes4\_prot\_fc} \cdot \text{d\_hes\_prot} \cdot \text{hes\_delay} \cdot \text{reg\_hes4\_fc} \cdot \text{s\_RNA}}$$
$$\text{init\_hes4\_d1} \rightarrow \frac{\text{init\_hes4}}{\text{reg\_hes4\_fc} \cdot \text{s\_RNA}}$$
$$\text{init\_hes4\_d2} \rightarrow \frac{\text{init\_hes4}}{\text{reg\_hes4\_fc} \cdot \text{s\_RNA}}$$
$$\text{init\_hes4\_d3} \rightarrow \frac{\text{init\_hes4}}{\text{reg\_hes4\_fc} \cdot \text{s\_RNA}}$$
$$\text{init\_hes5} \rightarrow \frac{\text{init\_hes5}}{\text{reg\_hes5\_fc} \cdot \text{s\_RNA}}$$
$$\text{init\_hes5\_d} \rightarrow \frac{4 \cdot \text{init\_hes5}}{\text{d\_hes5\_prot\_fc} \cdot \text{d\_hes\_prot} \cdot \text{hes\_delay} \cdot \text{reg\_hes5\_fc} \cdot \text{s\_RNA}}$$
$$\text{init\_hes5\_d1} \rightarrow \frac{\text{init\_hes5}}{\text{reg\_hes5\_fc} \cdot \text{s\_RNA}}$$
$$\text{init\_hes5\_d2} \rightarrow \frac{\text{init\_hes5}}{\text{reg\_hes5\_fc} \cdot \text{s\_RNA}}$$

$$\text{init\_hes5\_d3} \rightarrow \frac{\text{init\_hes5}}{\text{reg\_hes5\_fc} \cdot \text{s\_RNA}}$$

$$\text{init\_hes7} \rightarrow \frac{\text{init\_hes7}}{\text{reg\_hes7\_fc} \cdot \text{s\_RNA}}$$

$$\text{init\_hes7\_d} \rightarrow \frac{4 \cdot \text{init\_hes7} \cdot \text{transl\_hes7\_lof\_fc}}{\text{d\_hes7\_prot\_fc} \cdot \text{d\_hes\_prot} \cdot \text{hes\_delay} \cdot \text{reg\_hes7\_fc} \cdot \text{s\_RNA}}$$

$$\text{init\_hes7\_d1} \rightarrow \frac{\text{init\_hes7} \cdot \text{transl\_hes7\_lof\_fc}}{\text{reg\_hes7\_fc} \cdot \text{s\_RNA}}$$

$$\text{init\_hes7\_d2} \rightarrow \frac{\text{init\_hes7} \cdot \text{transl\_hes7\_lof\_fc}}{\text{reg\_hes7\_fc} \cdot \text{s\_RNA}}$$

$$\text{init\_hes7\_d3} \rightarrow \frac{\text{init\_hes7} \cdot \text{transl\_hes7\_lof\_fc}}{\text{reg\_hes7\_fc} \cdot \text{s\_RNA}}$$

$$\text{init\_isc} \rightarrow \text{init\_isc} \cdot \text{init\_isc\_lof\_hes7\_fc}$$

$$\text{init\_mcc} \rightarrow \text{init\_mcc} \cdot \text{init\_mcc\_lof\_hes7\_fc}$$

$$\text{init\_mpp} \rightarrow \text{init\_mpp} \cdot \text{init\_mpp\_lof\_hes7\_fc}$$

$$\text{init\_Spdef} \rightarrow \frac{\text{init\_spdef} \cdot \text{p\_Spdef}}{\text{d\_Spdef} \cdot \text{reg\_spdef\_fc} \cdot \text{s\_RNA}}$$

$$\text{init\_spdef} \rightarrow \frac{\text{init\_spdef}}{\text{reg\_spdef\_fc} \cdot \text{s\_RNA}}$$

$$\text{init\_tp63} \rightarrow \frac{\text{init\_tp63}}{\text{reg\_tp63\_fc} \cdot \text{s\_RNA}}$$

$$\text{k\_hes7\_i\_spdef} \rightarrow \frac{\text{init\_spdef} \cdot \text{k\_hes7\_i\_spdef\_fc} \cdot \text{p\_Spdef}}{\text{d\_Spdef} \cdot \text{reg\_spdef\_fc} \cdot \text{s\_RNA}}$$

$$\text{k\_ssc\_i\_spdef} \rightarrow \frac{\text{init\_spdef} \cdot \text{k\_ssc\_a\_spdef\_fc} \cdot \text{k\_ssc\_i\_spdef\_fc} \cdot \text{p\_Spdef}}{\text{d\_Spdef} \cdot \text{reg\_spdef\_fc} \cdot \text{s\_RNA}}$$

$$\text{k\_ssc\_a\_spdef} \rightarrow \frac{\text{init\_spdef} \cdot \text{k\_ssc\_a\_spdef\_fc} \cdot \text{p\_Spdef}}{\text{d\_Spdef} \cdot \text{reg\_spdef\_fc} \cdot \text{s\_RNA}}$$

$$\text{k\_tp63\_a\_spdef} \rightarrow \frac{\text{init\_spdef} \cdot \text{k\_tp63\_a\_spdef\_fc} \cdot \text{p\_Spdef}}{\text{d\_Spdef} \cdot \text{reg\_spdef\_fc} \cdot \text{s\_RNA}}$$

$$\text{transl\_hes4\_lof\_fc} \rightarrow 1$$

$$\text{transl\_hes5\_lof\_fc} \rightarrow 1$$

$$\text{transl\_hes7\_lof\_fc} \rightarrow \text{transl\_hes7\_lof\_fc}$$

- **Local condition #4 (global condition #3):**

$$\text{init\_hes4} \rightarrow \frac{\text{init\_hes4}}{\text{reg\_hes4\_fc} \cdot \text{s\_RNA}}$$

$$\text{init\_hes4\_d} \rightarrow \frac{4 \cdot \text{init\_hes4}}{\text{d\_hes4\_prot\_fc} \cdot \text{d\_hes\_prot} \cdot \text{hes\_delay} \cdot \text{reg\_hes4\_fc} \cdot \text{s\_RNA}}$$

$$\text{init\_hes4\_d1} \rightarrow \frac{\text{init\_hes4}}{\text{reg\_hes4\_fc} \cdot \text{s\_RNA}}$$

$$\text{init\_hes4\_d2} \rightarrow \frac{\text{init\_hes4}}{\text{reg\_hes4\_fc} \cdot \text{s\_RNA}}$$

$$\text{init\_hes4\_d3} \rightarrow \frac{\text{init\_hes4}}{\text{reg\_hes4\_fc} \cdot \text{s\_RNA}}$$

$$\mathrm{init\_hes5} \rightarrow \frac{\mathrm{init\_hes5}}{\mathrm{reg\_hes5\_fc} \cdot \mathrm{s\_RNA}}$$

$$\mathrm{init\_hes5\_d} \rightarrow \frac{4 \cdot \mathrm{init\_hes5} \cdot \mathrm{transl\_hes5\_lof\_fc}}{\mathrm{d\_hes5\_prot\_fc} \cdot \mathrm{d\_hes\_prot} \cdot \mathrm{hes\_delay} \cdot \mathrm{reg\_hes5\_fc} \cdot \mathrm{s\_RNA}}$$

$$\mathrm{init\_hes5\_d1} \rightarrow \frac{\mathrm{init\_hes5} \cdot \mathrm{transl\_hes5\_lof\_fc}}{\mathrm{reg\_hes5\_fc} \cdot \mathrm{s\_RNA}}$$

$$\mathrm{init\_hes5\_d2} \rightarrow \frac{\mathrm{init\_hes5} \cdot \mathrm{transl\_hes5\_lof\_fc}}{\mathrm{reg\_hes5\_fc} \cdot \mathrm{s\_RNA}}$$

$$\mathrm{init\_hes5\_d3} \rightarrow \frac{\mathrm{init\_hes5} \cdot \mathrm{transl\_hes5\_lof\_fc}}{\mathrm{reg\_hes5\_fc} \cdot \mathrm{s\_RNA}}$$

$$\mathrm{init\_hes7} \rightarrow \frac{\mathrm{init\_hes7}}{\mathrm{reg\_hes7\_fc} \cdot \mathrm{s\_RNA}}$$

$$\mathrm{init\_hes7\_d} \rightarrow \frac{4 \cdot \mathrm{init\_hes7}}{\mathrm{d\_hes7\_prot\_fc} \cdot \mathrm{d\_hes\_prot} \cdot \mathrm{hes\_delay} \cdot \mathrm{reg\_hes7\_fc} \cdot \mathrm{s\_RNA}}$$

$$\mathrm{init\_hes7\_d1} \rightarrow \frac{\mathrm{init\_hes7}}{\mathrm{reg\_hes7\_fc} \cdot \mathrm{s\_RNA}}$$

$$\mathrm{init\_hes7\_d2} \rightarrow \frac{\mathrm{init\_hes7}}{\mathrm{reg\_hes7\_fc} \cdot \mathrm{s\_RNA}}$$

$$\mathrm{init\_hes7\_d3} \rightarrow \frac{\mathrm{init\_hes7}}{\mathrm{reg\_hes7\_fc} \cdot \mathrm{s\_RNA}}$$

$$\mathrm{init\_isc} \rightarrow \mathrm{init\_isc} \cdot \mathrm{init\_isc\_lof\_hes5\_fc}$$

$$\mathrm{init\_mcc} \rightarrow \mathrm{init\_mcc} \cdot \mathrm{init\_mcc\_lof\_hes5\_fc}$$

$$\mathrm{init\_mpp} \rightarrow \mathrm{init\_mpp} \cdot \mathrm{init\_mpp\_lof\_hes5\_fc}$$

$$\mathrm{init\_Spdef} \rightarrow \frac{\mathrm{init\_spdef} \cdot \mathrm{p\_Spdef}}{\mathrm{d\_Spdef} \cdot \mathrm{reg\_spdef\_fc} \cdot \mathrm{s\_RNA}}$$

$$\mathrm{init\_spdef} \rightarrow \frac{\mathrm{init\_spdef}}{\mathrm{reg\_spdef\_fc} \cdot \mathrm{s\_RNA}}$$

$$\mathrm{init\_tp63} \rightarrow \frac{\mathrm{init\_tp63}}{\mathrm{reg\_tp63\_fc} \cdot \mathrm{s\_RNA}}$$

$$\mathrm{k\_hes7\_i\_spdef} \rightarrow \frac{\mathrm{init\_spdef} \cdot \mathrm{k\_hes7\_i\_spdef\_fc} \cdot \mathrm{p\_Spdef}}{\mathrm{d\_Spdef} \cdot \mathrm{reg\_spdef\_fc} \cdot \mathrm{s\_RNA}}$$

$$\mathrm{k\_ssc\_i\_spdef} \rightarrow \frac{\mathrm{init\_spdef} \cdot \mathrm{k\_ssc\_a\_spdef\_fc} \cdot \mathrm{k\_ssc\_i\_spdef\_fc} \cdot \mathrm{p\_Spdef}}{\mathrm{d\_Spdef} \cdot \mathrm{reg\_spdef\_fc} \cdot \mathrm{s\_RNA}}$$

$$\mathrm{k\_ssc\_a\_spdef} \rightarrow \frac{\mathrm{init\_spdef} \cdot \mathrm{k\_ssc\_a\_spdef\_fc} \cdot \mathrm{p\_Spdef}}{\mathrm{d\_Spdef} \cdot \mathrm{reg\_spdef\_fc} \cdot \mathrm{s\_RNA}}$$

$$\mathrm{k\_tp63\_a\_spdef} \rightarrow \frac{\mathrm{init\_spdef} \cdot \mathrm{k\_tp63\_a\_spdef\_fc} \cdot \mathrm{p\_Spdef}}{\mathrm{d\_Spdef} \cdot \mathrm{reg\_spdef\_fc} \cdot \mathrm{s\_RNA}}$$

$$\mathrm{transl\_hes4\_lof\_fc} \rightarrow 1$$

$$\mathrm{transl\_hes5\_lof\_fc} \rightarrow \mathrm{transl\_hes5\_lof\_fc}$$

$$\mathrm{transl\_hes7\_lof\_fc} \rightarrow 1$$

- **Local condition #5 (global condition #4):**

$$\mathrm{init\_hes4} \rightarrow \frac{\mathrm{init\_hes4}}{\mathrm{reg\_hes4\_fc} \cdot \mathrm{s\_RNA}}$$

$$\text{init\_hes4\_d} \rightarrow \frac{4 \cdot \text{init\_hes4} \cdot \text{transl\_hes4\_lof\_fc}}{\text{d\_hes4\_prot\_fc} \cdot \text{d\_hes\_prot} \cdot \text{hes\_delay} \cdot \text{reg\_hes4\_fc} \cdot \text{s\_RNA}}$$

$$\text{init\_hes4\_d1} \rightarrow \frac{\text{init\_hes4} \cdot \text{transl\_hes4\_lof\_fc}}{\text{reg\_hes4\_fc} \cdot \text{s\_RNA}}$$

$$\text{init\_hes4\_d2} \rightarrow \frac{\text{init\_hes4} \cdot \text{transl\_hes4\_lof\_fc}}{\text{reg\_hes4\_fc} \cdot \text{s\_RNA}}$$

$$\text{init\_hes4\_d3} \rightarrow \frac{\text{init\_hes4} \cdot \text{transl\_hes4\_lof\_fc}}{\text{reg\_hes4\_fc} \cdot \text{s\_RNA}}$$

$$\text{init\_hes5} \rightarrow \frac{\text{init\_hes5}}{\text{reg\_hes5\_fc} \cdot \text{s\_RNA}}$$

$$\text{init\_hes5\_d} \rightarrow \frac{4 \cdot \text{init\_hes5}}{\text{d\_hes5\_prot\_fc} \cdot \text{d\_hes\_prot} \cdot \text{hes\_delay} \cdot \text{reg\_hes5\_fc} \cdot \text{s\_RNA}}$$

$$\text{init\_hes5\_d1} \rightarrow \frac{\text{init\_hes5}}{\text{reg\_hes5\_fc} \cdot \text{s\_RNA}}$$

$$\text{init\_hes5\_d2} \rightarrow \frac{\text{init\_hes5}}{\text{reg\_hes5\_fc} \cdot \text{s\_RNA}}$$

$$\text{init\_hes5\_d3} \rightarrow \frac{\text{init\_hes5}}{\text{reg\_hes5\_fc} \cdot \text{s\_RNA}}$$

$$\text{init\_hes7} \rightarrow \frac{\text{init\_hes7}}{\text{reg\_hes7\_fc} \cdot \text{s\_RNA}}$$

$$\text{init\_hes7\_d} \rightarrow \frac{4 \cdot \text{init\_hes7}}{\text{d\_hes7\_prot\_fc} \cdot \text{d\_hes\_prot} \cdot \text{hes\_delay} \cdot \text{reg\_hes7\_fc} \cdot \text{s\_RNA}}$$

$$\text{init\_hes7\_d1} \rightarrow \frac{\text{init\_hes7}}{\text{reg\_hes7\_fc} \cdot \text{s\_RNA}}$$

$$\text{init\_hes7\_d2} \rightarrow \frac{\text{init\_hes7}}{\text{reg\_hes7\_fc} \cdot \text{s\_RNA}}$$

$$\text{init\_hes7\_d3} \rightarrow \frac{\text{init\_hes7}}{\text{reg\_hes7\_fc} \cdot \text{s\_RNA}}$$

$$\text{init\_isc} \rightarrow \text{init\_isc} \cdot \text{init\_isc\_lof\_hes4\_fc}$$

$$\text{init\_mcc} \rightarrow \text{init\_mcc} \cdot \text{init\_mcc\_lof\_hes4\_fc}$$

$$\text{init\_mpp} \rightarrow \text{init\_mpp} \cdot \text{init\_mpp\_lof\_hes4\_fc}$$

$$\text{init\_Spdef} \rightarrow \frac{\text{init\_spdef} \cdot \text{p\_Spdef}}{\text{d\_Spdef} \cdot \text{reg\_spdef\_fc} \cdot \text{s\_RNA}}$$

$$\text{init\_spdef} \rightarrow \frac{\text{init\_spdef}}{\text{reg\_spdef\_fc} \cdot \text{s\_RNA}}$$

$$\text{init\_tp63} \rightarrow \frac{\text{init\_tp63}}{\text{reg\_tp63\_fc} \cdot \text{s\_RNA}}$$

$$\text{k\_hes7\_i\_spdef} \rightarrow \frac{\text{init\_spdef} \cdot \text{k\_hes7\_i\_spdef\_fc} \cdot \text{p\_Spdef}}{\text{d\_Spdef} \cdot \text{reg\_spdef\_fc} \cdot \text{s\_RNA}}$$

$$\text{k\_ssc\_i\_spdef} \rightarrow \frac{\text{init\_spdef} \cdot \text{k\_ssc\_a\_spdef\_fc} \cdot \text{k\_ssc\_i\_spdef\_fc} \cdot \text{p\_Spdef}}{\text{d\_Spdef} \cdot \text{reg\_spdef\_fc} \cdot \text{s\_RNA}}$$

$$\text{k\_ssc\_a\_spdef} \rightarrow \frac{\text{init\_spdef} \cdot \text{k\_ssc\_a\_spdef\_fc} \cdot \text{p\_Spdef}}{\text{d\_Spdef} \cdot \text{reg\_spdef\_fc} \cdot \text{s\_RNA}}$$

$$\text{k\_tp63\_a\_spdef} \rightarrow \frac{\text{init\_spdef} \cdot \text{k\_tp63\_a\_spdef\_fc} \cdot \text{p\_Spdef}}{\text{d\_Spdef} \cdot \text{reg\_spdef\_fc} \cdot \text{s\_RNA}}$$

$$\text{transl\_hes4\_lof\_fc} \rightarrow \text{transl\_hes4\_lof\_fc}$$
$$\text{transl\_hes5\_lof\_fc} \rightarrow 1$$
$$\text{transl\_hes7\_lof\_fc} \rightarrow 1$$

- **Local condition #6 (global condition #5):**

$$\text{init\_hes4} \rightarrow \frac{\text{init\_hes4}}{\text{reg\_hes4\_fc} \cdot \text{s\_RNA}}$$
$$\text{init\_hes4\_d} \rightarrow \frac{4 \cdot \text{init\_hes4}}{\text{d\_hes4\_prot\_fc} \cdot \text{d\_hes\_prot} \cdot \text{hes\_delay} \cdot \text{reg\_hes4\_fc} \cdot \text{s\_RNA}}$$
$$\text{init\_hes4\_d1} \rightarrow \frac{\text{init\_hes4}}{\text{reg\_hes4\_fc} \cdot \text{s\_RNA}}$$
$$\text{init\_hes4\_d2} \rightarrow \frac{\text{init\_hes4}}{\text{reg\_hes4\_fc} \cdot \text{s\_RNA}}$$
$$\text{init\_hes4\_d3} \rightarrow \frac{\text{init\_hes4}}{\text{reg\_hes4\_fc} \cdot \text{s\_RNA}}$$
$$\text{init\_hes5} \rightarrow \frac{\text{init\_hes5}}{\text{reg\_hes5\_fc} \cdot \text{s\_RNA}}$$
$$\text{init\_hes5\_d} \rightarrow \frac{4 \cdot \text{init\_hes5}}{\text{d\_hes5\_prot\_fc} \cdot \text{d\_hes\_prot} \cdot \text{hes\_delay} \cdot \text{reg\_hes5\_fc} \cdot \text{s\_RNA}}$$
$$\text{init\_hes5\_d1} \rightarrow \frac{\text{init\_hes5}}{\text{reg\_hes5\_fc} \cdot \text{s\_RNA}}$$
$$\text{init\_hes5\_d2} \rightarrow \frac{\text{init\_hes5}}{\text{reg\_hes5\_fc} \cdot \text{s\_RNA}}$$
$$\text{init\_hes5\_d3} \rightarrow \frac{\text{init\_hes5}}{\text{reg\_hes5\_fc} \cdot \text{s\_RNA}}$$
$$\text{init\_hes7} \rightarrow \frac{\text{init\_hes7} \cdot \text{init\_hes7\_gof\_lc\_fc}}{\text{reg\_hes7\_fc} \cdot \text{s\_RNA}}$$
$$\text{init\_hes7\_d} \rightarrow \frac{4 \cdot \text{init\_hes7} \cdot \text{init\_hes7\_gof\_lc\_fc}}{\text{d\_hes7\_prot\_fc} \cdot \text{d\_hes\_prot} \cdot \text{hes\_delay} \cdot \text{reg\_hes7\_fc} \cdot \text{s\_RNA}}$$
$$\text{init\_hes7\_d1} \rightarrow \frac{\text{init\_hes7} \cdot \text{init\_hes7\_gof\_lc\_fc}}{\text{reg\_hes7\_fc} \cdot \text{s\_RNA}}$$
$$\text{init\_hes7\_d2} \rightarrow \frac{\text{init\_hes7} \cdot \text{init\_hes7\_gof\_lc\_fc}}{\text{reg\_hes7\_fc} \cdot \text{s\_RNA}}$$
$$\text{init\_hes7\_d3} \rightarrow \frac{\text{init\_hes7} \cdot \text{init\_hes7\_gof\_lc\_fc}}{\text{reg\_hes7\_fc} \cdot \text{s\_RNA}}$$
$$\text{init\_isc} \rightarrow \text{init\_isc} \cdot \text{init\_isc\_gof\_lc\_hes7\_fc}$$
$$\text{init\_mcc} \rightarrow \text{init\_mcc} \cdot \text{init\_mcc\_gof\_lc\_hes7\_fc}$$
$$\text{init\_mpp} \rightarrow \text{init\_mpp} \cdot \text{init\_mpp\_gof\_lc\_hes7\_fc}$$
$$\text{init\_Spdef} \rightarrow \frac{\text{init\_spdef} \cdot \text{p\_Spdef}}{\text{d\_Spdef} \cdot \text{reg\_spdef\_fc} \cdot \text{s\_RNA}}$$
$$\text{init\_spdef} \rightarrow \frac{\text{init\_spdef}}{\text{reg\_spdef\_fc} \cdot \text{s\_RNA}}$$
$$\text{init\_tp63} \rightarrow \frac{\text{init\_tp63}}{\text{reg\_tp63\_fc} \cdot \text{s\_RNA}}$$

$$\text{k\_hes7\_i\_spdef} \rightarrow \frac{\text{init\_spdef} \cdot \text{k\_hes7\_i\_spdef\_fc} \cdot \text{p\_Spdef}}{\text{d\_Spdef} \cdot \text{reg\_spdef\_fc} \cdot \text{s\_RNA}}$$

$$\text{k\_ssc\_i\_spdef} \rightarrow \frac{\text{init\_spdef} \cdot \text{k\_ssc\_a\_spdef\_fc} \cdot \text{k\_ssc\_i\_spdef\_fc} \cdot \text{p\_Spdef}}{\text{d\_Spdef} \cdot \text{reg\_spdef\_fc} \cdot \text{s\_RNA}}$$

$$\text{k\_ssc\_a\_spdef} \rightarrow \frac{\text{init\_spdef} \cdot \text{k\_ssc\_a\_spdef\_fc} \cdot \text{p\_Spdef}}{\text{d\_Spdef} \cdot \text{reg\_spdef\_fc} \cdot \text{s\_RNA}}$$

$$\text{k\_tp63\_a\_spdef} \rightarrow \frac{\text{init\_spdef} \cdot \text{k\_tp63\_a\_spdef\_fc} \cdot \text{p\_Spdef}}{\text{d\_Spdef} \cdot \text{reg\_spdef\_fc} \cdot \text{s\_RNA}}$$

$$\text{transl\_hes4\_lof\_fc} \rightarrow 1$$

$$\text{transl\_hes5\_lof\_fc} \rightarrow 1$$

$$\text{transl\_hes7\_lof\_fc} \rightarrow 1$$

- **Local condition #7 (global condition #6):**

$$\text{init\_hes4} \rightarrow \frac{\text{init\_hes4}}{\text{reg\_hes4\_fc} \cdot \text{s\_RNA}}$$

$$\text{init\_hes4\_d} \rightarrow \frac{4 \cdot \text{init\_hes4}}{\text{d\_hes4\_prot\_fc} \cdot \text{d\_hes\_prot} \cdot \text{hes\_delay} \cdot \text{reg\_hes4\_fc} \cdot \text{s\_RNA}}$$

$$\text{init\_hes4\_d1} \rightarrow \frac{\text{init\_hes4}}{\text{reg\_hes4\_fc} \cdot \text{s\_RNA}}$$

$$\text{init\_hes4\_d2} \rightarrow \frac{\text{init\_hes4}}{\text{reg\_hes4\_fc} \cdot \text{s\_RNA}}$$

$$\text{init\_hes4\_d3} \rightarrow \frac{\text{init\_hes4}}{\text{reg\_hes4\_fc} \cdot \text{s\_RNA}}$$

$$\text{init\_hes5} \rightarrow \frac{\text{init\_hes5} \cdot \text{init\_hes5\_gof\_lc\_fc}}{\text{reg\_hes5\_fc} \cdot \text{s\_RNA}}$$

$$\text{init\_hes5\_d} \rightarrow \frac{4 \cdot \text{init\_hes5} \cdot \text{init\_hes5\_gof\_lc\_fc}}{\text{d\_hes5\_prot\_fc} \cdot \text{d\_hes\_prot} \cdot \text{hes\_delay} \cdot \text{reg\_hes5\_fc} \cdot \text{s\_RNA}}$$

$$\text{init\_hes5\_d1} \rightarrow \frac{\text{init\_hes5} \cdot \text{init\_hes5\_gof\_lc\_fc}}{\text{reg\_hes5\_fc} \cdot \text{s\_RNA}}$$

$$\text{init\_hes5\_d2} \rightarrow \frac{\text{init\_hes5} \cdot \text{init\_hes5\_gof\_lc\_fc}}{\text{reg\_hes5\_fc} \cdot \text{s\_RNA}}$$

$$\text{init\_hes5\_d3} \rightarrow \frac{\text{init\_hes5} \cdot \text{init\_hes5\_gof\_lc\_fc}}{\text{reg\_hes5\_fc} \cdot \text{s\_RNA}}$$

$$\text{init\_hes7} \rightarrow \frac{\text{init\_hes7}}{\text{reg\_hes7\_fc} \cdot \text{s\_RNA}}$$

$$\text{init\_hes7\_d} \rightarrow \frac{4 \cdot \text{init\_hes7}}{\text{d\_hes7\_prot\_fc} \cdot \text{d\_hes\_prot} \cdot \text{hes\_delay} \cdot \text{reg\_hes7\_fc} \cdot \text{s\_RNA}}$$

$$\text{init\_hes7\_d1} \rightarrow \frac{\text{init\_hes7}}{\text{reg\_hes7\_fc} \cdot \text{s\_RNA}}$$

$$\text{init\_hes7\_d2} \rightarrow \frac{\text{init\_hes7}}{\text{reg\_hes7\_fc} \cdot \text{s\_RNA}}$$

$$\text{init\_hes7\_d3} \rightarrow \frac{\text{init\_hes7}}{\text{reg\_hes7\_fc} \cdot \text{s\_RNA}}$$

$$\text{init\_isc} \rightarrow \text{init\_isc} \cdot \text{init\_isc\_gof\_lc\_hes5\_fc}$$

$$\mathrm{init\_mcc} \rightarrow \mathrm{init\_mcc} \cdot \mathrm{init\_mcc\_gof\_lc\_hes5\_fc}$$

$$\mathrm{init\_mpp} \rightarrow \mathrm{init\_mpp} \cdot \mathrm{init\_mpp\_gof\_lc\_hes5\_fc}$$

$$\mathrm{init\_Spdef} \rightarrow \frac{\mathrm{init\_spdef} \cdot \mathrm{p\_Spdef}}{\mathrm{d\_Spdef} \cdot \mathrm{reg\_spdef\_fc} \cdot \mathrm{s\_RNA}}$$

$$\mathrm{init\_spdef} \rightarrow \frac{\mathrm{init\_spdef}}{\mathrm{reg\_spdef\_fc} \cdot \mathrm{s\_RNA}}$$

$$\mathrm{init\_tp63} \rightarrow \frac{\mathrm{init\_tp63}}{\mathrm{reg\_tp63\_fc} \cdot \mathrm{s\_RNA}}$$

$$\mathrm{k\_hes7\_i\_spdef} \rightarrow \frac{\mathrm{init\_spdef} \cdot \mathrm{k\_hes7\_i\_spdef\_fc} \cdot \mathrm{p\_Spdef}}{\mathrm{d\_Spdef} \cdot \mathrm{reg\_spdef\_fc} \cdot \mathrm{s\_RNA}}$$

$$\mathrm{k\_ssc\_i\_spdef} \rightarrow \frac{\mathrm{init\_spdef} \cdot \mathrm{k\_ssc\_a\_spdef\_fc} \cdot \mathrm{k\_ssc\_i\_spdef\_fc} \cdot \mathrm{p\_Spdef}}{\mathrm{d\_Spdef} \cdot \mathrm{reg\_spdef\_fc} \cdot \mathrm{s\_RNA}}$$

$$\mathrm{k\_ssc\_a\_spdef} \rightarrow \frac{\mathrm{init\_spdef} \cdot \mathrm{k\_ssc\_a\_spdef\_fc} \cdot \mathrm{p\_Spdef}}{\mathrm{d\_Spdef} \cdot \mathrm{reg\_spdef\_fc} \cdot \mathrm{s\_RNA}}$$

$$\mathrm{k\_tp63\_a\_spdef} \rightarrow \frac{\mathrm{init\_spdef} \cdot \mathrm{k\_tp63\_a\_spdef\_fc} \cdot \mathrm{p\_Spdef}}{\mathrm{d\_Spdef} \cdot \mathrm{reg\_spdef\_fc} \cdot \mathrm{s\_RNA}}$$

$$\mathrm{transl\_hes4\_lof\_fc} \rightarrow 1$$

$$\mathrm{transl\_hes5\_lof\_fc} \rightarrow 1$$

$$\mathrm{transl\_hes7\_lof\_fc} \rightarrow 1$$

- **Local condition #8 (global condition #7):**

$$\mathrm{init\_hes4} \rightarrow \frac{\mathrm{init\_hes4} \cdot \mathrm{init\_hes4\_gof\_lc\_fc}}{\mathrm{reg\_hes4\_fc} \cdot \mathrm{s\_RNA}}$$

$$\mathrm{init\_hes4\_d} \rightarrow \frac{4 \cdot \mathrm{init\_hes4} \cdot \mathrm{init\_hes4\_gof\_lc\_fc}}{\mathrm{d\_hes4\_prot\_fc} \cdot \mathrm{d\_hes\_prot} \cdot \mathrm{hes\_delay} \cdot \mathrm{reg\_hes4\_fc} \cdot \mathrm{s\_RNA}}$$

$$\mathrm{init\_hes4\_d1} \rightarrow \frac{\mathrm{init\_hes4} \cdot \mathrm{init\_hes4\_gof\_lc\_fc}}{\mathrm{reg\_hes4\_fc} \cdot \mathrm{s\_RNA}}$$

$$\mathrm{init\_hes4\_d2} \rightarrow \frac{\mathrm{init\_hes4} \cdot \mathrm{init\_hes4\_gof\_lc\_fc}}{\mathrm{reg\_hes4\_fc} \cdot \mathrm{s\_RNA}}$$

$$\mathrm{init\_hes4\_d3} \rightarrow \frac{\mathrm{init\_hes4} \cdot \mathrm{init\_hes4\_gof\_lc\_fc}}{\mathrm{reg\_hes4\_fc} \cdot \mathrm{s\_RNA}}$$

$$\mathrm{init\_hes5} \rightarrow \frac{\mathrm{init\_hes5}}{\mathrm{reg\_hes5\_fc} \cdot \mathrm{s\_RNA}}$$

$$\mathrm{init\_hes5\_d} \rightarrow \frac{4 \cdot \mathrm{init\_hes5}}{\mathrm{d\_hes5\_prot\_fc} \cdot \mathrm{d\_hes\_prot} \cdot \mathrm{hes\_delay} \cdot \mathrm{reg\_hes5\_fc} \cdot \mathrm{s\_RNA}}$$

$$\mathrm{init\_hes5\_d1} \rightarrow \frac{\mathrm{init\_hes5}}{\mathrm{reg\_hes5\_fc} \cdot \mathrm{s\_RNA}}$$

$$\mathrm{init\_hes5\_d2} \rightarrow \frac{\mathrm{init\_hes5}}{\mathrm{reg\_hes5\_fc} \cdot \mathrm{s\_RNA}}$$

$$\mathrm{init\_hes5\_d3} \rightarrow \frac{\mathrm{init\_hes5}}{\mathrm{reg\_hes5\_fc} \cdot \mathrm{s\_RNA}}$$

$$\mathrm{init\_hes7} \rightarrow \frac{\mathrm{init\_hes7}}{\mathrm{reg\_hes7\_fc} \cdot \mathrm{s\_RNA}}$$

| time [min] | bc_frac number of cells |
|---|---|
| 124.03 | 0 |
| 124.03 | 0 |
| 124.03 | 0 |
| 124.03 | 0 |
| 124.03 | 0 |
| 124.03 | 0 |
| 124.03 | 0 |

**Table 2: Prior3**

$$\mathrm{init\_hes7\_d} \rightarrow \frac{4 \cdot \mathrm{init\_hes7}}{\mathrm{d\_hes7\_prot\_fc} \cdot \mathrm{d\_hes\_prot} \cdot \mathrm{hes\_delay} \cdot \mathrm{reg\_hes7\_fc} \cdot \mathrm{s\_RNA}}$$
$$\mathrm{init\_hes7\_d1} \rightarrow \frac{\mathrm{init\_hes7}}{\mathrm{reg\_hes7\_fc} \cdot \mathrm{s\_RNA}}$$
$$\mathrm{init\_hes7\_d2} \rightarrow \frac{\mathrm{init\_hes7}}{\mathrm{reg\_hes7\_fc} \cdot \mathrm{s\_RNA}}$$
$$\mathrm{init\_hes7\_d3} \rightarrow \frac{\mathrm{init\_hes7}}{\mathrm{reg\_hes7\_fc} \cdot \mathrm{s\_RNA}}$$
$$\mathrm{init\_isc} \rightarrow \mathrm{init\_isc} \cdot \mathrm{init\_isc\_gof\_lc\_hes4\_fc}$$
$$\mathrm{init\_mcc} \rightarrow \mathrm{init\_mcc} \cdot \mathrm{init\_mcc\_gof\_lc\_hes4\_fc}$$
$$\mathrm{init\_mpp} \rightarrow \mathrm{init\_mpp} \cdot \mathrm{init\_mpp\_gof\_lc\_hes4\_fc}$$
$$\mathrm{init\_Spdef} \rightarrow \frac{\mathrm{init\_spdef} \cdot \mathrm{p\_Spdef}}{\mathrm{d\_Spdef} \cdot \mathrm{reg\_spdef\_fc} \cdot \mathrm{s\_RNA}}$$
$$\mathrm{init\_spdef} \rightarrow \frac{\mathrm{init\_spdef}}{\mathrm{reg\_spdef\_fc} \cdot \mathrm{s\_RNA}}$$
$$\mathrm{init\_tp63} \rightarrow \frac{\mathrm{init\_tp63}}{\mathrm{reg\_tp63\_fc} \cdot \mathrm{s\_RNA}}$$
$$\mathrm{k\_hes7\_i\_spdef} \rightarrow \frac{\mathrm{init\_spdef} \cdot \mathrm{k\_hes7\_i\_spdef\_fc} \cdot \mathrm{p\_Spdef}}{\mathrm{d\_Spdef} \cdot \mathrm{reg\_spdef\_fc} \cdot \mathrm{s\_RNA}}$$
$$\mathrm{k\_ssc\_i\_spdef} \rightarrow \frac{\mathrm{init\_spdef} \cdot \mathrm{k\_ssc\_a\_spdef\_fc} \cdot \mathrm{k\_ssc\_i\_spdef\_fc} \cdot \mathrm{p\_Spdef}}{\mathrm{d\_Spdef} \cdot \mathrm{reg\_spdef\_fc} \cdot \mathrm{s\_RNA}}$$
$$\mathrm{k\_ssc\_a\_spdef} \rightarrow \frac{\mathrm{init\_spdef} \cdot \mathrm{k\_ssc\_a\_spdef\_fc} \cdot \mathrm{p\_Spdef}}{\mathrm{d\_Spdef} \cdot \mathrm{reg\_spdef\_fc} \cdot \mathrm{s\_RNA}}$$
$$\mathrm{k\_tp63\_a\_spdef} \rightarrow \frac{\mathrm{init\_spdef} \cdot \mathrm{k\_tp63\_a\_spdef\_fc} \cdot \mathrm{p\_Spdef}}{\mathrm{d\_Spdef} \cdot \mathrm{reg\_spdef\_fc} \cdot \mathrm{s\_RNA}}$$
$$\mathrm{transl\_hes4\_lof\_fc} \rightarrow 1$$
$$\mathrm{transl\_hes5\_lof\_fc} \rightarrow 1$$
$$\mathrm{transl\_hes7\_lof\_fc} \rightarrow 1$$

##### 1.8.4 Experimental data and model fit

#### 1.9 Experiment: Controls_Cells_aly_new

##### 1.9.1 Comments

Cell counts for RNA perturbation experiments

##### 1.9.2 Observables

The following observables are modified in this data set.

- **Observable:** isc_abs

$$\begin{aligned}&\mathrm{isc\_abs(t)}\\&=[\mathrm{isc}]\cdot(\mathrm{ex1}\cdot\mathrm{s\_cell1}+\mathrm{ex12}\cdot\mathrm{s\_cell12}+\mathrm{ex13}\cdot\mathrm{s\_cell13}+\mathrm{ex14}\cdot\mathrm{s\_cell14}+\mathrm{ex2}\cdot\mathrm{s\_cell2}+\mathrm{ex20}\cdot\mathrm{s\_cell20}+\mathrm{ex22}\cdot\mathrm{s\_cell22}+\mathrm{ex23}\cdot\mathrm{s\_cell23}+\mathrm{ex24}\cdot\mathrm{s\_cell24}\\&+\mathrm{ex25}\cdot\mathrm{s\_cell25}+\mathrm{ex3}\cdot\mathrm{s\_cell3}+\mathrm{ex4}\cdot\mathrm{s\_cell4}+\mathrm{ex5}\cdot\mathrm{s\_cell5}+\mathrm{ex8}\cdot\mathrm{s\_cell8}+\mathrm{ex9}\cdot\mathrm{s\_cell9}\end{aligned}\tag{101}$$

$$\sigma\{\mathrm{isc\_abs}\}(\mathrm{t})=\mathrm{sd\_cell}\tag{102}$$

- **Observable:** mcc_abs

$$\begin{aligned}&\mathrm{mcc\_abs(t)}\\&=[\mathrm{mcc\_total}]\cdot(\mathrm{ex1}\cdot\mathrm{s\_cell1}+\mathrm{ex12}\cdot\mathrm{s\_cell12}+\mathrm{ex13}\cdot\mathrm{s\_cell13}+\mathrm{ex14}\cdot\mathrm{s\_cell14}+\mathrm{ex2}\cdot\mathrm{s\_cell2}+\mathrm{ex20}\cdot\mathrm{s\_cell20}+\mathrm{ex22}\cdot\mathrm{s\_cell22}+\mathrm{ex23}\cdot\mathrm{s\_cell23}+\mathrm{ex24}\cdot\mathrm{s\_cell24}\\&+\mathrm{ex25}\cdot\mathrm{s\_cell25}+\mathrm{ex3}\cdot\mathrm{s\_cell3}+\mathrm{ex4}\cdot\mathrm{s\_cell4}+\mathrm{ex5}\cdot\mathrm{s\_cell5}+\mathrm{ex8}\cdot\mathrm{s\_cell8}+\mathrm{ex9}\cdot\mathrm{s\_cell9}\end{aligned}\tag{103}$$

$$\sigma\{\mathrm{mcc\_abs}\}(\mathrm{t})=\mathrm{sd\_cell}\tag{104}$$

- **Observable:** ssc_abs

$$\begin{aligned}\mathrm{ssc\_abs(t)}=[\mathrm{ssc}]\cdot(&\mathrm{ex1}\cdot\mathrm{s\_cell1}+\mathrm{ex12}\cdot\mathrm{s\_cell12}+\mathrm{ex13}\cdot\mathrm{s\_cell13}+\mathrm{ex14}\cdot\mathrm{s\_cell14}+\mathrm{ex2}\cdot\mathrm{s\_cell2}+\mathrm{ex20}\cdot\mathrm{s\_cell20}+\mathrm{ex22}\cdot\mathrm{s\_cell22}+\mathrm{ex23}\cdot\mathrm{s\_cell23}+\mathrm{ex24}\cdot\mathrm{s\_cell24}\\&+\mathrm{ex25}\cdot\mathrm{s\_cell25}+\mathrm{ex3}\cdot\mathrm{s\_cell3}+\mathrm{ex4}\cdot\mathrm{s\_cell4}+\mathrm{ex5}\cdot\mathrm{s\_cell5}+\mathrm{ex8}\cdot\mathrm{s\_cell8}+\mathrm{ex9}\cdot\mathrm{s\_cell9}\end{aligned}\tag{105}$$

$$\sigma\{\mathrm{ssc\_abs}\}(\mathrm{t})=\mathrm{sd\_cell}\tag{106}$$

##### 1.9.3 Experiment specific conditions

To evaluate the model for this experiment the following conditions are applied.

- **Local condition #9 (global condition #1):**

$$\begin{aligned}\mathrm{ex1}&\rightarrow 0\\\mathrm{ex12}&\rightarrow 0\\\mathrm{ex13}&\rightarrow 0\\\mathrm{ex14}&\rightarrow 0\\\mathrm{ex2}&\rightarrow 0\end{aligned}$$

$$\mathrm{ex20} \to 0$$
$$\mathrm{ex22} \to 0$$
$$\mathrm{ex23} \to 0$$
$$\mathrm{ex24} \to 0$$
$$\mathrm{ex25} \to 1$$
$$\mathrm{ex3} \to 0$$
$$\mathrm{ex4} \to 0$$
$$\mathrm{ex5} \to 0$$
$$\mathrm{ex8} \to 0$$
$$\mathrm{ex9} \to 0$$
$$\mathrm{init\_hes4} \to \frac{\mathrm{init\_hes4}}{\mathrm{reg\_hes4\_fc} \cdot \mathrm{s\_RNA}}$$
$$\mathrm{init\_hes4\_d} \to \frac{4 \cdot \mathrm{init\_hes4}}{\mathrm{d\_hes4\_prot\_fc} \cdot \mathrm{d\_hes\_prot} \cdot \mathrm{hes\_delay} \cdot \mathrm{reg\_hes4\_fc} \cdot \mathrm{s\_RNA}}$$
$$\mathrm{init\_hes4\_d1} \to \frac{\mathrm{init\_hes4}}{\mathrm{reg\_hes4\_fc} \cdot \mathrm{s\_RNA}}$$
$$\mathrm{init\_hes4\_d2} \to \frac{\mathrm{init\_hes4}}{\mathrm{reg\_hes4\_fc} \cdot \mathrm{s\_RNA}}$$
$$\mathrm{init\_hes4\_d3} \to \frac{\mathrm{init\_hes4}}{\mathrm{reg\_hes4\_fc} \cdot \mathrm{s\_RNA}}$$
$$\mathrm{init\_hes5} \to \frac{\mathrm{init\_hes5}}{\mathrm{reg\_hes5\_fc} \cdot \mathrm{s\_RNA}}$$
$$\mathrm{init\_hes5\_d} \to \frac{4 \cdot \mathrm{init\_hes5}}{\mathrm{d\_hes5\_prot\_fc} \cdot \mathrm{d\_hes\_prot} \cdot \mathrm{hes\_delay} \cdot \mathrm{reg\_hes5\_fc} \cdot \mathrm{s\_RNA}}$$
$$\mathrm{init\_hes5\_d1} \to \frac{\mathrm{init\_hes5}}{\mathrm{reg\_hes5\_fc} \cdot \mathrm{s\_RNA}}$$
$$\mathrm{init\_hes5\_d2} \to \frac{\mathrm{init\_hes5}}{\mathrm{reg\_hes5\_fc} \cdot \mathrm{s\_RNA}}$$
$$\mathrm{init\_hes5\_d3} \to \frac{\mathrm{init\_hes5}}{\mathrm{reg\_hes5\_fc} \cdot \mathrm{s\_RNA}}$$
$$\mathrm{init\_hes7} \to \frac{\mathrm{init\_hes7}}{\mathrm{reg\_hes7\_fc} \cdot \mathrm{s\_RNA}}$$
$$\mathrm{init\_hes7\_d} \to \frac{4 \cdot \mathrm{init\_hes7}}{\mathrm{d\_hes7\_prot\_fc} \cdot \mathrm{d\_hes\_prot} \cdot \mathrm{hes\_delay} \cdot \mathrm{reg\_hes7\_fc} \cdot \mathrm{s\_RNA}}$$
$$\mathrm{init\_hes7\_d1} \to \frac{\mathrm{init\_hes7}}{\mathrm{reg\_hes7\_fc} \cdot \mathrm{s\_RNA}}$$
$$\mathrm{init\_hes7\_d2} \to \frac{\mathrm{init\_hes7}}{\mathrm{reg\_hes7\_fc} \cdot \mathrm{s\_RNA}}$$
$$\mathrm{init\_hes7\_d3} \to \frac{\mathrm{init\_hes7}}{\mathrm{reg\_hes7\_fc} \cdot \mathrm{s\_RNA}}$$
$$\mathrm{init\_Spdef} \to \frac{\mathrm{init\_spdef} \cdot \mathrm{p\_Spdef}}{\mathrm{d\_Spdef} \cdot \mathrm{reg\_spdef\_fc} \cdot \mathrm{s\_RNA}}$$
$$\mathrm{init\_spdef} \to \frac{\mathrm{init\_spdef}}{\mathrm{reg\_spdef\_fc} \cdot \mathrm{s\_RNA}}$$

$$\text{init\_tp63} \rightarrow \frac{\text{init\_tp63}}{\text{reg\_tp63\_fc} \cdot \text{s\_RNA}}$$
$$\text{k\_hes7\_i\_spdef} \rightarrow \frac{\text{init\_spdef} \cdot \text{k\_hes7\_i\_spdef\_fc} \cdot \text{p\_Spdef}}{\text{d\_Spdef} \cdot \text{reg\_spdef\_fc} \cdot \text{s\_RNA}}$$
$$\text{k\_ssc\_i\_spdef} \rightarrow \frac{\text{init\_spdef} \cdot \text{k\_ssc\_a\_spdef\_fc} \cdot \text{k\_ssc\_i\_spdef\_fc} \cdot \text{p\_Spdef}}{\text{d\_Spdef} \cdot \text{reg\_spdef\_fc} \cdot \text{s\_RNA}}$$
$$\text{k\_ssc\_a\_spdef} \rightarrow \frac{\text{init\_spdef} \cdot \text{k\_ssc\_a\_spdef\_fc} \cdot \text{p\_Spdef}}{\text{d\_Spdef} \cdot \text{reg\_spdef\_fc} \cdot \text{s\_RNA}}$$
$$\text{k\_tp63\_a\_spdef} \rightarrow \frac{\text{init\_spdef} \cdot \text{k\_tp63\_a\_spdef\_fc} \cdot \text{p\_Spdef}}{\text{d\_Spdef} \cdot \text{reg\_spdef\_fc} \cdot \text{s\_RNA}}$$
$$\text{s\_cell1} \rightarrow \text{s\_cell1\_fc} \cdot \text{s\_cell\_mean}$$
$$\text{s\_cell12} \rightarrow \text{s\_cell12\_fc} \cdot \text{s\_cell\_mean}$$
$$\text{s\_cell13} \rightarrow \text{s\_cell13\_fc} \cdot \text{s\_cell\_mean}$$
$$\text{s\_cell14} \rightarrow \text{s\_cell14\_fc} \cdot \text{s\_cell\_mean}$$
$$\text{s\_cell2} \rightarrow \text{s\_cell2\_fc} \cdot \text{s\_cell\_mean}$$
$$\text{s\_cell20} \rightarrow \text{s\_cell20\_fc} \cdot \text{s\_cell\_mean}$$
$$\text{s\_cell22} \rightarrow \text{s\_cell22\_fc} \cdot \text{s\_cell\_mean}$$
$$\text{s\_cell23} \rightarrow \text{s\_cell23\_fc} \cdot \text{s\_cell\_mean}$$
$$\text{s\_cell24} \rightarrow \text{s\_cell24\_fc} \cdot \text{s\_cell\_mean}$$
$$\text{s\_cell25} \rightarrow \text{s\_cell25\_fc} \cdot \text{s\_cell\_mean}$$
$$\text{s\_cell3} \rightarrow \text{s\_cell3\_fc} \cdot \text{s\_cell\_mean}$$
$$\text{s\_cell4} \rightarrow \text{s\_cell4\_fc} \cdot \text{s\_cell\_mean}$$
$$\text{s\_cell5} \rightarrow \text{s\_cell5\_fc} \cdot \text{s\_cell\_mean}$$
$$\text{s\_cell8} \rightarrow \text{s\_cell8\_fc} \cdot \text{s\_cell\_mean}$$
$$\text{s\_cell9} \rightarrow \text{s\_cell9\_fc} \cdot \text{s\_cell\_mean}$$
$$\text{transl\_hes4\_lof\_fc} \rightarrow 1$$
$$\text{transl\_hes5\_lof\_fc} \rightarrow 1$$
$$\text{transl\_hes7\_lof\_fc} \rightarrow 1$$

- **Local condition #10 (global condition #1):**

$$\text{ex1} \rightarrow 0$$
$$\text{ex12} \rightarrow 0$$
$$\text{ex13} \rightarrow 0$$
$$\text{ex14} \rightarrow 0$$
$$\text{ex2} \rightarrow 0$$
$$\text{ex20} \rightarrow 0$$
$$\text{ex22} \rightarrow 0$$
$$\text{ex23} \rightarrow 0$$
$$\text{ex24} \rightarrow 1$$
$$\text{ex25} \rightarrow 0$$
$$\text{ex3} \rightarrow 0$$
$$\text{ex4} \rightarrow 0$$

$$\mathrm{ex5} \rightarrow 0$$
$$\mathrm{ex8} \rightarrow 0$$
$$\mathrm{ex9} \rightarrow 0$$
$$\mathrm{init\_hes4} \rightarrow \frac{\mathrm{init\_hes4}}{\mathrm{reg\_hes4\_fc} \cdot \mathrm{s\_RNA}}$$
$$\mathrm{init\_hes4\_d} \rightarrow \frac{4 \cdot \mathrm{init\_hes4}}{\mathrm{d\_hes4\_prot\_fc} \cdot \mathrm{d\_hes\_prot} \cdot \mathrm{hes\_delay} \cdot \mathrm{reg\_hes4\_fc} \cdot \mathrm{s\_RNA}}$$
$$\mathrm{init\_hes4\_d1} \rightarrow \frac{\mathrm{init\_hes4}}{\mathrm{reg\_hes4\_fc} \cdot \mathrm{s\_RNA}}$$
$$\mathrm{init\_hes4\_d2} \rightarrow \frac{\mathrm{init\_hes4}}{\mathrm{reg\_hes4\_fc} \cdot \mathrm{s\_RNA}}$$
$$\mathrm{init\_hes4\_d3} \rightarrow \frac{\mathrm{init\_hes4}}{\mathrm{reg\_hes4\_fc} \cdot \mathrm{s\_RNA}}$$
$$\mathrm{init\_hes5} \rightarrow \frac{\mathrm{init\_hes5}}{\mathrm{reg\_hes5\_fc} \cdot \mathrm{s\_RNA}}$$
$$\mathrm{init\_hes5\_d} \rightarrow \frac{4 \cdot \mathrm{init\_hes5}}{\mathrm{d\_hes5\_prot\_fc} \cdot \mathrm{d\_hes\_prot} \cdot \mathrm{hes\_delay} \cdot \mathrm{reg\_hes5\_fc} \cdot \mathrm{s\_RNA}}$$
$$\mathrm{init\_hes5\_d1} \rightarrow \frac{\mathrm{init\_hes5}}{\mathrm{reg\_hes5\_fc} \cdot \mathrm{s\_RNA}}$$
$$\mathrm{init\_hes5\_d2} \rightarrow \frac{\mathrm{init\_hes5}}{\mathrm{reg\_hes5\_fc} \cdot \mathrm{s\_RNA}}$$
$$\mathrm{init\_hes5\_d3} \rightarrow \frac{\mathrm{init\_hes5}}{\mathrm{reg\_hes5\_fc} \cdot \mathrm{s\_RNA}}$$
$$\mathrm{init\_hes7} \rightarrow \frac{\mathrm{init\_hes7}}{\mathrm{reg\_hes7\_fc} \cdot \mathrm{s\_RNA}}$$
$$\mathrm{init\_hes7\_d} \rightarrow \frac{4 \cdot \mathrm{init\_hes7}}{\mathrm{d\_hes7\_prot\_fc} \cdot \mathrm{d\_hes\_prot} \cdot \mathrm{hes\_delay} \cdot \mathrm{reg\_hes7\_fc} \cdot \mathrm{s\_RNA}}$$
$$\mathrm{init\_hes7\_d1} \rightarrow \frac{\mathrm{init\_hes7}}{\mathrm{reg\_hes7\_fc} \cdot \mathrm{s\_RNA}}$$
$$\mathrm{init\_hes7\_d2} \rightarrow \frac{\mathrm{init\_hes7}}{\mathrm{reg\_hes7\_fc} \cdot \mathrm{s\_RNA}}$$
$$\mathrm{init\_hes7\_d3} \rightarrow \frac{\mathrm{init\_hes7}}{\mathrm{reg\_hes7\_fc} \cdot \mathrm{s\_RNA}}$$
$$\mathrm{init\_Spdef} \rightarrow \frac{\mathrm{init\_spdef} \cdot \mathrm{p\_Spdef}}{\mathrm{d\_Spdef} \cdot \mathrm{reg\_spdef\_fc} \cdot \mathrm{s\_RNA}}$$
$$\mathrm{init\_spdef} \rightarrow \frac{\mathrm{init\_spdef}}{\mathrm{reg\_spdef\_fc} \cdot \mathrm{s\_RNA}}$$
$$\mathrm{init\_tp63} \rightarrow \frac{\mathrm{init\_tp63}}{\mathrm{reg\_tp63\_fc} \cdot \mathrm{s\_RNA}}$$
$$\mathrm{k\_hes7\_i\_spdef} \rightarrow \frac{\mathrm{init\_spdef} \cdot \mathrm{k\_hes7\_i\_spdef\_fc} \cdot \mathrm{p\_Spdef}}{\mathrm{d\_Spdef} \cdot \mathrm{reg\_spdef\_fc} \cdot \mathrm{s\_RNA}}$$
$$\mathrm{k\_ssc\_i\_spdef} \rightarrow \frac{\mathrm{init\_spdef} \cdot \mathrm{k\_ssc\_a\_spdef\_fc} \cdot \mathrm{k\_ssc\_i\_spdef\_fc} \cdot \mathrm{p\_Spdef}}{\mathrm{d\_Spdef} \cdot \mathrm{reg\_spdef\_fc} \cdot \mathrm{s\_RNA}}$$
$$\mathrm{k\_ssc\_a\_spdef} \rightarrow \frac{\mathrm{init\_spdef} \cdot \mathrm{k\_ssc\_a\_spdef\_fc} \cdot \mathrm{p\_Spdef}}{\mathrm{d\_Spdef} \cdot \mathrm{reg\_spdef\_fc} \cdot \mathrm{s\_RNA}}$$

$$\text{k\_tp63\_a\_spdef} \to \frac{\text{init\_spdef} \cdot \text{k\_tp63\_a\_spdef\_fc} \cdot \text{p\_Spdef}}{\text{d\_Spdef} \cdot \text{reg\_spdef\_fc} \cdot \text{s\_RNA}}$$

$$\text{s\_cell1} \to \text{s\_cell1\_fc} \cdot \text{s\_cell\_mean}$$

$$\text{s\_cell12} \to \text{s\_cell12\_fc} \cdot \text{s\_cell\_mean}$$

$$\text{s\_cell13} \to \text{s\_cell13\_fc} \cdot \text{s\_cell\_mean}$$

$$\text{s\_cell14} \to \text{s\_cell14\_fc} \cdot \text{s\_cell\_mean}$$

$$\text{s\_cell2} \to \text{s\_cell2\_fc} \cdot \text{s\_cell\_mean}$$

$$\text{s\_cell20} \to \text{s\_cell20\_fc} \cdot \text{s\_cell\_mean}$$

$$\text{s\_cell22} \to \text{s\_cell22\_fc} \cdot \text{s\_cell\_mean}$$

$$\text{s\_cell23} \to \text{s\_cell23\_fc} \cdot \text{s\_cell\_mean}$$

$$\text{s\_cell24} \to \text{s\_cell24\_fc} \cdot \text{s\_cell\_mean}$$

$$\text{s\_cell25} \to \text{s\_cell25\_fc} \cdot \text{s\_cell\_mean}$$

$$\text{s\_cell3} \to \text{s\_cell3\_fc} \cdot \text{s\_cell\_mean}$$

$$\text{s\_cell4} \to \text{s\_cell4\_fc} \cdot \text{s\_cell\_mean}$$

$$\text{s\_cell5} \to \text{s\_cell5\_fc} \cdot \text{s\_cell\_mean}$$

$$\text{s\_cell8} \to \text{s\_cell8\_fc} \cdot \text{s\_cell\_mean}$$

$$\text{s\_cell9} \to \text{s\_cell9\_fc} \cdot \text{s\_cell\_mean}$$

$$\text{transl\_hes4\_lof\_fc} \to 1$$

$$\text{transl\_hes5\_lof\_fc} \to 1$$

$$\text{transl\_hes7\_lof\_fc} \to 1$$

- **Local condition #11 (global condition #1):**

$$\text{ex1} \to 0$$

$$\text{ex12} \to 0$$

$$\text{ex13} \to 0$$

$$\text{ex14} \to 0$$

$$\text{ex2} \to 0$$

$$\text{ex20} \to 0$$

$$\text{ex22} \to 0$$

$$\text{ex23} \to 1$$

$$\text{ex24} \to 0$$

$$\text{ex25} \to 0$$

$$\text{ex3} \to 0$$

$$\text{ex4} \to 0$$

$$\text{ex5} \to 0$$

$$\text{ex8} \to 0$$

$$\text{ex9} \to 0$$

$$\text{init\_hes4} \to \frac{\text{init\_hes4}}{\text{reg\_hes4\_fc} \cdot \text{s\_RNA}}$$

$$\text{init\_hes4\_d} \to \frac{4 \cdot \text{init\_hes4}}{\text{d\_hes4\_prot\_fc} \cdot \text{d\_hes\_prot} \cdot \text{hes\_delay} \cdot \text{reg\_hes4\_fc} \cdot \text{s\_RNA}}$$

$$\text{init\_hes4\_d1} \rightarrow \frac{\text{init\_hes4}}{\text{reg\_hes4\_fc} \cdot \text{s\_RNA}}$$
$$\text{init\_hes4\_d2} \rightarrow \frac{\text{init\_hes4}}{\text{reg\_hes4\_fc} \cdot \text{s\_RNA}}$$
$$\text{init\_hes4\_d3} \rightarrow \frac{\text{init\_hes4}}{\text{reg\_hes4\_fc} \cdot \text{s\_RNA}}$$
$$\text{init\_hes5} \rightarrow \frac{\text{init\_hes5}}{\text{reg\_hes5\_fc} \cdot \text{s\_RNA}}$$
$$\text{init\_hes5\_d} \rightarrow \frac{4 \cdot \text{init\_hes5}}{\text{d\_hes5\_prot\_fc} \cdot \text{d\_hes\_prot} \cdot \text{hes\_delay} \cdot \text{reg\_hes5\_fc} \cdot \text{s\_RNA}}$$
$$\text{init\_hes5\_d1} \rightarrow \frac{\text{init\_hes5}}{\text{reg\_hes5\_fc} \cdot \text{s\_RNA}}$$
$$\text{init\_hes5\_d2} \rightarrow \frac{\text{init\_hes5}}{\text{reg\_hes5\_fc} \cdot \text{s\_RNA}}$$
$$\text{init\_hes5\_d3} \rightarrow \frac{\text{init\_hes5}}{\text{reg\_hes5\_fc} \cdot \text{s\_RNA}}$$
$$\text{init\_hes7} \rightarrow \frac{\text{init\_hes7}}{\text{reg\_hes7\_fc} \cdot \text{s\_RNA}}$$
$$\text{init\_hes7\_d} \rightarrow \frac{4 \cdot \text{init\_hes7}}{\text{d\_hes7\_prot\_fc} \cdot \text{d\_hes\_prot} \cdot \text{hes\_delay} \cdot \text{reg\_hes7\_fc} \cdot \text{s\_RNA}}$$
$$\text{init\_hes7\_d1} \rightarrow \frac{\text{init\_hes7}}{\text{reg\_hes7\_fc} \cdot \text{s\_RNA}}$$
$$\text{init\_hes7\_d2} \rightarrow \frac{\text{init\_hes7}}{\text{reg\_hes7\_fc} \cdot \text{s\_RNA}}$$
$$\text{init\_hes7\_d3} \rightarrow \frac{\text{init\_hes7}}{\text{reg\_hes7\_fc} \cdot \text{s\_RNA}}$$
$$\text{init\_Spdef} \rightarrow \frac{\text{init\_spdef} \cdot \text{p\_Spdef}}{\text{d\_Spdef} \cdot \text{reg\_spdef\_fc} \cdot \text{s\_RNA}}$$
$$\text{init\_spdef} \rightarrow \frac{\text{init\_spdef}}{\text{reg\_spdef\_fc} \cdot \text{s\_RNA}}$$
$$\text{init\_tp63} \rightarrow \frac{\text{init\_tp63}}{\text{reg\_tp63\_fc} \cdot \text{s\_RNA}}$$
$$\text{k\_hes7\_i\_spdef} \rightarrow \frac{\text{init\_spdef} \cdot \text{k\_hes7\_i\_spdef\_fc} \cdot \text{p\_Spdef}}{\text{d\_Spdef} \cdot \text{reg\_spdef\_fc} \cdot \text{s\_RNA}}$$
$$\text{k\_ssc\_i\_spdef} \rightarrow \frac{\text{init\_spdef} \cdot \text{k\_ssc\_a\_spdef\_fc} \cdot \text{k\_ssc\_i\_spdef\_fc} \cdot \text{p\_Spdef}}{\text{d\_Spdef} \cdot \text{reg\_spdef\_fc} \cdot \text{s\_RNA}}$$
$$\text{k\_ssc\_a\_spdef} \rightarrow \frac{\text{init\_spdef} \cdot \text{k\_ssc\_a\_spdef\_fc} \cdot \text{p\_Spdef}}{\text{d\_Spdef} \cdot \text{reg\_spdef\_fc} \cdot \text{s\_RNA}}$$
$$\text{k\_tp63\_a\_spdef} \rightarrow \frac{\text{init\_spdef} \cdot \text{k\_tp63\_a\_spdef\_fc} \cdot \text{p\_Spdef}}{\text{d\_Spdef} \cdot \text{reg\_spdef\_fc} \cdot \text{s\_RNA}}$$
$$\text{s\_cell1} \rightarrow \text{s\_cell1\_fc} \cdot \text{s\_cell\_mean}$$
$$\text{s\_cell12} \rightarrow \text{s\_cell12\_fc} \cdot \text{s\_cell\_mean}$$
$$\text{s\_cell13} \rightarrow \text{s\_cell13\_fc} \cdot \text{s\_cell\_mean}$$
$$\text{s\_cell14} \rightarrow \text{s\_cell14\_fc} \cdot \text{s\_cell\_mean}$$
$$\text{s\_cell2} \rightarrow \text{s\_cell2\_fc} \cdot \text{s\_cell\_mean}$$

$$s\_cell20 \rightarrow s\_cell20\_fc \cdot s\_cell\_mean$$
$$s\_cell22 \rightarrow s\_cell22\_fc \cdot s\_cell\_mean$$
$$s\_cell23 \rightarrow s\_cell23\_fc \cdot s\_cell\_mean$$
$$s\_cell24 \rightarrow s\_cell24\_fc \cdot s\_cell\_mean$$
$$s\_cell25 \rightarrow s\_cell25\_fc \cdot s\_cell\_mean$$
$$s\_cell3 \rightarrow s\_cell3\_fc \cdot s\_cell\_mean$$
$$s\_cell4 \rightarrow s\_cell4\_fc \cdot s\_cell\_mean$$
$$s\_cell5 \rightarrow s\_cell5\_fc \cdot s\_cell\_mean$$
$$s\_cell8 \rightarrow s\_cell8\_fc \cdot s\_cell\_mean$$
$$s\_cell9 \rightarrow s\_cell9\_fc \cdot s\_cell\_mean$$
$$transl\_hes4\_lof\_fc \rightarrow 1$$
$$transl\_hes5\_lof\_fc \rightarrow 1$$
$$transl\_hes7\_lof\_fc \rightarrow 1$$

- **Local condition #12 (global condition #1):**

$$ex1 \rightarrow 0$$
$$ex12 \rightarrow 0$$
$$ex13 \rightarrow 0$$
$$ex14 \rightarrow 0$$
$$ex2 \rightarrow 0$$
$$ex20 \rightarrow 0$$
$$ex22 \rightarrow 1$$
$$ex23 \rightarrow 0$$
$$ex24 \rightarrow 0$$
$$ex25 \rightarrow 0$$
$$ex3 \rightarrow 0$$
$$ex4 \rightarrow 0$$
$$ex5 \rightarrow 0$$
$$ex8 \rightarrow 0$$
$$ex9 \rightarrow 0$$
$$init\_hes4 \rightarrow \frac{init\_hes4}{reg\_hes4\_fc \cdot s\_RNA}$$
$$init\_hes4\_d \rightarrow \frac{4 \cdot init\_hes4}{d\_hes4\_prot\_fc \cdot d\_hes\_prot \cdot hes\_delay \cdot reg\_hes4\_fc \cdot s\_RNA}$$
$$init\_hes4\_d1 \rightarrow \frac{init\_hes4}{reg\_hes4\_fc \cdot s\_RNA}$$
$$init\_hes4\_d2 \rightarrow \frac{init\_hes4}{reg\_hes4\_fc \cdot s\_RNA}$$
$$init\_hes4\_d3 \rightarrow \frac{init\_hes4}{reg\_hes4\_fc \cdot s\_RNA}$$
$$init\_hes5 \rightarrow \frac{init\_hes5}{reg\_hes5\_fc \cdot s\_RNA}$$

$$\text{init\_hes5\_d} \to \frac{4 \cdot \text{init\_hes5}}{\text{d\_hes5\_prot\_fc} \cdot \text{d\_hes\_prot} \cdot \text{hes\_delay} \cdot \text{reg\_hes5\_fc} \cdot \text{s\_RNA}}$$

$$\text{init\_hes5\_d1} \to \frac{\text{init\_hes5}}{\text{reg\_hes5\_fc} \cdot \text{s\_RNA}}$$

$$\text{init\_hes5\_d2} \to \frac{\text{init\_hes5}}{\text{reg\_hes5\_fc} \cdot \text{s\_RNA}}$$

$$\text{init\_hes5\_d3} \to \frac{\text{init\_hes5}}{\text{reg\_hes5\_fc} \cdot \text{s\_RNA}}$$

$$\text{init\_hes7} \to \frac{\text{init\_hes7}}{\text{reg\_hes7\_fc} \cdot \text{s\_RNA}}$$

$$\text{init\_hes7\_d} \to \frac{4 \cdot \text{init\_hes7}}{\text{d\_hes7\_prot\_fc} \cdot \text{d\_hes\_prot} \cdot \text{hes\_delay} \cdot \text{reg\_hes7\_fc} \cdot \text{s\_RNA}}$$

$$\text{init\_hes7\_d1} \to \frac{\text{init\_hes7}}{\text{reg\_hes7\_fc} \cdot \text{s\_RNA}}$$

$$\text{init\_hes7\_d2} \to \frac{\text{init\_hes7}}{\text{reg\_hes7\_fc} \cdot \text{s\_RNA}}$$

$$\text{init\_hes7\_d3} \to \frac{\text{init\_hes7}}{\text{reg\_hes7\_fc} \cdot \text{s\_RNA}}$$

$$\text{init\_Spdef} \to \frac{\text{init\_spdef} \cdot \text{p\_Spdef}}{\text{d\_Spdef} \cdot \text{reg\_spdef\_fc} \cdot \text{s\_RNA}}$$

$$\text{init\_spdef} \to \frac{\text{init\_spdef}}{\text{reg\_spdef\_fc} \cdot \text{s\_RNA}}$$

$$\text{init\_tp63} \to \frac{\text{init\_tp63}}{\text{reg\_tp63\_fc} \cdot \text{s\_RNA}}$$

$$\text{k\_hes7\_i\_spdef} \to \frac{\text{init\_spdef} \cdot \text{k\_hes7\_i\_spdef\_fc} \cdot \text{p\_Spdef}}{\text{d\_Spdef} \cdot \text{reg\_spdef\_fc} \cdot \text{s\_RNA}}$$

$$\text{k\_ssc\_i\_spdef} \to \frac{\text{init\_spdef} \cdot \text{k\_ssc\_a\_spdef\_fc} \cdot \text{k\_ssc\_i\_spdef\_fc} \cdot \text{p\_Spdef}}{\text{d\_Spdef} \cdot \text{reg\_spdef\_fc} \cdot \text{s\_RNA}}$$

$$\text{k\_ssc\_a\_spdef} \to \frac{\text{init\_spdef} \cdot \text{k\_ssc\_a\_spdef\_fc} \cdot \text{p\_Spdef}}{\text{d\_Spdef} \cdot \text{reg\_spdef\_fc} \cdot \text{s\_RNA}}$$

$$\text{k\_tp63\_a\_spdef} \to \frac{\text{init\_spdef} \cdot \text{k\_tp63\_a\_spdef\_fc} \cdot \text{p\_Spdef}}{\text{d\_Spdef} \cdot \text{reg\_spdef\_fc} \cdot \text{s\_RNA}}$$

$$\text{s\_cell1} \to \text{s\_cell1\_fc} \cdot \text{s\_cell\_mean}$$

$$\text{s\_cell12} \to \text{s\_cell12\_fc} \cdot \text{s\_cell\_mean}$$

$$\text{s\_cell13} \to \text{s\_cell13\_fc} \cdot \text{s\_cell\_mean}$$

$$\text{s\_cell14} \to \text{s\_cell14\_fc} \cdot \text{s\_cell\_mean}$$

$$\text{s\_cell2} \to \text{s\_cell2\_fc} \cdot \text{s\_cell\_mean}$$

$$\text{s\_cell20} \to \text{s\_cell20\_fc} \cdot \text{s\_cell\_mean}$$

$$\text{s\_cell22} \to \text{s\_cell22\_fc} \cdot \text{s\_cell\_mean}$$

$$\text{s\_cell23} \to \text{s\_cell23\_fc} \cdot \text{s\_cell\_mean}$$

$$\text{s\_cell24} \to \text{s\_cell24\_fc} \cdot \text{s\_cell\_mean}$$

$$\text{s\_cell25} \to \text{s\_cell25\_fc} \cdot \text{s\_cell\_mean}$$

$$\text{s\_cell3} \to \text{s\_cell3\_fc} \cdot \text{s\_cell\_mean}$$

$$\text{s\_cell4} \to \text{s\_cell4\_fc} \cdot \text{s\_cell\_mean}$$

$$\mathrm{s\_cell5} \rightarrow \mathrm{s\_cell5\_fc} \cdot \mathrm{s\_cell\_mean}$$
$$\mathrm{s\_cell8} \rightarrow \mathrm{s\_cell8\_fc} \cdot \mathrm{s\_cell\_mean}$$
$$\mathrm{s\_cell9} \rightarrow \mathrm{s\_cell9\_fc} \cdot \mathrm{s\_cell\_mean}$$
$$\mathrm{transl\_hes4\_lof\_fc} \rightarrow 1$$
$$\mathrm{transl\_hes5\_lof\_fc} \rightarrow 1$$
$$\mathrm{transl\_hes7\_lof\_fc} \rightarrow 1$$

- **Local condition #13 (global condition #1):**

$$\mathrm{ex1} \rightarrow 0$$
$$\mathrm{ex12} \rightarrow 0$$
$$\mathrm{ex13} \rightarrow 0$$
$$\mathrm{ex14} \rightarrow 0$$
$$\mathrm{ex2} \rightarrow 0$$
$$\mathrm{ex20} \rightarrow 1$$
$$\mathrm{ex22} \rightarrow 0$$
$$\mathrm{ex23} \rightarrow 0$$
$$\mathrm{ex24} \rightarrow 0$$
$$\mathrm{ex25} \rightarrow 0$$
$$\mathrm{ex3} \rightarrow 0$$
$$\mathrm{ex4} \rightarrow 0$$
$$\mathrm{ex5} \rightarrow 0$$
$$\mathrm{ex8} \rightarrow 0$$
$$\mathrm{ex9} \rightarrow 0$$
$$\mathrm{init\_hes4} \rightarrow \frac{\mathrm{init\_hes4}}{\mathrm{reg\_hes4\_fc} \cdot \mathrm{s\_RNA}}$$
$$\mathrm{init\_hes4\_d} \rightarrow \frac{4 \cdot \mathrm{init\_hes4}}{\mathrm{d\_hes4\_prot\_fc} \cdot \mathrm{d\_hes\_prot} \cdot \mathrm{hes\_delay} \cdot \mathrm{reg\_hes4\_fc} \cdot \mathrm{s\_RNA}}$$
$$\mathrm{init\_hes4\_d1} \rightarrow \frac{\mathrm{init\_hes4}}{\mathrm{reg\_hes4\_fc} \cdot \mathrm{s\_RNA}}$$
$$\mathrm{init\_hes4\_d2} \rightarrow \frac{\mathrm{init\_hes4}}{\mathrm{reg\_hes4\_fc} \cdot \mathrm{s\_RNA}}$$
$$\mathrm{init\_hes4\_d3} \rightarrow \frac{\mathrm{init\_hes4}}{\mathrm{reg\_hes4\_fc} \cdot \mathrm{s\_RNA}}$$
$$\mathrm{init\_hes5} \rightarrow \frac{\mathrm{init\_hes5}}{\mathrm{reg\_hes5\_fc} \cdot \mathrm{s\_RNA}}$$
$$\mathrm{init\_hes5\_d} \rightarrow \frac{4 \cdot \mathrm{init\_hes5}}{\mathrm{d\_hes5\_prot\_fc} \cdot \mathrm{d\_hes\_prot} \cdot \mathrm{hes\_delay} \cdot \mathrm{reg\_hes5\_fc} \cdot \mathrm{s\_RNA}}$$
$$\mathrm{init\_hes5\_d1} \rightarrow \frac{\mathrm{init\_hes5}}{\mathrm{reg\_hes5\_fc} \cdot \mathrm{s\_RNA}}$$
$$\mathrm{init\_hes5\_d2} \rightarrow \frac{\mathrm{init\_hes5}}{\mathrm{reg\_hes5\_fc} \cdot \mathrm{s\_RNA}}$$
$$\mathrm{init\_hes5\_d3} \rightarrow \frac{\mathrm{init\_hes5}}{\mathrm{reg\_hes5\_fc} \cdot \mathrm{s\_RNA}}$$

$$\mathrm{init\_hes7} \rightarrow \frac{\mathrm{init\_hes7}}{\mathrm{reg\_hes7\_fc} \cdot \mathrm{s\_RNA}}$$
$$\mathrm{init\_hes7\_d} \rightarrow \frac{4 \cdot \mathrm{init\_hes7}}{\mathrm{d\_hes7\_prot\_fc} \cdot \mathrm{d\_hes\_prot} \cdot \mathrm{hes\_delay} \cdot \mathrm{reg\_hes7\_fc} \cdot \mathrm{s\_RNA}}$$
$$\mathrm{init\_hes7\_d1} \rightarrow \frac{\mathrm{init\_hes7}}{\mathrm{reg\_hes7\_fc} \cdot \mathrm{s\_RNA}}$$
$$\mathrm{init\_hes7\_d2} \rightarrow \frac{\mathrm{init\_hes7}}{\mathrm{reg\_hes7\_fc} \cdot \mathrm{s\_RNA}}$$
$$\mathrm{init\_hes7\_d3} \rightarrow \frac{\mathrm{init\_hes7}}{\mathrm{reg\_hes7\_fc} \cdot \mathrm{s\_RNA}}$$
$$\mathrm{init\_Spdef} \rightarrow \frac{\mathrm{init\_spdef} \cdot \mathrm{p\_Spdef}}{\mathrm{d\_Spdef} \cdot \mathrm{reg\_spdef\_fc} \cdot \mathrm{s\_RNA}}$$
$$\mathrm{init\_spdef} \rightarrow \frac{\mathrm{init\_spdef}}{\mathrm{reg\_spdef\_fc} \cdot \mathrm{s\_RNA}}$$
$$\mathrm{init\_tp63} \rightarrow \frac{\mathrm{init\_tp63}}{\mathrm{reg\_tp63\_fc} \cdot \mathrm{s\_RNA}}$$
$$\mathrm{k\_hes7\_i\_spdef} \rightarrow \frac{\mathrm{init\_spdef} \cdot \mathrm{k\_hes7\_i\_spdef\_fc} \cdot \mathrm{p\_Spdef}}{\mathrm{d\_Spdef} \cdot \mathrm{reg\_spdef\_fc} \cdot \mathrm{s\_RNA}}$$
$$\mathrm{k\_ssc\_i\_spdef} \rightarrow \frac{\mathrm{init\_spdef} \cdot \mathrm{k\_ssc\_a\_spdef\_fc} \cdot \mathrm{k\_ssc\_i\_spdef\_fc} \cdot \mathrm{p\_Spdef}}{\mathrm{d\_Spdef} \cdot \mathrm{reg\_spdef\_fc} \cdot \mathrm{s\_RNA}}$$
$$\mathrm{k\_ssc\_a\_spdef} \rightarrow \frac{\mathrm{init\_spdef} \cdot \mathrm{k\_ssc\_a\_spdef\_fc} \cdot \mathrm{p\_Spdef}}{\mathrm{d\_Spdef} \cdot \mathrm{reg\_spdef\_fc} \cdot \mathrm{s\_RNA}}$$
$$\mathrm{k\_tp63\_a\_spdef} \rightarrow \frac{\mathrm{init\_spdef} \cdot \mathrm{k\_tp63\_a\_spdef\_fc} \cdot \mathrm{p\_Spdef}}{\mathrm{d\_Spdef} \cdot \mathrm{reg\_spdef\_fc} \cdot \mathrm{s\_RNA}}$$
$$\mathrm{s\_cell1} \rightarrow \mathrm{s\_cell1\_fc} \cdot \mathrm{s\_cell\_mean}$$
$$\mathrm{s\_cell12} \rightarrow \mathrm{s\_cell12\_fc} \cdot \mathrm{s\_cell\_mean}$$
$$\mathrm{s\_cell13} \rightarrow \mathrm{s\_cell13\_fc} \cdot \mathrm{s\_cell\_mean}$$
$$\mathrm{s\_cell14} \rightarrow \mathrm{s\_cell14\_fc} \cdot \mathrm{s\_cell\_mean}$$
$$\mathrm{s\_cell2} \rightarrow \mathrm{s\_cell2\_fc} \cdot \mathrm{s\_cell\_mean}$$
$$\mathrm{s\_cell20} \rightarrow \mathrm{s\_cell20\_fc} \cdot \mathrm{s\_cell\_mean}$$
$$\mathrm{s\_cell22} \rightarrow \mathrm{s\_cell22\_fc} \cdot \mathrm{s\_cell\_mean}$$
$$\mathrm{s\_cell23} \rightarrow \mathrm{s\_cell23\_fc} \cdot \mathrm{s\_cell\_mean}$$
$$\mathrm{s\_cell24} \rightarrow \mathrm{s\_cell24\_fc} \cdot \mathrm{s\_cell\_mean}$$
$$\mathrm{s\_cell25} \rightarrow \mathrm{s\_cell25\_fc} \cdot \mathrm{s\_cell\_mean}$$
$$\mathrm{s\_cell3} \rightarrow \mathrm{s\_cell3\_fc} \cdot \mathrm{s\_cell\_mean}$$
$$\mathrm{s\_cell4} \rightarrow \mathrm{s\_cell4\_fc} \cdot \mathrm{s\_cell\_mean}$$
$$\mathrm{s\_cell5} \rightarrow \mathrm{s\_cell5\_fc} \cdot \mathrm{s\_cell\_mean}$$
$$\mathrm{s\_cell8} \rightarrow \mathrm{s\_cell8\_fc} \cdot \mathrm{s\_cell\_mean}$$
$$\mathrm{s\_cell9} \rightarrow \mathrm{s\_cell9\_fc} \cdot \mathrm{s\_cell\_mean}$$
$$\mathrm{transl\_hes4\_lof\_fc} \rightarrow 1$$
$$\mathrm{transl\_hes5\_lof\_fc} \rightarrow 1$$
$$\mathrm{transl\_hes7\_lof\_fc} \rightarrow 1$$

- **Local condition #14 (global condition #1):**

$$
\begin{aligned}
\mathrm{ex1} &\to 0\\
\mathrm{ex12} &\to 0\\
\mathrm{ex13} &\to 0\\
\mathrm{ex14} &\to 1\\
\mathrm{ex2} &\to 0\\
\mathrm{ex20} &\to 0\\
\mathrm{ex22} &\to 0\\
\mathrm{ex23} &\to 0\\
\mathrm{ex24} &\to 0\\
\mathrm{ex25} &\to 0\\
\mathrm{ex3} &\to 0\\
\mathrm{ex4} &\to 0\\
\mathrm{ex5} &\to 0\\
\mathrm{ex8} &\to 0\\
\mathrm{ex9} &\to 0\\
\mathrm{init\_hes4} &\to \frac{\mathrm{init\_hes4}}{\mathrm{reg\_hes4\_fc} \cdot \mathrm{s\_RNA}}\\
\mathrm{init\_hes4\_d} &\to \frac{4 \cdot \mathrm{init\_hes4}}{\mathrm{d\_hes4\_prot\_fc} \cdot \mathrm{d\_hes\_prot} \cdot \mathrm{hes\_delay} \cdot \mathrm{reg\_hes4\_fc} \cdot \mathrm{s\_RNA}}\\
\mathrm{init\_hes4\_d1} &\to \frac{\mathrm{init\_hes4}}{\mathrm{reg\_hes4\_fc} \cdot \mathrm{s\_RNA}}\\
\mathrm{init\_hes4\_d2} &\to \frac{\mathrm{init\_hes4}}{\mathrm{reg\_hes4\_fc} \cdot \mathrm{s\_RNA}}\\
\mathrm{init\_hes4\_d3} &\to \frac{\mathrm{init\_hes4}}{\mathrm{reg\_hes4\_fc} \cdot \mathrm{s\_RNA}}\\
\mathrm{init\_hes5} &\to \frac{\mathrm{init\_hes5}}{\mathrm{reg\_hes5\_fc} \cdot \mathrm{s\_RNA}}\\
\mathrm{init\_hes5\_d} &\to \frac{4 \cdot \mathrm{init\_hes5}}{\mathrm{d\_hes5\_prot\_fc} \cdot \mathrm{d\_hes\_prot} \cdot \mathrm{hes\_delay} \cdot \mathrm{reg\_hes5\_fc} \cdot \mathrm{s\_RNA}}\\
\mathrm{init\_hes5\_d1} &\to \frac{\mathrm{init\_hes5}}{\mathrm{reg\_hes5\_fc} \cdot \mathrm{s\_RNA}}\\
\mathrm{init\_hes5\_d2} &\to \frac{\mathrm{init\_hes5}}{\mathrm{reg\_hes5\_fc} \cdot \mathrm{s\_RNA}}\\
\mathrm{init\_hes5\_d3} &\to \frac{\mathrm{init\_hes5}}{\mathrm{reg\_hes5\_fc} \cdot \mathrm{s\_RNA}}\\
\mathrm{init\_hes7} &\to \frac{\mathrm{init\_hes7}}{\mathrm{reg\_hes7\_fc} \cdot \mathrm{s\_RNA}}\\
\mathrm{init\_hes7\_d} &\to \frac{4 \cdot \mathrm{init\_hes7}}{\mathrm{d\_hes7\_prot\_fc} \cdot \mathrm{d\_hes\_prot} \cdot \mathrm{hes\_delay} \cdot \mathrm{reg\_hes7\_fc} \cdot \mathrm{s\_RNA}}\\
\mathrm{init\_hes7\_d1} &\to \frac{\mathrm{init\_hes7}}{\mathrm{reg\_hes7\_fc} \cdot \mathrm{s\_RNA}}\\
\mathrm{init\_hes7\_d2} &\to \frac{\mathrm{init\_hes7}}{\mathrm{reg\_hes7\_fc} \cdot \mathrm{s\_RNA}}
\end{aligned}
$$

$$\text{init\_hes7\_d3} \rightarrow \frac{\text{init\_hes7}}{\text{reg\_hes7\_fc} \cdot \text{s\_RNA}}$$

$$\text{init\_Spdef} \rightarrow \frac{\text{init\_spdef} \cdot \text{p\_Spdef}}{\text{d\_Spdef} \cdot \text{reg\_spdef\_fc} \cdot \text{s\_RNA}}$$

$$\text{init\_spdef} \rightarrow \frac{\text{init\_spdef}}{\text{reg\_spdef\_fc} \cdot \text{s\_RNA}}$$

$$\text{init\_tp63} \rightarrow \frac{\text{init\_tp63}}{\text{reg\_tp63\_fc} \cdot \text{s\_RNA}}$$

$$\text{k\_hes7\_i\_spdef} \rightarrow \frac{\text{init\_spdef} \cdot \text{k\_hes7\_i\_spdef\_fc} \cdot \text{p\_Spdef}}{\text{d\_Spdef} \cdot \text{reg\_spdef\_fc} \cdot \text{s\_RNA}}$$

$$\text{k\_ssc\_i\_spdef} \rightarrow \frac{\text{init\_spdef} \cdot \text{k\_ssc\_a\_spdef\_fc} \cdot \text{k\_ssc\_i\_spdef\_fc} \cdot \text{p\_Spdef}}{\text{d\_Spdef} \cdot \text{reg\_spdef\_fc} \cdot \text{s\_RNA}}$$

$$\text{k\_ssc\_a\_spdef} \rightarrow \frac{\text{init\_spdef} \cdot \text{k\_ssc\_a\_spdef\_fc} \cdot \text{p\_Spdef}}{\text{d\_Spdef} \cdot \text{reg\_spdef\_fc} \cdot \text{s\_RNA}}$$

$$\text{k\_tp63\_a\_spdef} \rightarrow \frac{\text{init\_spdef} \cdot \text{k\_tp63\_a\_spdef\_fc} \cdot \text{p\_Spdef}}{\text{d\_Spdef} \cdot \text{reg\_spdef\_fc} \cdot \text{s\_RNA}}$$

$$\text{s\_cell1} \rightarrow \text{s\_cell1\_fc} \cdot \text{s\_cell\_mean}$$

$$\text{s\_cell12} \rightarrow \text{s\_cell12\_fc} \cdot \text{s\_cell\_mean}$$

$$\text{s\_cell13} \rightarrow \text{s\_cell13\_fc} \cdot \text{s\_cell\_mean}$$

$$\text{s\_cell14} \rightarrow \text{s\_cell14\_fc} \cdot \text{s\_cell\_mean}$$

$$\text{s\_cell2} \rightarrow \text{s\_cell2\_fc} \cdot \text{s\_cell\_mean}$$

$$\text{s\_cell20} \rightarrow \text{s\_cell20\_fc} \cdot \text{s\_cell\_mean}$$

$$\text{s\_cell22} \rightarrow \text{s\_cell22\_fc} \cdot \text{s\_cell\_mean}$$

$$\text{s\_cell23} \rightarrow \text{s\_cell23\_fc} \cdot \text{s\_cell\_mean}$$

$$\text{s\_cell24} \rightarrow \text{s\_cell24\_fc} \cdot \text{s\_cell\_mean}$$

$$\text{s\_cell25} \rightarrow \text{s\_cell25\_fc} \cdot \text{s\_cell\_mean}$$

$$\text{s\_cell3} \rightarrow \text{s\_cell3\_fc} \cdot \text{s\_cell\_mean}$$

$$\text{s\_cell4} \rightarrow \text{s\_cell4\_fc} \cdot \text{s\_cell\_mean}$$

$$\text{s\_cell5} \rightarrow \text{s\_cell5\_fc} \cdot \text{s\_cell\_mean}$$

$$\text{s\_cell8} \rightarrow \text{s\_cell8\_fc} \cdot \text{s\_cell\_mean}$$

$$\text{s\_cell9} \rightarrow \text{s\_cell9\_fc} \cdot \text{s\_cell\_mean}$$

$$\text{transl\_hes4\_lof\_fc} \rightarrow 1$$

$$\text{transl\_hes5\_lof\_fc} \rightarrow 1$$

$$\text{transl\_hes7\_lof\_fc} \rightarrow 1$$

- **Local condition #15 (global condition #1):**

$$\text{ex1} \rightarrow 0$$

$$\text{ex12} \rightarrow 0$$

$$\text{ex13} \rightarrow 1$$

$$\text{ex14} \rightarrow 0$$

$$\text{ex2} \rightarrow 0$$

$$\text{ex20} \rightarrow 0$$

$$\mathrm{ex22} \rightarrow 0$$
$$\mathrm{ex23} \rightarrow 0$$
$$\mathrm{ex24} \rightarrow 0$$
$$\mathrm{ex25} \rightarrow 0$$
$$\mathrm{ex3} \rightarrow 0$$
$$\mathrm{ex4} \rightarrow 0$$
$$\mathrm{ex5} \rightarrow 0$$
$$\mathrm{ex8} \rightarrow 0$$
$$\mathrm{ex9} \rightarrow 0$$
$$\mathrm{init\_hes4} \rightarrow \frac{\mathrm{init\_hes4}}{\mathrm{reg\_hes4\_fc} \cdot \mathrm{s\_RNA}}$$
$$\mathrm{init\_hes4\_d} \rightarrow \frac{4 \cdot \mathrm{init\_hes4}}{\mathrm{d\_hes4\_prot\_fc} \cdot \mathrm{d\_hes\_prot} \cdot \mathrm{hes\_delay} \cdot \mathrm{reg\_hes4\_fc} \cdot \mathrm{s\_RNA}}$$
$$\mathrm{init\_hes4\_d1} \rightarrow \frac{\mathrm{init\_hes4}}{\mathrm{reg\_hes4\_fc} \cdot \mathrm{s\_RNA}}$$
$$\mathrm{init\_hes4\_d2} \rightarrow \frac{\mathrm{init\_hes4}}{\mathrm{reg\_hes4\_fc} \cdot \mathrm{s\_RNA}}$$
$$\mathrm{init\_hes4\_d3} \rightarrow \frac{\mathrm{init\_hes4}}{\mathrm{reg\_hes4\_fc} \cdot \mathrm{s\_RNA}}$$
$$\mathrm{init\_hes5} \rightarrow \frac{\mathrm{init\_hes5}}{\mathrm{reg\_hes5\_fc} \cdot \mathrm{s\_RNA}}$$
$$\mathrm{init\_hes5\_d} \rightarrow \frac{4 \cdot \mathrm{init\_hes5}}{\mathrm{d\_hes5\_prot\_fc} \cdot \mathrm{d\_hes\_prot} \cdot \mathrm{hes\_delay} \cdot \mathrm{reg\_hes5\_fc} \cdot \mathrm{s\_RNA}}$$
$$\mathrm{init\_hes5\_d1} \rightarrow \frac{\mathrm{init\_hes5}}{\mathrm{reg\_hes5\_fc} \cdot \mathrm{s\_RNA}}$$
$$\mathrm{init\_hes5\_d2} \rightarrow \frac{\mathrm{init\_hes5}}{\mathrm{reg\_hes5\_fc} \cdot \mathrm{s\_RNA}}$$
$$\mathrm{init\_hes5\_d3} \rightarrow \frac{\mathrm{init\_hes5}}{\mathrm{reg\_hes5\_fc} \cdot \mathrm{s\_RNA}}$$
$$\mathrm{init\_hes7} \rightarrow \frac{\mathrm{init\_hes7}}{\mathrm{reg\_hes7\_fc} \cdot \mathrm{s\_RNA}}$$
$$\mathrm{init\_hes7\_d} \rightarrow \frac{4 \cdot \mathrm{init\_hes7}}{\mathrm{d\_hes7\_prot\_fc} \cdot \mathrm{d\_hes\_prot} \cdot \mathrm{hes\_delay} \cdot \mathrm{reg\_hes7\_fc} \cdot \mathrm{s\_RNA}}$$
$$\mathrm{init\_hes7\_d1} \rightarrow \frac{\mathrm{init\_hes7}}{\mathrm{reg\_hes7\_fc} \cdot \mathrm{s\_RNA}}$$
$$\mathrm{init\_hes7\_d2} \rightarrow \frac{\mathrm{init\_hes7}}{\mathrm{reg\_hes7\_fc} \cdot \mathrm{s\_RNA}}$$
$$\mathrm{init\_hes7\_d3} \rightarrow \frac{\mathrm{init\_hes7}}{\mathrm{reg\_hes7\_fc} \cdot \mathrm{s\_RNA}}$$
$$\mathrm{init\_Spdef} \rightarrow \frac{\mathrm{init\_spdef} \cdot \mathrm{p\_Spdef}}{\mathrm{d\_Spdef} \cdot \mathrm{reg\_spdef\_fc} \cdot \mathrm{s\_RNA}}$$
$$\mathrm{init\_spdef} \rightarrow \frac{\mathrm{init\_spdef}}{\mathrm{reg\_spdef\_fc} \cdot \mathrm{s\_RNA}}$$

$$\text{init\_tp63} \rightarrow \frac{\text{init\_tp63}}{\text{reg\_tp63\_fc} \cdot \text{s\_RNA}}$$

$$\text{k\_hes7\_i\_spdef} \rightarrow \frac{\text{init\_spdef} \cdot \text{k\_hes7\_i\_spdef\_fc} \cdot \text{p\_Spdef}}{\text{d\_Spdef} \cdot \text{reg\_spdef\_fc} \cdot \text{s\_RNA}}$$

$$\text{k\_ssc\_i\_spdef} \rightarrow \frac{\text{init\_spdef} \cdot \text{k\_ssc\_a\_spdef\_fc} \cdot \text{k\_ssc\_i\_spdef\_fc} \cdot \text{p\_Spdef}}{\text{d\_Spdef} \cdot \text{reg\_spdef\_fc} \cdot \text{s\_RNA}}$$

$$\text{k\_ssc\_a\_spdef} \rightarrow \frac{\text{init\_spdef} \cdot \text{k\_ssc\_a\_spdef\_fc} \cdot \text{p\_Spdef}}{\text{d\_Spdef} \cdot \text{reg\_spdef\_fc} \cdot \text{s\_RNA}}$$

$$\text{k\_tp63\_a\_spdef} \rightarrow \frac{\text{init\_spdef} \cdot \text{k\_tp63\_a\_spdef\_fc} \cdot \text{p\_Spdef}}{\text{d\_Spdef} \cdot \text{reg\_spdef\_fc} \cdot \text{s\_RNA}}$$

$$\text{s\_cell1} \rightarrow \text{s\_cell1\_fc} \cdot \text{s\_cell\_mean}$$

$$\text{s\_cell12} \rightarrow \text{s\_cell12\_fc} \cdot \text{s\_cell\_mean}$$

$$\text{s\_cell13} \rightarrow \text{s\_cell13\_fc} \cdot \text{s\_cell\_mean}$$

$$\text{s\_cell14} \rightarrow \text{s\_cell14\_fc} \cdot \text{s\_cell\_mean}$$

$$\text{s\_cell2} \rightarrow \text{s\_cell2\_fc} \cdot \text{s\_cell\_mean}$$

$$\text{s\_cell20} \rightarrow \text{s\_cell20\_fc} \cdot \text{s\_cell\_mean}$$

$$\text{s\_cell22} \rightarrow \text{s\_cell22\_fc} \cdot \text{s\_cell\_mean}$$

$$\text{s\_cell23} \rightarrow \text{s\_cell23\_fc} \cdot \text{s\_cell\_mean}$$

$$\text{s\_cell24} \rightarrow \text{s\_cell24\_fc} \cdot \text{s\_cell\_mean}$$

$$\text{s\_cell25} \rightarrow \text{s\_cell25\_fc} \cdot \text{s\_cell\_mean}$$

$$\text{s\_cell3} \rightarrow \text{s\_cell3\_fc} \cdot \text{s\_cell\_mean}$$

$$\text{s\_cell4} \rightarrow \text{s\_cell4\_fc} \cdot \text{s\_cell\_mean}$$

$$\text{s\_cell5} \rightarrow \text{s\_cell5\_fc} \cdot \text{s\_cell\_mean}$$

$$\text{s\_cell8} \rightarrow \text{s\_cell8\_fc} \cdot \text{s\_cell\_mean}$$

$$\text{s\_cell9} \rightarrow \text{s\_cell9\_fc} \cdot \text{s\_cell\_mean}$$

$$\text{transl\_hes4\_lof\_fc} \rightarrow 1$$

$$\text{transl\_hes5\_lof\_fc} \rightarrow 1$$

$$\text{transl\_hes7\_lof\_fc} \rightarrow 1$$

- **Local condition #16 (global condition #1):**

$$\text{ex1} \rightarrow 0$$

$$\text{ex12} \rightarrow 1$$

$$\text{ex13} \rightarrow 0$$

$$\text{ex14} \rightarrow 0$$

$$\text{ex2} \rightarrow 0$$

$$\text{ex20} \rightarrow 0$$

$$\text{ex22} \rightarrow 0$$

$$\text{ex23} \rightarrow 0$$

$$\text{ex24} \rightarrow 0$$

$$\text{ex25} \rightarrow 0$$

$$\text{ex3} \rightarrow 0$$

$$\text{ex4} \rightarrow 0$$

$$\mathrm{ex5} \rightarrow 0$$

$$\mathrm{ex8} \rightarrow 0$$

$$\mathrm{ex9} \rightarrow 0$$

$$\mathrm{init\_hes4} \rightarrow \frac{\mathrm{init\_hes4}}{\mathrm{reg\_hes4\_fc} \cdot \mathrm{s\_RNA}}$$

$$\mathrm{init\_hes4\_d} \rightarrow \frac{4 \cdot \mathrm{init\_hes4}}{\mathrm{d\_hes4\_prot\_fc} \cdot \mathrm{d\_hes\_prot} \cdot \mathrm{hes\_delay} \cdot \mathrm{reg\_hes4\_fc} \cdot \mathrm{s\_RNA}}$$

$$\mathrm{init\_hes4\_d1} \rightarrow \frac{\mathrm{init\_hes4}}{\mathrm{reg\_hes4\_fc} \cdot \mathrm{s\_RNA}}$$

$$\mathrm{init\_hes4\_d2} \rightarrow \frac{\mathrm{init\_hes4}}{\mathrm{reg\_hes4\_fc} \cdot \mathrm{s\_RNA}}$$

$$\mathrm{init\_hes4\_d3} \rightarrow \frac{\mathrm{init\_hes4}}{\mathrm{reg\_hes4\_fc} \cdot \mathrm{s\_RNA}}$$

$$\mathrm{init\_hes5} \rightarrow \frac{\mathrm{init\_hes5}}{\mathrm{reg\_hes5\_fc} \cdot \mathrm{s\_RNA}}$$

$$\mathrm{init\_hes5\_d} \rightarrow \frac{4 \cdot \mathrm{init\_hes5}}{\mathrm{d\_hes5\_prot\_fc} \cdot \mathrm{d\_hes\_prot} \cdot \mathrm{hes\_delay} \cdot \mathrm{reg\_hes5\_fc} \cdot \mathrm{s\_RNA}}$$

$$\mathrm{init\_hes5\_d1} \rightarrow \frac{\mathrm{init\_hes5}}{\mathrm{reg\_hes5\_fc} \cdot \mathrm{s\_RNA}}$$

$$\mathrm{init\_hes5\_d2} \rightarrow \frac{\mathrm{init\_hes5}}{\mathrm{reg\_hes5\_fc} \cdot \mathrm{s\_RNA}}$$

$$\mathrm{init\_hes5\_d3} \rightarrow \frac{\mathrm{init\_hes5}}{\mathrm{reg\_hes5\_fc} \cdot \mathrm{s\_RNA}}$$

$$\mathrm{init\_hes7} \rightarrow \frac{\mathrm{init\_hes7}}{\mathrm{reg\_hes7\_fc} \cdot \mathrm{s\_RNA}}$$

$$\mathrm{init\_hes7\_d} \rightarrow \frac{4 \cdot \mathrm{init\_hes7}}{\mathrm{d\_hes7\_prot\_fc} \cdot \mathrm{d\_hes\_prot} \cdot \mathrm{hes\_delay} \cdot \mathrm{reg\_hes7\_fc} \cdot \mathrm{s\_RNA}}$$

$$\mathrm{init\_hes7\_d1} \rightarrow \frac{\mathrm{init\_hes7}}{\mathrm{reg\_hes7\_fc} \cdot \mathrm{s\_RNA}}$$

$$\mathrm{init\_hes7\_d2} \rightarrow \frac{\mathrm{init\_hes7}}{\mathrm{reg\_hes7\_fc} \cdot \mathrm{s\_RNA}}$$

$$\mathrm{init\_hes7\_d3} \rightarrow \frac{\mathrm{init\_hes7}}{\mathrm{reg\_hes7\_fc} \cdot \mathrm{s\_RNA}}$$

$$\mathrm{init\_Spdef} \rightarrow \frac{\mathrm{init\_spdef} \cdot \mathrm{p\_Spdef}}{\mathrm{d\_Spdef} \cdot \mathrm{reg\_spdef\_fc} \cdot \mathrm{s\_RNA}}$$

$$\mathrm{init\_spdef} \rightarrow \frac{\mathrm{init\_spdef}}{\mathrm{reg\_spdef\_fc} \cdot \mathrm{s\_RNA}}$$

$$\mathrm{init\_tp63} \rightarrow \frac{\mathrm{init\_tp63}}{\mathrm{reg\_tp63\_fc} \cdot \mathrm{s\_RNA}}$$

$$\mathrm{k\_hes7\_i\_spdef} \rightarrow \frac{\mathrm{init\_spdef} \cdot \mathrm{k\_hes7\_i\_spdef\_fc} \cdot \mathrm{p\_Spdef}}{\mathrm{d\_Spdef} \cdot \mathrm{reg\_spdef\_fc} \cdot \mathrm{s\_RNA}}$$

$$\mathrm{k\_ssc\_i\_spdef} \rightarrow \frac{\mathrm{init\_spdef} \cdot \mathrm{k\_ssc\_a\_spdef\_fc} \cdot \mathrm{k\_ssc\_i\_spdef\_fc} \cdot \mathrm{p\_Spdef}}{\mathrm{d\_Spdef} \cdot \mathrm{reg\_spdef\_fc} \cdot \mathrm{s\_RNA}}$$

$$\mathrm{k\_ssc\_a\_spdef} \rightarrow \frac{\mathrm{init\_spdef} \cdot \mathrm{k\_ssc\_a\_spdef\_fc} \cdot \mathrm{p\_Spdef}}{\mathrm{d\_Spdef} \cdot \mathrm{reg\_spdef\_fc} \cdot \mathrm{s\_RNA}}$$

$$\text{k\_tp63\_a\_spdef} \to \frac{\text{init\_spdef} \cdot \text{k\_tp63\_a\_spdef\_fc} \cdot \text{p\_Spdef}}{\text{d\_Spdef} \cdot \text{reg\_spdef\_fc} \cdot \text{s\_RNA}}$$

$$\text{s\_cell1} \to \text{s\_cell1\_fc} \cdot \text{s\_cell\_mean}$$

$$\text{s\_cell12} \to \text{s\_cell12\_fc} \cdot \text{s\_cell\_mean}$$

$$\text{s\_cell13} \to \text{s\_cell13\_fc} \cdot \text{s\_cell\_mean}$$

$$\text{s\_cell14} \to \text{s\_cell14\_fc} \cdot \text{s\_cell\_mean}$$

$$\text{s\_cell2} \to \text{s\_cell2\_fc} \cdot \text{s\_cell\_mean}$$

$$\text{s\_cell20} \to \text{s\_cell20\_fc} \cdot \text{s\_cell\_mean}$$

$$\text{s\_cell22} \to \text{s\_cell22\_fc} \cdot \text{s\_cell\_mean}$$

$$\text{s\_cell23} \to \text{s\_cell23\_fc} \cdot \text{s\_cell\_mean}$$

$$\text{s\_cell24} \to \text{s\_cell24\_fc} \cdot \text{s\_cell\_mean}$$

$$\text{s\_cell25} \to \text{s\_cell25\_fc} \cdot \text{s\_cell\_mean}$$

$$\text{s\_cell3} \to \text{s\_cell3\_fc} \cdot \text{s\_cell\_mean}$$

$$\text{s\_cell4} \to \text{s\_cell4\_fc} \cdot \text{s\_cell\_mean}$$

$$\text{s\_cell5} \to \text{s\_cell5\_fc} \cdot \text{s\_cell\_mean}$$

$$\text{s\_cell8} \to \text{s\_cell8\_fc} \cdot \text{s\_cell\_mean}$$

$$\text{s\_cell9} \to \text{s\_cell9\_fc} \cdot \text{s\_cell\_mean}$$

$$\text{transl\_hes4\_lof\_fc} \to 1$$

$$\text{transl\_hes5\_lof\_fc} \to 1$$

$$\text{transl\_hes7\_lof\_fc} \to 1$$

- **Local condition #17 (global condition #1):**

$$\text{ex1} \to 0$$

$$\text{ex12} \to 0$$

$$\text{ex13} \to 0$$

$$\text{ex14} \to 0$$

$$\text{ex2} \to 0$$

$$\text{ex20} \to 0$$

$$\text{ex22} \to 0$$

$$\text{ex23} \to 0$$

$$\text{ex24} \to 0$$

$$\text{ex25} \to 0$$

$$\text{ex3} \to 0$$

$$\text{ex4} \to 0$$

$$\text{ex5} \to 0$$

$$\text{ex8} \to 0$$

$$\text{ex9} \to 1$$

$$\text{init\_hes4} \to \frac{\text{init\_hes4}}{\text{reg\_hes4\_fc} \cdot \text{s\_RNA}}$$

$$\text{init\_hes4\_d} \to \frac{4 \cdot \text{init\_hes4}}{\text{d\_hes4\_prot\_fc} \cdot \text{d\_hes\_prot} \cdot \text{hes\_delay} \cdot \text{reg\_hes4\_fc} \cdot \text{s\_RNA}}$$

$$\text{init\_hes4\_d1} \rightarrow \frac{\text{init\_hes4}}{\text{reg\_hes4\_fc} \cdot \text{s\_RNA}}$$
$$\text{init\_hes4\_d2} \rightarrow \frac{\text{init\_hes4}}{\text{reg\_hes4\_fc} \cdot \text{s\_RNA}}$$
$$\text{init\_hes4\_d3} \rightarrow \frac{\text{init\_hes4}}{\text{reg\_hes4\_fc} \cdot \text{s\_RNA}}$$
$$\text{init\_hes5} \rightarrow \frac{\text{init\_hes5}}{\text{reg\_hes5\_fc} \cdot \text{s\_RNA}}$$
$$\text{init\_hes5\_d} \rightarrow \frac{4 \cdot \text{init\_hes5}}{\text{d\_hes5\_prot\_fc} \cdot \text{d\_hes\_prot} \cdot \text{hes\_delay} \cdot \text{reg\_hes5\_fc} \cdot \text{s\_RNA}}$$
$$\text{init\_hes5\_d1} \rightarrow \frac{\text{init\_hes5}}{\text{reg\_hes5\_fc} \cdot \text{s\_RNA}}$$
$$\text{init\_hes5\_d2} \rightarrow \frac{\text{init\_hes5}}{\text{reg\_hes5\_fc} \cdot \text{s\_RNA}}$$
$$\text{init\_hes5\_d3} \rightarrow \frac{\text{init\_hes5}}{\text{reg\_hes5\_fc} \cdot \text{s\_RNA}}$$
$$\text{init\_hes7} \rightarrow \frac{\text{init\_hes7}}{\text{reg\_hes7\_fc} \cdot \text{s\_RNA}}$$
$$\text{init\_hes7\_d} \rightarrow \frac{4 \cdot \text{init\_hes7}}{\text{d\_hes7\_prot\_fc} \cdot \text{d\_hes\_prot} \cdot \text{hes\_delay} \cdot \text{reg\_hes7\_fc} \cdot \text{s\_RNA}}$$
$$\text{init\_hes7\_d1} \rightarrow \frac{\text{init\_hes7}}{\text{reg\_hes7\_fc} \cdot \text{s\_RNA}}$$
$$\text{init\_hes7\_d2} \rightarrow \frac{\text{init\_hes7}}{\text{reg\_hes7\_fc} \cdot \text{s\_RNA}}$$
$$\text{init\_hes7\_d3} \rightarrow \frac{\text{init\_hes7}}{\text{reg\_hes7\_fc} \cdot \text{s\_RNA}}$$
$$\text{init\_Spdef} \rightarrow \frac{\text{init\_spdef} \cdot \text{p\_Spdef}}{\text{d\_Spdef} \cdot \text{reg\_spdef\_fc} \cdot \text{s\_RNA}}$$
$$\text{init\_spdef} \rightarrow \frac{\text{init\_spdef}}{\text{reg\_spdef\_fc} \cdot \text{s\_RNA}}$$
$$\text{init\_tp63} \rightarrow \frac{\text{init\_tp63}}{\text{reg\_tp63\_fc} \cdot \text{s\_RNA}}$$
$$\text{k\_hes7\_i\_spdef} \rightarrow \frac{\text{init\_spdef} \cdot \text{k\_hes7\_i\_spdef\_fc} \cdot \text{p\_Spdef}}{\text{d\_Spdef} \cdot \text{reg\_spdef\_fc} \cdot \text{s\_RNA}}$$
$$\text{k\_ssc\_i\_spdef} \rightarrow \frac{\text{init\_spdef} \cdot \text{k\_ssc\_a\_spdef\_fc} \cdot \text{k\_ssc\_i\_spdef\_fc} \cdot \text{p\_Spdef}}{\text{d\_Spdef} \cdot \text{reg\_spdef\_fc} \cdot \text{s\_RNA}}$$
$$\text{k\_ssc\_a\_spdef} \rightarrow \frac{\text{init\_spdef} \cdot \text{k\_ssc\_a\_spdef\_fc} \cdot \text{p\_Spdef}}{\text{d\_Spdef} \cdot \text{reg\_spdef\_fc} \cdot \text{s\_RNA}}$$
$$\text{k\_tp63\_a\_spdef} \rightarrow \frac{\text{init\_spdef} \cdot \text{k\_tp63\_a\_spdef\_fc} \cdot \text{p\_Spdef}}{\text{d\_Spdef} \cdot \text{reg\_spdef\_fc} \cdot \text{s\_RNA}}$$
$$\text{s\_cell1} \rightarrow \text{s\_cell1\_fc} \cdot \text{s\_cell\_mean}$$
$$\text{s\_cell12} \rightarrow \text{s\_cell12\_fc} \cdot \text{s\_cell\_mean}$$
$$\text{s\_cell13} \rightarrow \text{s\_cell13\_fc} \cdot \text{s\_cell\_mean}$$
$$\text{s\_cell14} \rightarrow \text{s\_cell14\_fc} \cdot \text{s\_cell\_mean}$$
$$\text{s\_cell2} \rightarrow \text{s\_cell2\_fc} \cdot \text{s\_cell\_mean}$$

$$s\_cell20 \to s\_cell20\_fc \cdot s\_cell\_mean$$
$$s\_cell22 \to s\_cell22\_fc \cdot s\_cell\_mean$$
$$s\_cell23 \to s\_cell23\_fc \cdot s\_cell\_mean$$
$$s\_cell24 \to s\_cell24\_fc \cdot s\_cell\_mean$$
$$s\_cell25 \to s\_cell25\_fc \cdot s\_cell\_mean$$
$$s\_cell3 \to s\_cell3\_fc \cdot s\_cell\_mean$$
$$s\_cell4 \to s\_cell4\_fc \cdot s\_cell\_mean$$
$$s\_cell5 \to s\_cell5\_fc \cdot s\_cell\_mean$$
$$s\_cell8 \to s\_cell8\_fc \cdot s\_cell\_mean$$
$$s\_cell9 \to s\_cell9\_fc \cdot s\_cell\_mean$$
$$transl\_hes4\_lof\_fc \to 1$$
$$transl\_hes5\_lof\_fc \to 1$$
$$transl\_hes7\_lof\_fc \to 1$$

- **Local condition #18 (global condition #1):**

$$ex1 \to 0$$
$$ex12 \to 0$$
$$ex13 \to 0$$
$$ex14 \to 0$$
$$ex2 \to 0$$
$$ex20 \to 0$$
$$ex22 \to 0$$
$$ex23 \to 0$$
$$ex24 \to 0$$
$$ex25 \to 0$$
$$ex3 \to 0$$
$$ex4 \to 0$$
$$ex5 \to 0$$
$$ex8 \to 1$$
$$ex9 \to 0$$
$$init\_hes4 \to \frac{init\_hes4}{reg\_hes4\_fc \cdot s\_RNA}$$
$$init\_hes4\_d \to \frac{4 \cdot init\_hes4}{d\_hes4\_prot\_fc \cdot d\_hes\_prot \cdot hes\_delay \cdot reg\_hes4\_fc \cdot s\_RNA}$$
$$init\_hes4\_d1 \to \frac{init\_hes4}{reg\_hes4\_fc \cdot s\_RNA}$$
$$init\_hes4\_d2 \to \frac{init\_hes4}{reg\_hes4\_fc \cdot s\_RNA}$$
$$init\_hes4\_d3 \to \frac{init\_hes4}{reg\_hes4\_fc \cdot s\_RNA}$$
$$init\_hes5 \to \frac{init\_hes5}{reg\_hes5\_fc \cdot s\_RNA}$$

$$\mathrm{init\_hes5\_d} \rightarrow \frac{4 \cdot \mathrm{init\_hes5}}{\mathrm{d\_hes5\_prot\_fc} \cdot \mathrm{d\_hes\_prot} \cdot \mathrm{hes\_delay} \cdot \mathrm{reg\_hes5\_fc} \cdot \mathrm{s\_RNA}}$$

$$\mathrm{init\_hes5\_d1} \rightarrow \frac{\mathrm{init\_hes5}}{\mathrm{reg\_hes5\_fc} \cdot \mathrm{s\_RNA}}$$

$$\mathrm{init\_hes5\_d2} \rightarrow \frac{\mathrm{init\_hes5}}{\mathrm{reg\_hes5\_fc} \cdot \mathrm{s\_RNA}}$$

$$\mathrm{init\_hes5\_d3} \rightarrow \frac{\mathrm{init\_hes5}}{\mathrm{reg\_hes5\_fc} \cdot \mathrm{s\_RNA}}$$

$$\mathrm{init\_hes7} \rightarrow \frac{\mathrm{init\_hes7}}{\mathrm{reg\_hes7\_fc} \cdot \mathrm{s\_RNA}}$$

$$\mathrm{init\_hes7\_d} \rightarrow \frac{4 \cdot \mathrm{init\_hes7}}{\mathrm{d\_hes7\_prot\_fc} \cdot \mathrm{d\_hes\_prot} \cdot \mathrm{hes\_delay} \cdot \mathrm{reg\_hes7\_fc} \cdot \mathrm{s\_RNA}}$$

$$\mathrm{init\_hes7\_d1} \rightarrow \frac{\mathrm{init\_hes7}}{\mathrm{reg\_hes7\_fc} \cdot \mathrm{s\_RNA}}$$

$$\mathrm{init\_hes7\_d2} \rightarrow \frac{\mathrm{init\_hes7}}{\mathrm{reg\_hes7\_fc} \cdot \mathrm{s\_RNA}}$$

$$\mathrm{init\_hes7\_d3} \rightarrow \frac{\mathrm{init\_hes7}}{\mathrm{reg\_hes7\_fc} \cdot \mathrm{s\_RNA}}$$

$$\mathrm{init\_Spdef} \rightarrow \frac{\mathrm{init\_spdef} \cdot \mathrm{p\_Spdef}}{\mathrm{d\_Spdef} \cdot \mathrm{reg\_spdef\_fc} \cdot \mathrm{s\_RNA}}$$

$$\mathrm{init\_spdef} \rightarrow \frac{\mathrm{init\_spdef}}{\mathrm{reg\_spdef\_fc} \cdot \mathrm{s\_RNA}}$$

$$\mathrm{init\_tp63} \rightarrow \frac{\mathrm{init\_tp63}}{\mathrm{reg\_tp63\_fc} \cdot \mathrm{s\_RNA}}$$

$$\mathrm{k\_hes7\_i\_spdef} \rightarrow \frac{\mathrm{init\_spdef} \cdot \mathrm{k\_hes7\_i\_spdef\_fc} \cdot \mathrm{p\_Spdef}}{\mathrm{d\_Spdef} \cdot \mathrm{reg\_spdef\_fc} \cdot \mathrm{s\_RNA}}$$

$$\mathrm{k\_ssc\_i\_spdef} \rightarrow \frac{\mathrm{init\_spdef} \cdot \mathrm{k\_ssc\_a\_spdef\_fc} \cdot \mathrm{k\_ssc\_i\_spdef\_fc} \cdot \mathrm{p\_Spdef}}{\mathrm{d\_Spdef} \cdot \mathrm{reg\_spdef\_fc} \cdot \mathrm{s\_RNA}}$$

$$\mathrm{k\_ssc\_a\_spdef} \rightarrow \frac{\mathrm{init\_spdef} \cdot \mathrm{k\_ssc\_a\_spdef\_fc} \cdot \mathrm{p\_Spdef}}{\mathrm{d\_Spdef} \cdot \mathrm{reg\_spdef\_fc} \cdot \mathrm{s\_RNA}}$$

$$\mathrm{k\_tp63\_a\_spdef} \rightarrow \frac{\mathrm{init\_spdef} \cdot \mathrm{k\_tp63\_a\_spdef\_fc} \cdot \mathrm{p\_Spdef}}{\mathrm{d\_Spdef} \cdot \mathrm{reg\_spdef\_fc} \cdot \mathrm{s\_RNA}}$$

$$\mathrm{s\_cell1} \rightarrow \mathrm{s\_cell1\_fc} \cdot \mathrm{s\_cell\_mean}$$

$$\mathrm{s\_cell12} \rightarrow \mathrm{s\_cell12\_fc} \cdot \mathrm{s\_cell\_mean}$$

$$\mathrm{s\_cell13} \rightarrow \mathrm{s\_cell13\_fc} \cdot \mathrm{s\_cell\_mean}$$

$$\mathrm{s\_cell14} \rightarrow \mathrm{s\_cell14\_fc} \cdot \mathrm{s\_cell\_mean}$$

$$\mathrm{s\_cell2} \rightarrow \mathrm{s\_cell2\_fc} \cdot \mathrm{s\_cell\_mean}$$

$$\mathrm{s\_cell20} \rightarrow \mathrm{s\_cell20\_fc} \cdot \mathrm{s\_cell\_mean}$$

$$\mathrm{s\_cell22} \rightarrow \mathrm{s\_cell22\_fc} \cdot \mathrm{s\_cell\_mean}$$

$$\mathrm{s\_cell23} \rightarrow \mathrm{s\_cell23\_fc} \cdot \mathrm{s\_cell\_mean}$$

$$\mathrm{s\_cell24} \rightarrow \mathrm{s\_cell24\_fc} \cdot \mathrm{s\_cell\_mean}$$

$$\mathrm{s\_cell25} \rightarrow \mathrm{s\_cell25\_fc} \cdot \mathrm{s\_cell\_mean}$$

$$\mathrm{s\_cell3} \rightarrow \mathrm{s\_cell3\_fc} \cdot \mathrm{s\_cell\_mean}$$

$$\mathrm{s\_cell4} \rightarrow \mathrm{s\_cell4\_fc} \cdot \mathrm{s\_cell\_mean}$$

$$\text{s\_cell5} \to \text{s\_cell5\_fc} \cdot \text{s\_cell\_mean}$$
$$\text{s\_cell8} \to \text{s\_cell8\_fc} \cdot \text{s\_cell\_mean}$$
$$\text{s\_cell9} \to \text{s\_cell9\_fc} \cdot \text{s\_cell\_mean}$$
$$\text{transl\_hes4\_lof\_fc} \to 1$$
$$\text{transl\_hes5\_lof\_fc} \to 1$$
$$\text{transl\_hes7\_lof\_fc} \to 1$$

- **Local condition #19 (global condition #1):**

$$\text{ex1} \to 0$$
$$\text{ex12} \to 0$$
$$\text{ex13} \to 0$$
$$\text{ex14} \to 0$$
$$\text{ex2} \to 0$$
$$\text{ex20} \to 0$$
$$\text{ex22} \to 0$$
$$\text{ex23} \to 0$$
$$\text{ex24} \to 0$$
$$\text{ex25} \to 0$$
$$\text{ex3} \to 0$$
$$\text{ex4} \to 0$$
$$\text{ex5} \to 1$$
$$\text{ex8} \to 0$$
$$\text{ex9} \to 0$$
$$\text{init\_hes4} \to \frac{\text{init\_hes4}}{\text{reg\_hes4\_fc} \cdot \text{s\_RNA}}$$
$$\text{init\_hes4\_d} \to \frac{4 \cdot \text{init\_hes4}}{\text{d\_hes4\_prot\_fc} \cdot \text{d\_hes\_prot} \cdot \text{hes\_delay} \cdot \text{reg\_hes4\_fc} \cdot \text{s\_RNA}}$$
$$\text{init\_hes4\_d1} \to \frac{\text{init\_hes4}}{\text{reg\_hes4\_fc} \cdot \text{s\_RNA}}$$
$$\text{init\_hes4\_d2} \to \frac{\text{init\_hes4}}{\text{reg\_hes4\_fc} \cdot \text{s\_RNA}}$$
$$\text{init\_hes4\_d3} \to \frac{\text{init\_hes4}}{\text{reg\_hes4\_fc} \cdot \text{s\_RNA}}$$
$$\text{init\_hes5} \to \frac{\text{init\_hes5}}{\text{reg\_hes5\_fc} \cdot \text{s\_RNA}}$$
$$\text{init\_hes5\_d} \to \frac{4 \cdot \text{init\_hes5}}{\text{d\_hes5\_prot\_fc} \cdot \text{d\_hes\_prot} \cdot \text{hes\_delay} \cdot \text{reg\_hes5\_fc} \cdot \text{s\_RNA}}$$
$$\text{init\_hes5\_d1} \to \frac{\text{init\_hes5}}{\text{reg\_hes5\_fc} \cdot \text{s\_RNA}}$$
$$\text{init\_hes5\_d2} \to \frac{\text{init\_hes5}}{\text{reg\_hes5\_fc} \cdot \text{s\_RNA}}$$
$$\text{init\_hes5\_d3} \to \frac{\text{init\_hes5}}{\text{reg\_hes5\_fc} \cdot \text{s\_RNA}}$$

$$\text{init\_hes7} \rightarrow \frac{\text{init\_hes7}}{\text{reg\_hes7\_fc} \cdot \text{s\_RNA}}$$

$$\text{init\_hes7\_d} \rightarrow \frac{4 \cdot \text{init\_hes7}}{\text{d\_hes7\_prot\_fc} \cdot \text{d\_hes\_prot} \cdot \text{hes\_delay} \cdot \text{reg\_hes7\_fc} \cdot \text{s\_RNA}}$$

$$\text{init\_hes7\_d1} \rightarrow \frac{\text{init\_hes7}}{\text{reg\_hes7\_fc} \cdot \text{s\_RNA}}$$

$$\text{init\_hes7\_d2} \rightarrow \frac{\text{init\_hes7}}{\text{reg\_hes7\_fc} \cdot \text{s\_RNA}}$$

$$\text{init\_hes7\_d3} \rightarrow \frac{\text{init\_hes7}}{\text{reg\_hes7\_fc} \cdot \text{s\_RNA}}$$

$$\text{init\_Spdef} \rightarrow \frac{\text{init\_spdef} \cdot \text{p\_Spdef}}{\text{d\_Spdef} \cdot \text{reg\_spdef\_fc} \cdot \text{s\_RNA}}$$

$$\text{init\_spdef} \rightarrow \frac{\text{init\_spdef}}{\text{reg\_spdef\_fc} \cdot \text{s\_RNA}}$$

$$\text{init\_tp63} \rightarrow \frac{\text{init\_tp63}}{\text{reg\_tp63\_fc} \cdot \text{s\_RNA}}$$

$$\text{k\_hes7\_i\_spdef} \rightarrow \frac{\text{init\_spdef} \cdot \text{k\_hes7\_i\_spdef\_fc} \cdot \text{p\_Spdef}}{\text{d\_Spdef} \cdot \text{reg\_spdef\_fc} \cdot \text{s\_RNA}}$$

$$\text{k\_ssc\_i\_spdef} \rightarrow \frac{\text{init\_spdef} \cdot \text{k\_ssc\_a\_spdef\_fc} \cdot \text{k\_ssc\_i\_spdef\_fc} \cdot \text{p\_Spdef}}{\text{d\_Spdef} \cdot \text{reg\_spdef\_fc} \cdot \text{s\_RNA}}$$

$$\text{k\_ssc\_a\_spdef} \rightarrow \frac{\text{init\_spdef} \cdot \text{k\_ssc\_a\_spdef\_fc} \cdot \text{p\_Spdef}}{\text{d\_Spdef} \cdot \text{reg\_spdef\_fc} \cdot \text{s\_RNA}}$$

$$\text{k\_tp63\_a\_spdef} \rightarrow \frac{\text{init\_spdef} \cdot \text{k\_tp63\_a\_spdef\_fc} \cdot \text{p\_Spdef}}{\text{d\_Spdef} \cdot \text{reg\_spdef\_fc} \cdot \text{s\_RNA}}$$

$$\text{s\_cell1} \rightarrow \text{s\_cell1\_fc} \cdot \text{s\_cell\_mean}$$

$$\text{s\_cell12} \rightarrow \text{s\_cell12\_fc} \cdot \text{s\_cell\_mean}$$

$$\text{s\_cell13} \rightarrow \text{s\_cell13\_fc} \cdot \text{s\_cell\_mean}$$

$$\text{s\_cell14} \rightarrow \text{s\_cell14\_fc} \cdot \text{s\_cell\_mean}$$

$$\text{s\_cell2} \rightarrow \text{s\_cell2\_fc} \cdot \text{s\_cell\_mean}$$

$$\text{s\_cell20} \rightarrow \text{s\_cell20\_fc} \cdot \text{s\_cell\_mean}$$

$$\text{s\_cell22} \rightarrow \text{s\_cell22\_fc} \cdot \text{s\_cell\_mean}$$

$$\text{s\_cell23} \rightarrow \text{s\_cell23\_fc} \cdot \text{s\_cell\_mean}$$

$$\text{s\_cell24} \rightarrow \text{s\_cell24\_fc} \cdot \text{s\_cell\_mean}$$

$$\text{s\_cell25} \rightarrow \text{s\_cell25\_fc} \cdot \text{s\_cell\_mean}$$

$$\text{s\_cell3} \rightarrow \text{s\_cell3\_fc} \cdot \text{s\_cell\_mean}$$

$$\text{s\_cell4} \rightarrow \text{s\_cell4\_fc} \cdot \text{s\_cell\_mean}$$

$$\text{s\_cell5} \rightarrow \text{s\_cell5\_fc} \cdot \text{s\_cell\_mean}$$

$$\text{s\_cell8} \rightarrow \text{s\_cell8\_fc} \cdot \text{s\_cell\_mean}$$

$$\text{s\_cell9} \rightarrow \text{s\_cell9\_fc} \cdot \text{s\_cell\_mean}$$

$$\text{transl\_hes4\_lof\_fc} \rightarrow 1$$

$$\text{transl\_hes5\_lof\_fc} \rightarrow 1$$

$$\text{transl\_hes7\_lof\_fc} \rightarrow 1$$

- **Local condition #20 (global condition #1):**

$$\begin{aligned}
\text{ex1} &\to 0\\
\text{ex12} &\to 0\\
\text{ex13} &\to 0\\
\text{ex14} &\to 0\\
\text{ex2} &\to 0\\
\text{ex20} &\to 0\\
\text{ex22} &\to 0\\
\text{ex23} &\to 0\\
\text{ex24} &\to 0\\
\text{ex25} &\to 0\\
\text{ex3} &\to 0\\
\text{ex4} &\to 1\\
\text{ex5} &\to 0\\
\text{ex8} &\to 0\\
\text{ex9} &\to 0\\
\text{init_hes4} &\to \frac{\text{init_hes4}}{\text{reg_hes4_fc} \cdot \text{s_RNA}}\\
\text{init_hes4_d} &\to \frac{4 \cdot \text{init_hes4}}{\text{d_hes4_prot_fc} \cdot \text{d_hes_prot} \cdot \text{hes_delay} \cdot \text{reg_hes4_fc} \cdot \text{s_RNA}}\\
\text{init_hes4_d1} &\to \frac{\text{init_hes4}}{\text{reg_hes4_fc} \cdot \text{s_RNA}}\\
\text{init_hes4_d2} &\to \frac{\text{init_hes4}}{\text{reg_hes4_fc} \cdot \text{s_RNA}}\\
\text{init_hes4_d3} &\to \frac{\text{init_hes4}}{\text{reg_hes4_fc} \cdot \text{s_RNA}}\\
\text{init_hes5} &\to \frac{\text{init_hes5}}{\text{reg_hes5_fc} \cdot \text{s_RNA}}\\
\text{init_hes5_d} &\to \frac{4 \cdot \text{init_hes5}}{\text{d_hes5_prot_fc} \cdot \text{d_hes_prot} \cdot \text{hes_delay} \cdot \text{reg_hes5_fc} \cdot \text{s_RNA}}\\
\text{init_hes5_d1} &\to \frac{\text{init_hes5}}{\text{reg_hes5_fc} \cdot \text{s_RNA}}\\
\text{init_hes5_d2} &\to \frac{\text{init_hes5}}{\text{reg_hes5_fc} \cdot \text{s_RNA}}\\
\text{init_hes5_d3} &\to \frac{\text{init_hes5}}{\text{reg_hes5_fc} \cdot \text{s_RNA}}\\
\text{init_hes7} &\to \frac{\text{init_hes7}}{\text{reg_hes7_fc} \cdot \text{s_RNA}}\\
\text{init_hes7_d} &\to \frac{4 \cdot \text{init_hes7}}{\text{d_hes7_prot_fc} \cdot \text{d_hes_prot} \cdot \text{hes_delay} \cdot \text{reg_hes7_fc} \cdot \text{s_RNA}}\\
\text{init_hes7_d1} &\to \frac{\text{init_hes7}}{\text{reg_hes7_fc} \cdot \text{s_RNA}}\\
\text{init_hes7_d2} &\to \frac{\text{init_hes7}}{\text{reg_hes7_fc} \cdot \text{s_RNA}}
\end{aligned}$$

$$\text{init\_hes7\_d3} \rightarrow \frac{\text{init\_hes7}}{\text{reg\_hes7\_fc} \cdot \text{s\_RNA}}$$
$$\text{init\_Spdef} \rightarrow \frac{\text{init\_spdef} \cdot \text{p\_Spdef}}{\text{d\_Spdef} \cdot \text{reg\_spdef\_fc} \cdot \text{s\_RNA}}$$
$$\text{init\_spdef} \rightarrow \frac{\text{init\_spdef}}{\text{reg\_spdef\_fc} \cdot \text{s\_RNA}}$$
$$\text{init\_tp63} \rightarrow \frac{\text{init\_tp63}}{\text{reg\_tp63\_fc} \cdot \text{s\_RNA}}$$
$$\text{k\_hes7\_i\_spdef} \rightarrow \frac{\text{init\_spdef} \cdot \text{k\_hes7\_i\_spdef\_fc} \cdot \text{p\_Spdef}}{\text{d\_Spdef} \cdot \text{reg\_spdef\_fc} \cdot \text{s\_RNA}}$$
$$\text{k\_ssc\_i\_spdef} \rightarrow \frac{\text{init\_spdef} \cdot \text{k\_ssc\_a\_spdef\_fc} \cdot \text{k\_ssc\_i\_spdef\_fc} \cdot \text{p\_Spdef}}{\text{d\_Spdef} \cdot \text{reg\_spdef\_fc} \cdot \text{s\_RNA}}$$
$$\text{k\_ssc\_a\_spdef} \rightarrow \frac{\text{init\_spdef} \cdot \text{k\_ssc\_a\_spdef\_fc} \cdot \text{p\_Spdef}}{\text{d\_Spdef} \cdot \text{reg\_spdef\_fc} \cdot \text{s\_RNA}}$$
$$\text{k\_tp63\_a\_spdef} \rightarrow \frac{\text{init\_spdef} \cdot \text{k\_tp63\_a\_spdef\_fc} \cdot \text{p\_Spdef}}{\text{d\_Spdef} \cdot \text{reg\_spdef\_fc} \cdot \text{s\_RNA}}$$
$$\text{s\_cell1} \rightarrow \text{s\_cell1\_fc} \cdot \text{s\_cell\_mean}$$
$$\text{s\_cell12} \rightarrow \text{s\_cell12\_fc} \cdot \text{s\_cell\_mean}$$
$$\text{s\_cell13} \rightarrow \text{s\_cell13\_fc} \cdot \text{s\_cell\_mean}$$
$$\text{s\_cell14} \rightarrow \text{s\_cell14\_fc} \cdot \text{s\_cell\_mean}$$
$$\text{s\_cell2} \rightarrow \text{s\_cell2\_fc} \cdot \text{s\_cell\_mean}$$
$$\text{s\_cell20} \rightarrow \text{s\_cell20\_fc} \cdot \text{s\_cell\_mean}$$
$$\text{s\_cell22} \rightarrow \text{s\_cell22\_fc} \cdot \text{s\_cell\_mean}$$
$$\text{s\_cell23} \rightarrow \text{s\_cell23\_fc} \cdot \text{s\_cell\_mean}$$
$$\text{s\_cell24} \rightarrow \text{s\_cell24\_fc} \cdot \text{s\_cell\_mean}$$
$$\text{s\_cell25} \rightarrow \text{s\_cell25\_fc} \cdot \text{s\_cell\_mean}$$
$$\text{s\_cell3} \rightarrow \text{s\_cell3\_fc} \cdot \text{s\_cell\_mean}$$
$$\text{s\_cell4} \rightarrow \text{s\_cell4\_fc} \cdot \text{s\_cell\_mean}$$
$$\text{s\_cell5} \rightarrow \text{s\_cell5\_fc} \cdot \text{s\_cell\_mean}$$
$$\text{s\_cell8} \rightarrow \text{s\_cell8\_fc} \cdot \text{s\_cell\_mean}$$
$$\text{s\_cell9} \rightarrow \text{s\_cell9\_fc} \cdot \text{s\_cell\_mean}$$
$$\text{transl\_hes4\_lof\_fc} \rightarrow 1$$
$$\text{transl\_hes5\_lof\_fc} \rightarrow 1$$
$$\text{transl\_hes7\_lof\_fc} \rightarrow 1$$

- **Local condition #21 (global condition #1):**

$$\text{ex1} \rightarrow 0$$
$$\text{ex12} \rightarrow 0$$
$$\text{ex13} \rightarrow 0$$
$$\text{ex14} \rightarrow 0$$
$$\text{ex2} \rightarrow 0$$
$$\text{ex20} \rightarrow 0$$

$$\mathrm{ex22} \rightarrow 0$$
$$\mathrm{ex23} \rightarrow 0$$
$$\mathrm{ex24} \rightarrow 0$$
$$\mathrm{ex25} \rightarrow 0$$
$$\mathrm{ex3} \rightarrow 1$$
$$\mathrm{ex4} \rightarrow 0$$
$$\mathrm{ex5} \rightarrow 0$$
$$\mathrm{ex8} \rightarrow 0$$
$$\mathrm{ex9} \rightarrow 0$$
$$\mathrm{init\_hes4} \rightarrow \frac{\mathrm{init\_hes4}}{\mathrm{reg\_hes4\_fc} \cdot \mathrm{s\_RNA}}$$
$$\mathrm{init\_hes4\_d} \rightarrow \frac{4 \cdot \mathrm{init\_hes4}}{\mathrm{d\_hes4\_prot\_fc} \cdot \mathrm{d\_hes\_prot} \cdot \mathrm{hes\_delay} \cdot \mathrm{reg\_hes4\_fc} \cdot \mathrm{s\_RNA}}$$
$$\mathrm{init\_hes4\_d1} \rightarrow \frac{\mathrm{init\_hes4}}{\mathrm{reg\_hes4\_fc} \cdot \mathrm{s\_RNA}}$$
$$\mathrm{init\_hes4\_d2} \rightarrow \frac{\mathrm{init\_hes4}}{\mathrm{reg\_hes4\_fc} \cdot \mathrm{s\_RNA}}$$
$$\mathrm{init\_hes4\_d3} \rightarrow \frac{\mathrm{init\_hes4}}{\mathrm{reg\_hes4\_fc} \cdot \mathrm{s\_RNA}}$$
$$\mathrm{init\_hes5} \rightarrow \frac{\mathrm{init\_hes5}}{\mathrm{reg\_hes5\_fc} \cdot \mathrm{s\_RNA}}$$
$$\mathrm{init\_hes5\_d} \rightarrow \frac{4 \cdot \mathrm{init\_hes5}}{\mathrm{d\_hes5\_prot\_fc} \cdot \mathrm{d\_hes\_prot} \cdot \mathrm{hes\_delay} \cdot \mathrm{reg\_hes5\_fc} \cdot \mathrm{s\_RNA}}$$
$$\mathrm{init\_hes5\_d1} \rightarrow \frac{\mathrm{init\_hes5}}{\mathrm{reg\_hes5\_fc} \cdot \mathrm{s\_RNA}}$$
$$\mathrm{init\_hes5\_d2} \rightarrow \frac{\mathrm{init\_hes5}}{\mathrm{reg\_hes5\_fc} \cdot \mathrm{s\_RNA}}$$
$$\mathrm{init\_hes5\_d3} \rightarrow \frac{\mathrm{init\_hes5}}{\mathrm{reg\_hes5\_fc} \cdot \mathrm{s\_RNA}}$$
$$\mathrm{init\_hes7} \rightarrow \frac{\mathrm{init\_hes7}}{\mathrm{reg\_hes7\_fc} \cdot \mathrm{s\_RNA}}$$
$$\mathrm{init\_hes7\_d} \rightarrow \frac{4 \cdot \mathrm{init\_hes7}}{\mathrm{d\_hes7\_prot\_fc} \cdot \mathrm{d\_hes\_prot} \cdot \mathrm{hes\_delay} \cdot \mathrm{reg\_hes7\_fc} \cdot \mathrm{s\_RNA}}$$
$$\mathrm{init\_hes7\_d1} \rightarrow \frac{\mathrm{init\_hes7}}{\mathrm{reg\_hes7\_fc} \cdot \mathrm{s\_RNA}}$$
$$\mathrm{init\_hes7\_d2} \rightarrow \frac{\mathrm{init\_hes7}}{\mathrm{reg\_hes7\_fc} \cdot \mathrm{s\_RNA}}$$
$$\mathrm{init\_hes7\_d3} \rightarrow \frac{\mathrm{init\_hes7}}{\mathrm{reg\_hes7\_fc} \cdot \mathrm{s\_RNA}}$$
$$\mathrm{init\_Spdef} \rightarrow \frac{\mathrm{init\_spdef} \cdot \mathrm{p\_Spdef}}{\mathrm{d\_Spdef} \cdot \mathrm{reg\_spdef\_fc} \cdot \mathrm{s\_RNA}}$$
$$\mathrm{init\_spdef} \rightarrow \frac{\mathrm{init\_spdef}}{\mathrm{reg\_spdef\_fc} \cdot \mathrm{s\_RNA}}$$

$$\mathrm{init\_tp63} \to \frac{\mathrm{init\_tp63}}{\mathrm{reg\_tp63\_fc} \cdot \mathrm{s\_RNA}}$$
$$\mathrm{k\_hes7\_i\_spdef} \to \frac{\mathrm{init\_spdef} \cdot \mathrm{k\_hes7\_i\_spdef\_fc} \cdot \mathrm{p\_Spdef}}{\mathrm{d\_Spdef} \cdot \mathrm{reg\_spdef\_fc} \cdot \mathrm{s\_RNA}}$$
$$\mathrm{k\_ssc\_i\_spdef} \to \frac{\mathrm{init\_spdef} \cdot \mathrm{k\_ssc\_a\_spdef\_fc} \cdot \mathrm{k\_ssc\_i\_spdef\_fc} \cdot \mathrm{p\_Spdef}}{\mathrm{d\_Spdef} \cdot \mathrm{reg\_spdef\_fc} \cdot \mathrm{s\_RNA}}$$
$$\mathrm{k\_ssc\_a\_spdef} \to \frac{\mathrm{init\_spdef} \cdot \mathrm{k\_ssc\_a\_spdef\_fc} \cdot \mathrm{p\_Spdef}}{\mathrm{d\_Spdef} \cdot \mathrm{reg\_spdef\_fc} \cdot \mathrm{s\_RNA}}$$
$$\mathrm{k\_tp63\_a\_spdef} \to \frac{\mathrm{init\_spdef} \cdot \mathrm{k\_tp63\_a\_spdef\_fc} \cdot \mathrm{p\_Spdef}}{\mathrm{d\_Spdef} \cdot \mathrm{reg\_spdef\_fc} \cdot \mathrm{s\_RNA}}$$
$$\mathrm{s\_cell1} \to \mathrm{s\_cell1\_fc} \cdot \mathrm{s\_cell\_mean}$$
$$\mathrm{s\_cell12} \to \mathrm{s\_cell12\_fc} \cdot \mathrm{s\_cell\_mean}$$
$$\mathrm{s\_cell13} \to \mathrm{s\_cell13\_fc} \cdot \mathrm{s\_cell\_mean}$$
$$\mathrm{s\_cell14} \to \mathrm{s\_cell14\_fc} \cdot \mathrm{s\_cell\_mean}$$
$$\mathrm{s\_cell2} \to \mathrm{s\_cell2\_fc} \cdot \mathrm{s\_cell\_mean}$$
$$\mathrm{s\_cell20} \to \mathrm{s\_cell20\_fc} \cdot \mathrm{s\_cell\_mean}$$
$$\mathrm{s\_cell22} \to \mathrm{s\_cell22\_fc} \cdot \mathrm{s\_cell\_mean}$$
$$\mathrm{s\_cell23} \to \mathrm{s\_cell23\_fc} \cdot \mathrm{s\_cell\_mean}$$
$$\mathrm{s\_cell24} \to \mathrm{s\_cell24\_fc} \cdot \mathrm{s\_cell\_mean}$$
$$\mathrm{s\_cell25} \to \mathrm{s\_cell25\_fc} \cdot \mathrm{s\_cell\_mean}$$
$$\mathrm{s\_cell3} \to \mathrm{s\_cell3\_fc} \cdot \mathrm{s\_cell\_mean}$$
$$\mathrm{s\_cell4} \to \mathrm{s\_cell4\_fc} \cdot \mathrm{s\_cell\_mean}$$
$$\mathrm{s\_cell5} \to \mathrm{s\_cell5\_fc} \cdot \mathrm{s\_cell\_mean}$$
$$\mathrm{s\_cell8} \to \mathrm{s\_cell8\_fc} \cdot \mathrm{s\_cell\_mean}$$
$$\mathrm{s\_cell9} \to \mathrm{s\_cell9\_fc} \cdot \mathrm{s\_cell\_mean}$$
$$\mathrm{transl\_hes4\_lof\_fc} \to 1$$
$$\mathrm{transl\_hes5\_lof\_fc} \to 1$$
$$\mathrm{transl\_hes7\_lof\_fc} \to 1$$

- **Local condition #22 (global condition #1):**

$$\mathrm{ex1} \to 0$$
$$\mathrm{ex12} \to 0$$
$$\mathrm{ex13} \to 0$$
$$\mathrm{ex14} \to 0$$
$$\mathrm{ex2} \to 1$$
$$\mathrm{ex20} \to 0$$
$$\mathrm{ex22} \to 0$$
$$\mathrm{ex23} \to 0$$
$$\mathrm{ex24} \to 0$$
$$\mathrm{ex25} \to 0$$
$$\mathrm{ex3} \to 0$$
$$\mathrm{ex4} \to 0$$

$$\mathrm{ex5} \rightarrow 0$$
$$\mathrm{ex8} \rightarrow 0$$
$$\mathrm{ex9} \rightarrow 0$$
$$\mathrm{init\_hes4} \rightarrow \frac{\mathrm{init\_hes4}}{\mathrm{reg\_hes4\_fc} \cdot \mathrm{s\_RNA}}$$
$$\mathrm{init\_hes4\_d} \rightarrow \frac{4 \cdot \mathrm{init\_hes4}}{\mathrm{d\_hes4\_prot\_fc} \cdot \mathrm{d\_hes\_prot} \cdot \mathrm{hes\_delay} \cdot \mathrm{reg\_hes4\_fc} \cdot \mathrm{s\_RNA}}$$
$$\mathrm{init\_hes4\_d1} \rightarrow \frac{\mathrm{init\_hes4}}{\mathrm{reg\_hes4\_fc} \cdot \mathrm{s\_RNA}}$$
$$\mathrm{init\_hes4\_d2} \rightarrow \frac{\mathrm{init\_hes4}}{\mathrm{reg\_hes4\_fc} \cdot \mathrm{s\_RNA}}$$
$$\mathrm{init\_hes4\_d3} \rightarrow \frac{\mathrm{init\_hes4}}{\mathrm{reg\_hes4\_fc} \cdot \mathrm{s\_RNA}}$$
$$\mathrm{init\_hes5} \rightarrow \frac{\mathrm{init\_hes5}}{\mathrm{reg\_hes5\_fc} \cdot \mathrm{s\_RNA}}$$
$$\mathrm{init\_hes5\_d} \rightarrow \frac{4 \cdot \mathrm{init\_hes5}}{\mathrm{d\_hes5\_prot\_fc} \cdot \mathrm{d\_hes\_prot} \cdot \mathrm{hes\_delay} \cdot \mathrm{reg\_hes5\_fc} \cdot \mathrm{s\_RNA}}$$
$$\mathrm{init\_hes5\_d1} \rightarrow \frac{\mathrm{init\_hes5}}{\mathrm{reg\_hes5\_fc} \cdot \mathrm{s\_RNA}}$$
$$\mathrm{init\_hes5\_d2} \rightarrow \frac{\mathrm{init\_hes5}}{\mathrm{reg\_hes5\_fc} \cdot \mathrm{s\_RNA}}$$
$$\mathrm{init\_hes5\_d3} \rightarrow \frac{\mathrm{init\_hes5}}{\mathrm{reg\_hes5\_fc} \cdot \mathrm{s\_RNA}}$$
$$\mathrm{init\_hes7} \rightarrow \frac{\mathrm{init\_hes7}}{\mathrm{reg\_hes7\_fc} \cdot \mathrm{s\_RNA}}$$
$$\mathrm{init\_hes7\_d} \rightarrow \frac{4 \cdot \mathrm{init\_hes7}}{\mathrm{d\_hes7\_prot\_fc} \cdot \mathrm{d\_hes\_prot} \cdot \mathrm{hes\_delay} \cdot \mathrm{reg\_hes7\_fc} \cdot \mathrm{s\_RNA}}$$
$$\mathrm{init\_hes7\_d1} \rightarrow \frac{\mathrm{init\_hes7}}{\mathrm{reg\_hes7\_fc} \cdot \mathrm{s\_RNA}}$$
$$\mathrm{init\_hes7\_d2} \rightarrow \frac{\mathrm{init\_hes7}}{\mathrm{reg\_hes7\_fc} \cdot \mathrm{s\_RNA}}$$
$$\mathrm{init\_hes7\_d3} \rightarrow \frac{\mathrm{init\_hes7}}{\mathrm{reg\_hes7\_fc} \cdot \mathrm{s\_RNA}}$$
$$\mathrm{init\_Spdef} \rightarrow \frac{\mathrm{init\_spdef} \cdot \mathrm{p\_Spdef}}{\mathrm{d\_Spdef} \cdot \mathrm{reg\_spdef\_fc} \cdot \mathrm{s\_RNA}}$$
$$\mathrm{init\_spdef} \rightarrow \frac{\mathrm{init\_spdef}}{\mathrm{reg\_spdef\_fc} \cdot \mathrm{s\_RNA}}$$
$$\mathrm{init\_tp63} \rightarrow \frac{\mathrm{init\_tp63}}{\mathrm{reg\_tp63\_fc} \cdot \mathrm{s\_RNA}}$$
$$\mathrm{k\_hes7\_i\_spdef} \rightarrow \frac{\mathrm{init\_spdef} \cdot \mathrm{k\_hes7\_i\_spdef\_fc} \cdot \mathrm{p\_Spdef}}{\mathrm{d\_Spdef} \cdot \mathrm{reg\_spdef\_fc} \cdot \mathrm{s\_RNA}}$$
$$\mathrm{k\_ssc\_i\_spdef} \rightarrow \frac{\mathrm{init\_spdef} \cdot \mathrm{k\_ssc\_a\_spdef\_fc} \cdot \mathrm{k\_ssc\_i\_spdef\_fc} \cdot \mathrm{p\_Spdef}}{\mathrm{d\_Spdef} \cdot \mathrm{reg\_spdef\_fc} \cdot \mathrm{s\_RNA}}$$
$$\mathrm{k\_ssc\_a\_spdef} \rightarrow \frac{\mathrm{init\_spdef} \cdot \mathrm{k\_ssc\_a\_spdef\_fc} \cdot \mathrm{p\_Spdef}}{\mathrm{d\_Spdef} \cdot \mathrm{reg\_spdef\_fc} \cdot \mathrm{s\_RNA}}$$

$$\text{k\_tp63\_a\_spdef} \rightarrow \frac{\text{init\_spdef} \cdot \text{k\_tp63\_a\_spdef\_fc} \cdot \text{p\_Spdef}}{\text{d\_Spdef} \cdot \text{reg\_spdef\_fc} \cdot \text{s\_RNA}}$$

$$\text{s\_cell1} \rightarrow \text{s\_cell1\_fc} \cdot \text{s\_cell\_mean}$$

$$\text{s\_cell12} \rightarrow \text{s\_cell12\_fc} \cdot \text{s\_cell\_mean}$$

$$\text{s\_cell13} \rightarrow \text{s\_cell13\_fc} \cdot \text{s\_cell\_mean}$$

$$\text{s\_cell14} \rightarrow \text{s\_cell14\_fc} \cdot \text{s\_cell\_mean}$$

$$\text{s\_cell2} \rightarrow \text{s\_cell2\_fc} \cdot \text{s\_cell\_mean}$$

$$\text{s\_cell20} \rightarrow \text{s\_cell20\_fc} \cdot \text{s\_cell\_mean}$$

$$\text{s\_cell22} \rightarrow \text{s\_cell22\_fc} \cdot \text{s\_cell\_mean}$$

$$\text{s\_cell23} \rightarrow \text{s\_cell23\_fc} \cdot \text{s\_cell\_mean}$$

$$\text{s\_cell24} \rightarrow \text{s\_cell24\_fc} \cdot \text{s\_cell\_mean}$$

$$\text{s\_cell25} \rightarrow \text{s\_cell25\_fc} \cdot \text{s\_cell\_mean}$$

$$\text{s\_cell3} \rightarrow \text{s\_cell3\_fc} \cdot \text{s\_cell\_mean}$$

$$\text{s\_cell4} \rightarrow \text{s\_cell4\_fc} \cdot \text{s\_cell\_mean}$$

$$\text{s\_cell5} \rightarrow \text{s\_cell5\_fc} \cdot \text{s\_cell\_mean}$$

$$\text{s\_cell8} \rightarrow \text{s\_cell8\_fc} \cdot \text{s\_cell\_mean}$$

$$\text{s\_cell9} \rightarrow \text{s\_cell9\_fc} \cdot \text{s\_cell\_mean}$$

$$\text{transl\_hes4\_lof\_fc} \rightarrow 1$$

$$\text{transl\_hes5\_lof\_fc} \rightarrow 1$$

$$\text{transl\_hes7\_lof\_fc} \rightarrow 1$$

- **Local condition #23 (global condition #1):**

$$\text{ex1} \rightarrow 1$$

$$\text{ex12} \rightarrow 0$$

$$\text{ex13} \rightarrow 0$$

$$\text{ex14} \rightarrow 0$$

$$\text{ex2} \rightarrow 0$$

$$\text{ex20} \rightarrow 0$$

$$\text{ex22} \rightarrow 0$$

$$\text{ex23} \rightarrow 0$$

$$\text{ex24} \rightarrow 0$$

$$\text{ex25} \rightarrow 0$$

$$\text{ex3} \rightarrow 0$$

$$\text{ex4} \rightarrow 0$$

$$\text{ex5} \rightarrow 0$$

$$\text{ex8} \rightarrow 0$$

$$\text{ex9} \rightarrow 0$$

$$\text{init\_hes4} \rightarrow \frac{\text{init\_hes4}}{\text{reg\_hes4\_fc} \cdot \text{s\_RNA}}$$

$$\text{init\_hes4\_d} \rightarrow \frac{4 \cdot \text{init\_hes4}}{\text{d\_hes4\_prot\_fc} \cdot \text{d\_hes\_prot} \cdot \text{hes\_delay} \cdot \text{reg\_hes4\_fc} \cdot \text{s\_RNA}}$$

$$\text{init\_hes4\_d1} \rightarrow \frac{\text{init\_hes4}}{\text{reg\_hes4\_fc} \cdot \text{s\_RNA}}$$
$$\text{init\_hes4\_d2} \rightarrow \frac{\text{init\_hes4}}{\text{reg\_hes4\_fc} \cdot \text{s\_RNA}}$$
$$\text{init\_hes4\_d3} \rightarrow \frac{\text{init\_hes4}}{\text{reg\_hes4\_fc} \cdot \text{s\_RNA}}$$
$$\text{init\_hes5} \rightarrow \frac{\text{init\_hes5}}{\text{reg\_hes5\_fc} \cdot \text{s\_RNA}}$$
$$\text{init\_hes5\_d} \rightarrow \frac{4 \cdot \text{init\_hes5}}{\text{d\_hes5\_prot\_fc} \cdot \text{d\_hes\_prot} \cdot \text{hes\_delay} \cdot \text{reg\_hes5\_fc} \cdot \text{s\_RNA}}$$
$$\text{init\_hes5\_d1} \rightarrow \frac{\text{init\_hes5}}{\text{reg\_hes5\_fc} \cdot \text{s\_RNA}}$$
$$\text{init\_hes5\_d2} \rightarrow \frac{\text{init\_hes5}}{\text{reg\_hes5\_fc} \cdot \text{s\_RNA}}$$
$$\text{init\_hes5\_d3} \rightarrow \frac{\text{init\_hes5}}{\text{reg\_hes5\_fc} \cdot \text{s\_RNA}}$$
$$\text{init\_hes7} \rightarrow \frac{\text{init\_hes7}}{\text{reg\_hes7\_fc} \cdot \text{s\_RNA}}$$
$$\text{init\_hes7\_d} \rightarrow \frac{4 \cdot \text{init\_hes7}}{\text{d\_hes7\_prot\_fc} \cdot \text{d\_hes\_prot} \cdot \text{hes\_delay} \cdot \text{reg\_hes7\_fc} \cdot \text{s\_RNA}}$$
$$\text{init\_hes7\_d1} \rightarrow \frac{\text{init\_hes7}}{\text{reg\_hes7\_fc} \cdot \text{s\_RNA}}$$
$$\text{init\_hes7\_d2} \rightarrow \frac{\text{init\_hes7}}{\text{reg\_hes7\_fc} \cdot \text{s\_RNA}}$$
$$\text{init\_hes7\_d3} \rightarrow \frac{\text{init\_hes7}}{\text{reg\_hes7\_fc} \cdot \text{s\_RNA}}$$
$$\text{init\_Spdef} \rightarrow \frac{\text{init\_spdef} \cdot \text{p\_Spdef}}{\text{d\_Spdef} \cdot \text{reg\_spdef\_fc} \cdot \text{s\_RNA}}$$
$$\text{init\_spdef} \rightarrow \frac{\text{init\_spdef}}{\text{reg\_spdef\_fc} \cdot \text{s\_RNA}}$$
$$\text{init\_tp63} \rightarrow \frac{\text{init\_tp63}}{\text{reg\_tp63\_fc} \cdot \text{s\_RNA}}$$
$$\text{k\_hes7\_i\_spdef} \rightarrow \frac{\text{init\_spdef} \cdot \text{k\_hes7\_i\_spdef\_fc} \cdot \text{p\_Spdef}}{\text{d\_Spdef} \cdot \text{reg\_spdef\_fc} \cdot \text{s\_RNA}}$$
$$\text{k\_ssc\_i\_spdef} \rightarrow \frac{\text{init\_spdef} \cdot \text{k\_ssc\_a\_spdef\_fc} \cdot \text{k\_ssc\_i\_spdef\_fc} \cdot \text{p\_Spdef}}{\text{d\_Spdef} \cdot \text{reg\_spdef\_fc} \cdot \text{s\_RNA}}$$
$$\text{k\_ssc\_a\_spdef} \rightarrow \frac{\text{init\_spdef} \cdot \text{k\_ssc\_a\_spdef\_fc} \cdot \text{p\_Spdef}}{\text{d\_Spdef} \cdot \text{reg\_spdef\_fc} \cdot \text{s\_RNA}}$$
$$\text{k\_tp63\_a\_spdef} \rightarrow \frac{\text{init\_spdef} \cdot \text{k\_tp63\_a\_spdef\_fc} \cdot \text{p\_Spdef}}{\text{d\_Spdef} \cdot \text{reg\_spdef\_fc} \cdot \text{s\_RNA}}$$
$$\text{s\_cell1} \rightarrow \text{s\_cell1\_fc} \cdot \text{s\_cell\_mean}$$
$$\text{s\_cell12} \rightarrow \text{s\_cell12\_fc} \cdot \text{s\_cell\_mean}$$
$$\text{s\_cell13} \rightarrow \text{s\_cell13\_fc} \cdot \text{s\_cell\_mean}$$
$$\text{s\_cell14} \rightarrow \text{s\_cell14\_fc} \cdot \text{s\_cell\_mean}$$
$$\text{s\_cell2} \rightarrow \text{s\_cell2\_fc} \cdot \text{s\_cell\_mean}$$

$$s\_cell20 \rightarrow s\_cell20\_fc \cdot s\_cell\_mean$$
$$s\_cell22 \rightarrow s\_cell22\_fc \cdot s\_cell\_mean$$
$$s\_cell23 \rightarrow s\_cell23\_fc \cdot s\_cell\_mean$$
$$s\_cell24 \rightarrow s\_cell24\_fc \cdot s\_cell\_mean$$
$$s\_cell25 \rightarrow s\_cell25\_fc \cdot s\_cell\_mean$$
$$s\_cell3 \rightarrow s\_cell3\_fc \cdot s\_cell\_mean$$
$$s\_cell4 \rightarrow s\_cell4\_fc \cdot s\_cell\_mean$$
$$s\_cell5 \rightarrow s\_cell5\_fc \cdot s\_cell\_mean$$
$$s\_cell8 \rightarrow s\_cell8\_fc \cdot s\_cell\_mean$$
$$s\_cell9 \rightarrow s\_cell9\_fc \cdot s\_cell\_mean$$
$$transl\_hes4\_lof\_fc \rightarrow 1$$
$$transl\_hes5\_lof\_fc \rightarrow 1$$
$$transl\_hes7\_lof\_fc \rightarrow 1$$

##### 1.9.4 Experimental data and model fit

| time [min] | isc_abs [a.u.] | mcc_abs [a.u.] | ssc_abs [a.u.] |
|---|---|---|---|
| 124.03 | 47 | 57 | 71 |
| 124.03 | 59 | 58 | 70 |
| 124.03 | 49 | 52 | 79 |
| 124.03 | 55 | 47 | 64 |
| 124.03 | 68 | 71 | 58 |
| 124.03 | 57 | 87 | 80 |
| 124.03 | 64 | 64 | 84 |
| 124.03 | 59 | 58 | 84 |
| 124.03 | 67 | 59 | 67 |
| 124.03 | 67 | 81 | 63 |
| 124.03 | 44 | 59 | 72 |
| 124.03 | 70 | 67 | 58 |
| 124.03 | 55 | 56 | 54 |
| 124.03 | 71 | 69 | 67 |
| 124.03 | 74 | 86 | 87 |
| 124.03 | 39 | 50 | 44 |
| 124.03 | 80 | 70 | 71 |
| 124.03 | 37 | 55 | 51 |
| 124.03 | 67 | 58 | 43 |
| 124.03 | 58 | 46 | 62 |
| 124.03 | 61 | 33 | 37 |
| 124.03 | 38 | 47 | 25 |
| 124.03 | 41 | 43 | 23 |
| 124.03 | 63 | 36 | 13 |
| 124.03 | 50 | 39 | 41 |
| 124.03 | 62 | 43 | 13 |
| 124.03 | 55 | 55 | 43 |
| 124.03 | 46 | 60 | 43 |
| 124.03 | 45 | 49 | 43 |
| 124.03 | 56 | 53 | 32 |
| 124.03 | 65 | 43 | 52 |
| 124.03 | 47 | 47 | 51 |
| 124.03 | 49 | 42 | 36 |
| 124.03 | 84 | 64 | 38 |
| 124.03 | 77 | 86 | 71 |
| 124.03 | 49 | 45 | 52 |
| 124.03 | 65 | 65 | 63 |
| 124.03 | 81 | 72 | 66 |
| 124.03 | 76 | 64 | 42 |
| 124.03 | 66 | 44 | 58 |
| 124.03 | 103 | 105 | 6 |
| 124.03 | 30 | 45 | 35 |
| 124.03 | 47 | 50 | 44 |
| 124.03 | 34 | 52 | 27 |
| 124.03 | 61 | 68 | 45 |
| 124.03 | 40 | 47 | 53 |
| 124.03 | 87 | 81 | 65 |
| 124.03 | 101 | 93 | 47 |
| 124.03 | 82 | 81 | 58 |
| 124.03 | 91 | 72 | 56 |
| 124.03 | 57 | 52 | 32 |
| 124.03 | 76 | 64 | 48 |
| 124.03 | 76 | 50 | 56 |

**Table 3: Experimental data for the experiment Controls_Cells_aly_new, part 1.**

| time [min] | isc_abs [a.u.] | mcc_abs [a.u.] | ssc_abs [a.u.] |
|---|---|---|---|
| 124.03 | 56 | 57 | 42 |
| 124.03 | 67 | 71 | 61 |
| 124.03 | 99 | 83 | 51 |
| 124.03 | 56 | 43 | 41 |
| 124.03 | 61 | 79 | 59 |
| 124.03 | 67 | 70 | 39 |
| 124.03 | 38 | 53 | 23 |
| 124.03 | 81 | 55 | 47 |
| 124.03 | 63 | 69 | 78 |
| 124.03 | 70 | 51 | 49 |
| 124.03 | 74 | 90 | 71 |

**Table 4: Experimental data for the experiment Controls_Cells_aly_new, part 2.**

#### 1.10 Experiment: GainOfFunction_Cells_hes4_15conc

##### 1.10.1 Comments

Cell counts for RNA perturbation experiments

##### 1.10.2 Observables

The following observables are modified in this data set.

- **Observable:** isc_abs

$$\text{isc\_abs}(t) = [\text{isc}] \cdot (\text{ex12} \cdot \text{s\_cell12} + \text{ex13} \cdot \text{s\_cell13} + \text{ex14} \cdot \text{s\_cell14} + \text{ex20} \cdot \text{s\_cell20} + \text{ex22} \cdot \text{s\_cell22} + \text{ex23} \cdot \text{s\_cell23} + \text{ex24} \cdot \text{s\_cell24} + \text{ex9} \cdot \text{s\_cell9}) \quad (107)$$

$$\sigma\{\text{isc\_abs}\}(t) = \text{sd\_cell} \quad (108)$$

- **Observable:** mcc_abs

$$\text{mcc\_abs}(t) = [\text{mcc\_total}] \cdot (\text{ex12} \cdot \text{s\_cell12} + \text{ex13} \cdot \text{s\_cell13} + \text{ex14} \cdot \text{s\_cell14} + \text{ex20} \cdot \text{s\_cell20} + \text{ex22} \cdot \text{s\_cell22} + \text{ex23} \cdot \text{s\_cell23} + \text{ex24} \cdot \text{s\_cell24} + \text{ex9} \cdot \text{s\_cell9}) \quad (109)$$

$$\sigma\{\text{mcc\_abs}\}(t) = \text{sd\_cell} \quad (110)$$

- **Observable:** ssc_abs

$$\text{ssc\_abs}(t) = [\text{ssc}] \cdot (\text{ex12} \cdot \text{s\_cell12} + \text{ex13} \cdot \text{s\_cell13} + \text{ex14} \cdot \text{s\_cell14} + \text{ex20} \cdot \text{s\_cell20} + \text{ex22} \cdot \text{s\_cell22} + \text{ex23} \cdot \text{s\_cell23} + \text{ex24} \cdot \text{s\_cell24} + \text{ex9} \cdot \text{s\_cell9}) \quad (111)$$

$$\sigma\{\text{ssc\_abs}\}(t) = \text{sd\_cell} \quad (112)$$

##### 1.10.3 Experiment specific conditions

To evaluate the model for this experiment the following conditions are applied.

- **Local condition #24 (global condition #7):**

$$\text{ex12} \rightarrow 0$$
$$\text{ex13} \rightarrow 0$$
$$\text{ex14} \rightarrow 0$$
$$\text{ex20} \rightarrow 0$$
$$\text{ex22} \rightarrow 0$$
$$\text{ex23} \rightarrow 1$$
$$\text{ex24} \rightarrow 0$$
$$\text{ex9} \rightarrow 0$$
$$\text{init\_hes4} \rightarrow \frac{\text{init\_hes4} \cdot \text{init\_hes4\_gof\_lc\_fc}}{\text{reg\_hes4\_fc} \cdot \text{s\_RNA}}$$
$$\text{init\_hes4\_d} \rightarrow \frac{4 \cdot \text{init\_hes4} \cdot \text{init\_hes4\_gof\_lc\_fc}}{\text{d\_hes4\_prot\_fc} \cdot \text{d\_hes\_prot} \cdot \text{hes\_delay} \cdot \text{reg\_hes4\_fc} \cdot \text{s\_RNA}}$$
$$\text{init\_hes4\_d1} \rightarrow \frac{\text{init\_hes4} \cdot \text{init\_hes4\_gof\_lc\_fc}}{\text{reg\_hes4\_fc} \cdot \text{s\_RNA}}$$
$$\text{init\_hes4\_d2} \rightarrow \frac{\text{init\_hes4} \cdot \text{init\_hes4\_gof\_lc\_fc}}{\text{reg\_hes4\_fc} \cdot \text{s\_RNA}}$$

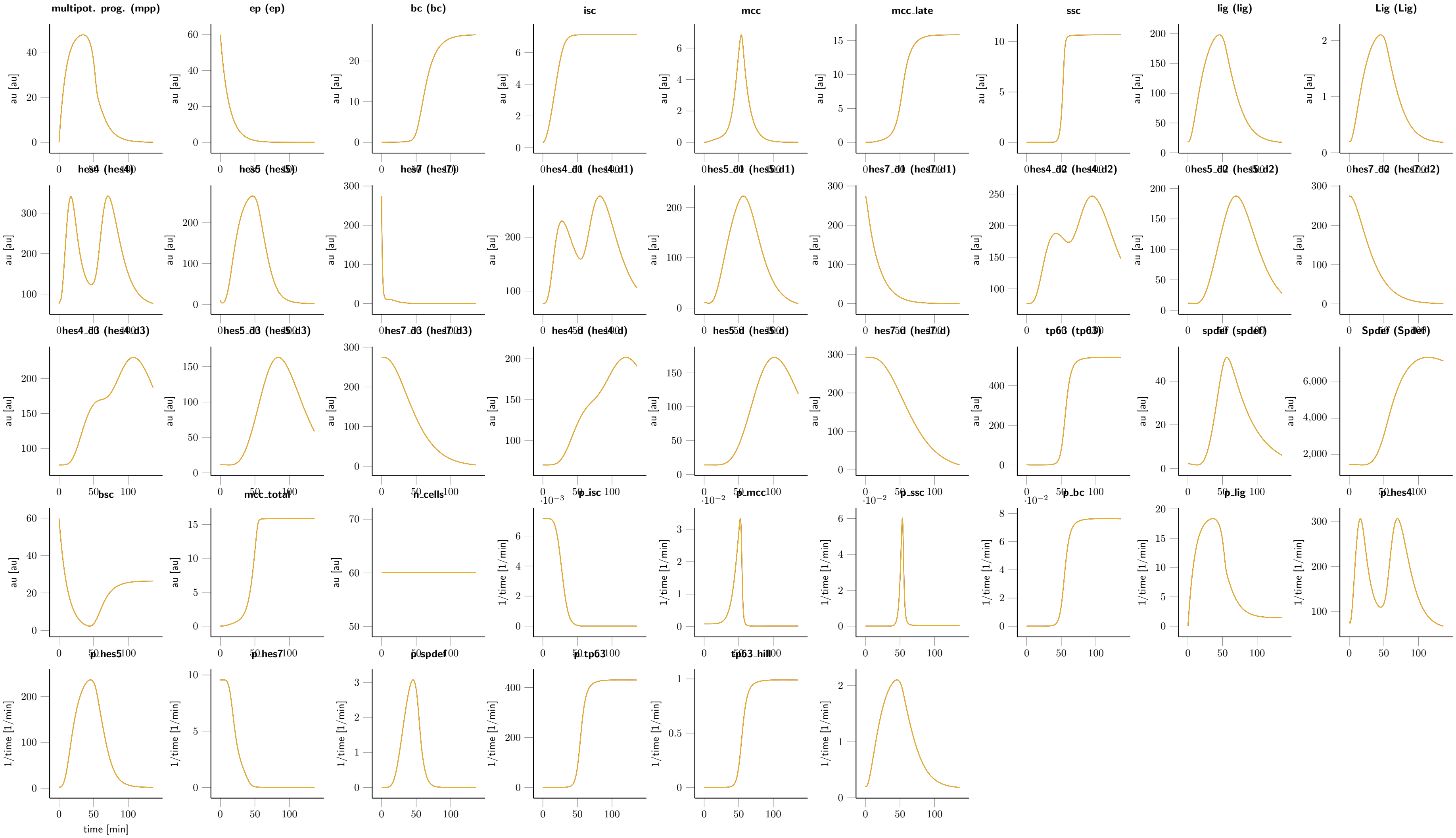

**Figure 3:** Fitted dynamics of hes4 gain of the function condition.

$$\mathrm{init\_hes4\_d3} \rightarrow \frac{\mathrm{init\_hes4} \cdot \mathrm{init\_hes4\_gof\_lc\_fc}}{\mathrm{reg\_hes4\_fc} \cdot \mathrm{s\_RNA}}$$

$$\mathrm{init\_hes5} \rightarrow \frac{\mathrm{init\_hes5}}{\mathrm{reg\_hes5\_fc} \cdot \mathrm{s\_RNA}}$$

$$\mathrm{init\_hes5\_d} \rightarrow \frac{4 \cdot \mathrm{init\_hes5}}{\mathrm{d\_hes5\_prot\_fc} \cdot \mathrm{d\_hes\_prot} \cdot \mathrm{hes\_delay} \cdot \mathrm{reg\_hes5\_fc} \cdot \mathrm{s\_RNA}}$$

$$\mathrm{init\_hes5\_d1} \rightarrow \frac{\mathrm{init\_hes5}}{\mathrm{reg\_hes5\_fc} \cdot \mathrm{s\_RNA}}$$

$$\mathrm{init\_hes5\_d2} \rightarrow \frac{\mathrm{init\_hes5}}{\mathrm{reg\_hes5\_fc} \cdot \mathrm{s\_RNA}}$$

$$\mathrm{init\_hes5\_d3} \rightarrow \frac{\mathrm{init\_hes5}}{\mathrm{reg\_hes5\_fc} \cdot \mathrm{s\_RNA}}$$

$$\mathrm{init\_hes7} \rightarrow \frac{\mathrm{init\_hes7}}{\mathrm{reg\_hes7\_fc} \cdot \mathrm{s\_RNA}}$$

$$\mathrm{init\_hes7\_d} \rightarrow \frac{4 \cdot \mathrm{init\_hes7}}{\mathrm{d\_hes7\_prot\_fc} \cdot \mathrm{d\_hes\_prot} \cdot \mathrm{hes\_delay} \cdot \mathrm{reg\_hes7\_fc} \cdot \mathrm{s\_RNA}}$$

$$\mathrm{init\_hes7\_d1} \rightarrow \frac{\mathrm{init\_hes7}}{\mathrm{reg\_hes7\_fc} \cdot \mathrm{s\_RNA}}$$

$$\mathrm{init\_hes7\_d2} \rightarrow \frac{\mathrm{init\_hes7}}{\mathrm{reg\_hes7\_fc} \cdot \mathrm{s\_RNA}}$$

$$\mathrm{init\_hes7\_d3} \rightarrow \frac{\mathrm{init\_hes7}}{\mathrm{reg\_hes7\_fc} \cdot \mathrm{s\_RNA}}$$

$$\mathrm{init\_isc} \rightarrow \mathrm{init\_isc} \cdot \mathrm{init\_isc\_gof\_lc\_hes4\_fc}$$

$$\mathrm{init\_mcc} \rightarrow \mathrm{init\_mcc} \cdot \mathrm{init\_mcc\_gof\_lc\_hes4\_fc}$$

$$\mathrm{init\_mpp} \rightarrow \mathrm{init\_mpp} \cdot \mathrm{init\_mpp\_gof\_lc\_hes4\_fc}$$

$$\mathrm{init\_Spdef} \rightarrow \frac{\mathrm{init\_spdef} \cdot \mathrm{p\_Spdef}}{\mathrm{d\_Spdef} \cdot \mathrm{reg\_spdef\_fc} \cdot \mathrm{s\_RNA}}$$

$$\mathrm{init\_spdef} \rightarrow \frac{\mathrm{init\_spdef}}{\mathrm{reg\_spdef\_fc} \cdot \mathrm{s\_RNA}}$$

$$\mathrm{init\_tp63} \rightarrow \frac{\mathrm{init\_tp63}}{\mathrm{reg\_tp63\_fc} \cdot \mathrm{s\_RNA}}$$

$$\mathrm{k\_hes7\_i\_spdef} \rightarrow \frac{\mathrm{init\_spdef} \cdot \mathrm{k\_hes7\_i\_spdef\_fc} \cdot \mathrm{p\_Spdef}}{\mathrm{d\_Spdef} \cdot \mathrm{reg\_spdef\_fc} \cdot \mathrm{s\_RNA}}$$

$$\mathrm{k\_ssc\_i\_spdef} \rightarrow \frac{\mathrm{init\_spdef} \cdot \mathrm{k\_ssc\_a\_spdef\_fc} \cdot \mathrm{k\_ssc\_i\_spdef\_fc} \cdot \mathrm{p\_Spdef}}{\mathrm{d\_Spdef} \cdot \mathrm{reg\_spdef\_fc} \cdot \mathrm{s\_RNA}}$$

$$\mathrm{k\_ssc\_a\_spdef} \rightarrow \frac{\mathrm{init\_spdef} \cdot \mathrm{k\_ssc\_a\_spdef\_fc} \cdot \mathrm{p\_Spdef}}{\mathrm{d\_Spdef} \cdot \mathrm{reg\_spdef\_fc} \cdot \mathrm{s\_RNA}}$$

$$\mathrm{k\_tp63\_a\_spdef} \rightarrow \frac{\mathrm{init\_spdef} \cdot \mathrm{k\_tp63\_a\_spdef\_fc} \cdot \mathrm{p\_Spdef}}{\mathrm{d\_Spdef} \cdot \mathrm{reg\_spdef\_fc} \cdot \mathrm{s\_RNA}}$$

$$\mathrm{s\_cell12} \rightarrow \mathrm{s\_cell12\_fc} \cdot \mathrm{s\_cell\_mean}$$

$$\mathrm{s\_cell13} \rightarrow \mathrm{s\_cell13\_fc} \cdot \mathrm{s\_cell\_mean}$$

$$\mathrm{s\_cell14} \rightarrow \mathrm{s\_cell14\_fc} \cdot \mathrm{s\_cell\_mean}$$

$$\mathrm{s\_cell20} \rightarrow \mathrm{s\_cell20\_fc} \cdot \mathrm{s\_cell\_mean}$$

$$\mathrm{s\_cell22} \rightarrow \mathrm{s\_cell22\_fc} \cdot \mathrm{s\_cell\_mean}$$

$$\mathrm{s\_cell23} \to \mathrm{s\_cell23\_fc} \cdot \mathrm{s\_cell\_mean}$$
$$\mathrm{s\_cell24} \to \mathrm{s\_cell24\_fc} \cdot \mathrm{s\_cell\_mean}$$
$$\mathrm{s\_cell9} \to \mathrm{s\_cell9\_fc} \cdot \mathrm{s\_cell\_mean}$$
$$\mathrm{transl\_hes4\_lof\_fc} \to 1$$
$$\mathrm{transl\_hes5\_lof\_fc} \to 1$$
$$\mathrm{transl\_hes7\_lof\_fc} \to 1$$

- **Local condition #25 (global condition #7):**

$$\mathrm{ex12} \to 0$$
$$\mathrm{ex13} \to 0$$
$$\mathrm{ex14} \to 0$$
$$\mathrm{ex20} \to 0$$
$$\mathrm{ex22} \to 1$$
$$\mathrm{ex23} \to 0$$
$$\mathrm{ex24} \to 0$$
$$\mathrm{ex9} \to 0$$
$$\mathrm{init\_hes4} \to \frac{\mathrm{init\_hes4} \cdot \mathrm{init\_hes4\_gof\_lc\_fc}}{\mathrm{reg\_hes4\_fc} \cdot \mathrm{s\_RNA}}$$
$$\mathrm{init\_hes4\_d} \to \frac{4 \cdot \mathrm{init\_hes4} \cdot \mathrm{init\_hes4\_gof\_lc\_fc}}{\mathrm{d\_hes4\_prot\_fc} \cdot \mathrm{d\_hes\_prot} \cdot \mathrm{hes\_delay} \cdot \mathrm{reg\_hes4\_fc} \cdot \mathrm{s\_RNA}}$$
$$\mathrm{init\_hes4\_d1} \to \frac{\mathrm{init\_hes4} \cdot \mathrm{init\_hes4\_gof\_lc\_fc}}{\mathrm{reg\_hes4\_fc} \cdot \mathrm{s\_RNA}}$$
$$\mathrm{init\_hes4\_d2} \to \frac{\mathrm{init\_hes4} \cdot \mathrm{init\_hes4\_gof\_lc\_fc}}{\mathrm{reg\_hes4\_fc} \cdot \mathrm{s\_RNA}}$$
$$\mathrm{init\_hes4\_d3} \to \frac{\mathrm{init\_hes4} \cdot \mathrm{init\_hes4\_gof\_lc\_fc}}{\mathrm{reg\_hes4\_fc} \cdot \mathrm{s\_RNA}}$$
$$\mathrm{init\_hes5} \to \frac{\mathrm{init\_hes5}}{\mathrm{reg\_hes5\_fc} \cdot \mathrm{s\_RNA}}$$
$$\mathrm{init\_hes5\_d} \to \frac{4 \cdot \mathrm{init\_hes5}}{\mathrm{d\_hes5\_prot\_fc} \cdot \mathrm{d\_hes\_prot} \cdot \mathrm{hes\_delay} \cdot \mathrm{reg\_hes5\_fc} \cdot \mathrm{s\_RNA}}$$
$$\mathrm{init\_hes5\_d1} \to \frac{\mathrm{init\_hes5}}{\mathrm{reg\_hes5\_fc} \cdot \mathrm{s\_RNA}}$$
$$\mathrm{init\_hes5\_d2} \to \frac{\mathrm{init\_hes5}}{\mathrm{reg\_hes5\_fc} \cdot \mathrm{s\_RNA}}$$
$$\mathrm{init\_hes5\_d3} \to \frac{\mathrm{init\_hes5}}{\mathrm{reg\_hes5\_fc} \cdot \mathrm{s\_RNA}}$$
$$\mathrm{init\_hes7} \to \frac{\mathrm{init\_hes7}}{\mathrm{reg\_hes7\_fc} \cdot \mathrm{s\_RNA}}$$
$$\mathrm{init\_hes7\_d} \to \frac{4 \cdot \mathrm{init\_hes7}}{\mathrm{d\_hes7\_prot\_fc} \cdot \mathrm{d\_hes\_prot} \cdot \mathrm{hes\_delay} \cdot \mathrm{reg\_hes7\_fc} \cdot \mathrm{s\_RNA}}$$
$$\mathrm{init\_hes7\_d1} \to \frac{\mathrm{init\_hes7}}{\mathrm{reg\_hes7\_fc} \cdot \mathrm{s\_RNA}}$$

$$\text{init\_hes7\_d2} \to \frac{\text{init\_hes7}}{\text{reg\_hes7\_fc} \cdot \text{s\_RNA}}$$

$$\text{init\_hes7\_d3} \to \frac{\text{init\_hes7}}{\text{reg\_hes7\_fc} \cdot \text{s\_RNA}}$$

$$\text{init\_isc} \to \text{init\_isc} \cdot \text{init\_isc\_gof\_lc\_hes4\_fc}$$

$$\text{init\_mcc} \to \text{init\_mcc} \cdot \text{init\_mcc\_gof\_lc\_hes4\_fc}$$

$$\text{init\_mpp} \to \text{init\_mpp} \cdot \text{init\_mpp\_gof\_lc\_hes4\_fc}$$

$$\text{init\_Spdef} \to \frac{\text{init\_spdef} \cdot \text{p\_Spdef}}{\text{d\_Spdef} \cdot \text{reg\_spdef\_fc} \cdot \text{s\_RNA}}$$

$$\text{init\_spdef} \to \frac{\text{init\_spdef}}{\text{reg\_spdef\_fc} \cdot \text{s\_RNA}}$$

$$\text{init\_tp63} \to \frac{\text{init\_tp63}}{\text{reg\_tp63\_fc} \cdot \text{s\_RNA}}$$

$$\text{k\_hes7\_i\_spdef} \to \frac{\text{init\_spdef} \cdot \text{k\_hes7\_i\_spdef\_fc} \cdot \text{p\_Spdef}}{\text{d\_Spdef} \cdot \text{reg\_spdef\_fc} \cdot \text{s\_RNA}}$$

$$\text{k\_ssc\_i\_spdef} \to \frac{\text{init\_spdef} \cdot \text{k\_ssc\_a\_spdef\_fc} \cdot \text{k\_ssc\_i\_spdef\_fc} \cdot \text{p\_Spdef}}{\text{d\_Spdef} \cdot \text{reg\_spdef\_fc} \cdot \text{s\_RNA}}$$

$$\text{k\_ssc\_a\_spdef} \to \frac{\text{init\_spdef} \cdot \text{k\_ssc\_a\_spdef\_fc} \cdot \text{p\_Spdef}}{\text{d\_Spdef} \cdot \text{reg\_spdef\_fc} \cdot \text{s\_RNA}}$$

$$\text{k\_tp63\_a\_spdef} \to \frac{\text{init\_spdef} \cdot \text{k\_tp63\_a\_spdef\_fc} \cdot \text{p\_Spdef}}{\text{d\_Spdef} \cdot \text{reg\_spdef\_fc} \cdot \text{s\_RNA}}$$

$$\text{s\_cell12} \to \text{s\_cell12\_fc} \cdot \text{s\_cell\_mean}$$

$$\text{s\_cell13} \to \text{s\_cell13\_fc} \cdot \text{s\_cell\_mean}$$

$$\text{s\_cell14} \to \text{s\_cell14\_fc} \cdot \text{s\_cell\_mean}$$

$$\text{s\_cell20} \to \text{s\_cell20\_fc} \cdot \text{s\_cell\_mean}$$

$$\text{s\_cell22} \to \text{s\_cell22\_fc} \cdot \text{s\_cell\_mean}$$

$$\text{s\_cell23} \to \text{s\_cell23\_fc} \cdot \text{s\_cell\_mean}$$

$$\text{s\_cell24} \to \text{s\_cell24\_fc} \cdot \text{s\_cell\_mean}$$

$$\text{s\_cell9} \to \text{s\_cell9\_fc} \cdot \text{s\_cell\_mean}$$

$$\text{transl\_hes4\_lof\_fc} \to 1$$

$$\text{transl\_hes5\_lof\_fc} \to 1$$

$$\text{transl\_hes7\_lof\_fc} \to 1$$

- **Local condition #26 (global condition #7):**

$\text{s\_cell12} \to \text{s\_cell12\_fc} \cdot \text{s\_cell\_mean}$
$\text{s\_cell13} \to \text{s\_cell13\_fc} \cdot \text{s\_cell\_mean}$
$\text{s\_cell14} \to \text{s\_cell14\_fc} \cdot \text{s\_cell\_mean}$
$\text{s\_cell20} \to \text{s\_cell20\_fc} \cdot \text{s\_cell\_mean}$
$\text{s\_cell22} \to \text{s\_cell22\_fc} \cdot \text{s\_cell\_mean}$
$\text{s\_cell23} \to \text{s\_cell23\_fc} \cdot \text{s\_cell\_mean}$
$\text{s\_cell24} \to \text{s\_cell24\_fc} \cdot \text{s\_cell\_mean}$
$\text{s\_cell9} \to \text{s\_cell9\_fc} \cdot \text{s\_cell\_mean}$
$\text{transl\_hes4\_lof\_fc} \to 1$

| time [min] | isc_abs [a.u.] | mcc_abs [a.u.] | ssc_abs [a.u.] |
|---|---|---|---|
| 124.03 | 5 | 38 | 1 |
| 124.03 | 38 | 62 | 41 |
| 124.03 | 3 | 82 | 29 |
| 124.03 | 9 | 24 | 19 |
| 124.03 | 20 | 18 | 40 |
| 124.03 | 46 | 48 | 65 |
| 124.03 | 27 | 19 | 61 |
| 124.03 | 8 | 95 | 7 |
| 124.03 | 33 | 53 | 53 |
| 124.03 | 50 | 52 | 29 |
| 124.03 | 33 | 49 | 58 |
| 124.03 | 11 | 76 | 8 |

**Table 5: Experimental data for the experiment GainOfFunction_Cells_hes4_15conc**

transl_hes5_lof_fc $\rightarrow 1$
transl_hes7_lof_fc $\rightarrow 1$

##### 1.10.4 Experimental data and model fit

#### 1.11 Experiment: GainOfFunction_Cells_hes5_15conc

##### 1.11.1 Comments

Cell counts for RNA perturbation experiments

##### 1.11.2 Observables

The following observables are modified in this data set.

- **Observable:** isc_abs

$$\mathrm{isc\_abs}(t) = [\mathrm{isc}] \cdot (\mathrm{ex12} \cdot \mathrm{s\_cell12} + \mathrm{ex13} \cdot \mathrm{s\_cell13} + \mathrm{ex14} \cdot \mathrm{s\_cell14} + \mathrm{ex20} \cdot \mathrm{s\_cell20} + \mathrm{ex22} \cdot \mathrm{s\_cell22} + \mathrm{ex23} \cdot \mathrm{s\_cell23} + \mathrm{ex24} \cdot \mathrm{s\_cell24} + \mathrm{ex9} \cdot \mathrm{s\_cell9}) \quad (113)$$
$$\sigma\{\mathrm{isc\_abs}\}(t) = \mathrm{sd\_cell} \quad (114)$$

- **Observable:** mcc_abs

$$\mathrm{mcc\_abs}(t) = [\mathrm{mcc\_total}] \cdot (\mathrm{ex12} \cdot \mathrm{s\_cell12} + \mathrm{ex13} \cdot \mathrm{s\_cell13} + \mathrm{ex14} \cdot \mathrm{s\_cell14} + \mathrm{ex20} \cdot \mathrm{s\_cell20} + \mathrm{ex22} \cdot \mathrm{s\_cell22} + \mathrm{ex23} \cdot \mathrm{s\_cell23} + \mathrm{ex24} \cdot \mathrm{s\_cell24} + \mathrm{ex9} \cdot \mathrm{s\_cell9}) \quad (115)$$
$$\sigma\{\mathrm{mcc\_abs}\}(t) = \mathrm{sd\_cell} \quad (116)$$

- **Observable:** ssc_abs

$$\mathrm{ssc\_abs}(t) = [\mathrm{ssc}] \cdot (\mathrm{ex12} \cdot \mathrm{s\_cell12} + \mathrm{ex13} \cdot \mathrm{s\_cell13} + \mathrm{ex14} \cdot \mathrm{s\_cell14} + \mathrm{ex20} \cdot \mathrm{s\_cell20} + \mathrm{ex22} \cdot \mathrm{s\_cell22} + \mathrm{ex23} \cdot \mathrm{s\_cell23} + \mathrm{ex24} \cdot \mathrm{s\_cell24} + \mathrm{ex9} \cdot \mathrm{s\_cell9}) \quad (117)$$
$$\sigma\{\mathrm{ssc\_abs}\}(t) = \mathrm{sd\_cell} \quad (118)$$

##### 1.11.3 Experiment specific conditions

To evaluate the model for this experiment the following conditions are applied.

- **Local condition #27 (global condition #6):**

s_cell12 $\rightarrow \mathrm{s\_cell12\_fc} \cdot \mathrm{s\_cell\_mean}$
$\mathrm{s\_cell13} \rightarrow \mathrm{s\_cell13\_fc} \cdot \mathrm{s\_cell\_mean}$
$\mathrm{s\_cell14} \rightarrow \mathrm{s\_cell14\_fc} \cdot \mathrm{s\_cell\_mean}$
$\mathrm{s\_cell20} \rightarrow \mathrm{s\_cell20\_fc} \cdot \mathrm{s\_cell\_mean}$
$\mathrm{s\_cell22} \rightarrow \mathrm{s\_cell22\_fc} \cdot \mathrm{s\_cell\_mean}$
$\mathrm{s\_cell23} \rightarrow \mathrm{s\_cell23\_fc} \cdot \mathrm{s\_cell\_mean}$
$\mathrm{s\_cell24} \rightarrow \mathrm{s\_cell24\_fc} \cdot \mathrm{s\_cell\_mean}$
$\mathrm{s\_cell9} \rightarrow \mathrm{s\_cell9\_fc} \cdot \mathrm{s\_cell\_mean}$
$\mathrm{transl\_hes4\_lof\_fc} \rightarrow 1$
$\mathrm{transl\_hes5\_lof\_fc} \rightarrow 1$
$\mathrm{transl\_hes7\_lof\_fc} \rightarrow 1$

- **Local condition #28 (global condition #6):**

s_cell12 $\rightarrow \mathrm{s\_cell12\_fc} \cdot \mathrm{s\_cell\_mean}$
$\mathrm{s\_cell13} \rightarrow \mathrm{s\_cell13\_fc} \cdot \mathrm{s\_cell\_mean}$

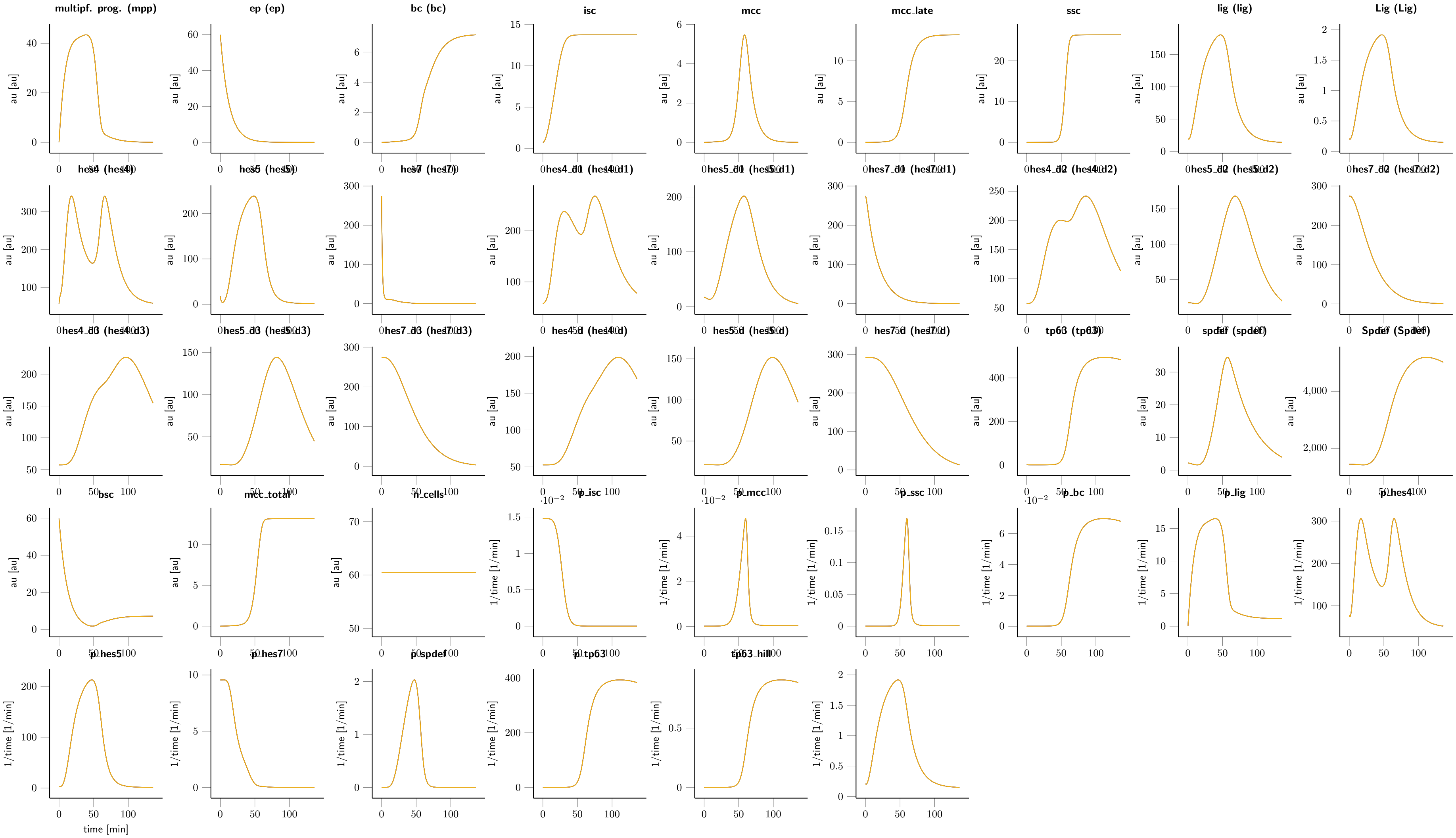

**Figure 4:** Fitted dynamics of hes5 gain of the function condition.

s_cell14 → s_cell14_fc · s_cell_mean
s_cell20 → s_cell20_fc · s_cell_mean
s_cell22 → s_cell22_fc · s_cell_mean
s_cell23 → s_cell23_fc · s_cell_mean
s_cell24 → s_cell24_fc · s_cell_mean
s_cell9 → s_cell9_fc · s_cell_mean
transl_hes4_lof_fc → 1
transl_hes5_lof_fc → 1
transl_hes7_lof_fc → 1

- **Local condition #29 (global condition #6):**

s_cell12 → s_cell12_fc · s_cell_mean
s_cell13 → s_cell13_fc · s_cell_mean
s_cell14 → s_cell14_fc · s_cell_mean
s_cell20 → s_cell20_fc · s_cell_mean
s_cell22 → s_cell22_fc · s_cell_mean
s_cell23 → s_cell23_fc · s_cell_mean
s_cell24 → s_cell24_fc · s_cell_mean
s_cell9 → s_cell9_fc · s_cell_mean
transl_hes4_lof_fc → 1
transl_hes5_lof_fc → 1
transl_hes7_lof_fc → 1

##### 1.11.4 Experimental data and model fit

| time [min] | isc_abs [a.u.] | mcc_abs [a.u.] | ssc_abs [a.u.] |
|---|---|---|---|
| 124.03 | 10 | 27 | 88 |
| 124.03 | 61 | 56 | 107 |
| 124.03 | 62 | 40 | 90 |
| 124.03 | 44 | 49 | 85 |
| 124.03 | 17 | 36 | 82 |
| 124.03 | 4 | 43 | 23 |
| 124.03 | 24 | 27 | 21 |
| 124.03 | 15 | 23 | 35 |
| 124.03 | 49 | 35 | 26 |
| 124.03 | 12 | 26 | 47 |
| 124.03 | 30 | 31 | 46 |
| 124.03 | 2 | 29 | 45 |
| 124.03 | 19 | 26 | 41 |
| 124.03 | 15 | 27 | 59 |

**Table 6: Experimental data for the experiment GainOfFunction_Cells_hes5_15conc**

#### 1.12 Experiment: GainOfFunction_Cells_hes7_15conc

##### 1.12.1 Comments

Cell counts for RNA perturbation experiments

##### 1.12.2 Observables

The following observables are modified in this data set.

- **Observable:** isc_abs

$$\text{isc\_abs}(t) = [\text{isc}] \cdot (\text{ex12} \cdot \text{s\_cell12} + \text{ex13} \cdot \text{s\_cell13} + \text{ex14} \cdot \text{s\_cell14} + \text{ex20} \cdot \text{s\_cell20} + \text{ex22} \cdot \text{s\_cell22} + \text{ex23} \cdot \text{s\_cell23} + \text{ex24} \cdot \text{s\_cell24} + \text{ex9} \cdot \text{s\_cell9}) \tag{119}$$
$$\sigma\{\text{isc\_abs}\}(t) = \text{sd\_cell} \tag{120}$$

- **Observable:** mcc_abs

$$\text{mcc\_abs}(t) = [\text{mcc\_total}] \cdot (\text{ex12} \cdot \text{s\_cell12} + \text{ex13} \cdot \text{s\_cell13} + \text{ex14} \cdot \text{s\_cell14} + \text{ex20} \cdot \text{s\_cell20} + \text{ex22} \cdot \text{s\_cell22} + \text{ex23} \cdot \text{s\_cell23} + \text{ex24} \cdot \text{s\_cell24} + \text{ex9} \cdot \text{s\_cell9}) \tag{121}$$
$$\sigma\{\text{mcc\_abs}\}(t) = \text{sd\_cell} \tag{122}$$

- **Observable:** ssc_abs

$$\text{ssc\_abs}(t) = [\text{ssc}] \cdot (\text{ex12} \cdot \text{s\_cell12} + \text{ex13} \cdot \text{s\_cell13} + \text{ex14} \cdot \text{s\_cell14} + \text{ex20} \cdot \text{s\_cell20} + \text{ex22} \cdot \text{s\_cell22} + \text{ex23} \cdot \text{s\_cell23} + \text{ex24} \cdot \text{s\_cell24} + \text{ex9} \cdot \text{s\_cell9}) \tag{123}$$
$$\sigma\{\text{ssc\_abs}\}(t) = \text{sd\_cell} \tag{124}$$

##### 1.12.3 Experiment specific conditions

To evaluate the model for this experiment the following conditions are applied.

- **Local condition #30 (global condition #5):**

s_cell12 $\rightarrow$ s_cell12_fc $\cdot$ s_cell_mean
s_cell13 $\rightarrow$ s_cell13_fc $\cdot$ s_cell_mean
s_cell14 $\rightarrow$ s_cell14_fc $\cdot$ s_cell_mean
s_cell20 $\rightarrow$ s_cell20_fc $\cdot$ s_cell_mean
s_cell22 $\rightarrow$ s_cell22_fc $\cdot$ s_cell_mean
s_cell23 $\rightarrow$ s_cell23_fc $\cdot$ s_cell_mean
s_cell24 $\rightarrow$ s_cell24_fc $\cdot$ s_cell_mean
s_cell9 $\rightarrow$ s_cell9_fc $\cdot$ s_cell_mean
transl_hes4_lof_fc $\rightarrow$ 1
transl_hes5_lof_fc $\rightarrow$ 1
transl_hes7_lof_fc $\rightarrow$ 1

- **Local condition #31 (global condition #5):**

s_cell12 $\rightarrow$ s_cell12_fc $\cdot$ s_cell_mean
s_cell13 $\rightarrow$ s_cell13_fc $\cdot$ s_cell_mean

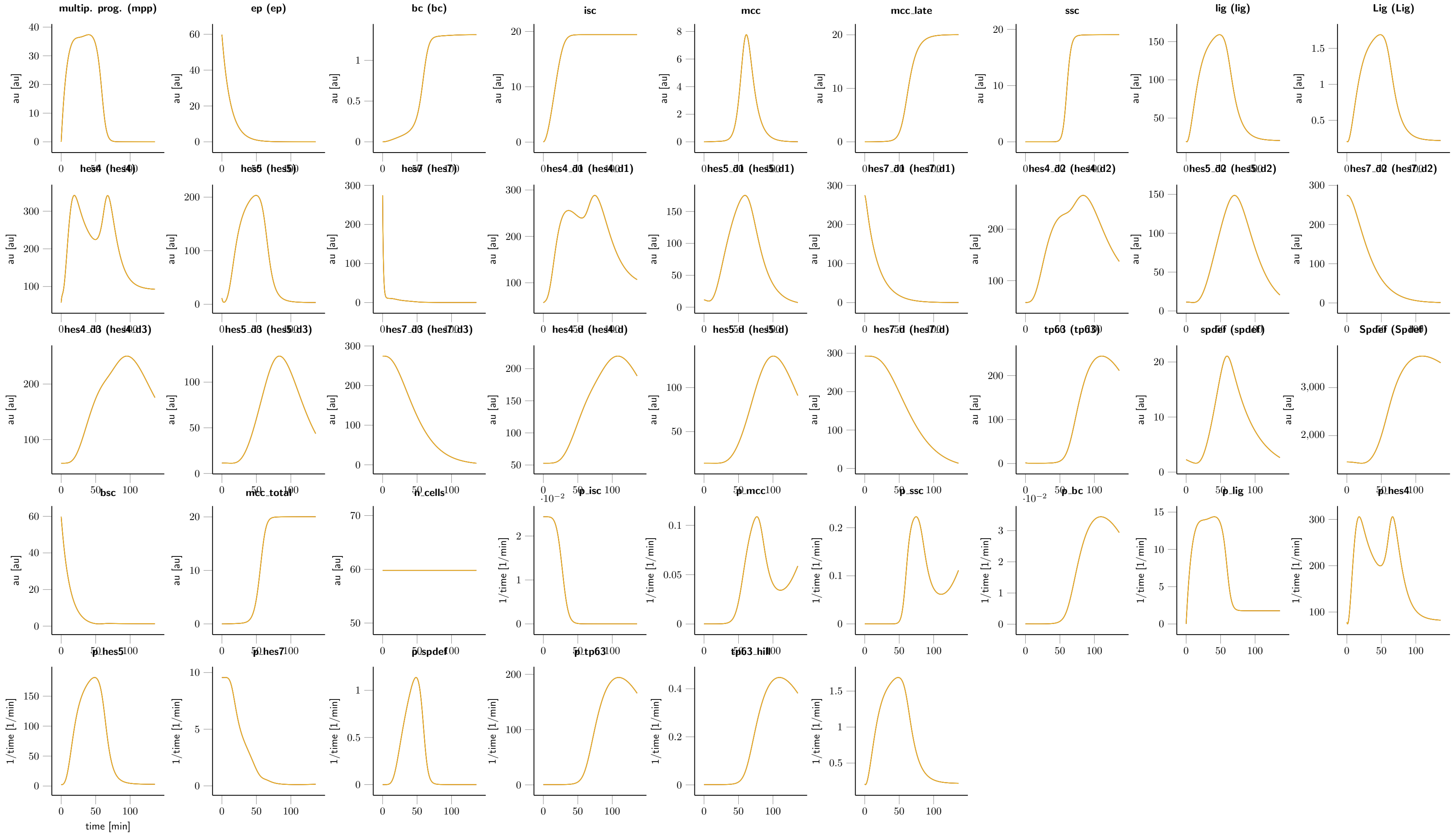

**Figure 5:** Fitted dynamics of hes7 gain of the function condition.

s_cell14 → s_cell14_fc · s_cell_mean
s_cell20 → s_cell20_fc · s_cell_mean
s_cell22 → s_cell22_fc · s_cell_mean
s_cell23 → s_cell23_fc · s_cell_mean
s_cell24 → s_cell24_fc · s_cell_mean
s_cell9 → s_cell9_fc · s_cell_mean
transl_hes4_lof_fc → 1
transl_hes5_lof_fc → 1
transl_hes7_lof_fc → 1

- **Local condition #32 (global condition #5):**

s_cell12 → s_cell12_fc · s_cell_mean
s_cell13 → s_cell13_fc · s_cell_mean
s_cell14 → s_cell14_fc · s_cell_mean
s_cell20 → s_cell20_fc · s_cell_mean
s_cell22 → s_cell22_fc · s_cell_mean
s_cell23 → s_cell23_fc · s_cell_mean
s_cell24 → s_cell24_fc · s_cell_mean
s_cell9 → s_cell9_fc · s_cell_mean
transl_hes4_lof_fc → 1
transl_hes5_lof_fc → 1
transl_hes7_lof_fc → 1

##### 1.12.4 Experimental data and model fit

| time [min] | isc_abs [a.u.] | mcc_abs [a.u.] | ssc_abs [a.u.] |
|---|---|---|---|
| 124.03 | 14 | 29 | 92 |
| 124.03 | 35 | 40 | 77 |
| 124.03 | 27 | 55 | 71 |
| 124.03 | 34 | 73 | 96 |
| 124.03 | 21 | 38 | 61 |
| 124.03 | 52 | 78 | 49 |
| 124.03 | 41 | 63 | 20 |
| 124.03 | 34 | 36 | 48 |
| 124.03 | 60 | 97 | 71 |
| 124.03 | 21 | 46 | 33 |
| 124.03 | 21 | 29 | 23 |
| 124.03 | 31 | 32 | 28 |
| 124.03 | 67 | 53 | 1 |
| 124.03 | 37 | 48 | 32 |

**Table 7: Experimental data for the experiment GainOfFunction_Cells_hes7_15conc**

#### 1.13 Experiment: LossOfFunction_Cells_hes4

##### 1.13.1 Comments

Cell counts for RNA perturbation experiments

##### 1.13.2 Observables

The following observables are modified in this data set.

- **Observable:** isc_abs

$$\text{isc\_abs}(t) = [\text{isc}] \cdot (\text{ex1} \cdot \text{s\_cell1} + \text{ex2} \cdot \text{s\_cell2} + \text{ex25} \cdot \text{s\_cell25} + \text{ex3} \cdot \text{s\_cell3} + \text{ex4} \cdot \text{s\_cell4} + \text{ex5} \cdot \text{s\_cell5} + \text{ex8} \cdot \text{s\_cell8}) \quad (125)$$
$$\sigma\{\text{isc\_abs}\}(t) = \text{sd\_cell} \quad (126)$$

- **Observable:** mcc_abs

$$\text{mcc\_abs}(t) = [\text{mcc\_total}] \cdot (\text{ex1} \cdot \text{s\_cell1} + \text{ex2} \cdot \text{s\_cell2} + \text{ex25} \cdot \text{s\_cell25} + \text{ex3} \cdot \text{s\_cell3} + \text{ex4} \cdot \text{s\_cell4} + \text{ex5} \cdot \text{s\_cell5} + \text{ex8} \cdot \text{s\_cell8}) \quad (127)$$
$$\sigma\{\text{mcc\_abs}\}(t) = \text{sd\_cell} \quad (128)$$

- **Observable:** ssc_abs

$$\text{ssc\_abs}(t) = [\text{ssc}] \cdot (\text{ex1} \cdot \text{s\_cell1} + \text{ex2} \cdot \text{s\_cell2} + \text{ex25} \cdot \text{s\_cell25} + \text{ex3} \cdot \text{s\_cell3} + \text{ex4} \cdot \text{s\_cell4} + \text{ex5} \cdot \text{s\_cell5} + \text{ex8} \cdot \text{s\_cell8}) \quad (129)$$
$$\sigma\{\text{ssc\_abs}\}(t) = \text{sd\_cell} \quad (130)$$

##### 1.13.3 Experiment specific conditions

To evaluate the model for this experiment the following conditions are applied.

- **Local condition #33 (global condition #4):**

s_cell1 $\rightarrow$ s_cell1_fc $\cdot$ s_cell_mean
s_cell2 $\rightarrow$ s_cell2_fc $\cdot$ s_cell_mean
s_cell25 $\rightarrow$ s_cell25_fc $\cdot$ s_cell_mean
s_cell3 $\rightarrow$ s_cell3_fc $\cdot$ s_cell_mean
s_cell4 $\rightarrow$ s_cell4_fc $\cdot$ s_cell_mean
s_cell5 $\rightarrow$ s_cell5_fc $\cdot$ s_cell_mean
s_cell8 $\rightarrow$ s_cell8_fc $\cdot$ s_cell_mean
transl_hes5_lof_fc $\rightarrow$ 1
transl_hes7_lof_fc $\rightarrow$ 1

- **Local condition #34 (global condition #4):**

s_cell1 $\rightarrow$ s_cell1_fc $\cdot$ s_cell_mean
s_cell2 $\rightarrow$ s_cell2_fc $\cdot$ s_cell_mean
s_cell25 $\rightarrow$ s_cell25_fc $\cdot$ s_cell_mean
s_cell3 $\rightarrow$ s_cell3_fc $\cdot$ s_cell_mean

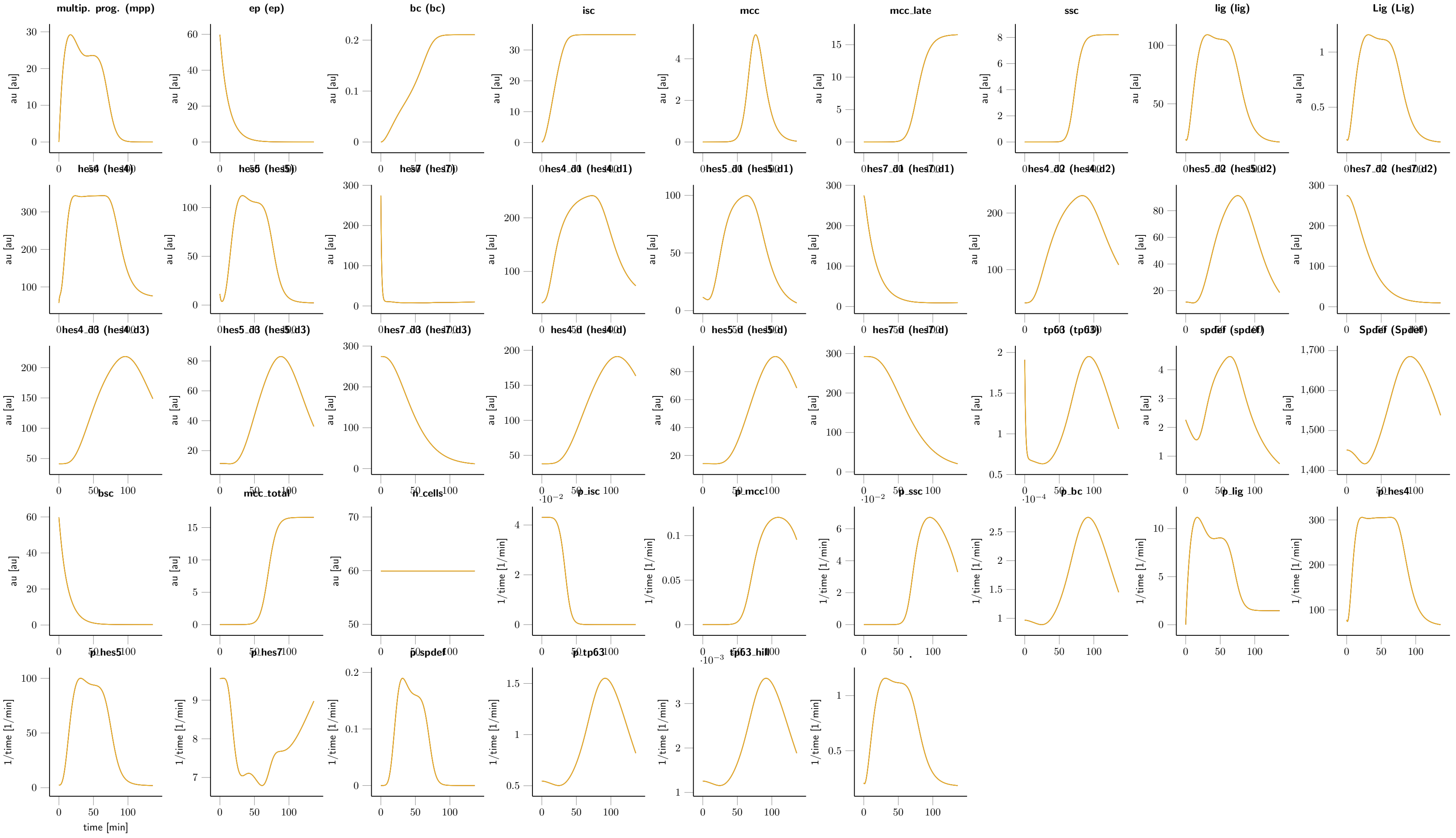

**Figure 6:** Fitted dynamics of hes4 loss of the function condition.

| time [min] | isc_abs [a.u.] | mcc_abs [a.u.] | ssc_abs [a.u.] |
|---|---|---|---|
| 124.03 | 41 | 56 | 9 |
| 124.03 | 73 | 23 | 10 |
| 124.03 | 122 | 26 | 23 |
| 124.03 | 127 | 96 | 7 |
| 124.03 | 82 | 60 | 12 |
| 124.03 | 87 | 40 | g 13 |
| 124.03 | 100 | 42 | 25 |
| 124.03 | 124 | 48 | 21 |
| 124.03 | 76 | 48 | 18 |
| 124.03 | 109 | 30 | 13 |
| 124.03 | 116 | 29 | 12 |
| 124.03 | 51 | 44 | 23 |
| 124.03 | 115 | 3 | 22 |

**Table 8: Experimental data for the experiment LossOfFunction_Cells_hes4**

$\text{s\_cell4} \rightarrow \text{s\_cell4\_fc} \cdot \text{s\_cell\_mean}$
$\text{s\_cell5} \rightarrow \text{s\_cell5\_fc} \cdot \text{s\_cell\_mean}$
$\text{s\_cell8} \rightarrow \text{s\_cell8\_fc} \cdot \text{s\_cell\_mean}$
$\text{transl\_hes5\_lof\_fc} \rightarrow 1$
$\text{transl\_hes7\_lof\_fc} \rightarrow 1$

- **Local condition #35 (global condition #4):**

s_cell1 $\rightarrow \text{s\_cell1\_fc} \cdot \text{s\_cell\_mean}$
$\text{s\_cell2} \rightarrow \text{s\_cell2\_fc} \cdot \text{s\_cell\_mean}$
$\text{s\_cell25} \rightarrow \text{s\_cell25\_fc} \cdot \text{s\_cell\_mean}$
$\text{s\_cell3} \rightarrow \text{s\_cell3\_fc} \cdot \text{s\_cell\_mean}$
$\text{s\_cell4} \rightarrow \text{s\_cell4\_fc} \cdot \text{s\_cell\_mean}$
$\text{s\_cell5} \rightarrow \text{s\_cell5\_fc} \cdot \text{s\_cell\_mean}$
$\text{s\_cell8} \rightarrow \text{s\_cell8\_fc} \cdot \text{s\_cell\_mean}$
$\text{transl\_hes5\_lof\_fc} \rightarrow 1$
$\text{transl\_hes7\_lof\_fc} \rightarrow 1$

##### 1.13.4 Experimental data and model fit

##### 1.14 Experiment: LossOfFunction_Cells_hes5

###### 1.14.1 Comments

Cell counts for RNA perturbation experiments

###### 1.14.2 Observables

The following observables are modified in this data set.

- **Observable:** isc_abs

$$\text{isc\_abs(t)} = [\text{isc}] \cdot (\text{ex1} \cdot \text{s\_cell1} + \text{ex2} \cdot \text{s\_cell2} + \text{ex25} \cdot \text{s\_cell25} + \text{ex3} \cdot \text{s\_cell3} + \text{ex4} \cdot \text{s\_cell4} + \text{ex5} \cdot \text{s\_cell5} + \text{ex8} \cdot \text{s\_cell8}) \tag{131}$$
$$\sigma\{\text{isc\_abs}\}(\text{t}) = \text{sd\_cell} \tag{132}$$

- **Observable:** mcc_abs

$$\text{mcc\_abs(t)} = [\text{mcc\_total}] \cdot (\text{ex1} \cdot \text{s\_cell1} + \text{ex2} \cdot \text{s\_cell2} + \text{ex25} \cdot \text{s\_cell25} + \text{ex3} \cdot \text{s\_cell3} + \text{ex4} \cdot \text{s\_cell4} + \text{ex5} \cdot \text{s\_cell5} + \text{ex8} \cdot \text{s\_cell8}) \tag{133}$$
$$\sigma\{\text{mcc\_abs}\}(\text{t}) = \text{sd\_cell} \tag{134}$$

- **Observable:** ssc_abs

$$\text{ssc\_abs(t)} = [\text{ssc}] \cdot (\text{ex1} \cdot \text{s\_cell1} + \text{ex2} \cdot \text{s\_cell2} + \text{ex25} \cdot \text{s\_cell25} + \text{ex3} \cdot \text{s\_cell3} + \text{ex4} \cdot \text{s\_cell4} + \text{ex5} \cdot \text{s\_cell5} + \text{ex8} \cdot \text{s\_cell8}) \tag{135}$$
$$\sigma\{\text{ssc\_abs}\}(\text{t}) = \text{sd\_cell} \tag{136}$$

###### 1.14.3 Experiment specific conditions

To evaluate the model for this experiment the following conditions are applied.

- **Local condition #36 (global condition #3):**

s_cell1 $\rightarrow \text{s\_cell1\_fc} \cdot \text{s\_cell\_mean}$
$\text{s\_cell2} \rightarrow \text{s\_cell2\_fc} \cdot \text{s\_cell\_mean}$
$\text{s\_cell25} \rightarrow \text{s\_cell25\_fc} \cdot \text{s\_cell\_mean}$
$\text{s\_cell3} \rightarrow \text{s\_cell3\_fc} \cdot \text{s\_cell\_mean}$
$\text{s\_cell4} \rightarrow \text{s\_cell4\_fc} \cdot \text{s\_cell\_mean}$
$\text{s\_cell5} \rightarrow \text{s\_cell5\_fc} \cdot \text{s\_cell\_mean}$
$\text{s\_cell8} \rightarrow \text{s\_cell8\_fc} \cdot \text{s\_cell\_mean}$
$\text{transl\_hes4\_lof\_fc} \rightarrow 1$
$\text{transl\_hes7\_lof\_fc} \rightarrow 1$

- **Local condition #37 (global condition #3):**

$$\text{ex1} \rightarrow 0$$
$$\text{ex2} \rightarrow 1$$
$$\text{ex25} \rightarrow 0$$
$$\text{ex3} \rightarrow 0$$

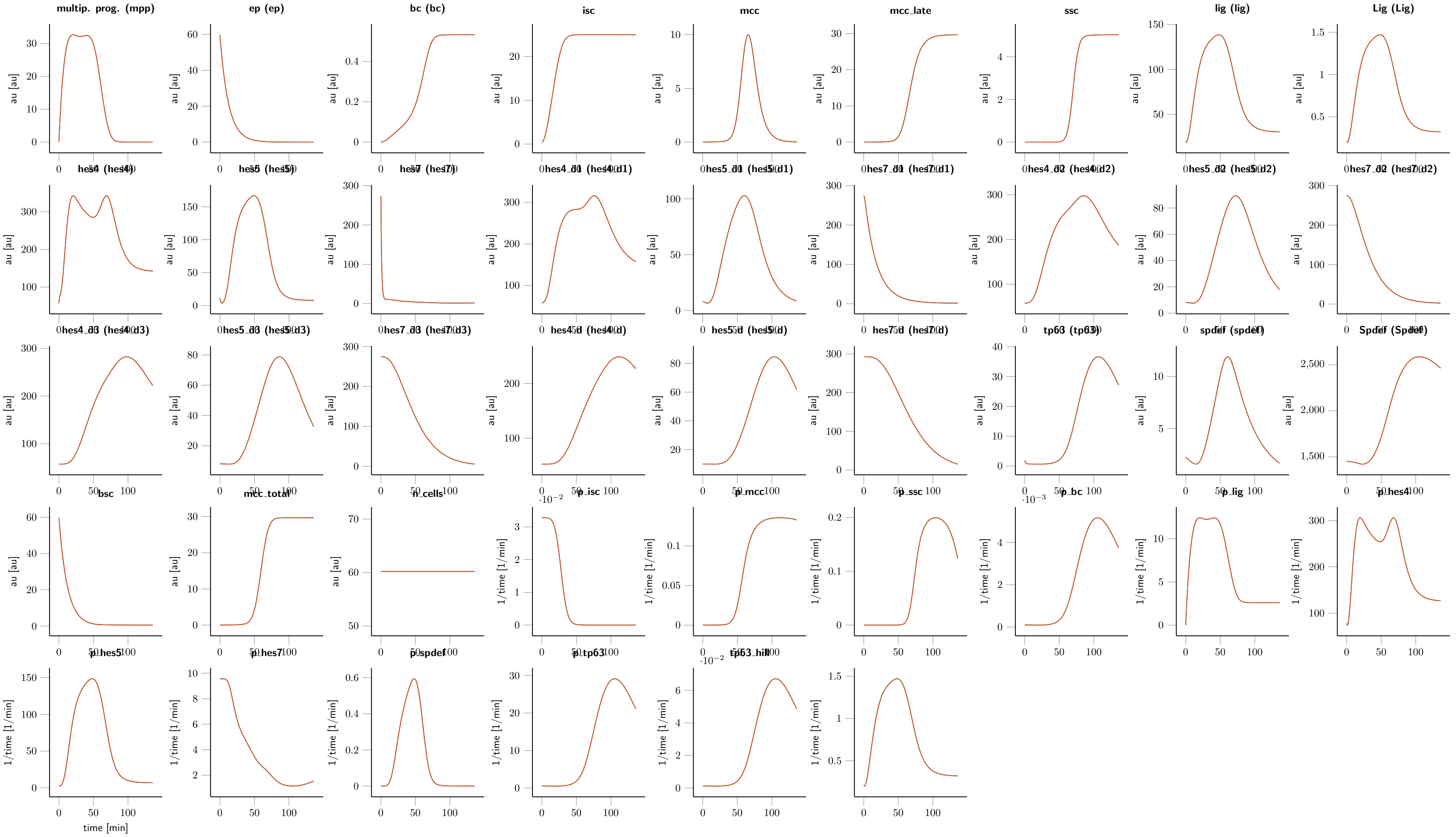

**Figure 7:** Fitted dynamics of hes5 loss of the function condition.

$$\mathrm{ex4} \to 0$$
$$\mathrm{ex5} \to 0$$
$$\mathrm{ex8} \to 0$$
$$\mathrm{init\_hes4} \to \frac{\mathrm{init\_hes4}}{\mathrm{reg\_hes4\_fc} \cdot \mathrm{s\_RNA}}$$
$$\mathrm{init\_hes4\_d} \to \frac{4 \cdot \mathrm{init\_hes4}}{\mathrm{d\_hes4\_prot\_fc} \cdot \mathrm{d\_hes\_prot} \cdot \mathrm{hes\_delay} \cdot \mathrm{reg\_hes4\_fc} \cdot \mathrm{s\_RNA}}$$
$$\mathrm{init\_hes4\_d1} \to \frac{\mathrm{init\_hes4}}{\mathrm{reg\_hes4\_fc} \cdot \mathrm{s\_RNA}}$$
$$\mathrm{init\_hes4\_d2} \to \frac{\mathrm{init\_hes4}}{\mathrm{reg\_hes4\_fc} \cdot \mathrm{s\_RNA}}$$
$$\mathrm{init\_hes4\_d3} \to \frac{\mathrm{init\_hes4}}{\mathrm{reg\_hes4\_fc} \cdot \mathrm{s\_RNA}}$$
$$\mathrm{init\_hes5} \to \frac{\mathrm{init\_hes5}}{\mathrm{reg\_hes5\_fc} \cdot \mathrm{s\_RNA}}$$
$$\mathrm{init\_hes5\_d} \to \frac{4 \cdot \mathrm{init\_hes5} \cdot \mathrm{transl\_hes5\_lof\_fc}}{\mathrm{d\_hes5\_prot\_fc} \cdot \mathrm{d\_hes\_prot} \cdot \mathrm{hes\_delay} \cdot \mathrm{reg\_hes5\_fc} \cdot \mathrm{s\_RNA}}$$
$$\mathrm{init\_hes5\_d1} \to \frac{\mathrm{init\_hes5} \cdot \mathrm{transl\_hes5\_lof\_fc}}{\mathrm{reg\_hes5\_fc} \cdot \mathrm{s\_RNA}}$$
$$\mathrm{init\_hes5\_d2} \to \frac{\mathrm{init\_hes5} \cdot \mathrm{transl\_hes5\_lof\_fc}}{\mathrm{reg\_hes5\_fc} \cdot \mathrm{s\_RNA}}$$
$$\mathrm{init\_hes5\_d3} \to \frac{\mathrm{init\_hes5} \cdot \mathrm{transl\_hes5\_lof\_fc}}{\mathrm{reg\_hes5\_fc} \cdot \mathrm{s\_RNA}}$$
$$\mathrm{init\_hes7} \to \frac{\mathrm{init\_hes7}}{\mathrm{reg\_hes7\_fc} \cdot \mathrm{s\_RNA}}$$
$$\mathrm{init\_hes7\_d} \to \frac{4 \cdot \mathrm{init\_hes7}}{\mathrm{d\_hes7\_prot\_fc} \cdot \mathrm{d\_hes\_prot} \cdot \mathrm{hes\_delay} \cdot \mathrm{reg\_hes7\_fc} \cdot \mathrm{s\_RNA}}$$
$$\mathrm{init\_hes7\_d1} \to \frac{\mathrm{init\_hes7}}{\mathrm{reg\_hes7\_fc} \cdot \mathrm{s\_RNA}}$$
$$\mathrm{init\_hes7\_d2} \to \frac{\mathrm{init\_hes7}}{\mathrm{reg\_hes7\_fc} \cdot \mathrm{s\_RNA}}$$
$$\mathrm{init\_hes7\_d3} \to \frac{\mathrm{init\_hes7}}{\mathrm{reg\_hes7\_fc} \cdot \mathrm{s\_RNA}}$$
$$\mathrm{init\_isc} \to \mathrm{init\_isc} \cdot \mathrm{init\_isc\_lof\_hes5\_fc}$$
$$\mathrm{init\_mcc} \to \mathrm{init\_mcc} \cdot \mathrm{init\_mcc\_lof\_hes5\_fc}$$
$$\mathrm{init\_mpp} \to \mathrm{init\_mpp} \cdot \mathrm{init\_mpp\_lof\_hes5\_fc}$$
$$\mathrm{init\_Spdef} \to \frac{\mathrm{init\_spdef} \cdot \mathrm{p\_Spdef}}{\mathrm{d\_Spdef} \cdot \mathrm{reg\_spdef\_fc} \cdot \mathrm{s\_RNA}}$$
$$\mathrm{init\_spdef} \to \frac{\mathrm{init\_spdef}}{\mathrm{reg\_spdef\_fc} \cdot \mathrm{s\_RNA}}$$
$$\mathrm{init\_tp63} \to \frac{\mathrm{init\_tp63}}{\mathrm{reg\_tp63\_fc} \cdot \mathrm{s\_RNA}}$$
$$\mathrm{k\_hes7\_i\_spdef} \to \frac{\mathrm{init\_spdef} \cdot \mathrm{k\_hes7\_i\_spdef\_fc} \cdot \mathrm{p\_Spdef}}{\mathrm{d\_Spdef} \cdot \mathrm{reg\_spdef\_fc} \cdot \mathrm{s\_RNA}}$$

| time [min] | isc_abs [a.u.] | mcc_abs [a.u.] | ssc_abs [a.u.] |
|---|---|---|---|
| 124.03 | 70 | 77 | 14 |
| 124.03 | 89 | 91 | 24 |
| 124.03 | 76 | 103 | 16 |
| 124.03 | 90 | 103 | 3 |
| 124.03 | 73 | 108 | 11 |
| 124.03 | 64 | 107 | 25 |
| 124.03 | 67 | 80 | 40 |
| 124.03 | 109 | 127 | 0 |
| 124.03 | 106 | 90 | 12 |

**Table 9: Experimental data for the experiment LossOfFunction_Cells_hes5**

$$\text{k\_ssc\_i\_spdef} \rightarrow \frac{\text{init\_spdef} \cdot \text{k\_ssc\_a\_spdef\_fc} \cdot \text{k\_ssc\_i\_spdef\_fc} \cdot \text{p\_Spdef}}{\text{d\_Spdef} \cdot \text{reg\_spdef\_fc} \cdot \text{s\_RNA}}$$

$$\text{k\_ssc\_a\_spdef} \rightarrow \frac{\text{init\_spdef} \cdot \text{k\_ssc\_a\_spdef\_fc} \cdot \text{p\_Spdef}}{\text{d\_Spdef} \cdot \text{reg\_spdef\_fc} \cdot \text{s\_RNA}}$$

$$\text{k\_tp63\_a\_spdef} \rightarrow \frac{\text{init\_spdef} \cdot \text{k\_tp63\_a\_spdef\_fc} \cdot \text{p\_Spdef}}{\text{d\_Spdef} \cdot \text{reg\_spdef\_fc} \cdot \text{s\_RNA}}$$

$$\text{s\_cell1} \rightarrow \text{s\_cell1\_fc} \cdot \text{s\_cell\_mean}$$

$$\text{s\_cell2} \rightarrow \text{s\_cell2\_fc} \cdot \text{s\_cell\_mean}$$

$$\text{s\_cell25} \rightarrow \text{s\_cell25\_fc} \cdot \text{s\_cell\_mean}$$

$$\text{s\_cell3} \rightarrow \text{s\_cell3\_fc} \cdot \text{s\_cell\_mean}$$

$$\text{s\_cell4} \rightarrow \text{s\_cell4\_fc} \cdot \text{s\_cell\_mean}$$

$$\text{s\_cell5} \rightarrow \text{s\_cell5\_fc} \cdot \text{s\_cell\_mean}$$

$$\text{s\_cell8} \rightarrow \text{s\_cell8\_fc} \cdot \text{s\_cell\_mean}$$

$$\text{transl\_hes4\_lof\_fc} \rightarrow 1$$

$$\text{transl\_hes7\_lof\_fc} \rightarrow 1$$

##### 1.14.4 Experimental data and model fit

#### 1.15 Experiment: LossOfFunction_Cells_hes7

##### 1.15.1 Comments

Cell counts for RNA perturbation experiments

##### 1.15.2 Observables

The following observables are modified in this data set.

- **Observable:** isc_abs

$$\text{isc\_abs}(t) = [\text{isc}] \cdot (\text{ex1} \cdot \text{s\_cell1} + \text{ex2} \cdot \text{s\_cell2} + \text{ex25} \cdot \text{s\_cell25} + \text{ex3} \cdot \text{s\_cell3} + \text{ex4} \cdot \text{s\_cell4} + \text{ex5} \cdot \text{s\_cell5} + \text{ex8} \cdot \text{s\_cell8}) \quad (137)$$
$$\sigma\{\text{isc\_abs}\}(t) = \text{sd\_cell} \quad (138)$$

- **Observable:** mcc_abs

$$\text{mcc\_abs}(t) = [\text{mcc\_total}] \cdot (\text{ex1} \cdot \text{s\_cell1} + \text{ex2} \cdot \text{s\_cell2} + \text{ex25} \cdot \text{s\_cell25} + \text{ex3} \cdot \text{s\_cell3} + \text{ex4} \cdot \text{s\_cell4} + \text{ex5} \cdot \text{s\_cell5} + \text{ex8} \cdot \text{s\_cell8}) \quad (139)$$
$$\sigma\{\text{mcc\_abs}\}(t) = \text{sd\_cell} \quad (140)$$

- **Observable:** ssc_abs

$$\text{ssc\_abs}(t) = [\text{ssc}] \cdot (\text{ex1} \cdot \text{s\_cell1} + \text{ex2} \cdot \text{s\_cell2} + \text{ex25} \cdot \text{s\_cell25} + \text{ex3} \cdot \text{s\_cell3} + \text{ex4} \cdot \text{s\_cell4} + \text{ex5} \cdot \text{s\_cell5} + \text{ex8} \cdot \text{s\_cell8}) \quad (141)$$
$$\sigma\{\text{ssc\_abs}\}(t) = \text{sd\_cell} \quad (142)$$

##### 1.15.3 Experiment specific conditions

To evaluate the model for this experiment the following conditions are applied.

- **Local condition #38 (global condition #2):**

s_cell1 $\rightarrow$ s_cell1_fc $\cdot$ s_cell_mean
s_cell2 $\rightarrow$ s_cell2_fc $\cdot$ s_cell_mean
s_cell25 $\rightarrow$ s_cell25_fc $\cdot$ s_cell_mean
s_cell3 $\rightarrow$ s_cell3_fc $\cdot$ s_cell_mean
s_cell4 $\rightarrow$ s_cell4_fc $\cdot$ s_cell_mean
s_cell5 $\rightarrow$ s_cell5_fc $\cdot$ s_cell_mean
s_cell8 $\rightarrow$ s_cell8_fc $\cdot$ s_cell_mean
transl_hes4_lof_fc $\rightarrow$ 1
transl_hes5_lof_fc $\rightarrow$ 1

- **Local condition #39 (global condition #2):**

s_cell1 $\rightarrow$ s_cell1_fc $\cdot$ s_cell_mean
s_cell2 $\rightarrow$ s_cell2_fc $\cdot$ s_cell_mean
s_cell25 $\rightarrow$ s_cell25_fc $\cdot$ s_cell_mean
s_cell3 $\rightarrow$ s_cell3_fc $\cdot$ s_cell_mean

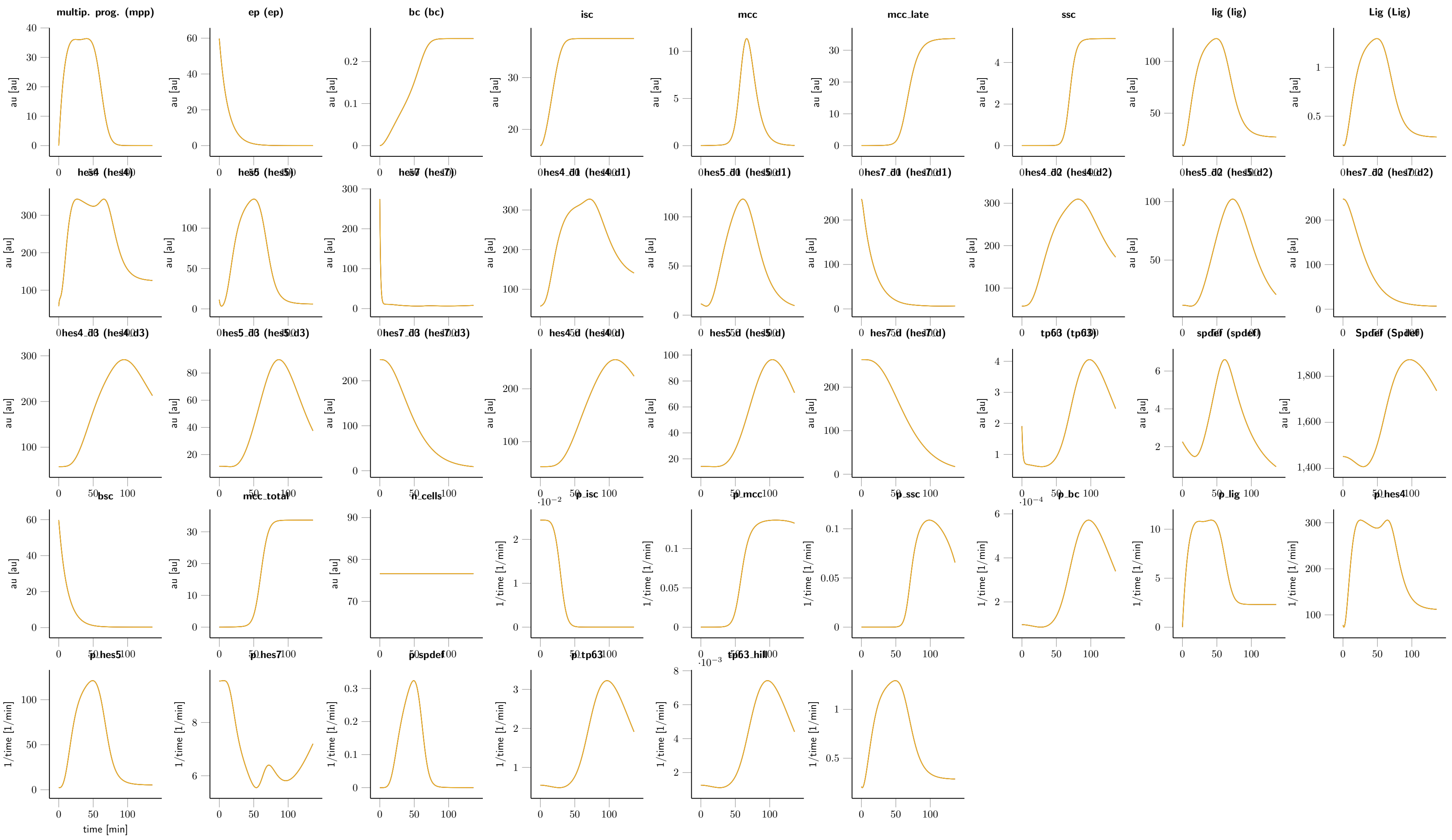

**Figure 8:** Fitted dynamics of hes7 loss of the function condition.

| time [min] | isc_abs [a.u.] | mcc_abs [a.u.] | ssc_abs [a.u.] |
|---|---|---|---|
| 124.03 | 122 | 73 | 22 |
| 124.03 | 99 | 60 | 7 |
| 124.03 | 103 | 105 | 6 |
| 124.03 | 79 | 97 | 4 |
| 124.03 | 98 | 51 | 3 |
| 124.03 | 104 | 118 | 3 |
| 124.03 | 119 | 86 | 11 |
| 124.03 | 80 | 74 | 8 |
| 124.03 | 88 | 69 | 18 |
| 124.03 | 125 | 87 | 3 |
| 124.03 | 116 | 123 | 25 |
| 124.03 | 78 | 113 | 48 |

**Table 10: Experimental data for the experiment LossOfFunction_Cells_hes7**

s_cell4 $\rightarrow$ s_cell4_fc $\cdot$ s_cell_mean
s_cell5 $\rightarrow$ s_cell5_fc $\cdot$ s_cell_mean
s_cell8 $\rightarrow$ s_cell8_fc $\cdot$ s_cell_mean
transl_hes4_lof_fc $\rightarrow$ 1
transl_hes5_lof_fc $\rightarrow$ 1

- **Local condition #40 (global condition #2):**

s_cell1 $\rightarrow$ s_cell1_fc $\cdot$ s_cell_mean
s_cell2 $\rightarrow$ s_cell2_fc $\cdot$ s_cell_mean
s_cell25 $\rightarrow$ s_cell25_fc $\cdot$ s_cell_mean
s_cell3 $\rightarrow$ s_cell3_fc $\cdot$ s_cell_mean
s_cell4 $\rightarrow$ s_cell4_fc $\cdot$ s_cell_mean
s_cell5 $\rightarrow$ s_cell5_fc $\cdot$ s_cell_mean
s_cell8 $\rightarrow$ s_cell8_fc $\cdot$ s_cell_mean
transl_hes4_lof_fc $\rightarrow$ 1
transl_hes5_lof_fc $\rightarrow$ 1

##### 1.15.4 Experimental data and model fit

#### 2 Estimated model parameters

In total 133 parameters are estimated from the experimental data. The model parameters were estimated by maximum likelihood estimation. The best fit yields a value of two time the negative log-likelihood that has been used as objective function $-2\log(L) = 3912.66$ for a total of 757 data points. In Table 11 – 13 the estimated parameter values are given. Parameters highlighted in red color indicate parameter values close to their bounds. The parameter name prefix init_ indicates the initial value of a dynamic variable.

#### References

[1] Alan C Hindmarsh, Peter N Brown, Keith E Grant, Steven L Lee, Radu Serban, Dan E Shumaker, and Carol S Woodward. SUNDIALS: Suite of nonlinear and differential/algebraic equation solvers. *ACM Transactions on Mathematical Software*, 31(3):363–396, sep 2005.

| | name | $\theta_{min}$ | $\hat{\theta}$ | $\theta_{max}$ | log | non-log $\hat{\theta}$ | estimated |
|---|---|---|---|---|---|---|---|
| 1 | d_hes4_prot_fc | -2 | +0.0649 | +2 | 1 | $+1.16 \cdot 1$ | 1 |
| 2 | d_hes4_fc | -2 | +0.0000 | +2 | 1 | $+1.00 \cdot 1$ | 0 |
| 3 | d_hes5_prot_fc | -2 | -0.0646 | +2 | 1 | $+8.62 \cdot 10^{-01}$ | 1 |
| 4 | d_hes5_fc | -2 | +0.0000 | +2 | 1 | $+1.00 \cdot 1$ | 0 |
| 5 | d_hes7_prot_fc | -2 | +0.0000 | +2 | 1 | $+1.00 \cdot 1$ | 1 |
| 6 | d_hes7_fc | -2 | +0.0000 | +2 | 1 | $+1.00 \cdot 1$ | 0 |
| 7 | d_hes_prot | -4 | -1.2453 | +1 | 1 | $+5.68 \cdot 10^{-02}$ | 1 |
| 8 | d_hes_rna | -4 | -0.0505 | +1 | 1 | $+8.90 \cdot 10^{-01}$ | 1 |
| 9 | d_lig_prot | -4 | +1.0000 | +1 | 1 | $+1.00 \cdot 10^{+01}$ | 1 |
| 10 | d_lig | -4 | -1.0622 | +1 | 1 | $+8.66 \cdot 10^{-02}$ | 1 |
| 11 | d_Spdef | -4 | -2.3068 | +1 | 1 | $+4.93 \cdot 10^{-03}$ | 1 |
| 12 | d_spdef | -4 | -1.5383 | +1 | 1 | $+2.90 \cdot 10^{-02}$ | 1 |
| 13 | d_tp63 | -4 | -0.0999 | +1 | 1 | $+7.95 \cdot 10^{-01}$ | 1 |
| 14 | hes_delay | -1 | +1.8196 | +4 | 1 | $+6.60 \cdot 10^{+01}$ | 1 |
| 15 | h_cell_i | +0 | +0.8451 | +0.8 | 1 | $+7.00 \cdot 1$ | 1 |
| 16 | h_hes4_i | +0 | +0.6710 | +0.8 | 1 | $+4.69 \cdot 1$ | 1 |
| 17 | h_hes4_a | +0 | +0.1324 | +0.8 | 1 | $+1.36 \cdot 1$ | 1 |
| 18 | h_hes5_a | +0 | +0.3560 | +0.8 | 1 | $+2.27 \cdot 1$ | 1 |
| 19 | h_hes7_i | +0 | +0.6620 | +0.8 | 1 | $+4.59 \cdot 1$ | 1 |
| 20 | h_hes7_i_spdef | +0 | +0.8451 | +0.8 | 1 | $+7.00 \cdot 1$ | 1 |
| 21 | h_isc_i_hes4 | +0 | +0.7877 | +0.8 | 1 | $+6.13 \cdot 1$ | 1 |
| 22 | h_isc_i_hes5 | +0 | +0.3142 | +0.8 | 1 | $+2.06 \cdot 1$ | 1 |
| 23 | h_isc_a | +0 | +0.8319 | +0.8 | 1 | $+6.79 \cdot 1$ | 1 |
| 24 | h_mcc_i_hes5 | +0 | +0.5720 | +0.8 | 1 | $+3.73 \cdot 1$ | 1 |
| 25 | h_mcc_a | +0 | +0.8451 | +0.8 | 1 | $+7.00 \cdot 1$ | 1 |
| 26 | h_spdef_a | +0 | +0.6817 | +0.8 | 1 | $+4.80 \cdot 1$ | 1 |
| 27 | h_ssc_i_hes4 | +0 | +0.7715 | +0.8 | 1 | $+5.91 \cdot 1$ | 1 |
| 28 | h_ssc_i_spdef | +0 | +0.7765 | +0.8 | 1 | $+5.98 \cdot 1$ | 1 |
| 29 | h_ssc_a | +0 | +0.8451 | +0.8 | 1 | $+7.00 \cdot 1$ | 1 |
| 30 | h_ssc_a_spdef | +0 | +0.8451 | +0.8 | 1 | $+7.00 \cdot 1$ | 1 |
| 31 | h_tp63_a | +0 | +0.3703 | +0.8 | 1 | $+2.35 \cdot 1$ | 1 |
| 32 | h_tp63_a_spdef | +0 | +0.8451 | +0.8 | 1 | $+7.00 \cdot 1$ | 1 |
| 33 | init_ep | +0 | +1.7764 | +4 | 1 | $+5.98 \cdot 10^{+01}$ | 1 |
| 34 | init_foxa1 | -0.5 | +0.2766 | +1 | 1 | $+1.89 \cdot 1$ | 1 |
| 35 | init_hes4 | +1 | +1.7596 | +2 | 1 | $+5.75 \cdot 10^{+01}$ | 1 |
| 36 | init_hes4_gof_lc_fc | +0 | +0.1246 | +3 | 1 | $+1.33 \cdot 1$ | 1 |
| 37 | init_hes5 | +0 | +1.0578 | +2 | 1 | $+1.14 \cdot 10^{+01}$ | 1 |
| 38 | init_hes5_gof_lc_fc | +0 | +0.1755 | +3 | 1 | $+1.50 \cdot 1$ | 1 |
| 39 | init_hes7 | +2 | +2.4384 | +3 | 1 | $+2.74 \cdot 10^{+02}$ | 1 |
| 40 | init_hes7_gof_lc_fc | +0 | +0.0001 | +3 | 1 | $+1.00 \cdot 1$ | 1 |
| 41 | init_isc | -4 | +0.0246 | +2 | 1 | $+1.06 \cdot 1$ | 1 |
| 42 | init_isc_gof_lc_hes4_fc | -3 | -0.4959 | +3 | 1 | $+3.19 \cdot 10^{-01}$ | 1 |
| 43 | init_isc_gof_lc_hes5_fc | -3 | -0.1626 | +3 | 1 | $+6.88 \cdot 10^{-01}$ | 1 |
| 44 | init_isc_gof_lc_hes7_fc | -3 | -1.3223 | +3 | 1 | $+4.76 \cdot 10^{-02}$ | 1 |
| 45 | init_isc_lof_hes4_fc | -3 | -0.7765 | +3 | 1 | $+1.67 \cdot 10^{-01}$ | 1 |
| 46 | init_isc_lof_hes5_fc | -3 | -0.3727 | +3 | 1 | $+4.24 \cdot 10^{-01}$ | 1 |
| 47 | init_isc_lof_hes7_fc | -3 | +1.2023 | +3 | 1 | $+1.59 \cdot 10^{+01}$ | 1 |
| 48 | init_ligand | +0.8 | +1.2937 | +2 | 1 | $+1.97 \cdot 10^{+01}$ | 1 |
| 49 | init_mcc | -4 | -3.8083 | +2 | 1 | $+1.55 \cdot 10^{-04}$ | 1 |

**Table 11: Estimated parameter values**
$\hat{\theta}$ indicates the estimated value of the parameters. $\theta_{min}$ and $\theta_{max}$ indicate the upper and lower bounds for the parameters. The log-column indicates if the value of a parameter was log-transformed. If log $\equiv$ 1 the non-log-column indicates the non-logarithmic value of the estimate. The estimated-column indicates if the parameter value was estimated (1), was temporarily fixed (0) or if its value was fixed to a constant value (2).

| | name | $\theta_{min}$ | $\hat{\theta}$ | $\theta_{max}$ | log | non-log $\hat{\theta}$ | estimated |
|---|---|---|---|---|---|---|---|
| 50 | init_mcc_gof_lc_hes4_fc | -3 | -0.0004 | +3 | 1 | $+9.99 \cdot 10^{-01}$ | 1 |
| 51 | init_mcc_gof_lc_hes5_fc | -3 | +0.0011 | +3 | 1 | $+1.00 \cdot 1$ | 1 |
| 52 | init_mcc_gof_lc_hes7_fc | -3 | -0.0007 | +3 | 1 | $+9.98 \cdot 10^{-01}$ | 1 |
| 53 | init_mcc_lof_hes4_fc | -3 | -0.0009 | +3 | 1 | $+9.98 \cdot 10^{-01}$ | 1 |
| 54 | init_mcc_lof_hes5_fc | -3 | -0.0001 | +3 | 1 | $+1.00 \cdot 1$ | 1 |
| 55 | init_mcc_lof_hes7_fc | -3 | -0.0003 | +3 | 1 | $+9.99 \cdot 10^{-01}$ | 1 |
| 56 | init_mcidas | -5 | +0.3867 | +1 | 1 | $+2.44 \cdot 1$ | 1 |
| 57 | init_mpp_gof_lc_hes4_fc | -3 | -0.0000 | +3 | 1 | $+1.00 \cdot 1$ | 1 |
| 58 | init_mpp_gof_lc_hes5_fc | -3 | +0.0000 | +3 | 1 | $+1.00 \cdot 1$ | 1 |
| 59 | init_mpp_gof_lc_hes7_fc | -3 | -0.0006 | +3 | 1 | $+9.99 \cdot 10^{-01}$ | 1 |
| 60 | init_mpp_lof_hes4_fc | -3 | -0.0007 | +3 | 1 | $+9.98 \cdot 10^{-01}$ | 1 |
| 61 | init_mpp_lof_hes5_fc | -3 | -0.0001 | +3 | 1 | $+1.00 \cdot 1$ | 1 |
| 62 | init_mpp_lof_hes7_fc | -3 | -0.0001 | +3 | 1 | $+1.00 \cdot 1$ | 1 |
| 63 | init_mpp | -4 | -3.9859 | +2 | 1 | $+1.03 \cdot 10^{-04}$ | 1 |
| 64 | init_spdef | -1 | +0.3542 | +0.5 | 1 | $+2.26 \cdot 1$ | 1 |
| 65 | init_tp63 | -1 | +0.2810 | +0.5 | 1 | $+1.91 \cdot 1$ | 1 |
| 66 | init_ubp1 | -5 | -4.9964 | +1 | 1 | $+1.01 \cdot 10^{-05}$ | 1 |
| 67 | k_cell_i_fc | +0 | +2.0425 | +3 | 1 | $+1.10 \cdot 10^{+02}$ | 1 |
| 68 | k_hes4_i_fc | +0 | +0.2828 | +2 | 1 | $+1.92 \cdot 1$ | 1 |
| 69 | k_hes4_a_fc | +0 | +0.6112 | +2 | 1 | $+4.09 \cdot 1$ | 1 |
| 70 | k_hes5_a_fc | +0 | +0.9823 | +3 | 1 | $+9.60 \cdot 1$ | 1 |
| 71 | foldraus | +0 | +0.8350 | +3 | 1 | $+6.84 \cdot 1$ | 1 |
| 72 | k_hes7_i_spdef_fc | +0 | +0.1182 | +3 | 1 | $+1.31 \cdot 1$ | 1 |
| 73 | k_isc_i_hes4_fc | +0 | +0.0000 | +3 | 1 | $+1.00 \cdot 1$ | 1 |
| 74 | k_isc_i_hes5_fc | +0 | +0.0000 | +3 | 1 | $+1.00 \cdot 1$ | 1 |
| 75 | k_isc_a_fc | -3 | -2.9620 | +0 | 1 | $+1.09 \cdot 10^{-03}$ | 1 |
| 76 | k_mcc_i_hes5_fc | +0 | +2.2420 | +3 | 1 | $+1.75 \cdot 10^{+02}$ | 1 |
| 77 | k_mcc_a_fc | +0 | +0.4421 | +3 | 1 | $+2.77 \cdot 1$ | 1 |
| 78 | k_spdef_a_fc | +0 | +1.2134 | +3 | 1 | $+1.63 \cdot 10^{+01}$ | 1 |
| 79 | k_ssc_i_hes4_fc | +0 | +2.9978 | +3 | 1 | $+9.95 \cdot 10^{+02}$ | 1 |
| 80 | k_ssc_i_spdef_fc | +0 | +1.7931 | +2 | 1 | $+6.21 \cdot 10^{+01}$ | 1 |
| 81 | k_ssc_a_fc | +0 | +0.6157 | +3 | 1 | $+4.13 \cdot 1$ | 1 |
| 82 | k_ssc_a_spdef_fc | +0 | +0.1182 | +2 | 1 | $+1.31 \cdot 1$ | 1 |
| 83 | k_tp63_a_fc | +0 | +2.4458 | +3 | 1 | $+2.79 \cdot 10^{+02}$ | 1 |
| 84 | k_tp63_a_spdef_fc | +0 | +0.4145 | +3 | 1 | $+2.60 \cdot 1$ | 1 |
| 85 | prior_p_hes4 | +0 | +1.0000 | +2 | 1 | $+1.00 \cdot 10^{+01}$ | 0 |
| 86 | prior_p_hes5 | +0 | +1.0000 | +2 | 1 | $+1.00 \cdot 10^{+01}$ | 0 |
| 87 | prior_p_hes7 | +0 | +1.0000 | +2 | 1 | $+1.00 \cdot 10^{+01}$ | 0 |
| 88 | prior_p_lig_mpp | +0 | +1.0000 | +2 | 1 | $+1.00 \cdot 10^{+01}$ | 0 |
| 89 | prior_p_spdef | +0 | +1.0000 | +2 | 1 | $+1.00 \cdot 10^{+01}$ | 0 |
| 90 | prior_p_tp63 | +0 | +1.0000 | +2 | 1 | $+1.00 \cdot 10^{+01}$ | 0 |
| 91 | p_lig_ep_fc | -4 | -3.3088 | +0 | 1 | $+4.91 \cdot 10^{-04}$ | 1 |
| 92 | p_lig_bc_fc | -4 | -2.9078 | +0 | 1 | $+1.24 \cdot 10^{-03}$ | 1 |
| 93 | p_lig_isc_fc | -4 | -3.7450 | +0 | 1 | $+1.80 \cdot 10^{-04}$ | 1 |
| 94 | p_lig_mcc_fc | -4 | -0.6378 | +0 | 1 | $+2.30 \cdot 10^{-01}$ | 1 |
| 95 | p_lig_ssc_fc | -4 | -3.6395 | +0 | 1 | $+2.29 \cdot 10^{-04}$ | 1 |
| 96 | p_mcc_late | -3 | -0.9863 | +1 | 1 | $+1.03 \cdot 10^{-01}$ | 1 |
| 97 | p_mpp | -3 | -1.0896 | +1 | 1 | $+8.14 \cdot 10^{-02}$ | 1 |
| 98 | p_Spdef | -4 | +0.5008 | +2 | 1 | $+3.17 \cdot 1$ | 1 |
| 99 | p_Lig | -4 | -0.9740 | +2 | 1 | $+1.06 \cdot 10^{-01}$ | 1 |

**Table 12: Estimated parameter values**
$\hat{\theta}$ indicates the estimated value of the parameters. $\theta_{min}$ and $\theta_{max}$ indicate the upper and lower bounds for the parameters. The log-column indicates if the value of a parameter was log-transformed. If log $\equiv$ 1 the non-log-column indicates the non-logarithmic value of the estimate. The estimated-column indicates if the parameter value was estimated (1), was temporarily fixed (0) or if its value was fixed to a constant value (2).

| | name | $\theta_{min}$ | $\hat{\theta}$ | $\theta_{max}$ | log | non-log $\hat{\theta}$ | estimated |
|---|---|---|---|---|---|---|---|
| 100 | reg_hes4_fc | -2 | -0.0000 | +2 | 1 | $+1.00 \cdot 1$ | 1 |
| 101 | reg_hes5_fc | -2 | +0.0000 | +2 | 1 | $+1.00 \cdot 1$ | 1 |
| 102 | reg_hes7_fc | -2 | +0.0000 | +2 | 1 | $+1.00 \cdot 1$ | 1 |
| 103 | reg_lig_fc | -2 | -0.0000 | +2 | 1 | $+1.00 \cdot 1$ | 1 |
| 104 | reg_p_hes4_fc | -2 | +1.7804 | +2 | 1 | $+6.03 \cdot 10^{+01}$ | 1 |
| 105 | reg_p_hes5_fc | -2 | +1.6515 | +2 | 1 | $+4.48 \cdot 10^{+01}$ | 1 |
| 106 | reg_p_hes7_fc | -2 | +0.0407 | +2 | 1 | $+1.10 \cdot 1$ | 1 |
| 107 | reg_p_lig_mpp_fc | -2 | +0.3599 | +2 | 1 | $+2.29 \cdot 1$ | 1 |
| 108 | reg_p_spdef_fc | -2 | +0.5374 | +2 | 1 | $+3.45 \cdot 1$ | 1 |
| 109 | reg_p_tp63_fc | -2 | +1.6381 | +2 | 1 | $+4.35 \cdot 10^{+01}$ | 1 |
| 110 | reg_spdef_fc | -2 | -0.0000 | +2 | 1 | $+1.00 \cdot 1$ | 1 |
| 111 | reg_tp63_fc | -2 | +0.0000 | +2 | 1 | $+1.00 \cdot 1$ | 1 |
| 112 | s_RNA | -2 | +0.0000 | +2 | 1 | $+1.00 \cdot 1$ | 0 |
| 113 | s_cell12_fc | -2 | -0.0825 | +2 | 1 | $+8.27 \cdot 10^{-01}$ | 1 |
| 114 | s_cell13_fc | -2 | -0.1107 | +2 | 1 | $+7.75 \cdot 10^{-01}$ | 1 |
| 115 | s_cell14_fc | -2 | -0.1948 | +2 | 1 | $+6.39 \cdot 10^{-01}$ | 1 |
| 116 | s_cell1_fc | -2 | +0.0525 | +2 | 1 | $+1.13 \cdot 1$ | 1 |
| 117 | s_cell20_fc | -2 | +0.0598 | +2 | 1 | $+1.15 \cdot 1$ | 1 |
| 118 | s_cell22_fc | -2 | +0.1135 | +2 | 1 | $+1.30 \cdot 1$ | 1 |
| 119 | s_cell23_fc | -2 | +0.0802 | +2 | 1 | $+1.20 \cdot 1$ | 1 |
| 120 | s_cell24_fc | -2 | +0.0651 | +2 | 1 | $+1.16 \cdot 1$ | 1 |
| 121 | s_cell25_fc | -2 | +0.0394 | +2 | 1 | $+1.09 \cdot 1$ | 1 |
| 122 | s_cell2_fc | -2 | +0.0925 | +2 | 1 | $+1.24 \cdot 1$ | 1 |
| 123 | s_cell3_fc | -2 | +0.0379 | +2 | 1 | $+1.09 \cdot 1$ | 1 |
| 124 | s_cell4_fc | -2 | +0.1298 | +2 | 1 | $+1.35 \cdot 1$ | 1 |
| 125 | s_cell5_fc | -2 | -0.0083 | +2 | 1 | $+9.81 \cdot 10^{-01}$ | 1 |
| 126 | s_cell8_fc | -2 | +0.0133 | +2 | 1 | $+1.03 \cdot 1$ | 1 |
| 127 | s_cell9_fc | -2 | +0.1215 | +2 | 1 | $+1.32 \cdot 1$ | 1 |
| 128 | s_cell_mean | -3 | +0.4092 | +3 | 1 | $+2.57 \cdot 1$ | 1 |
| 129 | s_isc | +0 | +2.8717 | +6 | 1 | $+7.44 \cdot 10^{+02}$ | 1 |
| 130 | s_isc_pure | +0 | +2.2581 | +6 | 1 | $+1.81 \cdot 10^{+02}$ | 1 |
| 131 | s_mcc | +0 | +3.8489 | +6 | 1 | $+7.06 \cdot 10^{+03}$ | 1 |
| 132 | s_mpp_foxi1_fc | -3 | -2.3341 | +3 | 1 | $+4.63 \cdot 10^{-03}$ | 1 |
| 133 | s_ssc | +0 | +1.8180 | +6 | 1 | $+6.58 \cdot 10^{+01}$ | 1 |
| 134 | sd_RNA | -3 | -0.4722 | +0 | 1 | $+3.37 \cdot 10^{-01}$ | 1 |
| 135 | sd_cell | +0 | +1.2720 | +2 | 1 | $+1.87 \cdot 10^{+01}$ | 1 |
| 136 | transl_hes4_lof_fc | -3 | -0.1430 | +0 | 1 | $+7.19 \cdot 10^{-01}$ | 1 |
| 137 | transl_hes5_lof_fc | -3 | -0.1517 | +0 | 1 | $+7.05 \cdot 10^{-01}$ | 1 |
| 138 | transl_hes7_lof_fc | -3 | -0.0456 | +0 | 1 | $+9.00 \cdot 10^{-01}$ | 1 |
| 139 | v_max_bc | -5 | -1.1132 | +2 | 1 | $+7.71 \cdot 10^{-02}$ | 1 |
| 140 | v_max_cell | -5 | -0.8297 | +2 | 1 | $+1.48 \cdot 10^{-01}$ | 1 |
| 141 | vmax_isc_fc | -2 | -0.1811 | +2 | 1 | $+6.59 \cdot 10^{-01}$ | 1 |
| 142 | vmax_mcc_fc | -2 | -0.0277 | +2 | 1 | $+9.38 \cdot 10^{-01}$ | 1 |
| 143 | vmax_ssc_fc | -2 | +0.2087 | +2 | 1 | $+1.62 \cdot 1$ | 1 |

**Table 13: Estimated parameter values**
$\hat{\theta}$ indicates the estimated value of the parameters. $\theta_{min}$ and $\theta_{max}$ indicate the upper and lower bounds for the parameters. The log-column indicates if the value of a parameter was log-transformed. If log $\equiv$ 1 the non-log-column indicates the non-logarithmic value of the estimate. The estimated-column indicates if the parameter value was estimated (1), was temporarily fixed (0) or if its value was fixed to a constant value (2).
